# A million orthopteran singers from a year of continuously operating field recorders detected, tracked and organized by song

**DOI:** 10.64898/2026.10.18.759993

**Authors:** Michael B. Eisen

## Abstract

Continuous acoustic monitoring has become an invaluable tool in ecology and conservation biology, with hundreds of thousands of automated recording units deployed worldwide. While semi-manual annotation is adequate for some uses, fully automated methods to segregate and identify the source of animal sounds are essential to utilize these data to their full extent. Significant progress has been made for birds, whose calls are generally discrete and structured, and where large annotated reference libraries are available. However these methods do not extend neatly to insects, whose choruses dominate nighttime, and often daytime, soundscapes across most of the world. While many approaches have been proposed to fill this gap, an efficient end-to-end system for annotating insect calls that can operate at the scale of available data has not emerged. Here we describe an unsupervised method that uses the intrinsic, highly repetitive structure of the songs of many insects, especially orthopterans, to identify and separate distinct songs in narrow (minute) time windows, connect them across windows for as long as entire nights as their carrier frequencies and syllable repeat rates vary with time, temperature and other factors, and assemble them across days and locations into a catalog of singers and song types across time, space and conditions. We apply this method to thirteen months (November 2024 to November 2025) of recordings from 27 autonomous recording units in different microhabitats at a field site in southwestern Arkansas, 175,896 recorder-hours in all. We detected 10.9 million singers in five-minute windows and linked them, within each night or day, into 911,005 tracks, of which 103,802 lasted two hours or more (71,770 at night, 32,032 by day), a total of 335,882 hours of song. In the plane of carrier frequency against syllable repeat rate these singers fall on discrete lines along which each slides with temperature, and we use this organization to define 45 song types, 37 trilling and 8 chirping, each composed of dozens to thousands of singer-nights. Within singers, syllable rate rises by about 3 percent per degree and carrier frequency by about 1.5 percent in nearly every type, recapitulating and extending classical observations on the temperature dependence of orthopteran song. Published and reference recordings placed on the same plane fall on these lines, and species of *Anaxipha* recorded by Walker and Funk land on single types at the right temperatures. More generally, this method and the data it produces provide a robust means of studying changes in insect song over time and space, and of understanding the behavioral and physical forces that shape its production.

## Introduction

The commercial availability of audio recorders designed to operate for long periods of time in field conditions (generally referred to as autonomous recording units, or ARUs), and their expanding use by researchers and amateur naturalists has inspired the development of tools designed to automatically annotate these recordings. The bulk of the effort has been directed at birds, and while methodological gaps remain, tools such as BirdNET (Kahl et al. 2021) and Perch (Ghani et al. 2023) allow for the algorithmic identification of most birds, along with some amphibians, mammals and insects (Stowell 2022).

However, in much of the world, a large fraction of the zoogenic energy in such recordings, especially at night, comes from Orthoptera – crickets, katydids and their relatives – and cicadas, who produce widely recognized songs consisting of rhythmic, repetitive sound pulses, grouped into chirps or extended into elongated trills or buzzes. Yet the species that produce these songs are largely absent from the inventories that acoustic monitoring produces.

Acoustic indices (Sueur et al. 2008; Pieretti et al. 2011; Sueur et al. 2014), the workhorse tool of soundscape ecology, capture some of this richness (Aide et al. 2017), and have shown promise as a tool for monitoring biodiversity recovery (Müller et al. 2023), a primary goal of ours. However, while they have the virtue of being easy to calculate, they lack taxonomic specificity, and the recent experience with avian acoustics clearly demonstrates that species and individual identification, and localization (Eisen 2026) should be the goal for insects.

There are several reasons why this does not yet exist. Deep-learning models like BirdNET and Perch, which are trained to recognize animal sounds from their characteristic patterns across time and frequency, generalize imperfectly to the continuous nature of orthopteran songs. There is also a far less robust collection of reference songs on which to draw. And, unlike birds and frogs, insect calls vary in syllable repeat rate and in some cases carrier frequency with temperature, rendering static acoustic signatures imprecise and unreliable.

Here, rather than attempting to associate audio signals with species up front, we describe a method that uses the shared characteristics of orthopteran songs to extract them from short (minutes) time windows, and to trace them across time for as long as entire nights or days. We then use the observed carrier frequencies, syllable repeat rates and temperature response to partition individual singers into candidate taxonomic units, and, where available, use reference data to suggest likely species or species complex identifications.

We were motivated to develop this method because we have generated over 400,000 hours of recordings from over three years of continuous operation of ARUs dispersed across a field site in southwestern Arkansas, where orthopteran choruses are complex and cacophonous from early spring to late fall. Although the data were collected to identify and localize birds, we lamented the lack of tools available to deal with the non-avian components of this soundscape.

Before describing the details of the approach we have taken, and the results of its application to our data, we first delve into the rich history of orthopteran acoustic research on which this work depends.

### History of Orthoptera song analysis

Humans have always been captivated by orthopteran singing. According to a myth recounted by Socrates in Plato’s Phaedrus, cicadas were once humans who became so enraptured when song first appeared that they forgot to eat and drink, and died. The Muses transformed them into cicadas, granting them the ability to sing throughout their lives without food or drink; after death, they return to the Muses and report which humans have honored each of them (Phaedrus 259b–d).

The description of insect song as a measurable signal traces at least as far back to the work of Alexandre Yersin in Switzerland in the 1850s (Yersin 1854), and subsequently Samuel Scudder, a noted American insect paleontologist, who in 1868 transcribed the songs of several New England orthopterans in musical notation (Scudder 1868a; Scudder 1868b). In 1881, Margarette Brooks, a young entomologist on Edward Morse’s staff at the Peabody Academy in Salem, prompted by a newspaper report that crickets could serve as thermometers, measured the relationship between chirp rate and temperature and published it in Popular Science Monthly. Then, in 1897 Amos Dolbear provided a simple formula for this relationship T(°F) = 50 + (N₆₀ − 40)/4, with N₆₀ the chirps per minute based on observations he made of (what is now believed to have been) the snowy tree cricket, *Oecanthus fultoni* (Dolbear 1897).

The publication of “Dolbear’s Law” inspired additional observations, which refined the formula and its accuracy to around one degree (Bessey and Bessey 1898; Edes 1899). Subsequent investigations (Shull 1907; Fulton 1925; Allard 1928; Allard 1930a; Allard 1930b) showed that there was considerable individual variation, and traced it to other environmental factors (such as humidity and wind currents) as well as to “cricket individuality”. Hayward (1901) extended the relation to a katydid, the true katydid *Pterophylla camellifolia*, whose phrase rate is several times more temperature-sensitive than a cricket’s. A relationship between temperature and pitch was also noted for many species (Fulton 1915).

The introduction of inexpensive oscilloscopes following the conclusion of WWII revolutionized the field. George Washington Pierce, a Harvard physicist known for work on electronic communication, spent his retirement recording crickets and katydids and published a lengthy book The Songs of Insects (Pierce 1948), the first quantitative account of carrier frequencies, pulse structure and their dependence on temperature across many species. His temperature work was extended to coneheads and meadow katydids by Frings and Frings (1957, 1962). Alexander used song as a taxonomic character, showing that the eastern Gryllus comprised several species distinguishable by song where morphology failed (Alexander 1957c; Alexander 1962) and doing the same for the ground crickets (Alexander 1957b).

In the early 1960s, Thomas Walker began to systematically characterize the carrier frequencies, syllable repeat rates, and temperature response of songs of US orthopterans, and used these data to revise the taxonomies of multiple groups (Walker 1962a; Walker 1962b; Walker 1963). In this work he established that in tree crickets both carrier frequency and pulse rate rise with temperature, and at a fixed ratio, while in field crickets the carrier is nearly temperature-independent. These data were subsequently compiled per species, along with the recordings that made them, in the online database Singing Insects of North America (Walker & Moore, SINA), and remain the reference data for the continent upon which this work draws heavily.

These field studies on the effects of temperature have been complemented by multiple studies in controlled conditions, including, for example, in *Gryllus bimaculatus* (Doherty 1985), *Gryllus firmus* (Pires and Hoy 1992), and *Gryllus intege*r (Martin, Gray and Cade 2000). Another result of Walker, that female tree crickets’ preference for pulse rate shifts with temperature in step with the males’ song (Walker 1957), has also been extensively studied subsequently, and largely confirmed (Doherty 1985; Pires and Hoy 1992; Gerhardt and Huber 2002), with some caveats about its efficacy in multispecies assemblages (Symes et al. 2017).

The mechanism was worked out in parallel. Orthopteran song is the product of a neural rhythm and a mechanical oscillator. Crickets and katydids sing by drawing a scraper on one forewing across a toothed file on the other (crickets on the closing stroke of each wing cycle), while slant-faced grasshoppers (Gomphocerinae) rub a row of pegs on the hind femur against a forewing vein (Alexander 1957a; Elsner & Popov 1978). Within a single stroke, each tooth impact delivers an impulse to a resonant region of the wing (the harp in crickets, the mirror in katydids) whose vibration is the carrier: its frequency is set jointly by the rate of tooth impacts and the resonator’s tuning, which in field crickets are locked together by an escapement so that one tooth is released per carrier cycle, the “clockwork cricket” (Elliott & Koch 1985; Koch et al. 1988; Bennet-Clark 1999; Bennet-Clark & Bailey 2002; Montealegre-Z et al. 2011; Montealegre-Z & Mason 2005 for katydids). The strokes themselves are driven by a central pattern generator in the thoracic and abdominal ganglia (Kutsch & Huber 1989; Hedwig 2000; Schöneich & Hedwig 2011, 2012), whose rhythm sets the repetition rate of the sound pulses (the syllables) and groups them into the species-specific patterns of chirps and trills (Alexander 1962). Neural and muscular activity thus governs the timing of song, while tooth-impact dynamics and wing resonance govern the sound wave within each syllable and hence its carrier frequency.

This separation helps explain why temperature can affect song timing and pitch differently. Temperature-sensitive neural and muscular processes generally accelerate song rhythms with warming (Walker 1975); selective-warming experiments implicate thoracic neural circuitry in this response, although additional body regions can contribute (Pires & Hoy 1992). Carrier frequency can be less temperature-sensitive because mechanical resonance constrains the oscillations within each pulse. In field crickets, an escapement-like interaction between the scraper, file, and vibrating wings couples tooth release to wing oscillation, helping maintain a relatively stable pitch despite changes in song tempo (Bennet-Clark & Bailey 2002). This constraint varies among taxa: tree crickets possess multiple nearby wing resonances that permit temperature-dependent changes in the driving mechanism to shift carrier frequency (Mhatre et al. 2012). Thermal variation in orthopteran songs thus reflects both the temperature sensitivity of motor-pattern generation and the mechanical constraints of the sound-producing apparatus.

### Automated acoustic monitoring of Orthopterans

Riede proposed in 1993 that acoustic inventories of orthopteran songs could be important tools in ecology and conservation biology, and demonstrated that plots of carrier frequency by syllable repeat rate had the information needed to discriminate individual species (Riede 1993; Riede 1998; Riede 2018).

Ganchev and Potamitis further established the discriminative value of this information on reference libraries of cricket, katydid, and cicada recordings (Potamitis et al. 2006; Ganchev and Potamitis 2007). But the most complete demonstration of what the simultaneous analysis of carrier frequency and syllable repeat rate can tell us came from Walker and Funk’s (Walker & Funk 2014) revision of the North American trigs of the genus *Anaxipha*. Working from 792 recordings of six sympatric, continuously trilling species (single males, recorded by hand with a thermometer in the vegetation beside them), they showed that when carrier frequency is plotted against pulse rate each species forms its own tilted line: a singer moves up and along the line at higher temperatures, and the lines of different species do not cross, so that a song can be assigned to species “with no need that the temperature of the calling male be known.”

The same paper assembled the temperature coefficients of pulse rate and carrier frequency across cricket taxa and made explicit the mechanistic reason the lines have the slopes they do: in *Gryllus*, whose wing resonator releases one file tooth per cycle, carrier frequency barely moves with temperature, whereas in tree crickets, trigs, and ground crickets the file drives the wing at the tooth-strike rate and carrier rises with pulse rate.

The work of Riede, Ganchev and Potamitis, and Walker and Funk shows both the value of these measurements and the feasibility of making them automatically, at scale, from unattended soundscape recordings. Riede went further and proposed a general “cricket detector” that would sort singers into “ethospecies” by their songs even where no reference recordings exist. There have since been signal-processing detectors for individual song types (Brandes 2008), automated monitoring of particular target taxa (Penone et al. 2013; Jeliazkov et al. 2016; Newson et al. 2017), deep-learning classifiers trained on curated insect libraries (Faiß and Stowell 2023; Isupova et al. 2026), and BirdNET now lists a few orthopterans among its species, though in our experience those detectors do not perform robustly in the field. To our knowledge, however, Riede’s idea has never been implemented or applied at scale.

We aim to do that here. We have developed a fully automated method, BROOKS (“Bioacoustic Records of Orthopteran Oscillators, Keyed by Song”), named for Margarette Brooks, who in 1881 first timed a cricket’s song against a thermometer, that identifies signals with the structure of orthopteran song in five-minute windows of continuous recording. From each singer’s carrier frequency, syllable rate and pattern of singing and silence we separate what we believe are individual singers and follow them for hours. The species-specific couplings that emerge between carrier frequency, syllable rate and temperature then define, and where possible let us name, distinct taxonomic units, Riede’s ethospecies.

### The structure of orthopteran calls

Figure 1 illustrates the basic architecture of the orthopteran songs we analyze here. Although there is wide variation in orthopteran songs, many of their calls have some common features that we rely on in our analysis. The basic unit is a <u>syllable (</u>Figure 1A), composed of a short millisecond burst in a narrow frequency band (the <u>carrier frequency</u>), that often sweeps across adjacent frequencies within the syllable. Carrier frequencies can range from around 600 Hz to above 100 kHz, with the main orthopteran groups (crickets, katydids and grasshoppers) tending to occupy different parts of carrier frequency space.

**Figure 1.**
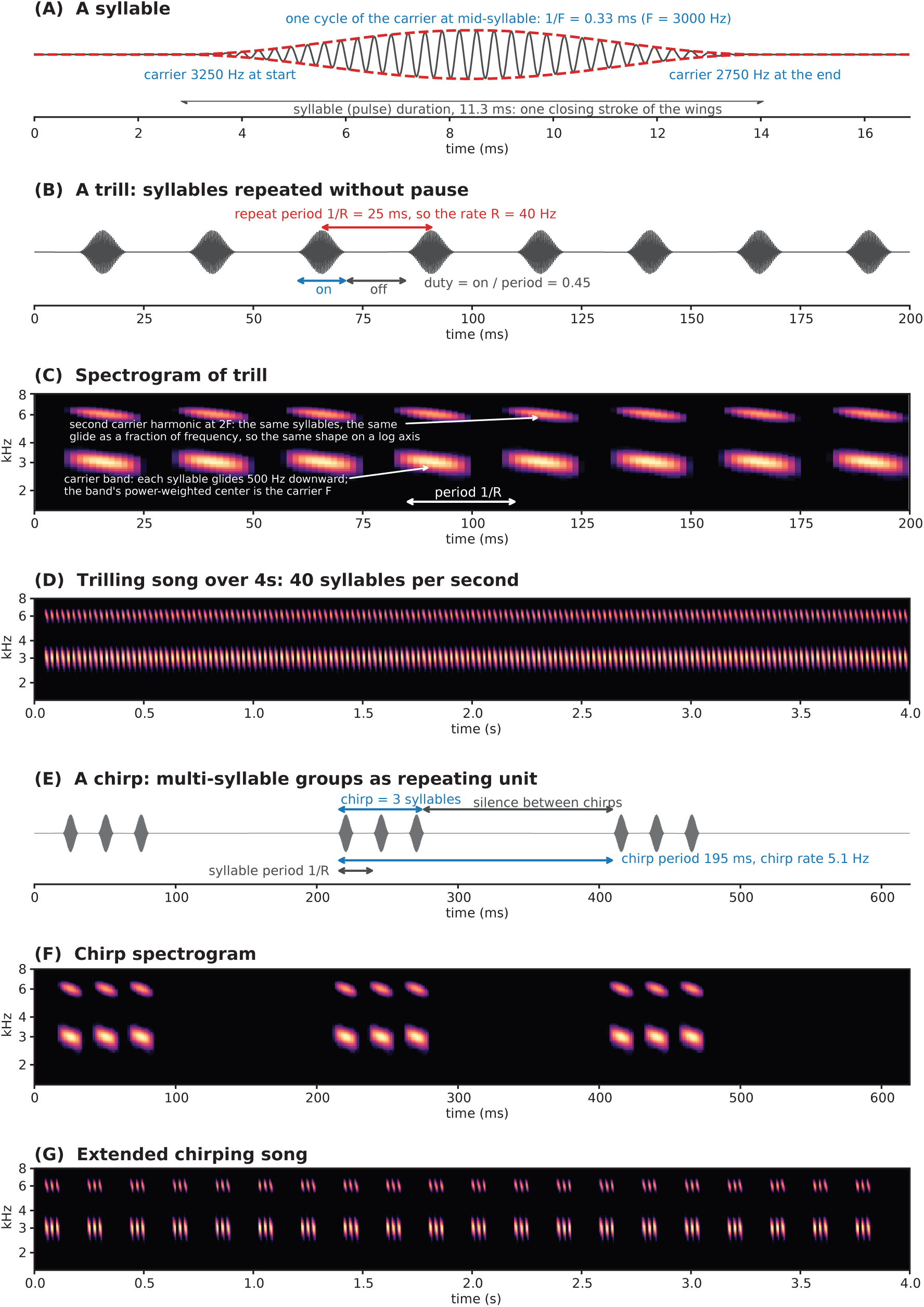
Anatomy of orthopteran songs Basic features of typical orthopteran songs including continuous trills with regularly repeating syllables in extended bouts and chirps with short bursts of repeated syllables are themselves repeated at a regular interval.

Syllables are repeated at a regular rate (the <u>syllable repeat rate</u>), and are organized into two main types of songs. <u>Trills</u> (Figure 1B-D) are long (usually multi-second to considerably longer) runs of syllables repeated at a regular rate. <u>Chirps</u> (Figure 1E-G) consist of groups of syllables strung together usually in a stereotyped way, and which are then themselves repeated in bursts, usually with regular spacing between chirps. (It was the inter-chirp interval that early orthopteran acoustic chronometrists like Brooks and Dolbear were using as their temperature-dependent timer, as this can be counted by ear, whereas the syllable repeat rate generally can only be studied electronically).

By and large, amongst North American orthopterans, the chirpers vs. trillers distinction follows taxonomy. Field crickets (*Gryllus*) are mostly chirpers, although some like *G. rubens* and *G. texensis* are trillers (Weissman and Gray 2019). In tree crickets (*Oecanthus*), the niveus group chirps, while the nigricornis group trills nearly continuously (Walker 1962a; Walker 1963). Ground crickets (Nemobiinae) are mostly trillers. Trigs (Trigonidiinae) are mostly trillers, with considerable variation in trill duration (Walker and Funk 2014). Mole crickets (Gryllotalpidae) are trillers. Katydids (Tettigoniidae) have more complex syllables, and their songs vary by taxa into a mix of chirp-like and trill-like. The primarily diurnal grasshoppers (Acrididae) tend to be more raspy, and are not a main focus of this analysis.

### Audio Data

The data we analyze here were collected using 27 <u>Solar BAR ARUs</u> (Frontier Labs, Brisbane, Australia) recording at 44,100 Hz between November 1, 2024 and November 30, 2025. The recorders were distributed across a 400 ha field site in southwestern Arkansas (USA) (see Figure 2A). This site was primarily used as pasture until 2022, and is now being used by us to conduct experiments on restoring native forests and the impact of doing so on biodiversity. The recorders were positioned at 11 stations: Back9 (7 recorders), NorthSide (5), EastCreek (4), House (4), and single recorders at Copse, EastMeander, FarNorth, MidCreek, Pond, SecretGarden and WestSideJunction. Four stations have multiple recorders to facilitate bird localization (Eisen 2026). The recorders were set to collect around the clock. Uptime was uneven (Figure 2B), so every figure that compares months or recorders carries the fraction of hours recorded in its panel title.

**Figure 2.**
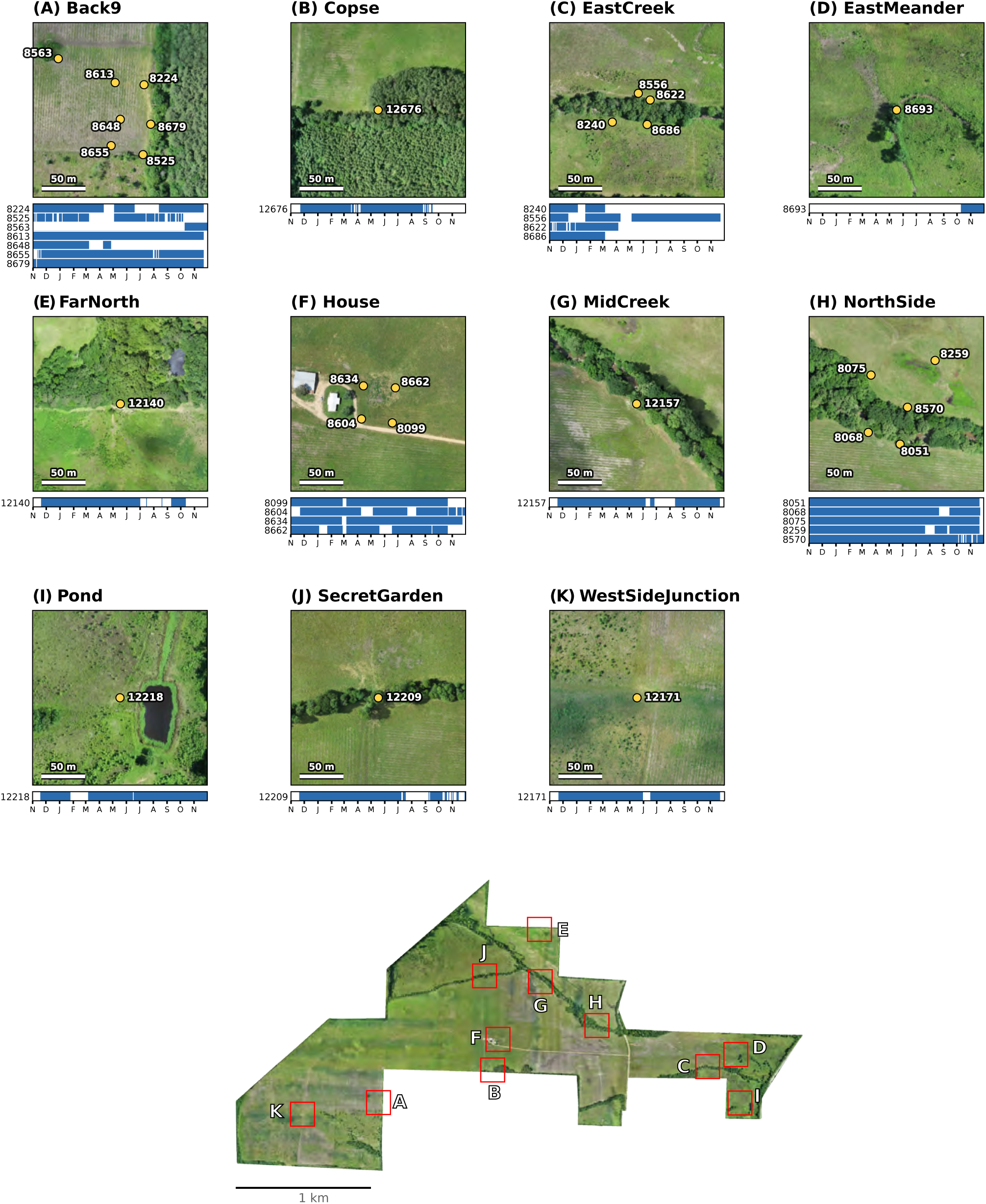
Recorder locations and uptime All recordings analyzed in this paper were made on Solar BAR (Frontier Labs) autonomous recording units distributed around a 400ha field site in the piney woods and rolling agricultural landscapes of the West Gulf Coastal Plain, in the northern part of the Ark-La-Tex region of the American South.

A representative night’s worth of data is shown in Figure 3, showing data from the night of September 20-21, 2025. Several prominent callers can be seen (Figure 3A) from dusk to dawn in the 6-8 kHz carrier frequency range, joined by additional callers in the 2-4 kHz carrier frequency range around sunset at 7:15 PM. The shift in carrier frequency over the night is shown in Figure 3B, along with hourly temperature estimates. The carrier frequencies are mostly stable in the one hour window shown in Figure 3C, allowing for a large number of candidate singers to be seen by eye. The bout pattern of individual callers can be seen better in Figure 3D, which covers 5 minutes, the window size used below to identify individual singers. Windows of 3s (Figure 3E-F) show the trilling pattern of a singer at a carrier frequency of 3.0 to 3.5 kHz (the spectrogram shown in Figure 3F is filtered to highlight the trill frequency). Figure 3G shows the shape of a single 34 ms syllable.

**Figure 3.**
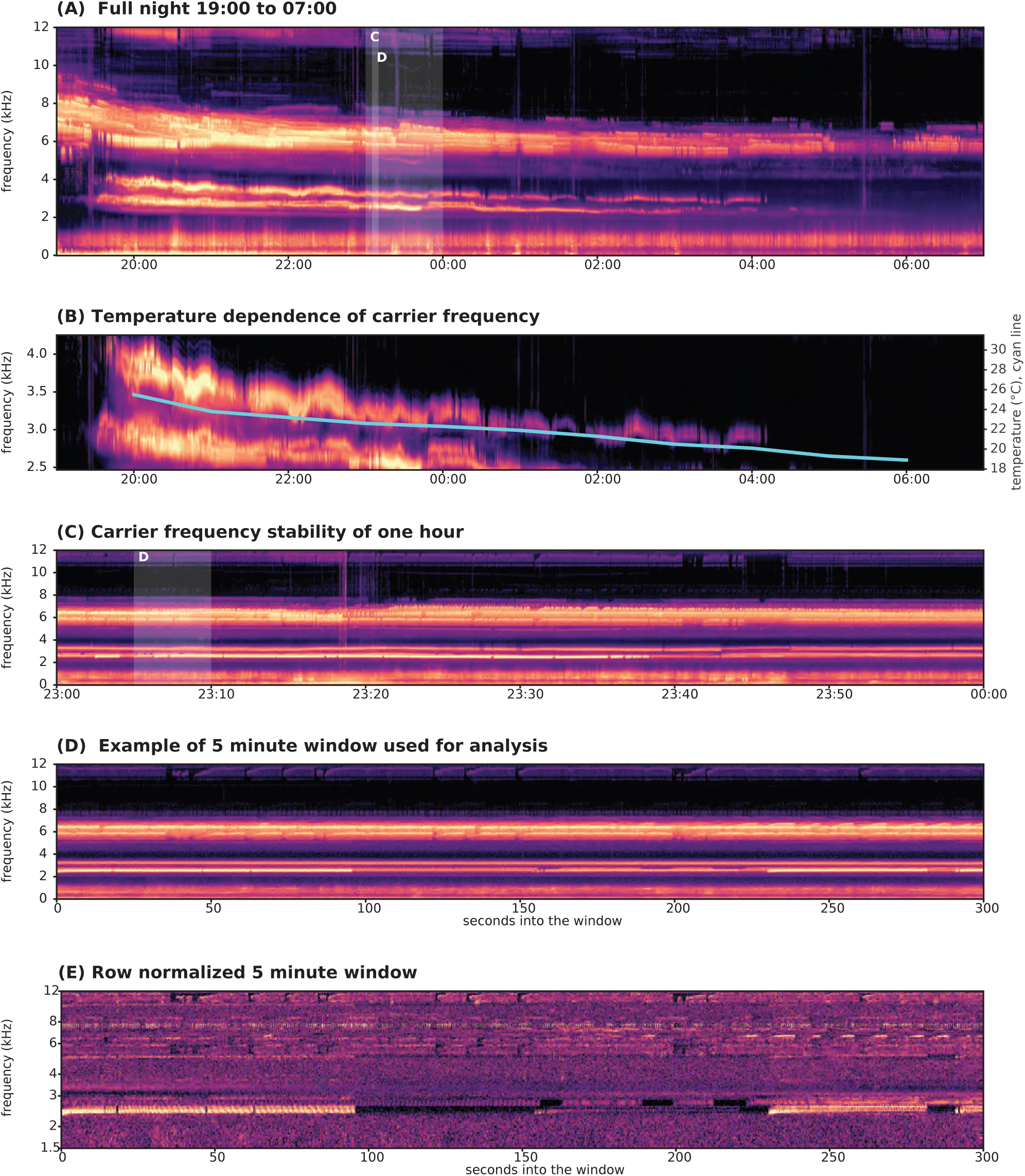
Example night audio. 13 hourlong audio files from recorder BARLT_00008662 on the night of September 20-21, 2025 were combined to cover the period of 7:00 PM to 7:00 AM local (UTC-6) time.

Where temperature data are used they are 2m above ground level (AGL) data from the High-Resolution Rapid Refresh (HRRR) atmospheric model produced by the US National Oceanic & Atmospheric Administration (NOAA) (Dowell et al. 2022).

### Identifying individual singers

As proposed by Riede (1993), we begin by identifying orthoptera-like songs in 5 minute windows by looking for discrete peaks in the three-dimensional space defined simultaneously by calling time, carrier frequency and modulation frequency. We use 5 minute windows because they are long enough to provide enough power to the modulation spectrum, but short enough that there is not significant drift in carrier frequency or modulation rate within the window. Many singers also exhibit a mix of bouts of singing and silence over the course of 5 minutes, something we make use of below in several ways. Windows sit on a fixed clock grid, every five minutes from 19:00 to 07:00 for a night and from 07:00 to 19:00 for a day, and each is read from the single hour-long file that covers its center, shifted inward when it would otherwise straddle two files.

#### Construction of a modulation map

For each 5 minute window, we begin by applying a short-time Fourier transform (STFT) to produce a raw spectrogram (using settings that produce 86 Hz frequency bins and 2.9 ms time bins, Figure 4A). We then take each frequency row, subtract its mean, and compute its Welch spectrum (at 0.17 Hz resolution) to reveal the periodicity at that carrier frequency, the modulation spectrum of that frequency band (Singh and Theunissen 2003; Miller-Viacava et al. 2026). (The Welch is time averaged power spectrum that trades lower noise for lower resolution). Highly repetitive calls like those of orthopteran singers appear in this Welch spectrum as a narrow peak at the pulse rate on a broad, sloping background. The background falls steeply below about 8 Hz and more gently above, so we measure the peak against it by subtracting a running median taken over 1.4 octaves of modulation rate, which follows the slope at every rate.

**Figure 4.**
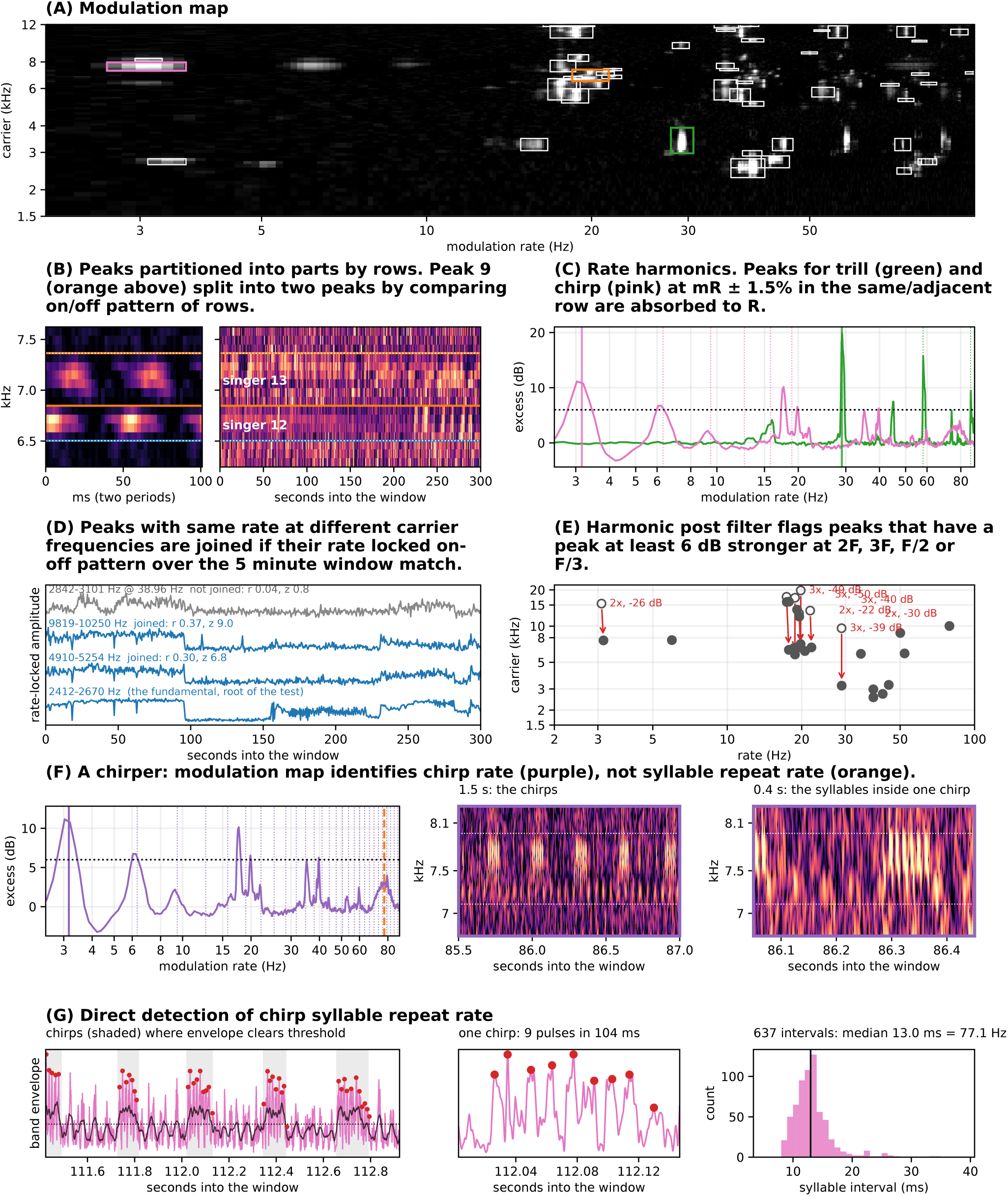
Overview of singer identification process Data shown are from five minute window starting at 23:05 local time (UTC-6) on September 20, 2025 on the House recorder BARLT_00008662.

After background subtraction, what remains is the excess modulation power over background in dB as a function of both carrier frequency (F) and modulation rate (R) at that frequency, which we plot as a “modulation map” (Figure 4B). The map we use spans carrier frequencies from 300 Hz to 22.05 kHz in 86 Hz bins, and modulation rates from 2 to 100 Hz in 0.17 Hz bins; the low end was chosen so that singers whose strongest rhythm is a chirp repeated a few times a second appear at that rate rather than only through its harmonics.

The modulation maps during times of orthopteran singing are generally well-resolved, and most consist of a small number of discrete peaks that represent one or more singers at the corresponding F and R. To enumerate these, we threshold the map at 6 dB above each row’s baseline, joining connected regions, and keeping any peak that spans at least two carrier frequency rows. In practice, these initial peaks, which span a compact set of carrier frequencies and modulation rates, often clearly contain pieces of the songs of multiple singers singing with nearly the same modulation rate in overlapping or adjacent frequency bands. Resolving them cleanly is the biggest challenge to automating the singer identification process.

#### Separating similar singers

As a first pass at separating individual singers, we group carrier frequency rows within the initial peak by their strongest rate bin, splitting the peak whenever there is a gap of more than one bin along the rate axis. This intentionally allows singers to have interleaved songs. We assign each resulting peak a coarsely estimated initial modulation rate R0 equal to the rate bin of the peak’s single strongest cell (at this point the resolution is within one bin of the modulation spectrum, in this case 0.17 Hz). We then find where in the 5 m window this rhythm is present, and measure the rate precisely from the audio data, to about 0.01 Hz.

For a singer that sings through most of the window this subwindow where we measure the rate is its loudest 30 seconds. For one that sings in bouts it is the set of runs of song of a second or more, each folded with its own phase and the runs aligned on their envelopes before they are summed, because a bout singer’s phase resets at every bout and a single fold over 30 seconds smears it. The refined rate is then checked against its own sub-multiples: if the fold at R/2 or R/3 is markedly deeper, we treat the peak as a harmonic of the true rhythm, and the rate is corrected. A peak below 8 Hz that shares its rows with a faster peak is treated not as a second singer but as that singer’s bout rhythm, the rate at which it groups its syllables into chirps, and is recorded as its chirp rate.

We next calculate two features of each frequency row within the peak and an additional 300 Hz in both directions. First, its syllable profile: we cut the singing time into periods of length 1/R, stack them, and average, so that the repeating unit stands out and everything else blurs away. Second, its singing pattern: for each half-second of the whole window, how strongly the row pulses at rate R, which quantifies when the singer is on and when it is silent.

We expect adjacent rows that belong to one singer to have nearly the same syllable profile and the same pattern of bouts. We therefore walk up the rows and start a new peak wherever either changes sharply. We then grow each peak outward into neighboring rows for as long as the profile stays continuous and the row still pulses at R, even faintly, since a louder steady neighbor can hide part of a singer’s band from the initial threshold.

#### Collapsing harmonics

A singer at carrier F and rate R produces further peaks at 2F, 3F (carrier frequency harmonics) and at 2R, 3R (modulation rate harmonics), and at combinations of the two. These harmonics are often detected as peaks on their own, but do not represent separate singers.

We absorb rate harmonics structurally: if there are peaks at R and at a multiple mR in the same rows, or within 300 Hz of them, we take the second to be a harmonic of the syllable rhythm rather than a second singer, since any repeating pulse produces such peaks. This is assumed from the mathematics rather than tested in each window; on a sample of 20 windows the absorbed harmonics switched on and off with their fundamentals as closely as a train’s own harmonic does, except where the fundamental was a chirp rate (this will be addressed in the next version of this analysis).

Two peaks at the same rate in different bands are joined into one singer only if they switch on and off together through the window, carrier harmonics included. A lone peak at a rate multiple of a singer in another band must also switch on and off with it, so that a chirper whose chirp rate is a sub-multiple of a trill’s rate is not absorbed into the trill. Every component is tested against the singer’s lowest-frequency component alone, never against the other components, so two singers cannot be merged through a third peak that resembles both.

Not every carrier harmonic is folded into its singer during the window analysis. After the window has been analyzed we therefore look for pairs of detections at the same rate whose carriers stand in a 2:1 or 3:1 ratio. If the upper member is at least 6 dB weaker it is flagged as the harmonic of the lower; if instead the lower member is the weaker one it is flagged as a shadow, the louder singer’s energy leaking into the band at half or a third of its carrier. Pairs of comparable level are left as two singers. Harmonics found at this stage are not folded into their singer; they are flagged and excluded from the analyses that follow. Where the fundamental was not resolved as a detection of its own, as happens in a dense chorus, its harmonics escape this flag. We also flag detections sitting on the fixed modulation rates of machinery, which we identify per recorder and month as rates that recur night after night with no change with temperature.

#### Differentiating trillers and chirpers

Some singers (chirpers) group a few syllables into chirps repeated a few times a second. Such a singer has a weak, broadened peak at its syllable rate, split into sidebands by the chirp rhythm, and a strong sharp peak at the chirp rate itself. Extending the map down to 2 Hz lets us see the chirp peak directly. A peak below 8 Hz that shares its rows with a faster peak is taken as that singer’s bout rhythm and recorded as its chirp rate (C) rather than as a second singer. The syllable rate inside the chirps is not read from the modulation spectrum, where a continuous neighbor in adjacent rows can leave a taller peak than the chirper’s own syllables, rather it is read in the time domain. We band-pass the audio to the chirper’s band, take its envelope, find the chirps as events where the envelope clears a threshold between its quiet and loud levels, and count the pulses inside each chirp. The syllable rate is one over the median interval between pulses, and a reading is kept only when the intervals are regular. Where the chirper’s own syllable peak was also detected as a separate singer in the same rows, the two are one singer and are merged. When no regular pulse train is found the singer is still recorded in the window, carrying its chirp rate alone. Because such a singer has no syllable rate it has no position in the carrier-by-rate plane used below and does not enter the tracks. Chirping singers whose syllable rate is read are tracked like any other, with the chirp rate carried alongside. Whether a tracked singer chirps or trills is decided again later, from its audio, when call types are defined, so that a chirping and a trilling species at the same carrier and rate are kept apart.

#### Single night outputs

The output of a window is thus a list of singers active in that window, along with several measures of the song and its behavior in the window. For the night shown in Figure 3, we identified a total of 7,739 peaks in the modulation map. Of these, 5,508 produced at least one part that passed the extent and modulation depth tests, yielding a total of 6,089 candidate singers. 1,888 of these were scored as rate harmonics and absorbed into a lower rate part, and the remaining 4,201 were joined into 3,065 singers by the co-variation test. Of these 2,359 consist of one fundamental part (a single carrier frequency range), while 706 join two or more carrier components. Of these 706, most (600) are carrier-rate harmonics, and the remainder are single-band songs stitched back together after having been separated in earlier steps, usually by an overlapping song. The postfilter flagged 3 pairs of songs as chirp-syllable pairs, and an additional 500 likely harmonics, yielding a total of 2,562 discrete songs from the 72 windows.

Figure 5A shows songs from the window analyzed in Figures 4, and Figure 5B shows the 2,562 from this one night run for one recorder plotted on a modulation rate vs. carrier frequency log-log axis, colored by the temperature at the corresponding hour (see below for where the temperature data comes from). The majority of points, extracted independently across the night, are clearly organized into multiple streaks where both the carrier frequency and modulation rate rise in tandem with temperature. Figures 5C-G show snippets and syllable profiles for 3 songs sampled from along the streaks (C-E) and chirp cluster (G) from Figure 5B, providing support for the hypothesis that they represent the same basic song.

**Figure 5.**
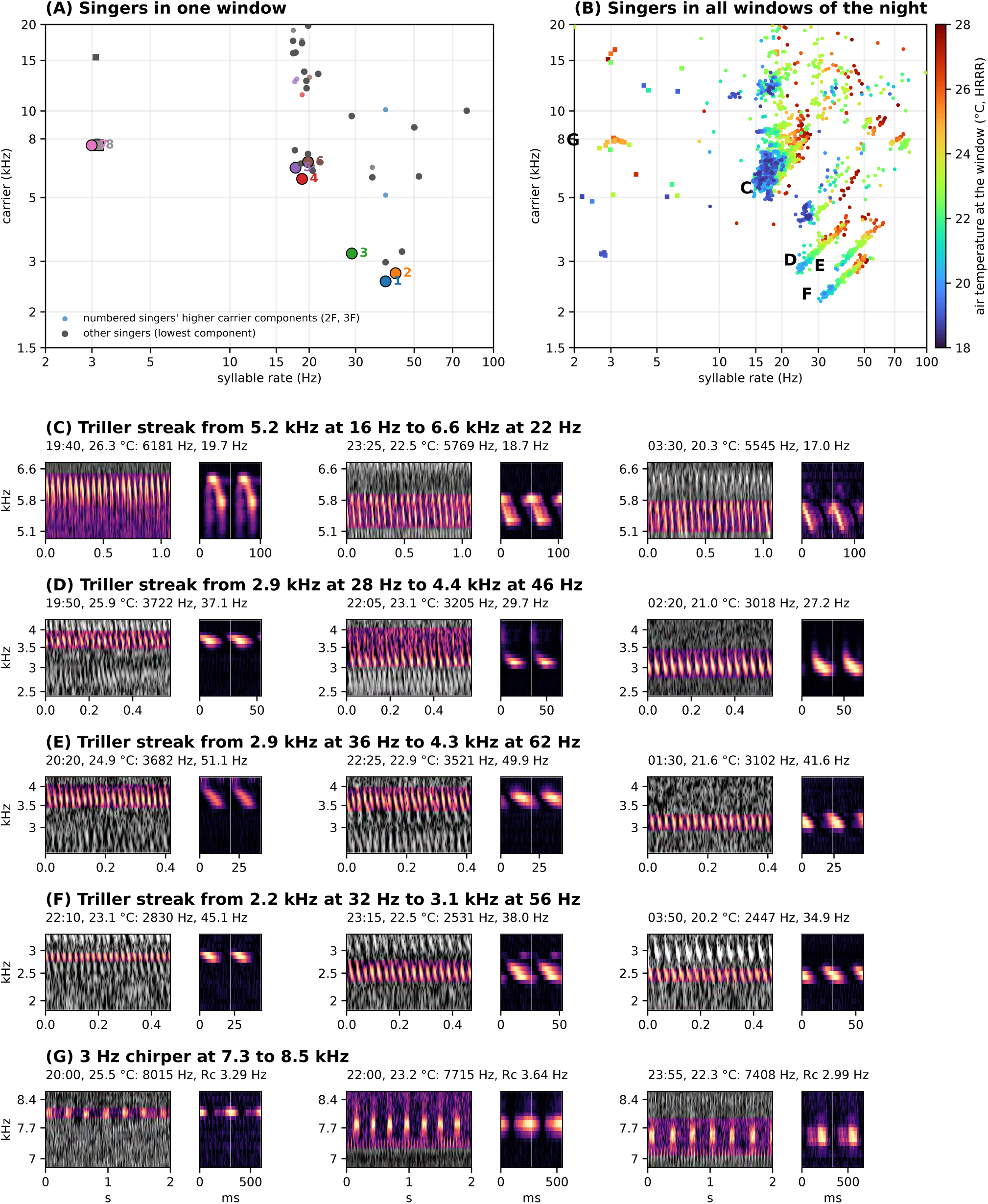
Coherent organization of individual songs across one night (A) Individual singers identified in window from five minute window starting at 23:05 local time (UTC-6) on September 20, 2025 on the House recorder BARLT_00008662. (B) Singers across night plotted with temperature. (C) to (G) show individual songs with their folded syllable.

#### Tracking singers across a night

Based on this observation, we next sought to link the singers found across the 12 hour night or day period of a recorder into tracks (the linking process and its application to the night of Figures 3 and 4 is presented in Figure 6). To do this, we first subtract the rate drift shared by all singers across the 12 hour period being analyzed, estimated window to window as the median change among matched singers, so that each singer only has to be followed through its own departure from that common drift. We consider a song in one window may then continue a track from an earlier window if its rate is within 2 % of the track’s (plus 0.4 % for each window skipped, up to six windows, or half an hour, since bout singers fall silent for minutes at a time and a faint singer drops below threshold on occasion) and its lowest-frequency band overlaps the track’s with 150 Hz of slack. Rather than pairing singers greedily window by window, we choose the set of tracks for the whole night at once, as the minimum-cost assignment in which each singer in each window continues at most one track: the cost of a link grows with the difference in rate and in carrier center, and with each skipped window, and ending one track to start another carries a fixed cost, so a new track begins only where no plausible continuation exists.

**Figure 6.**
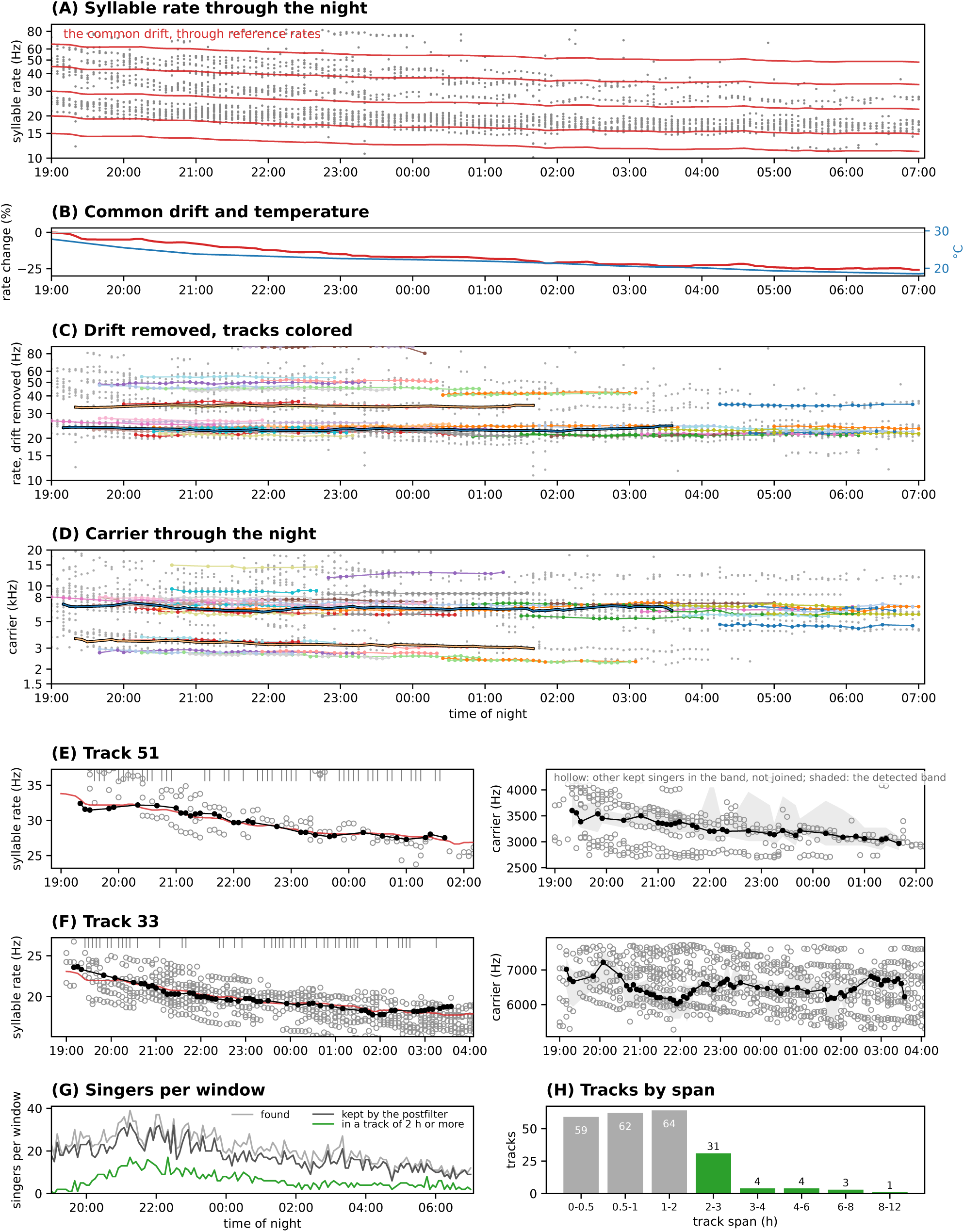
Linking singers across one night Overview of the process of linking singers including removal of common drift (A,B), connecting tracks by rate (C), and tracking carrier rates (D). Application to individual tracks (E,F), and results (G,H).

Because the rows assigned to a band change from window to window as faint edge rows cross or miss the threshold, a carrier measured from the band itself jumps by tens of Hz for reasons unrelated to the singer. We therefore fix, for each track, one reference frequency range, the lowest band present in at least a fifth of its windows, and in every window measure the center of energy over that same range, weighting each row only by the energy that pulses at the singer’s rate, so that a steady neighbor in the same rows does not change the estimate.

### A detailed record of orthopteran songs across a year

Over the thirteen months from November 2024 through November 2025, 27 recorders in 11 areas of the ranch recorded 176,184 hours of audio in 178,747 hour-long files, covering 395 calendar days and 7,782 recorder-days (the full archive back to October 2023 holds 403,465 hours). BROOKS analyzed this as 14,641 recorder-periods, 7,321 nights and 7,320 days, and 1,231,920 five-minute windows with audio. Those windows yielded 10.9 million window-singers, of which the postfilter kept 9.5 million: 8.95 million with a syllable rate and 0.53 million chirp-rate-only singers whose syllables were not resolved. Linking within each night or day joined them into 911,005 tracks, of which 103,802 lasted two hours or more (71,770 at night, 32,032 by day) and carry 335,882 track-hours. Of these, 26,116 night tracks and 5,427 day tracks became singer-nights and singer-days with a temperature at each hour, giving 139,171 night singer-hours and 27,673 day singer-hours, at air temperatures from −6 °C to 36 °C (median 22 °C).

These data, built progressively from 5 minute windows at specific recorders, provide a detailed record of the song and singing behavior of nearly a million orthopteran singers within range of one of these recorders across the year, linked to their location and the external conditions (especially temperature) in which the song occurred. The next question is whether we can efficiently group together singers of the same sort based on the measured attributes of their song, as suggested by Riede (1993), Walker and Funk (2014), and others.

#### Numerous types of trilling singers with distinct and strongly coupled carrier and rate variation with temperature

The obvious place to start is to look at where these songs fall in the plane of carrier frequency vs rate (syllable and/or chirp). Figure 7A shows the 135,124 night singer-hours of trilling, or mixed trilling-chirping, tracks plotted, as above, by their carrier frequency and syllable repeat rate, and colored by the temperature at which they were sung.

**Figure 7.**
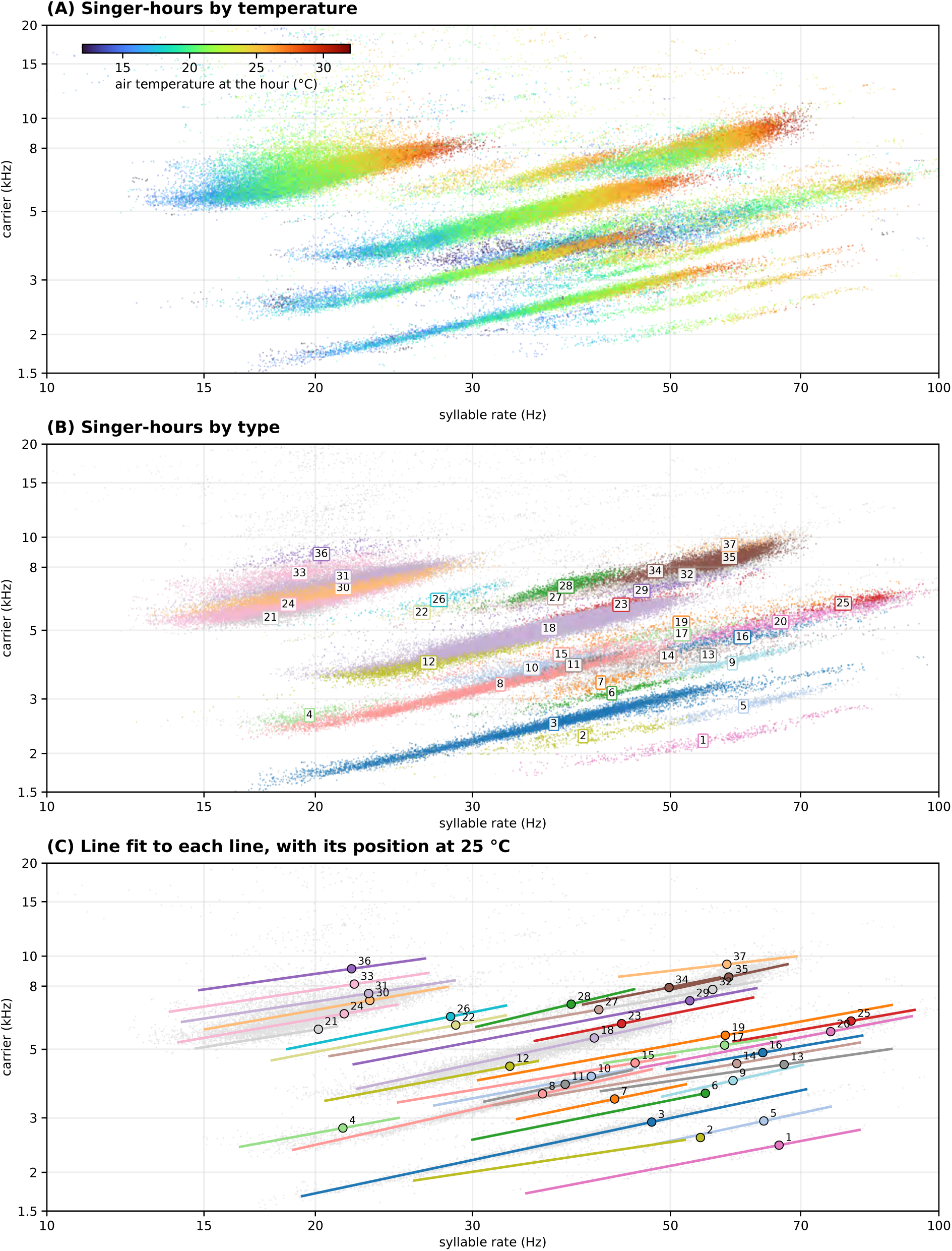
A year of trillers (A) Point for every triller singer-hour on carrier frequency by syllable-rate plane colored by HRRR 2m air temperature. (B) Classied triller plane types. (C) Lines fit to singers in each type.

The streaks evident in a single recorder-night dominate, and are refined, expanded and increased in number across the year. We enumerated streaks by first placing all singer-nights on the carrier frequency × syllable rate plane at their median values, brought to 25 °C from the singing period’s median temperature along the global direction of temperature dependence. Singer-nights were then grouped by average-linkage clustering in a frame rotated to that line, with a tolerance of 0.8 in log rate along the line and 0.06 in log carrier across it, so that a population spread along its line by individual variation and residual temperature effects falls in one cluster while a neighbor a few percent away in carrier does not. A cluster was split where its singer-nights formed two parallel bands against temperature, two clusters were merged where their scatter showed them to be one population. Where a cluster held both chirping and trilling singers, judged from their audio, it was divided into a chirp type and a trill type at the same position.

We then examined each cluster’s singer-nights on the plane, colored by temperature, together with plots of individual singers’ paths through their nights, and split a cluster where it held two populations that the clustering had merged: two parallel bands offset in carrier, two rate lines at the same temperature, or a group of singers whose rate at a given temperature differed from the rest. We required that individual singers not cross between the proposed halves within a night. In the clusters we split, fewer than 10 percent did, against 25 to 40 percent separation between the halves, whereas a cluster that was one population cut in two had a fifth of its singers crossing. Six clusters were split this way: three into two parts, one into three, one into four and one into five. Every resulting group of at least 40 singer-nights is a streak type (Figure 7B). In total, there are 37 trill types and 8 chirp types. Figure 7C shows the log-log slope for each type, and the inferred carrier frequency and syllable repeat rate at 25C.

The slope of a type’s streak is a population pattern. It is an aggregate of all the singer-nights of many animals at many temperatures laid on one plane, but it does not show us how individual singers move within it. To characterize that, we fitted log carrier on log syllable rate within every track of twelve windows or more whose rate moved by at least 2 percent over the night, 17,799 tracks in all. Figure 8A shows six of them, one from each of six types: the singer’s windows as carrier against rate about its night median, and beside them the paths of seven singers of the same type across the type’s streak (Figures S8-{type} in the Supplemental Material show a selection of individual singers for all types, including a snapshot of their song and the path they take through the type’s streak). Figure 8B gives the median and interquartile range of the within-singer slope for each of the 37 types, with the type’s streak slope from Figure 7C beside it.

**Figure 8.**
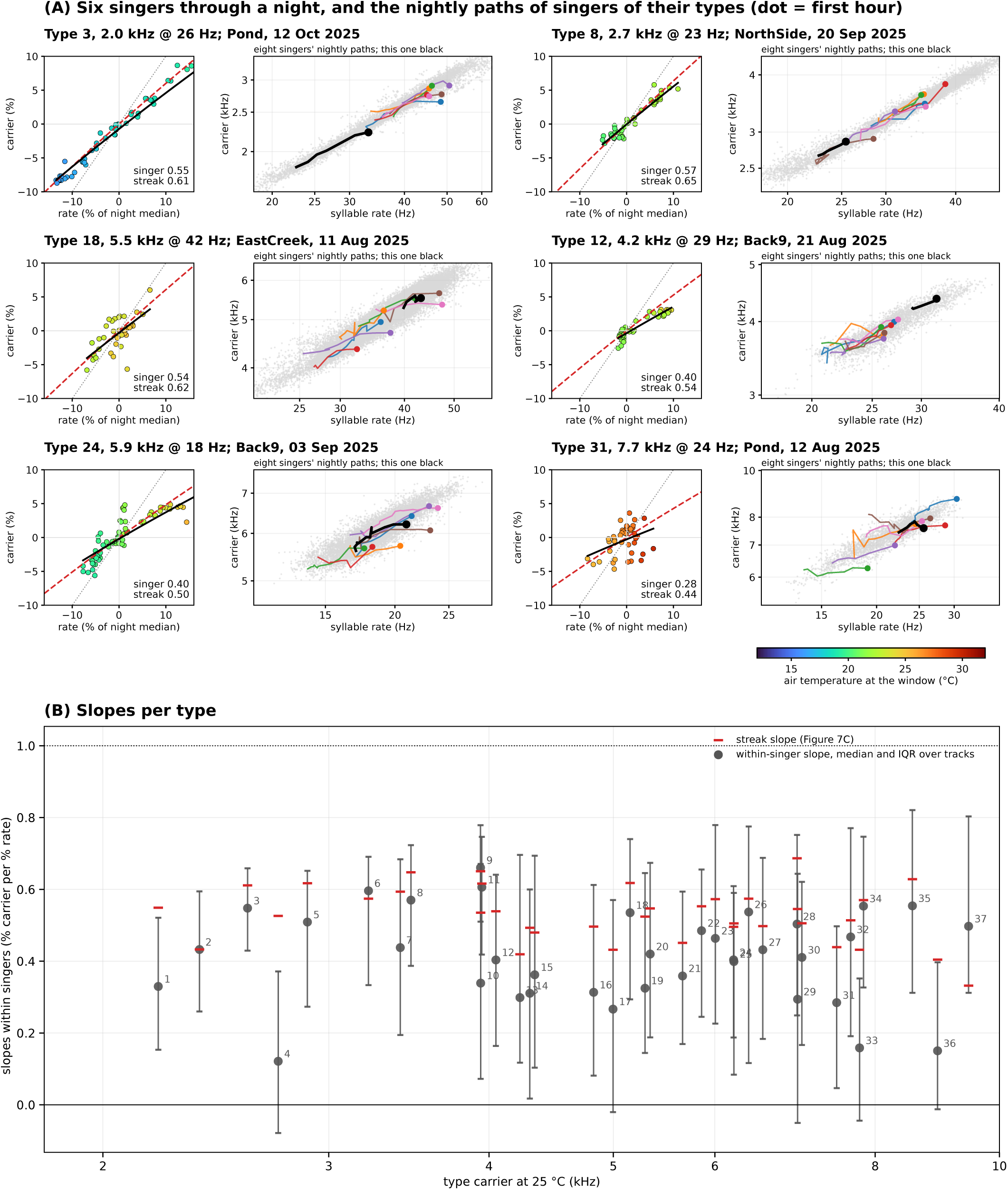
Carrier follows rate within a singer (A) Individual traced singers broadly follow type line on single nights, and have within night slopes similar to, but slightly lower than type. (B) Slopes of all singers within night by type.

The two slopes agree for many types but not all. The streak slopes lie between 0.33 and 0.69 (median 0.54), and for 16 of the 37 types the median singer’s slope is within 0.1 of its streak’s. Below 4 kHz the singers’ slopes (median 0.53) match their streaks; above 4 kHz they are generally lower (median 0.40), and in seven types, all above 4 kHz except type 4, the singer’s slope is below 0.3 while the streak’s is 0.4 to 0.55. The lower slope is due to generally smaller shifts in log frequency with temperature as opposed to an extended range of syllable rates. The streak’s slope would then have to come from differences between singers rather than from temperature.

#### Chirping singer types distinguished by carrier and chirp rate

Chirping singers carry two rhythms, the syllable rate inside the chirp and the rate at which chirps repeat. BROOKS reads the chirp rate directly from the modulation map, and the syllable rate from the audio inside the chirps where the syllables are regular enough to measure. Of the 7,837 tracks that chirped in at least half of their windows (32,908 singer-hours), 3,943 carried a syllable rate in at least half of their hours; these we placed on their own plane, carrier against chirp rate (Figure 9A), and against syllable rate (Figure 9B) for the 14,199 of their 15,547 hours that have one. The 3,894 tracks with a bout rhythm but no readable syllables are set aside: they include frogs and other sounds that pulse below 8 Hz, and we do not type them. The chirpers show the same picture as the trillers, parallel streaks rising with temperature, in the carrier against chirp-rate plane; the carrier against syllable-rate plane is less well resolved but divides into two groups, chirpers below 5 kHz singing 39 to 72 syllables per second and chirpers above 5 kHz singing 58 to 94. Clustering on the chirp-rate plane gave six types with at least 40 singer-nights; two of them held singers with two distinct syllable rates at the same temperature and were split on it, giving eight chirp types (Figure 9C) that hold 95 percent of the typed chirping singer-nights and are compact on the syllable-rate plane as well (Figure 9D). Chirp rate rises with temperature within a type by 1 to 7 percent per degree, the syllable rate by 4 percent or less, so the number of syllables per chirp period is not a constant of a type: in the 3.6 kHz chirper C1 it falls from about 30 on cold nights to 12 on warm ones, in the 3.9 kHz C2 from 14 to 10 (Figure 9E). The two groups differ in which rhythm they hold steady: the low-carrier chirpers keep their syllable rate nearly fixed and vary their chirp rate, while above 5 kHz the chirp rate hardly moves with temperature and the syllable rate varies between singers.

**Figure 9.**
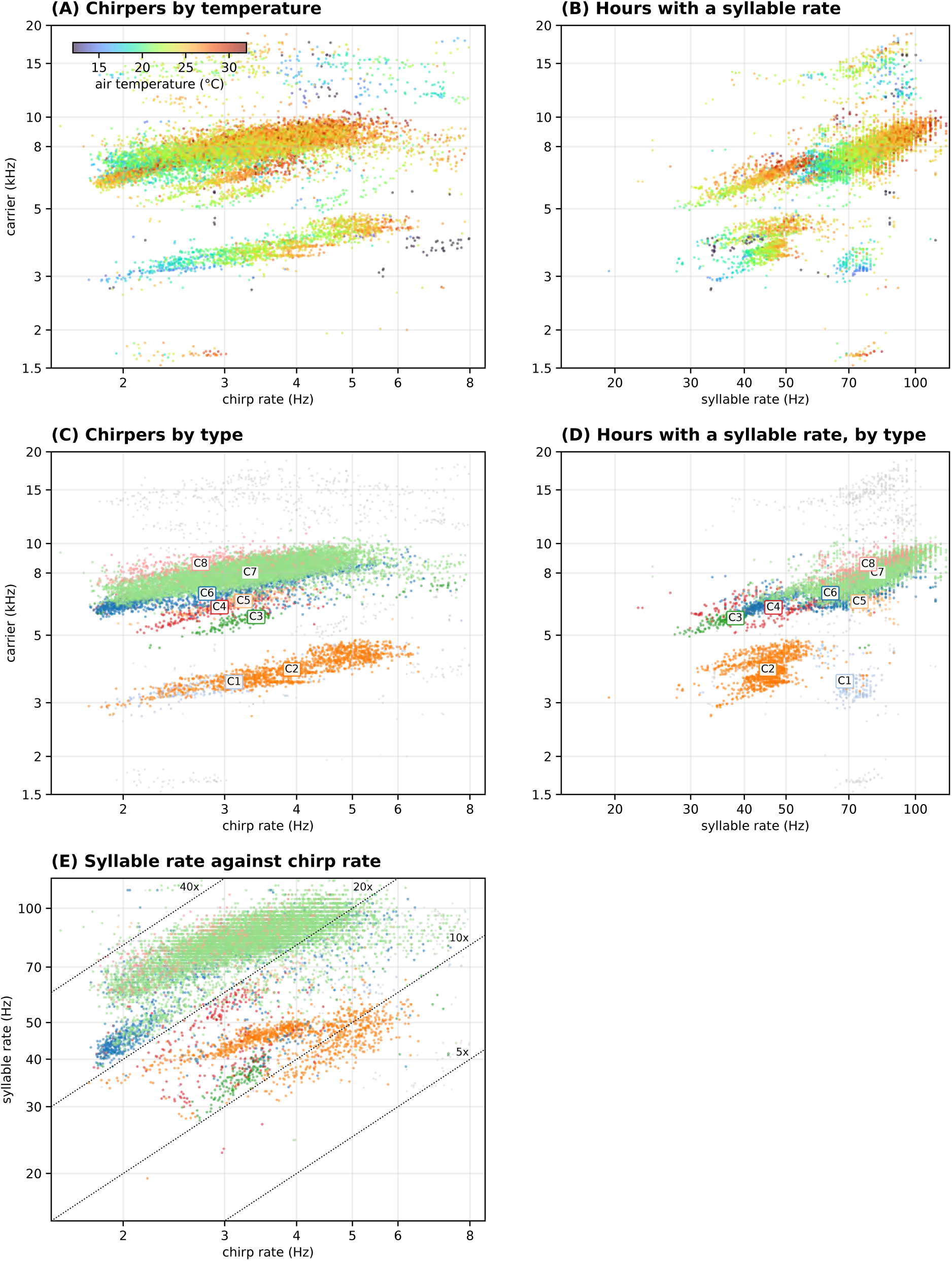
Identification of chirper types Chirpers have three measured rates: carrier frequency (F), syllable repeat rate (R) and chirp rate (C) and are plotted FxC (A) and FxR planes (B) to facilitate type identification (C-E).

#### Streak types hold consistent song types

To this point we have defined streaks solely on the basis of carrier frequency and rates extracted from 5 minute time windows, resolved by observed continuity and consistency in temperature responses. The similarity of this behavior to measurements from individual singers in the controlled settings makes it tempting to define these streak types as distinct biological entities. But before pursuing this idea further, we first wanted to make sure that songs sampled from different points along the streak axes defined above were consistent with having a similar biological origin. Figure 10 shows three types (two trillers and one chirper) as grids of half-second excerpts: six positions from the cold, slow end of the line to the warm, fast end, and at each position three singers from different recorder-nights, with the rows outside the detected band drawn in gray so that the band the detector found is visible against the rest of the spectrum. For type 8 (a 3.5 kHz triller), type 24 (a 6.2 kHz chorus triller) and type C2 (a 3.9 kHz chirper), the song form, syllable shape and glide are the same at every position. The same grids for every type with at least 40 singer-periods are in Figures S7-{type} in the Supplemental Material, together with a rate-filtered view that averages each excerpt over its own period so that the singer’s syllables stand out from its neighbors’.

**Figure 10.**
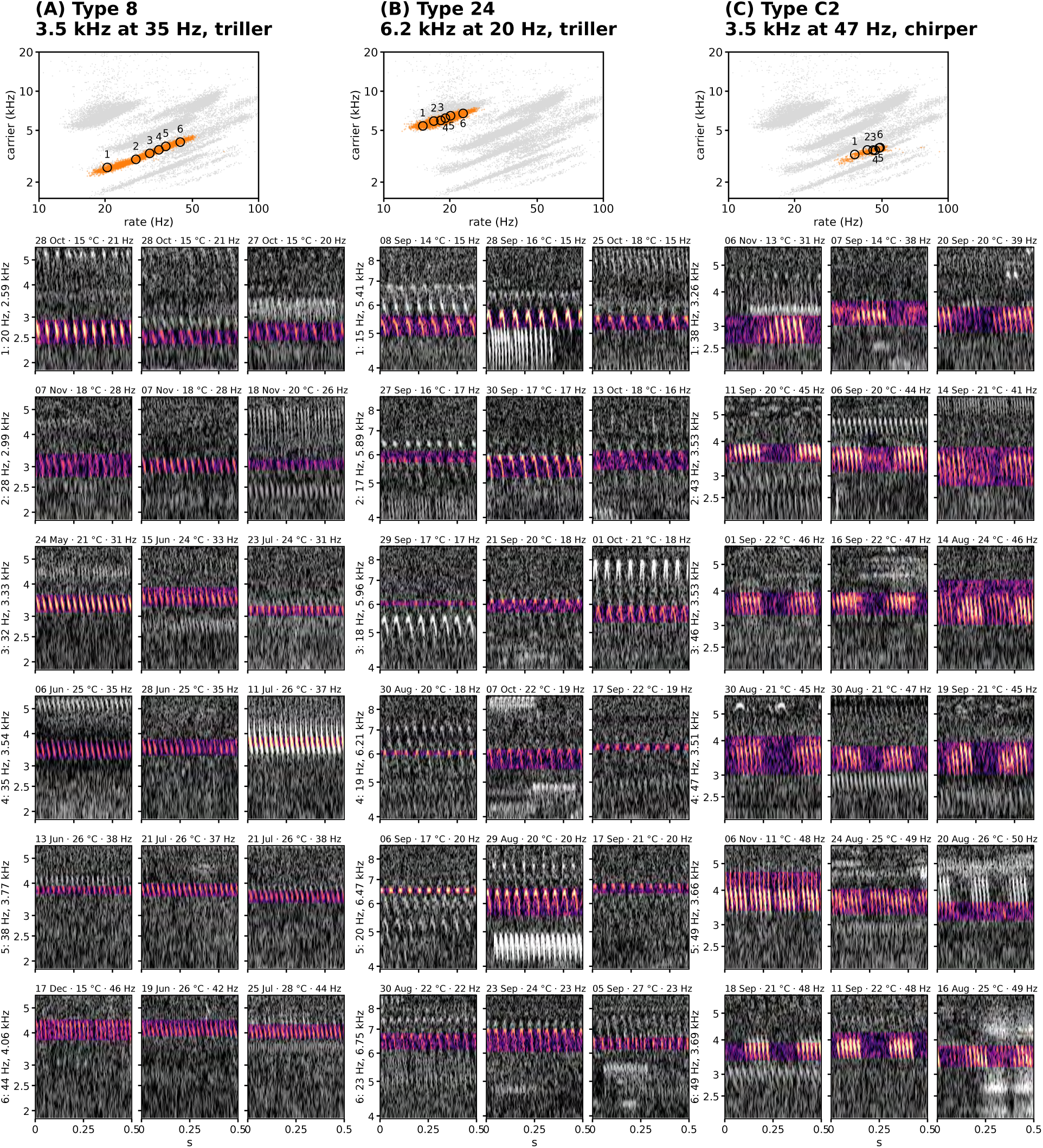
Coherence of three streaks across the year Three different streaks show coherent song types across temperature range. See also Figure 7S.

#### The mix of types through the year and across microhabitats

Because every type is tracked the same way on every recorder, the number of singers of a type tracked at once per recorded hour is a useful measure of its activity across seasons and geography. Figure 11A, which shows how the mix of types changes across the year, captures the seasonal variation well. The first November captures the tail end of a fall orthopteran chorus, before things go largely silent from December to March. There is a rise again in April and May carried by a few trilling types at about one singer per recorder-hour, before a late-summer peak in August and September when nearly five singers were tracked at once on an average recorder-hour, falling to nothing again by the end of November. Each type occupies its own window in the year; the 16 most active are shown by number and position, and three chirp types are among them.

**Figure 11.**
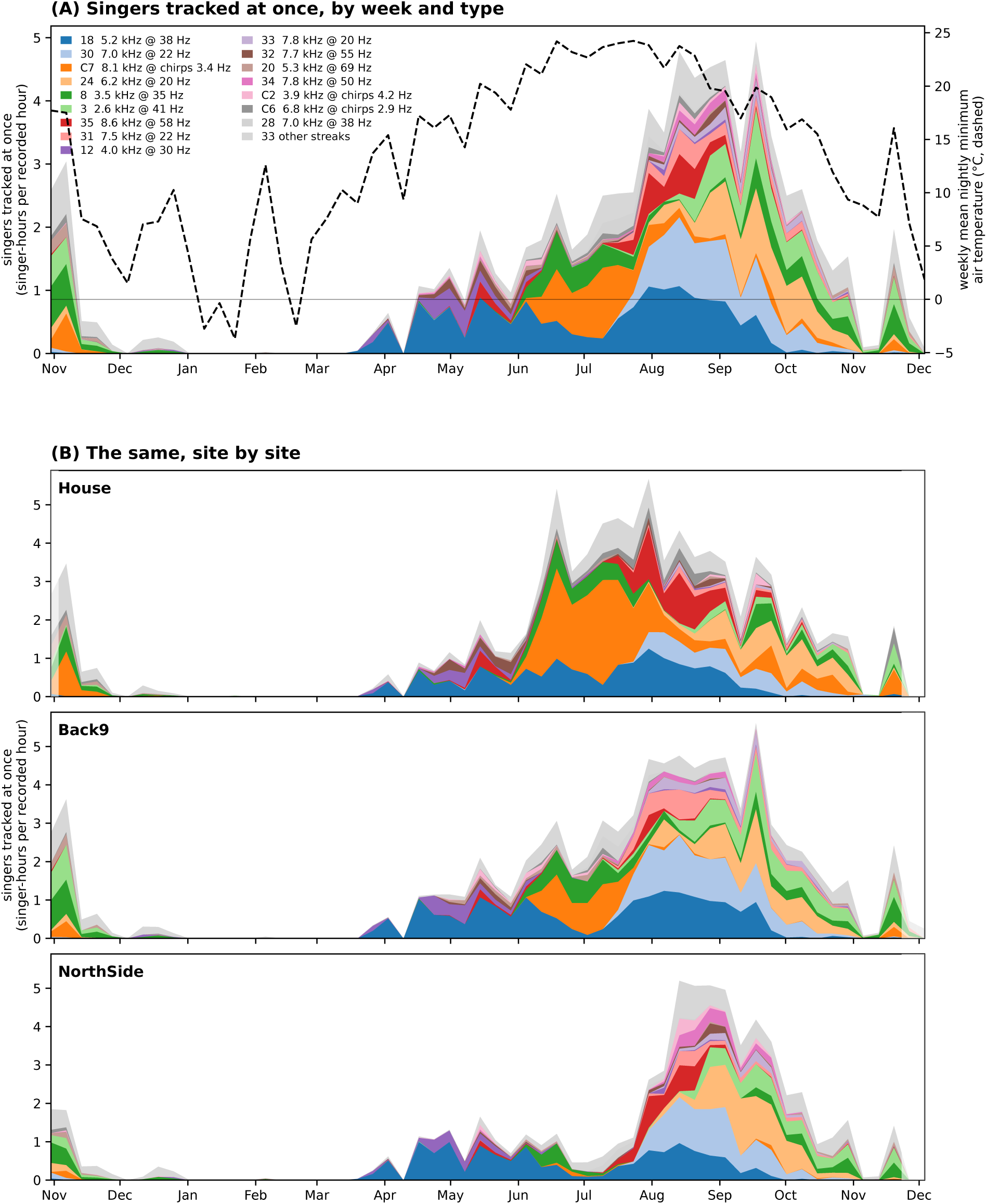
The mix of call types through the year and across sites Mix of different triller and chirper types across the year (A) and across three sites with multiple recorders (B). The sites are House, located in an open field, Back9, adjacent to a hardwood border to a loblolly pine plantation, and NorthSide in a field alongside tree-lined creek, and adjacent to season pond.

Figure 11B shows the same measure at three sites representing distinct microhabitats and robust recorder coverage. The Back9 recorders are in a field adjacent to a treeline bordering a mature commercial pine plantation. It is the highest and driest of the three sites. The House recorders are in a thicket of berry bushes and other annuals on a hill with only two small nearby trees. The NorthSide recorders are in the lowest and wettest area of these three, along the stand of trees that flank a large creek, and adjacent to a seasonal pond. The general shape of the year is the same at all three: the 5.2 kHz triller type 18 forms the base of the season from May onward at each, the 6.2 kHz tick type 24 and the 2.6 kHz triller type 3 close it in September and October, and the November chorus looks alike at every site in both years. But the summer peak is made of different types. At House the 8 kHz chirper C7 carries June through August, a quarter of all singer-hours, with the 8.6 kHz triller type 35 beside it, and the 7 kHz triller type 30 is scarce. At Back9 and NorthSide type 30 is the leading type of July and August and C7 is minor at Back9 and all but absent at NorthSide, where type 35 takes its place. The peak is reached in mid-August at House and NorthSide and a month later at Back9.

#### Identifying the singers

We are obviously ultimately interested in identifying the species producing each of these songs, and we have taken the first step towards doing this by comparing the song types we identified to reference recordings. The most comparable dataset covering species present in this region is that of Walker and Funk (2014), who measured the carrier frequency, pulse rate and temperature of 1,318 field and laboratory recordings of 13 North American species of *Anaxipha* at a range of temperatures. *Anaxipha* are small trigs of the kind that, we believe, dominate our data between 4 and 8 kHz, and several of the species have been described in our region.

We took Walker and Funk’s published per-recording measurements of pulse rate, carrier frequency and temperature and plotted them on the same plane as we used to identify singing types (Figure 12). Their recordings have the same general properties of our data, forming streaks of roughly the same slope, especially in the focal region of our triller analysis with rates over 15 Hz. Each species traces a line of the same slope as ours, 0.4 to 0.9 in log carrier on log pulse rate, and its temperature runs along the line in the same direction, cold at the slow end and warm at the fast end. Their pulse rates rise by 4 to 7 percent per degree and their carriers by 2.5 to 4 percent, the same regime as our types.

**Figure 12.**
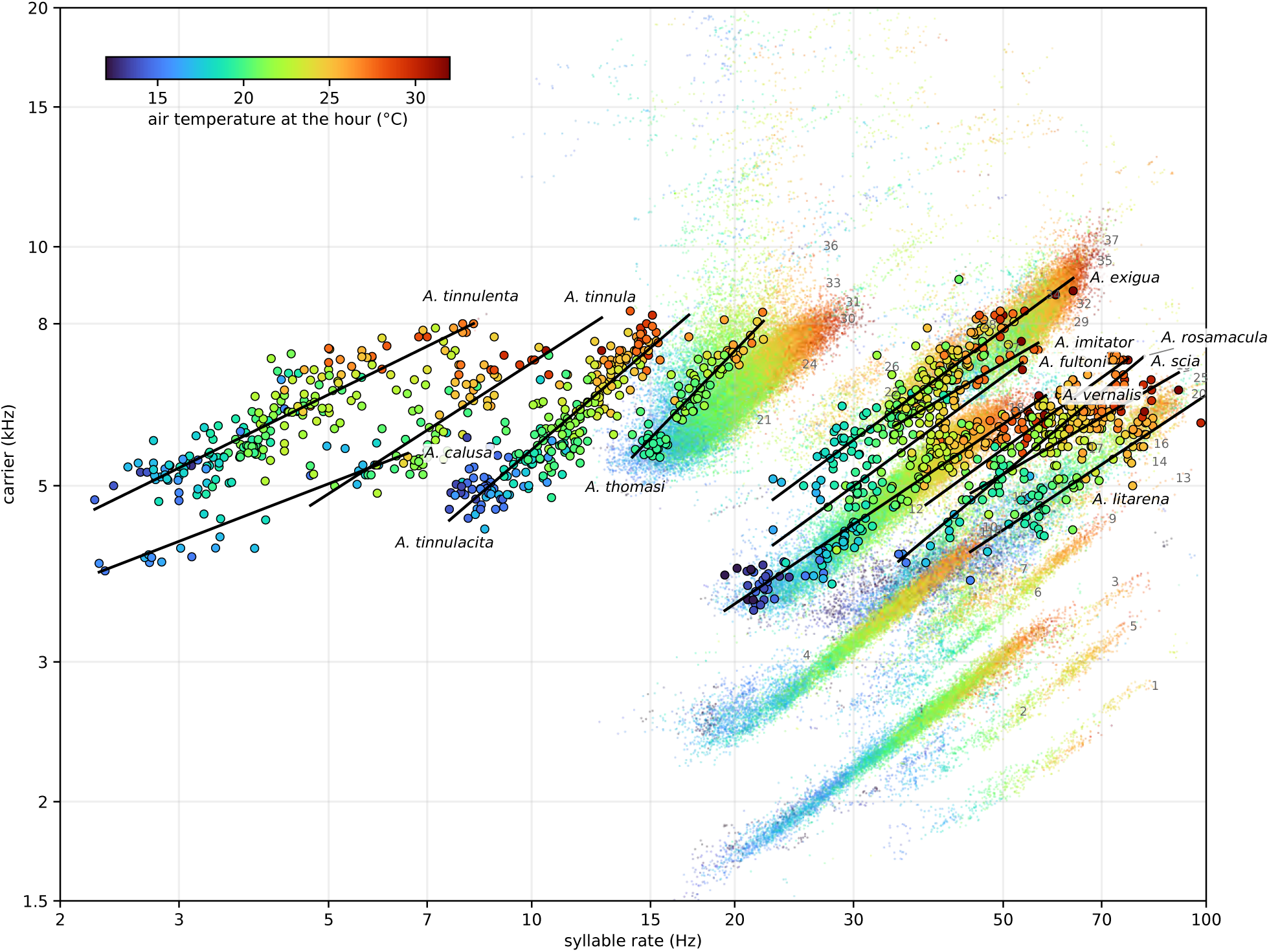
Published *Anaxipha* recordings on the year’s plane The year’s trill singer-hours colored by air temperature, as in Figure 7A, with 1,318 recordings of 13 *Anaxipha* species from Walker and Funk (2014; their supplementary song table) as rimmed circles colored by the recording’s own temperature on the same scale, each species with its fitted line and name; gray numbers mark the warm end of each type’s line.

Notably, several of their species overlap closely with our types. We tested each species against our types by calling a recording as lying on a type when its carrier is within 2.5 standard deviations of the type’s line at its pulse rate, and its pulse rate is within 2.5 standard deviations of the type’s rate at the recording’s temperature. By this test 94 percent of the 149 *A. vernalis* recordings lie on type 18 (5.2 kHz at 38 Hz at 25 °C), our most active type, and 79 percent of 193 *A. exigua* recordings lie on type 27 (6.5 kHz at 39 Hz), with the neighboring types 28 and 34 nearly as good a fit; *A. thomasi* lies on type 31 (7.5 kHz at 22 Hz), 62 percent of 79. Both *A. vernalis* and *A. exigua* are known from Arkansas. The agreement is not a matter of position alone: the recordings of these species sing at our types’ rates at the same temperatures, over the 12 to 35 °C their data span. Three species, *A. tinnula*, *A. tinnulacita* and *A. tinnulenta*, sing at 3 to 15 pulses per second, below the rate at which the modulation map detects a trill, and have no type here. Three others, *A. litarena*, *A. rosamacula* and *A. delicatula*, fall on types 19 and 20 as cleanly as *A. vernalis* falls on 18, but they are Gulf-coast and Florida species with few or no records near the site, highlighting that a match on this plane alone is insufficient evidence for species identification.

To expand this analysis beyond Anaxipha, we turned to Singing Insects of North America (SINA; Walker and Moore), created by Thomas Walker and maintained by the Orthopterist Society, which is the standard reference for orthopteran calls in the United States. We ran the SINA reference recordings through the same analysis we applied to our data, so that each recording becomes a point on the same plane measured in the same way; 157 recordings of 134 species yield such a point. 133 of these recordings, representing 111 species, are of species observed in Arkansas or neighboring states, and are thus plausible candidates for our field site.

In addition to running SINA recordings through our process, we also placed each species’ published pulse rate and dominant frequency, from the site’s text, on the plane as a second, independent reading. Where a species has both a recording and published numbers the two usually agree, within 5 percent in rate and carrier for most of the 54 species that have both. Where they disagree, it is by the structure of the song, a published pulse rate against a recording read at its chirp rate, and the published number is then not evidence about the same rhythm.

Figure 13A shows the 133 in-range recordings of 111 species over the year’s singer-hours and the fitted lines of the trilling types, with the published positions of the species nearest each line as diamonds. The geographically implausible species are included for reference, but not analyzed further. 70 recordings lie within 2.5 standard deviations of some line across it, and 53 of those also sit where the type sings at the recording’s temperature. Figure 13B lists, for each type, the recordings on its line placed by that along-line position, and the published position of the nearest species at 25 °C. Several lines carry one recording close to the line and to the expected rate, others carry several candidates that rate and carrier alone cannot separate, and the densest line of the summer, the 6.2 kHz chorus, matches recordings of several ground crickets.

**Figure 13.**
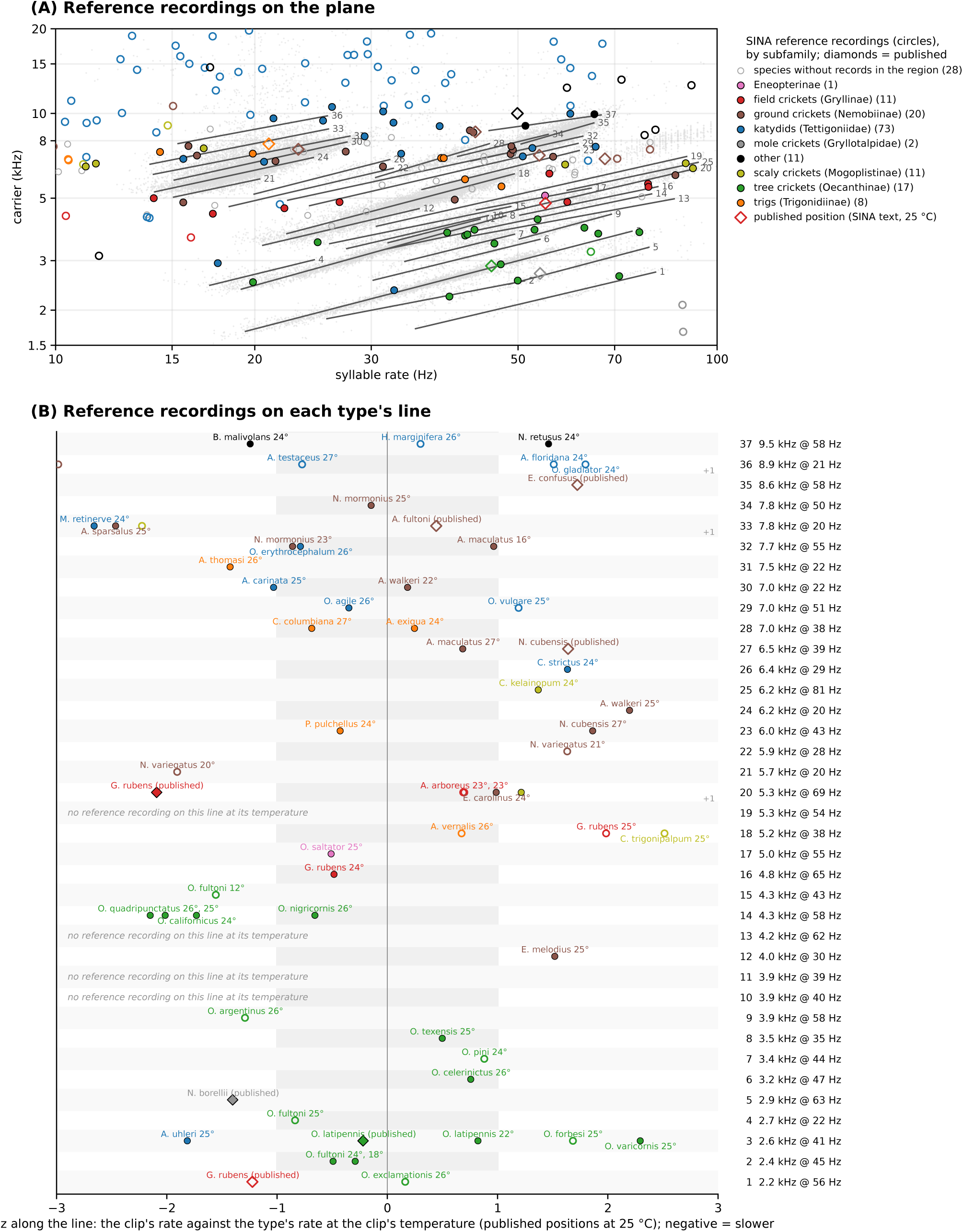
Reference recordings and published positions on the lines

In the absence of confirmed field identifications, we are reluctant to formally identify any of these singers. However, for most of the best candidates, there is a reassuring similarity between the reference call and samples from the corresponding streak. Figure 14 shows, for each of the matched types, a spectrogram of a half second sample from the reference recording along with four corresponding samples from the streak at approximately the same temperature as the reference. These by and large match, leading us to believe that, with additional reference recordings and input from experts, that automatic identification is feasible. For now, however, we leave these as candidate identifications.

**Figure 14.1.**
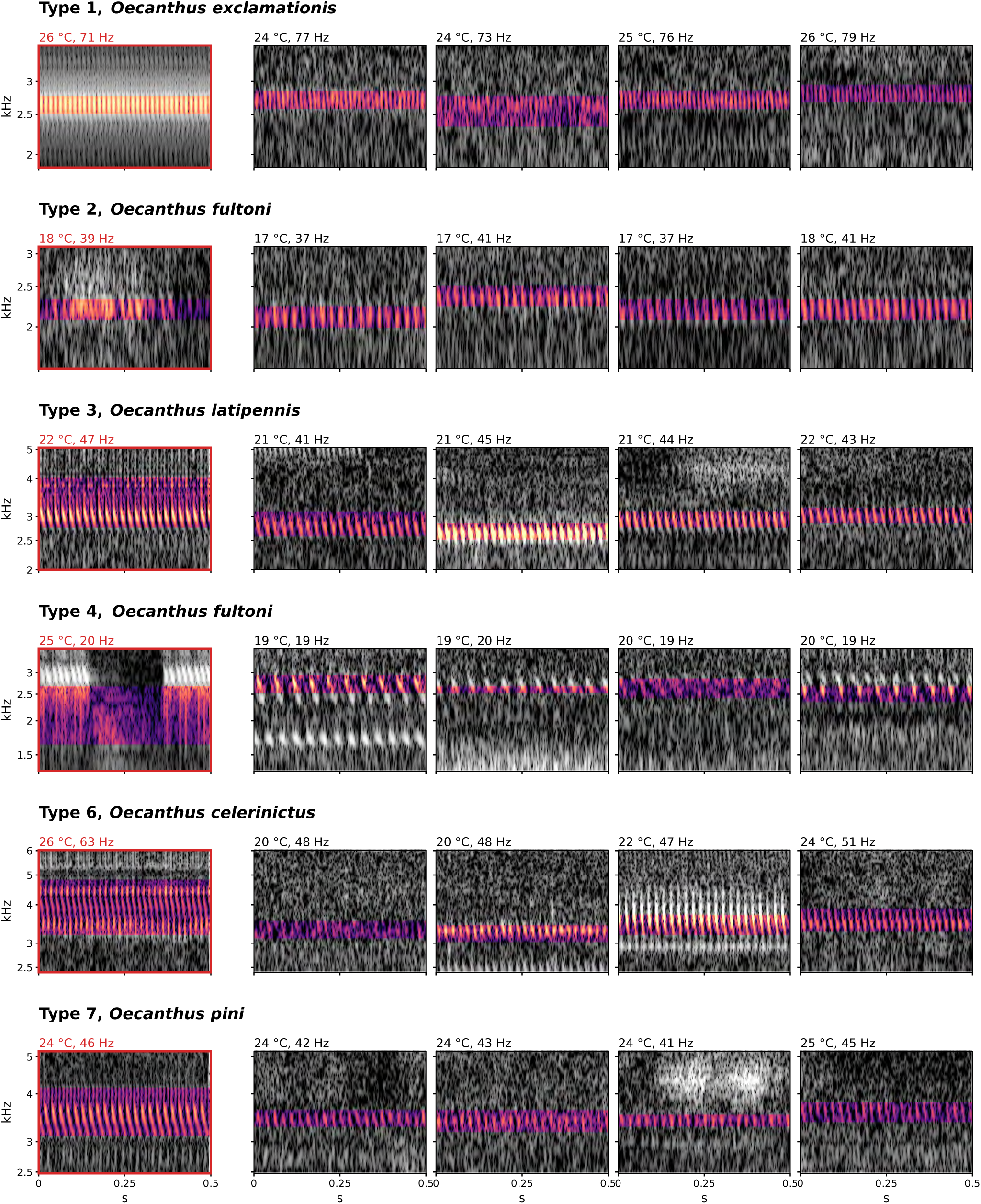
Reference recordings beside field singers of the type they match (1 of 7)

**Figure 14.2.**
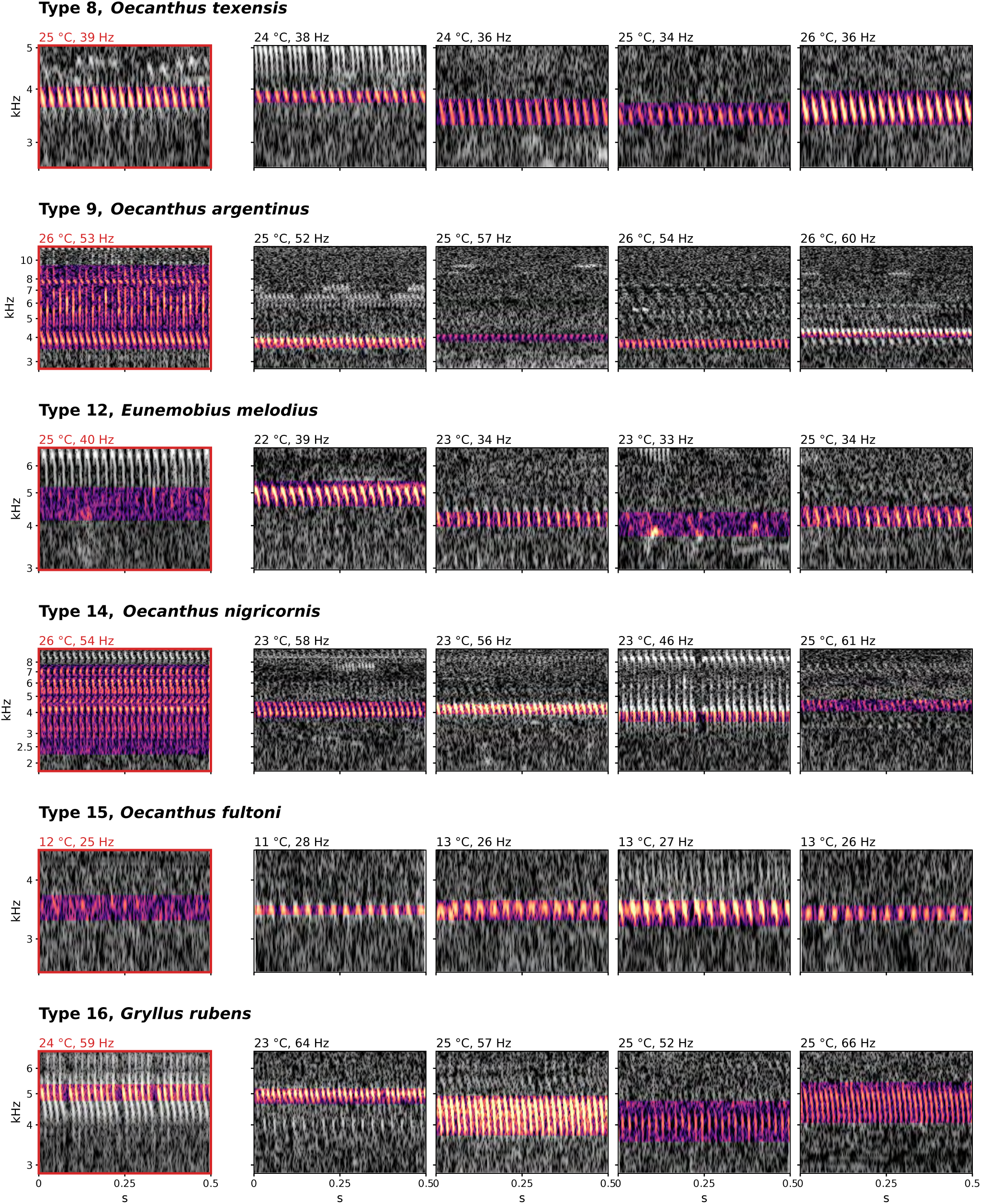
Reference recordings beside field singers of the type they match (2 of 7)

**Figure 14.3.**
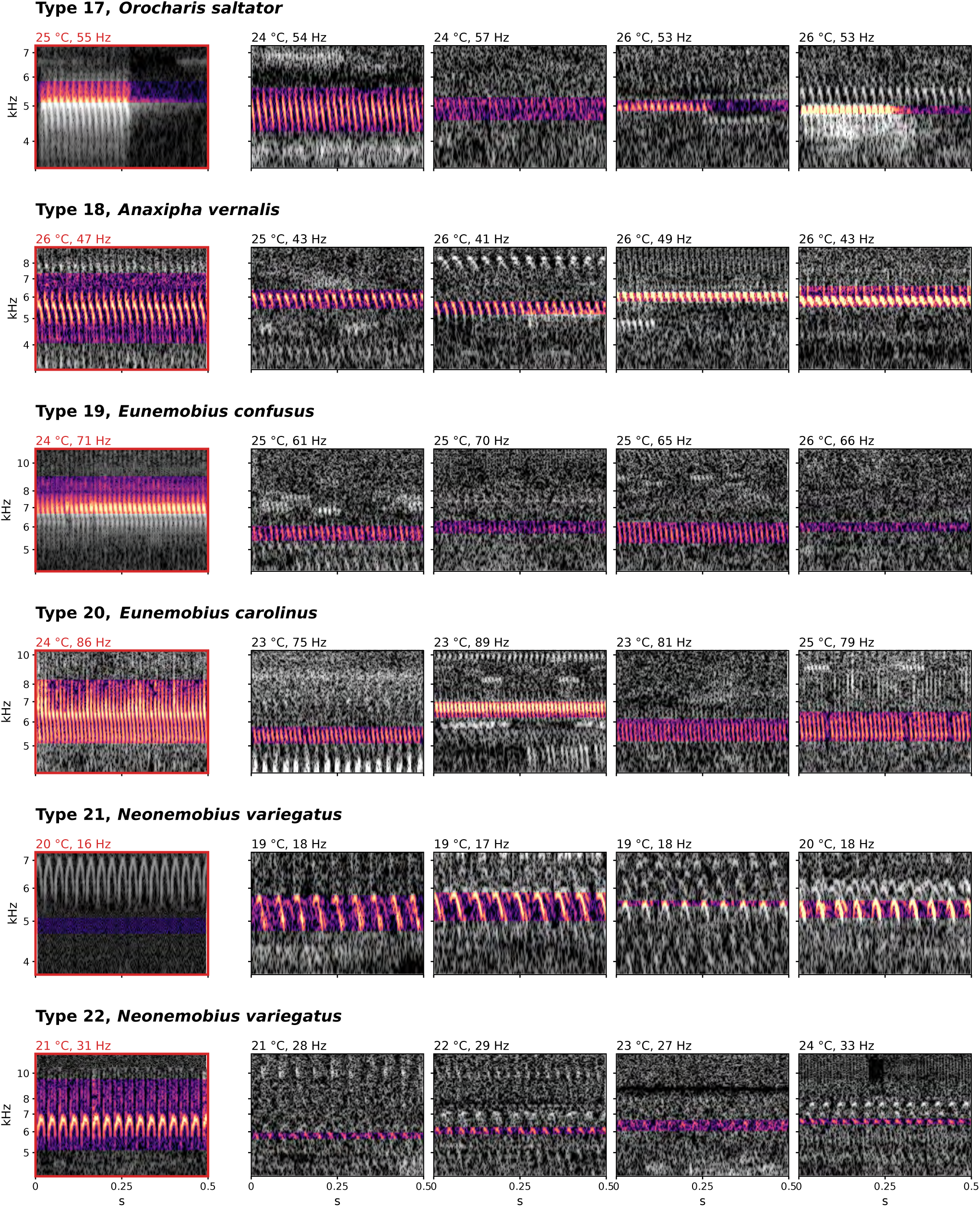
Reference recordings beside field singers of the type they match (3 of 7)

**Figure 14.4.**
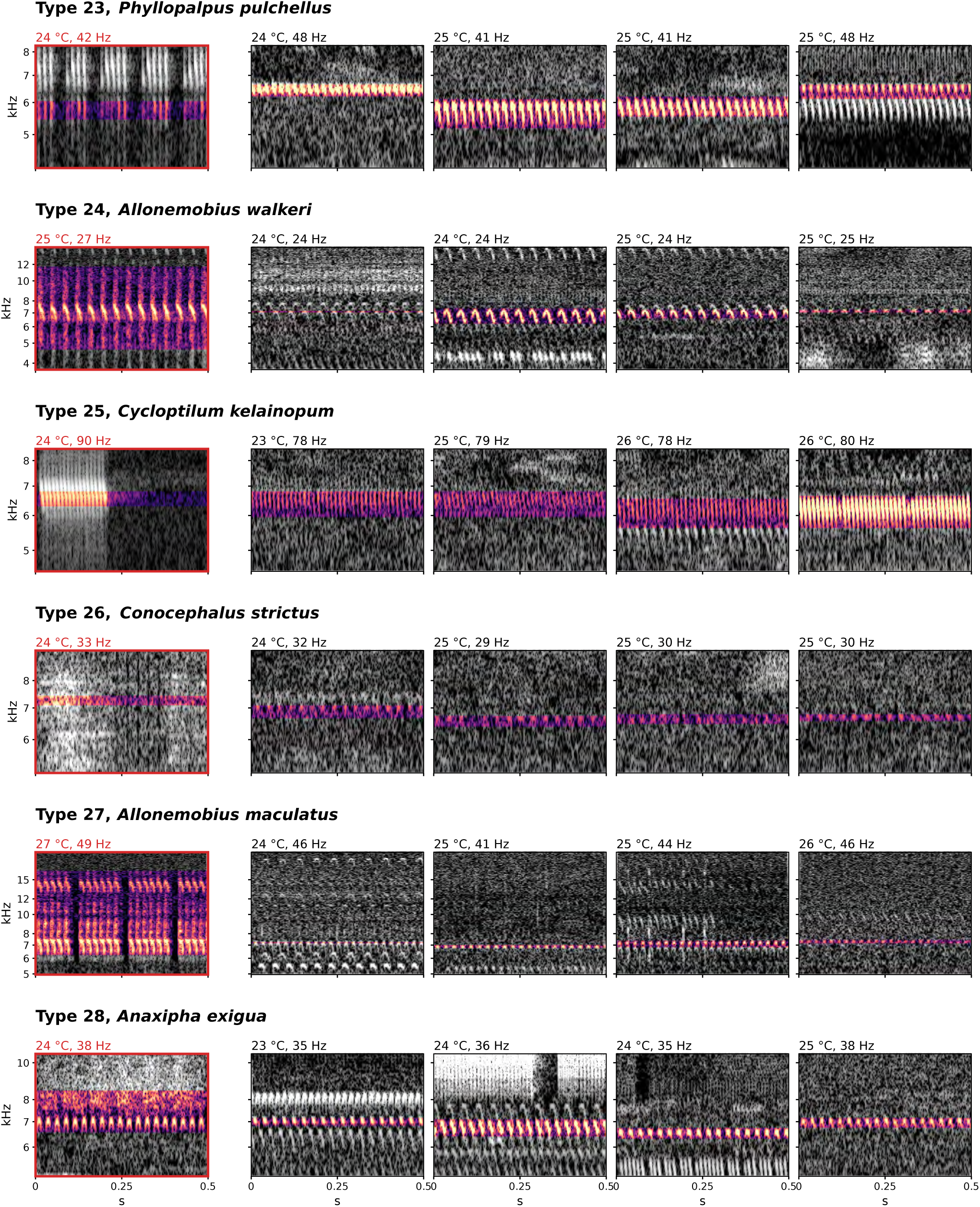
Reference recordings beside field singers of the type they match (4 of 7)

**Figure 14.5.**
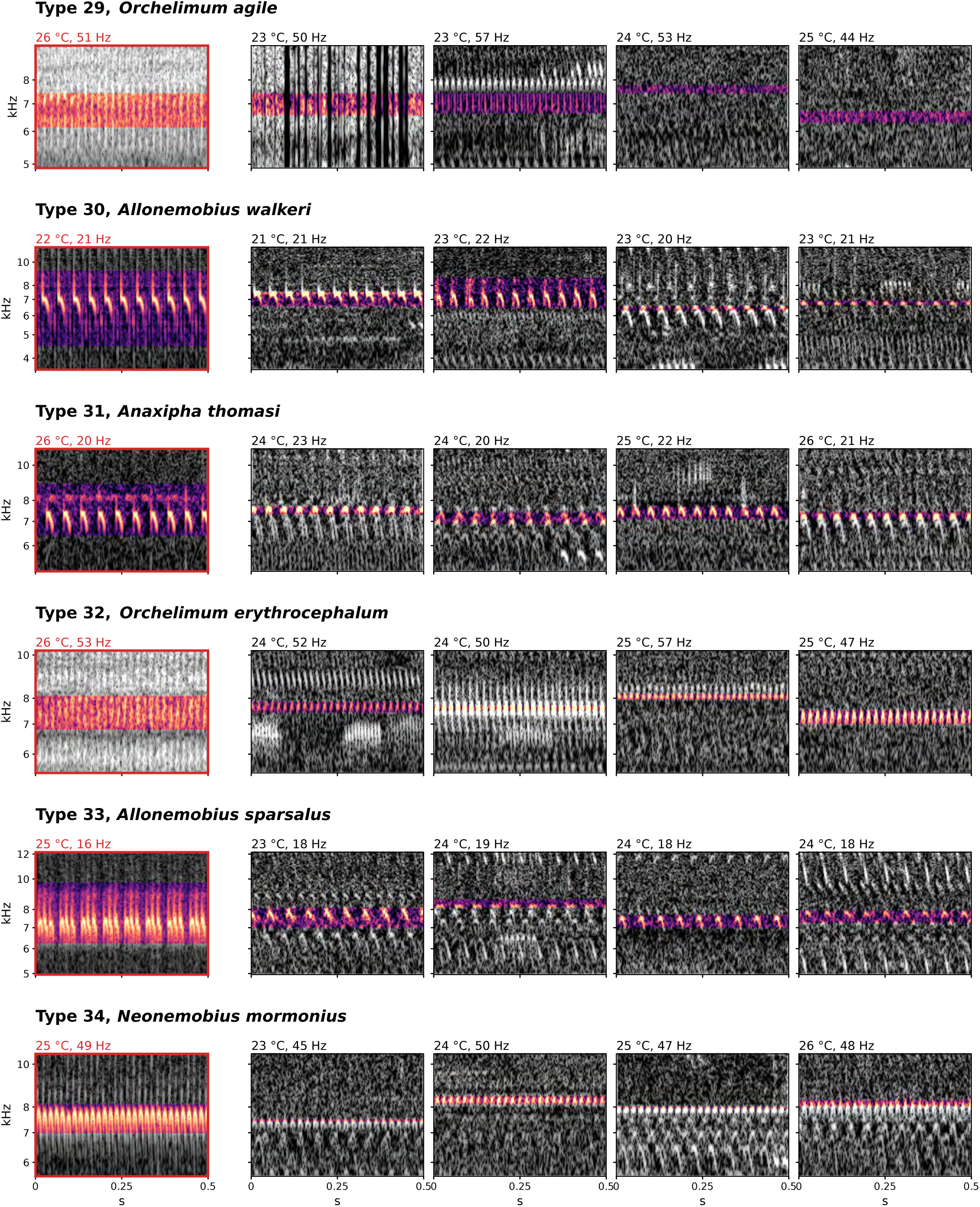
Reference recordings beside field singers of the type they match (5 of 7)

**Figure 14.6.**
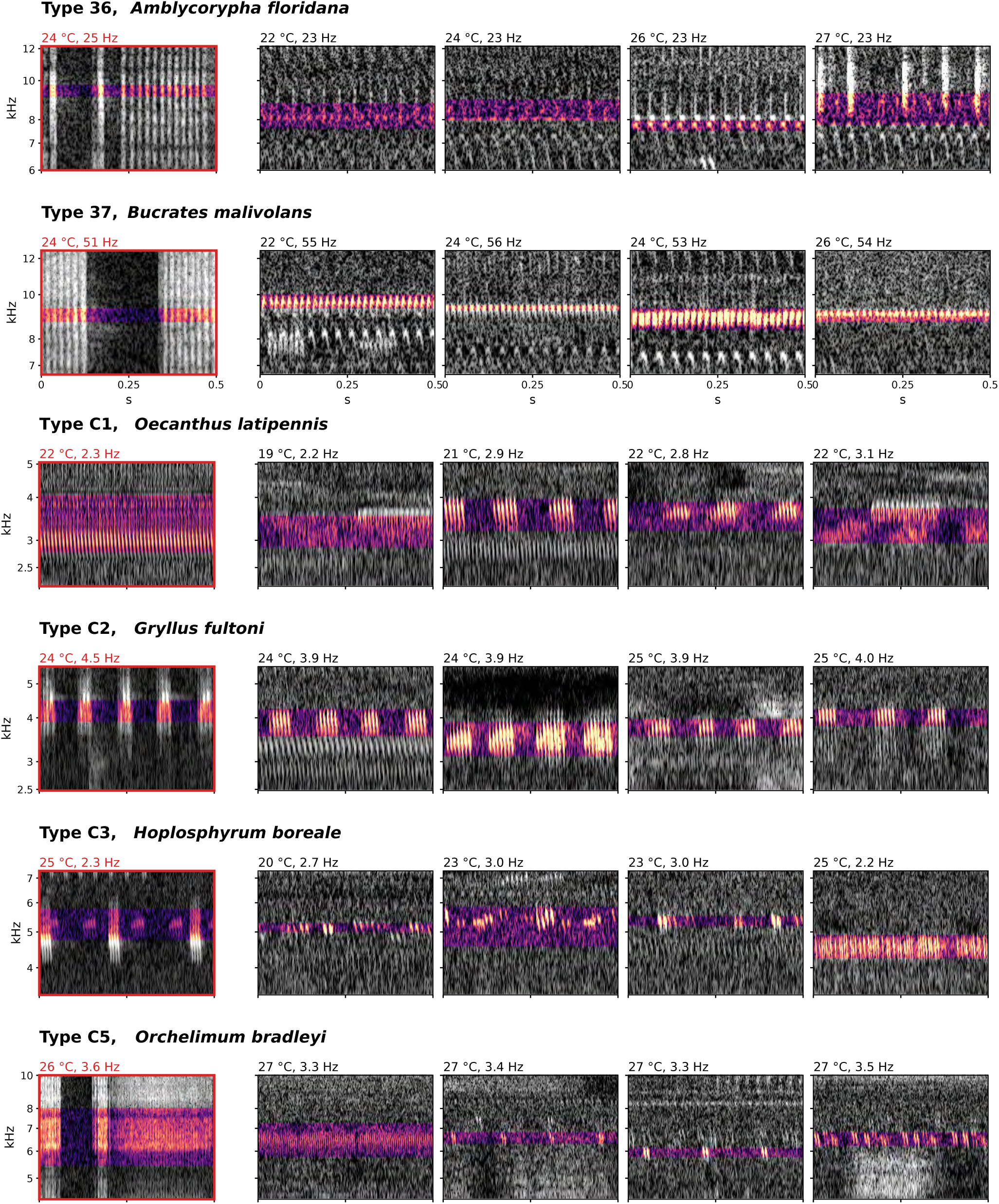
Reference recordings beside field singers of the type they match (6 of 7)

**Figure 14.7.**
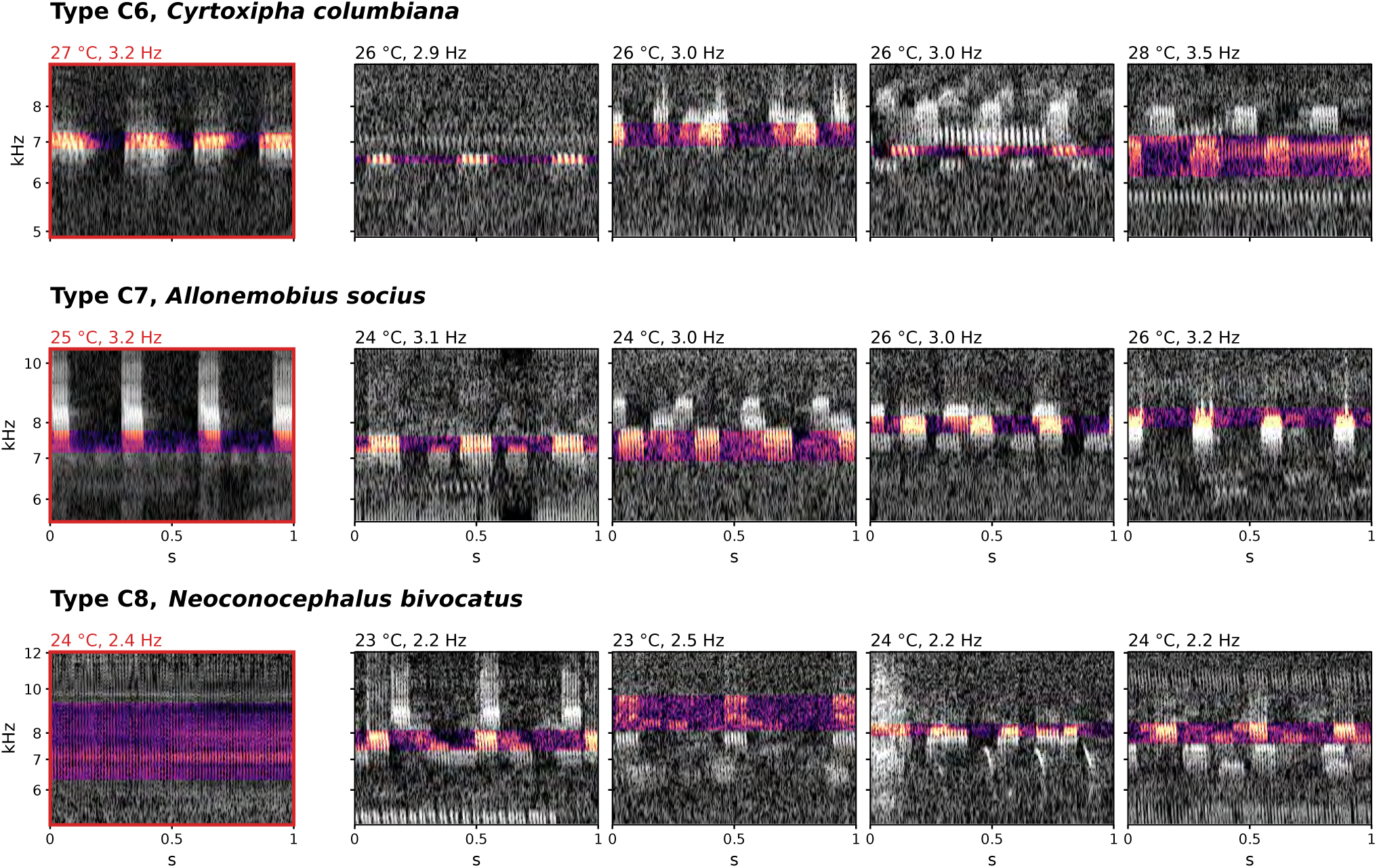
Reference recordings beside field singers of the type they match (7 of 7)

#### Songs during daytime

Although orthopteran songs are primarily nocturnal, we ran the complete pipeline on the daytime recordings, and placed tracks on the night catalog’s lines where they fall on one. Figure 15A shows the 27,671 day singer-hours of 5,442 tracks on the same plane as Figure 7. Most of the day singing comes from types that also sing at night (Figure 15B), but three lines above 9 kHz sing 88 to 94 percent of their singer-periods by day (Figure 15D) and so barely register in the night catalog: type 37 (9.5 kHz at 58 Hz), with 64 night singer-periods against 481 by day, and two lines at 10.2 and 10.7 kHz with 18 and 11 night singer-periods against 276 and 93 by day, too few at night to be night types at all. A fourth, at 8.2 kHz and 45 Hz, sings 80 percent of its periods by day. Three fifths of the 5,427 day singer-periods fall on a night line; the rest, the gray in Figure 15B and C, are day songs the night catalog has no line for, and we will build a day catalog to type them in a future version of this analysis.

**Figure 15.**
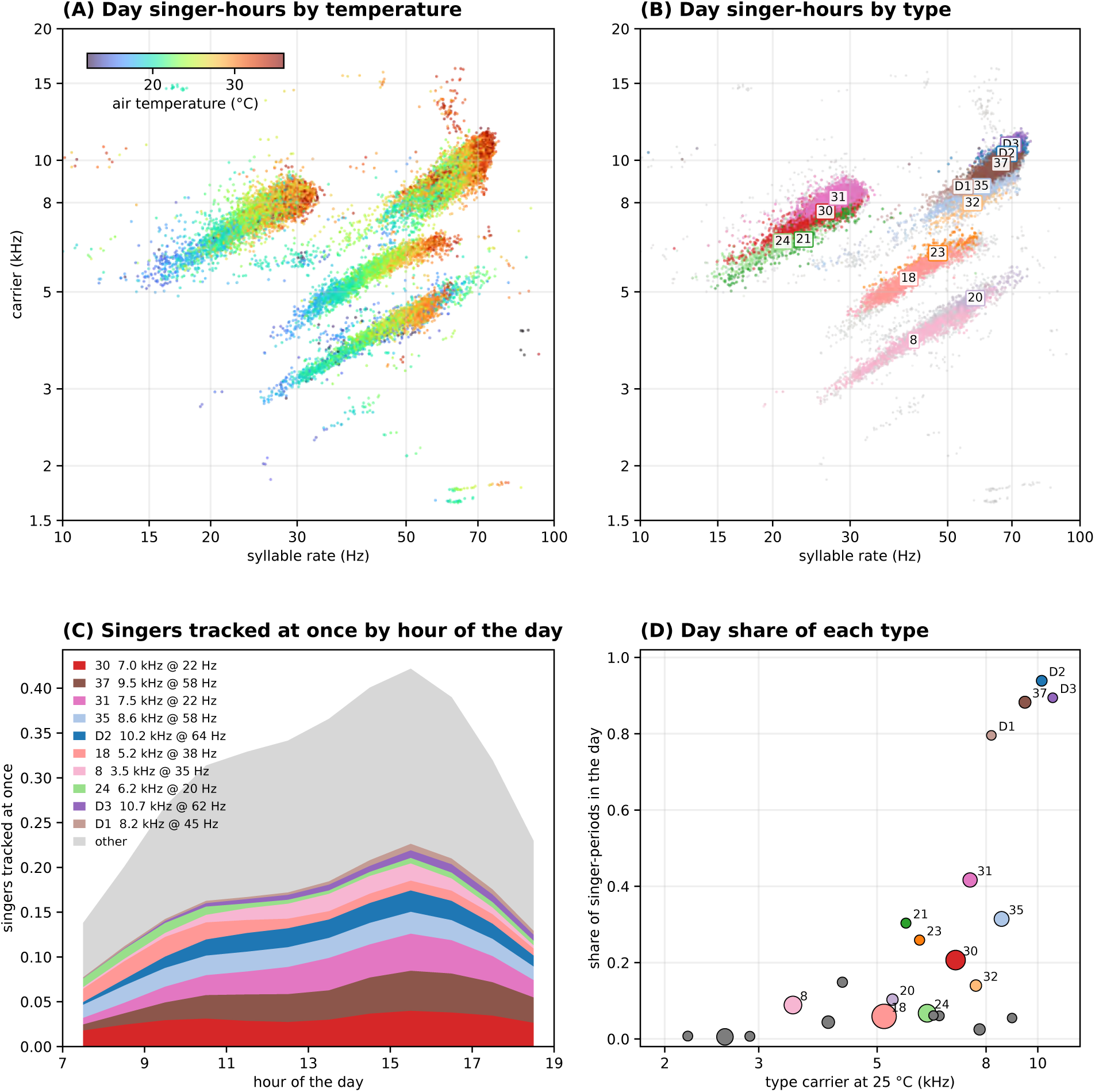
The day singers (A) Day trillers on carrier frequency x syllable rate plane. (B) Most overlap tracks from night. New tracks labeled with “D”. (C) Daily activity levels of day singers. (D) Fraction of songs of type in day vs night.

With the day singers added to the catalog, we can look at circadian rhythms in calls by type (Figure 16). As expected, the majority of types peak at night, with only type 37 peaking during the day.

**Figure 16.**
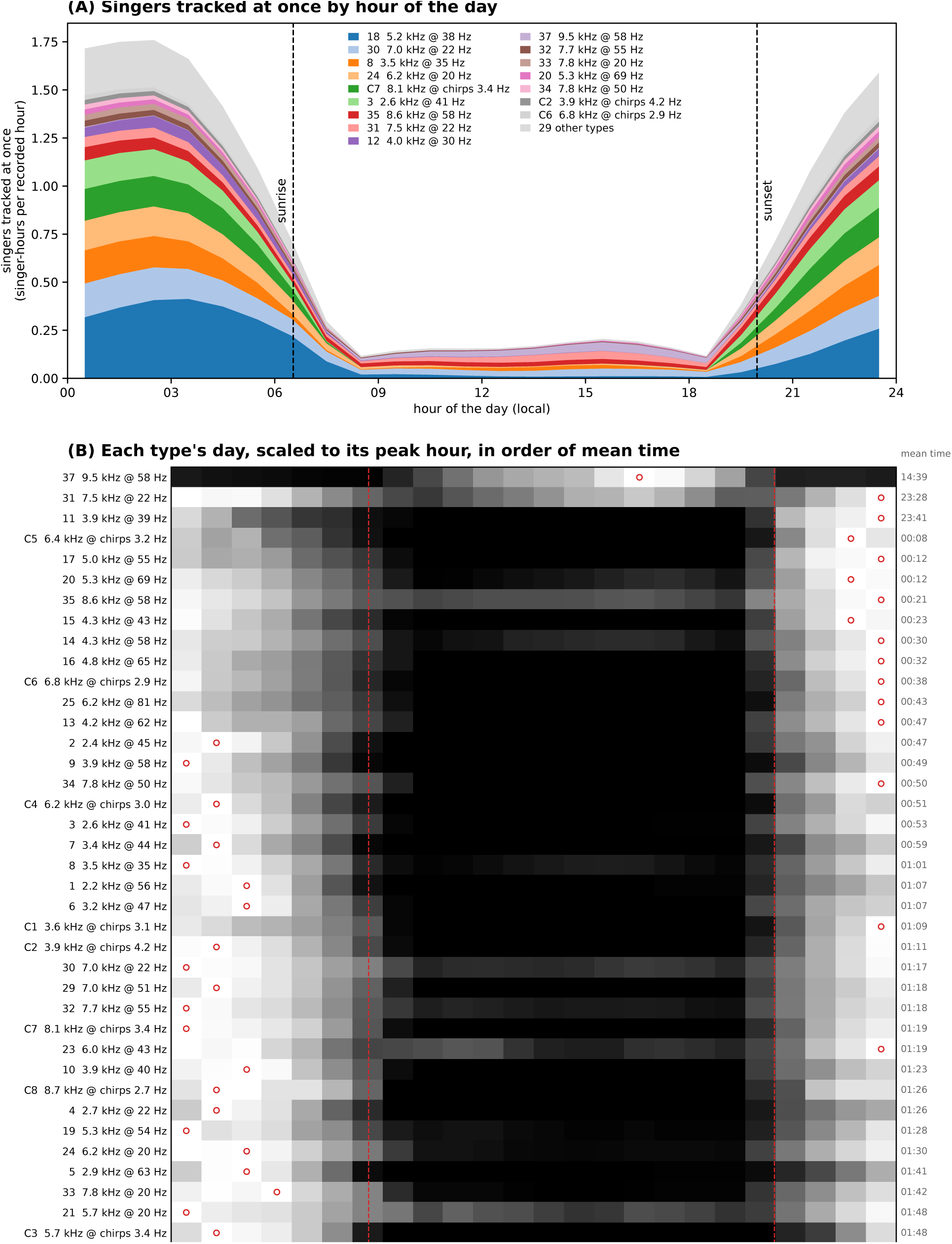
The types across the 24 hours of the day (A) Top singers tracked by hour across year. (B) Hourly patterns of dominant types with time of peak activity shown.

## Discussion

Our proximal purpose in developing and applying the approach described here was to see if it was practical to characterize the rich and fascinating and largely unannotated orthopteran chorus we could see in the roughly 160 TB of audio data we had collected to characterize birds. It is!

We have shown that five-minute windows from one recorder can be automatically resolved into its singers by projecting the data into the carrier frequency-rhythm plane; that individual singers can be tracked for hours across a night or day; that the accumulated tracks across a year resolve into dozens of narrow parallel lines spanning a wide range of carrier frequencies and syllable and/or chirp repeat rates; that these “streaks” can be readily resolved to distinguish several dozen distinct trilling and chirping types; that singers within these types sing the same song as they progress up the streak with increasing temperature in a predictable and repeatable way, from slower low-pitch songs on colder nights to faster higher-pitched songs when it is warmer. None of this used or requires reference recordings or a training set, though these can then be used to give names to each of the types.

That orthopteran songs move along such lines is not new. Brooks described it in 1881, Walker measured it species by species from the 1950s, and Walker and Funk drew exactly the same lines for six species of *Anaxipha* from 792 hand-made recordings. Indeed the concordance between our data and Walker and Funk’s *Anaxipha* is one of the things that gives us confidence we are on the right path here. What is new is the scale and the direction of inference. Walker and Funk’s lines were drawn for known species from single males recorded beside a thermometer. Ours emerge from 139,000 singer-hours of unattended recording, and the species, once they have been assigned, will have been inferred from the line rather than the line from the species.

Because this macroscopic, type-level view, is built progressively from 5 minute windows at specific recorders, it gives for every type, when it sings, where, at what temperatures, by day or night, with what syllable, and how its carrier and rate are linked, which can provide invaluable information about the species making the songs, its physiology, behavior and ecology, while also providing an important window into the state of the specific locations where each recorder is located as well as the larger ecosystems in which they are placed.

### Who are the singers?

Our analyses identify 45 distinct types that we recorded repeatedly across the analyzed year. This is a measure on its own of the orthopteran biodiversity at the site. But the goal is clearly to be able to put a species name on every one of the types, and to systematize the process for doing so. While the analyses we present in Figures 12, 13 and 14 give us confidence that this can be done, there are two major challenges to doing this fully. First, it still is not completely clear what our standards should be for confidently establishing a match. And second, the amount of high quality, annotated reference data remains limited.

The *Anaxipha* data are a best case scenario for now - many recordings for each species at a range of different temperatures. This allows us to not just try to place individual recordings on the type map, but to compare annotated types from the reference data to our types. This suggests a mapping of *A. vernalis* to type 18, and *A. exigua* to type 27, and both species have been recorded in Arkansas. However, the fact that two coastal species (*A. litarena*, and *A. delicatula*) and one whose range is not known to extend into Arkansas (*A. rosamacula*) would, by the same criteria, be assigned to types, merits caution.

Similarly, many of the candidate matches shown in Figures 13 and 14 appear reasonable on acoustic and geographic bases, but, especially in the absence of data from a range of temperatures, we are not confident enough to name names. Clearly this would benefit from input from someone with expertise in identifying these songs, which we will pursue as this work moves forward.

### Syllable shape

One feature of songs that we have not fully captured is the syllable shape - the way that the power sweeps across frequencies during a syllable - would likely help in refining both type segmentation and identification. Using terminology borrowed from birdsong description, we see at least five distinct syllable shapes (downslur, upslur, chevron, hockey stick, flat), examples of which are found in Figure 17, which displays a representative example for each of the 45 named types. The syllable types are mostly consistent within a song, and (by and large) among singers within a type (Figures S7-{type}). Indeed the consistency of syllable shape within a type is one of the reasons we are confident that they represent the songs of one biological entity.

**Figure 17.**
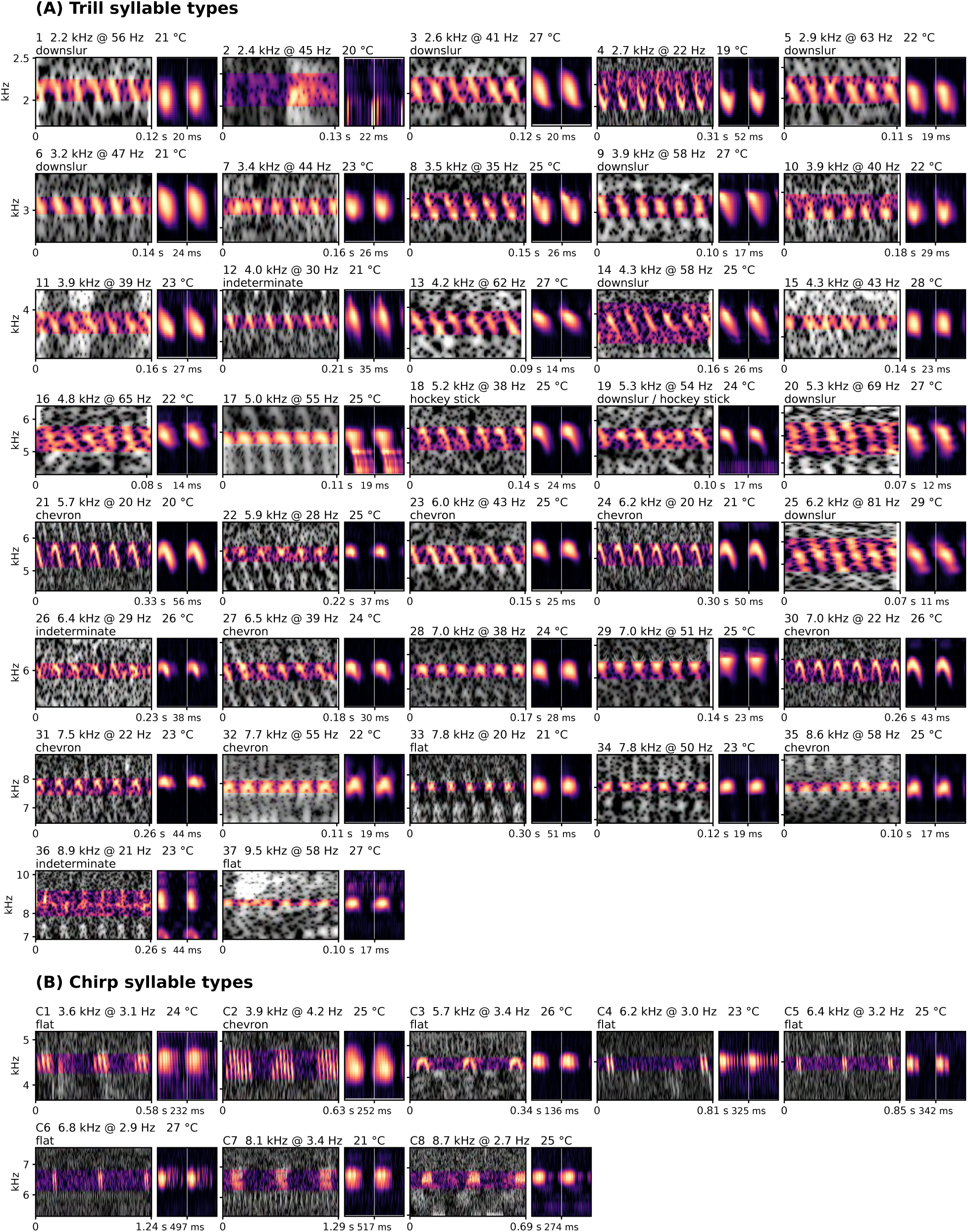
Syllable types across song types

The syllable shape labels in Figure 17 were done by eye based on appearance alone. We have been experimenting with ways to automate this classification, but are not satisfied with it yet, so only manual annotations of a small sample are included here. But we anticipate integrating syllable shape classification into the pipeline moving forward.

### What do the streaks tell us?

The work of Walker and Funk (2014) on North American *Anaxipha* primed us to expect to see syllable and chirp repeat rates as well as carrier frequency increase with temperature, and for singers of a species (or type) to fall along a compact tract in the carrier frequency – syllable repeat/chirp rate plane. As others have noted before, the fact that these data do fall along compact, separable lines says that the way that the syllable (and where it is present chirp) rate varies with frequency and both of these with temperature is a species/taxa specific character.

One way to think about the streaks on the plane is in terms of two temperature responses, that of syllable rate and of carrier frequency. To measure each on its own, and within singers rather than across them, we took every track of a type with at least four hours and a two-degree range of temperature within its night and expressed each of its hours as a deviation from that track’s own mean temperature and its own mean rate or carrier, in percent. Pooling these deviations over a type’s tracks gives the within-singer slope of the type (Figures 18 and 19; Table 1 gives the slopes, the interquartile range over singers, and the number of singers behind each). The same regressions run on the raw singer-hours give larger and more variable slopes, because they also take in differences between the singers that happen to sing on warm and on cold nights; Table 1 lists those too.

**Figure 18.**
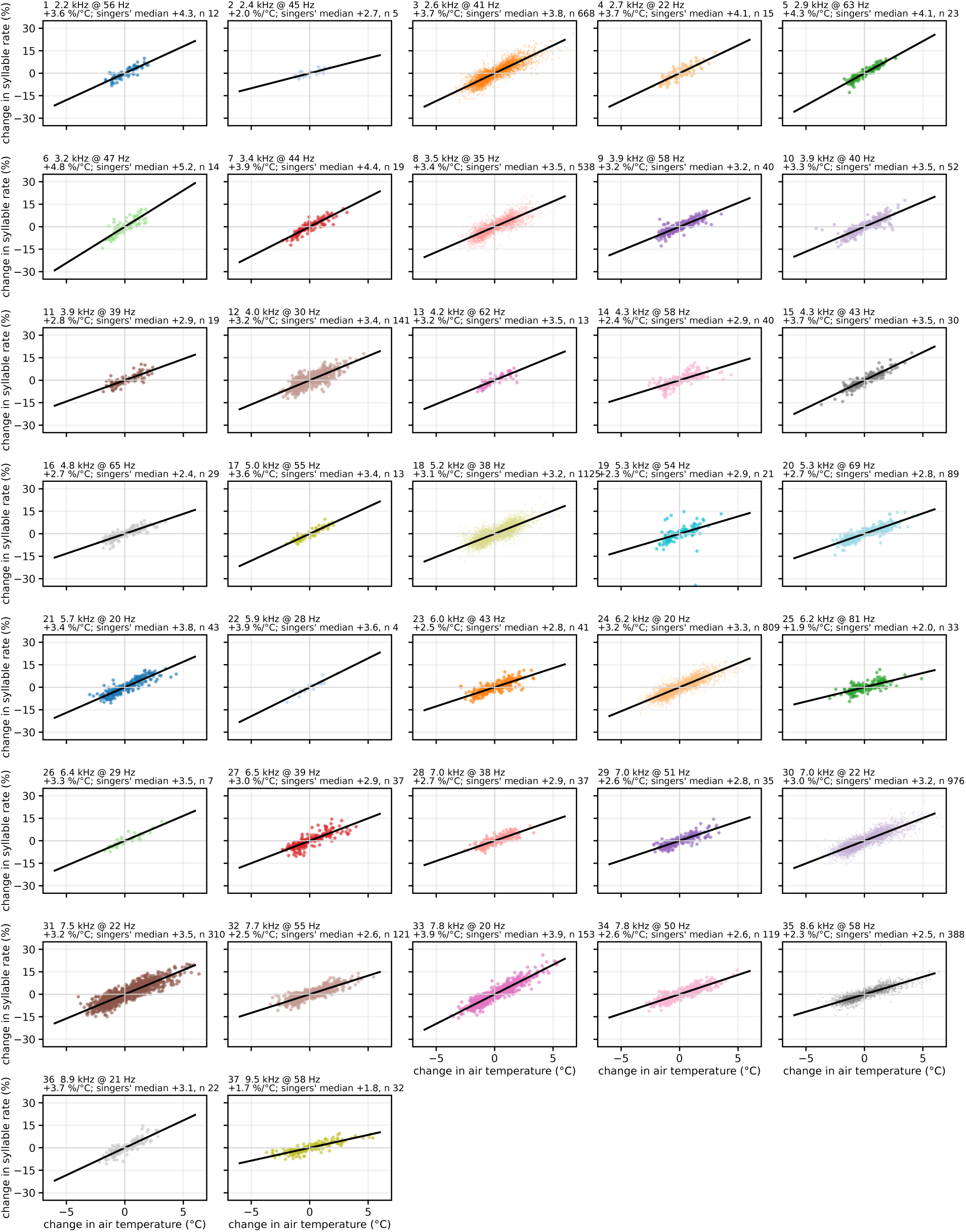
Syllable rate against air temperature within singers, type by type

**Figure 19.**
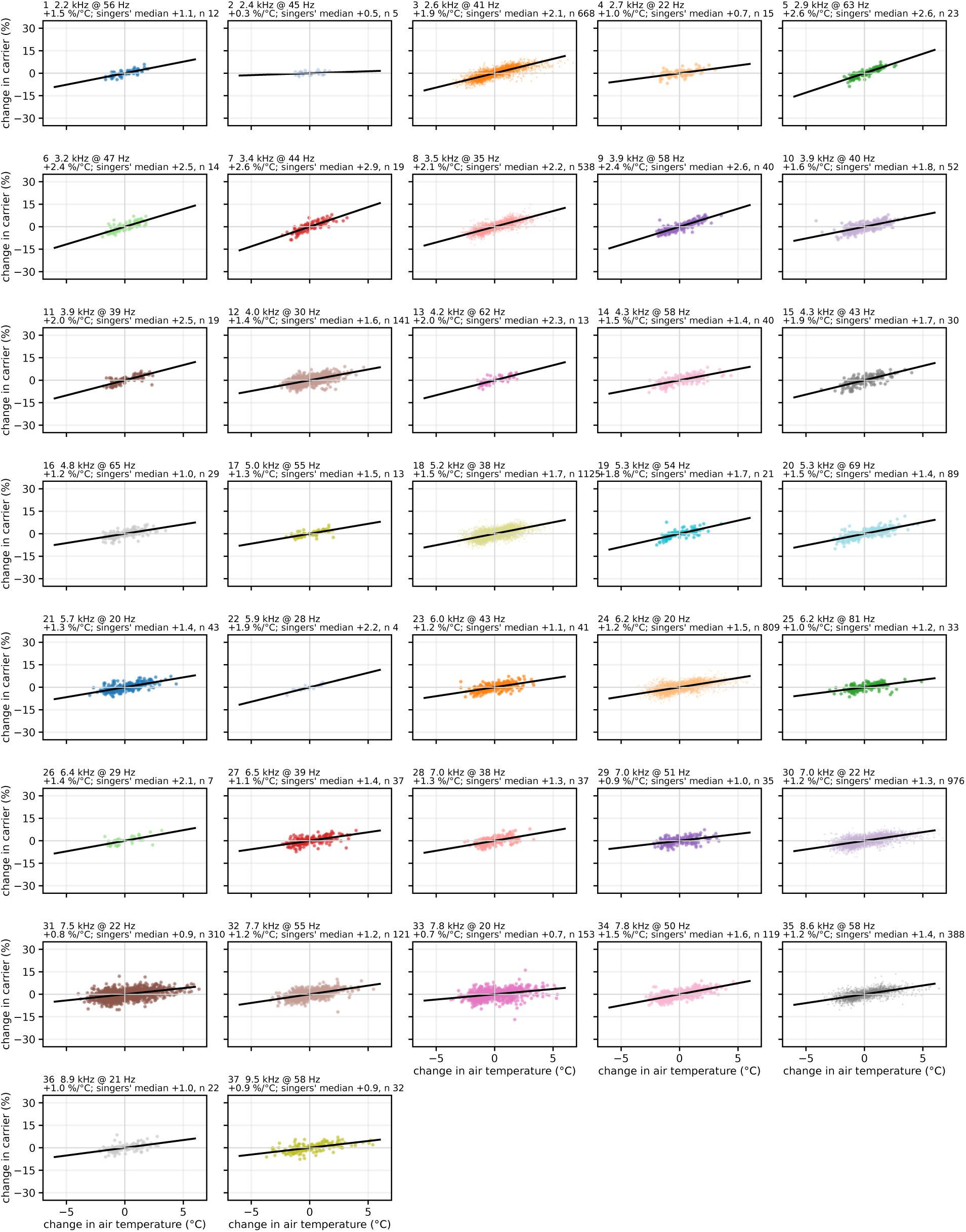
Carrier frequency against air temperature within singers, type by type

Within singers, the syllable rate rises with temperature in every type with rate of change of 1.7 to 4.9 percent per degree (median 3.2, and 30 of the 37 types fall between 2.3 and 4; Figure 18). The carrier also rises in every type, but by less and more variably: 0.3 to 2.6 percent per degree, median 1.4 (Figure 19). The ratio of the two slopes within a singer is the proportional response of carrier to rate, 0.48 in the median type. It is 0.5 to 0.8 in the types below 4 kHz and in several of the trills above 7 kHz, and falls below 0.3 in five types: 2, 4, 31, 33 and 36, in which the rate speeds up with the night’s warmth as everywhere else while the carrier barely moves.

These details, and especially the differences, in these responses likely reflect the different anatomical mechanisms used by different groups of orthopterans to generate their song, and are hard to fully deconstruct in the absence of type identifications. For the same reason, they also are likely taxonomically informative characters, and will be integrated into the naming step as we move forward.

### Why has relatively little attention been paid to carrier frequency variation relative to syllable/chip repeat rates?

The earliest observations about the temperature dependence of orthopteran calls focused on chirp rates, because those can be heard by ear. Dolbear’s cricket could be timed with a watch (Dolbear 1897), and for a century the pulse rate was the character that separated sibling species and the character whose match to female preference was studied under “temperature coupling.” Carrier frequency was harder to measure well until cheap digital spectrograms, and when it was measured it was usually reported as a single figure per species. The resonator model of cricket sound production (Bennet-Clark 1999; Bennet-Clark and Bailey 2002) then gave a reason to expect it to be constant: if the harp sets the pitch, temperature should move the rate and leave the carrier alone, and in the field crickets that were the model animals this is nearly what happens. The tree crickets, whose carrier does move (Sismondo 1979; Mhatre et al. 2011), were treated as the exception, with Walker’s work on *Oecanthus* the standard reference (Walker 1962a; Walker 1963).

Another part of the answer is that the relation is weaker in any one recording and only becomes a line when many recordings are placed together. A single animal at a single temperature gives one point; a laboratory series over temperature gives a short segment of its own line; a species line like the ones in Figure 7 needs hundreds of singers across a season, each with a temperature attached, which was not practical before continuous recording. Walker and Funk’s (2014) study of *Anaxipha* is the closest precedent we know: over 1,300 recordings of 13 species with temperatures, and the carrier and pulse rate of each. Their recordings fall on our lines (Figure 12), and they drew the same conclusion from them that we draw here, that carrier and pulse rate vary together within a species in a way the standard resonator account does not predict (see also Prestwich et al. 2000). Their data have been available for a decade. What has been missing is a way to produce such data for a whole assemblage rather than one genus at a time, which is what we have tried to supply.

Our own data add two things to this. The first is a correction to how we read the relation. The streaks on the plane are nearly parallel on log axes, and we were at first inclined to treat that as a finding. It is better seen as a ratio: within a singer the rate rises by about 3 percent per degree in every type we have, and the carrier by about 1.5 percent, so the slope of a type’s line is the ratio of its carrier’s thermal sensitivity to its rate’s, close to one half, and the parallelism says that this ratio varies less between types than either sensitivity alone. On linear axes, the ones Walker and Funk used, the same lines have slopes that differ several-fold because they scale with each type’s carrier-to-rate ratio.

As discussed above, the slope measured within a singer through its night and the slope of the type’s streak across the year are different quantities, and they do not always agree. In most types they are within a tenth of one another, but in five types the carrier barely moves while the rate follows the night’s cooling, and the streak nonetheless rises at the common slope. For those types the streak’s slope must come from differences between singers, and we find that singers of one type recorded at the same hour on the same recorder, at one temperature, differ from one another along a line of about the same slope. What distinguishes one singer of a type from its neighbor moves carrier and rate together in nearly the same proportion as temperature does. We do not know what that variable is (body size is the obvious candidate).

### What we learned about the method

Most of what we learned came from the failures, which we have kept in the record that accompanies the code. The modulation map is a good detector because it is relative: a faint singer that switches fully on and off scores as strongly as a loud one. For the same reason it is a poor judge of what a singer is, and every attempt to decide identity from the map alone, by the pattern of harmonics, by tests on the modulation spectrum, by a statistical test of whether a peak was a harmonic, invented animals or merged them. What worked was to read everything that is a property of the animal from the audio itself: the syllable rate inside a chirp from the envelope, the song form from whether the syllables follow the bout rhythm, the syllable’s shape from a fine fold at the refined rate. The second lesson was that greedy decisions do not survive a chorus. Linking window to window shredded tracks that a single assignment over the night kept whole; clustering with a tolerance that fit one season cut the long lines into temperature segments; and a chirp plane built from everything that pulsed below 8 Hz mixed crickets with frogs until we required that the syllables be readable. The third lesson was that the quality control is the grids. Looking at twelve positions by five singers of every type, with the audio behind each cell, found every error we have found, and we would not trust a type we had not looked at that way.

### Methodological Limitations

We succeeded in our overarching goal of being able to systematically extract orthopteran songs from years-long continuous audio recordings and to catalog the organization of this data across taxonomy, time and space. However, several obvious (and likely non-obvious) limitations are apparent.

The data look good (songs of a type look similar), and the agreement with hand measured and analyzed *Anaxipha* data as well as the broader literature serves as a sanity test. However, we haven’t done any kind of formal analysis of the sensitivity or specificity of the method at any time scale. This is almost entirely due to the lack of curated datasets against which to do this evaluation. We are very interested in pointers to datasets that might serve this purpose.

The assignment of frequency bands to an individual singer in a 5 minute window is heavily influenced by the intensity of the song (which is presumably related to distance from the recorder), the presence of other singers in nearby frequencies, and noise. This means that what appears to be the same song from the same singer can sometimes be fully captured in one window, and reduced to a few adjacent frequency rows in another. This may have effects on the way we extract numbers from each singer, but it definitely complicates our efforts to use syllable shape as a character both in taxonomy and quality control. We are very interested in developing methods to extract full syllables more reliably.

The method is designed to detect songs that have a high degree of regular, internal repetition, but not all orthoptera calls are like that. The detector reads rhythms between 2 and 100 Hz, so coneheads and other very fast tickers, and chirpers slower than two chirps a second, are outside it by construction. Furthermore, we made some choices in this version of the analysis that created some blind spots, especially with regards to trillers with syllable repeat rates below 10 Hz. The latter will be addressed in future versions of this analysis, the former will likely require a different approach.

### Data Limitations

For a variety of reasons, some having to do with equipment failures, but most having to do with our ability to go to the field site to collect SD cards, we had uninterrupted full year coverage for only a handful of recorders. But even if we did, the range of orthopteran calls is such that even the best possible arrangement of 27 recorders would not come close to capturing the full complexity of the orthopteran songs at our site. These recorders were positioned to capture bird songs primarily - which is why they are spaced at least 25m apart. As we begin to deploy a new generation of recorders, it is an open question how best to position them for orthopteran call analysis.

While our analyses don’t depend on highly accurate temperature measurements, having to rely on modeled temperatures with limited spatial resolution is a design failure on our part. We have now deployed a network of weather stations (METER group), one at each of the main recorder deployment sites, which will provide much better data going forward.

A particularly challenging aspect of the data is that there are a lot of orthopterans singing at this site. There are many time windows during times of year when orthopterans are most active when there are overlapping calls of both similar and very different types. Of course this is the most interesting time, but we are also interested in seeing how this method will perform on less complex soundscapes.

### From Soundscapes to Systematic Acoustic Inventories in Ecology

An acoustic inventory that needs no training set can be run anywhere orthopterans sing, which is most of the ice-free world, and it produces the things ecology needs from such an inventory: which types are present, when, where and in what numbers relative to one another. The measure we use, singers tracked at once per recorded hour, is comparable across recorders and years because every singer is found the same way. Given the large number of autonomous recording units already deployed to monitor birds, the application of acoustic inventory analysis to orthopteran ecology, and via orthoptera to other aspects of biodiversity and ecosystem state, was waiting for a tool to enable it. While there is a long way to go to perfect the approach we have presented here, we believe the ideas we have explored here, and the software we have developed are good enough to begin to pursue these goals at scale.

### Detailed Methods

#### Recordings and site

Recordings were made on a 400 ha property in Hempstead County, southwestern Arkansas, USA (33.75° N, 93.48° W), a mosaic of former pasture, creek-side woodland and ponds that has been under forest restoration since 2022. Twenty-seven Frontier Labs Solar BAR autonomous recorders at eleven stations recorded continuously at 44.1 kHz, 16-bit, mono, in hour-long files. Four stations carried several recorders a few tens of meters apart for a separate bird-localization study (Back9, seven; NorthSide, five; EastCreek, four; House, four); Copse, FarNorth, MidCreek, Pond, SecretGarden, WestSideJunction and, from October 9, 2025, EastMeander carried one each. The analysis covers November 1, 2024 to November 30, 2025. Each recorder’s calendar day was divided into a night, 19:00 to 07:00 local time, and a day, 07:00 to 19:00, and a recorder-period entered the analysis when at least six of its twelve hour files existed: 7,321 recorder-nights and 7,320 recorder-days, 87,935 recorded night-hours and 87,961 recorded day-hours, in all 175,896 hours were analyzed in five minute windows.

#### Temperature

Air temperature at 2 m above ground was taken from the National Oceanic and Atmospheric Administration’s High-Resolution Rapid Refresh model (HRRR, 3 km grid) at the grid cell containing the site, retrieved as hourly values through the Open-Meteo historical-forecast interface (Dowell et al. 2022; Zippenfenig 2023) and interpolated linearly to the center of each five-minute window. Five weather stations installed at recorder stations after the recording period (ATMOS 41, METER Group, from 9 September 2026) show that the gridded product runs about 2 °C warm on clear nights at this site with an hourly scatter of about 2 °C. The temperature slopes reported here are therefore slightly underestimated, and the along-line scatter of the types includes thermometer error. Where a night’s temperature is summarized it is the mean or minimum of the hourly HRRR values between 19:00 and 07:00.

#### Overview

The analysis has four stages, each reading only the output of the one before. A five-minute window of one recorder is decomposed into its singers in the plane of carrier frequency against amplitude-modulation rate, without reference recordings or training data. The windows of one recorder-period are linked into tracks, one per singer, by a single assignment over the whole period. Tracks lasting two hours or more are pooled over the year and clustered into call types, each a line in the carrier-rate plane after correction to a common temperature. Each type is then described (season, thermal range, sites, song form, syllable shape, carrier-rate coupling, chirp structure) and compared with reference recordings placed on the same plane by the same measurement.

#### One window: the modulation map

A window is 300 s of audio, placed on a fixed clock grid every five minutes and read from the hour file that covers its center. Its short-time Fourier transform uses 512-point Hann frames with a 128-point hop (86 Hz rows, 2.9 ms frames). For every carrier row above 300 Hz the time series of level in decibels is de-meaned and its Welch spectrum taken (2048-point segments, 0.17 Hz resolution) over modulation rates of 2 to 100 Hz. A running median over 1.4 octaves of log rate is subtracted from each row’s own spectrum, so that a row that is merely noisy scores zero at every rate and a row modulated at rate R scores at R and its harmonics. The result, the excess modulation power as a function of carrier frequency and modulation rate, is the modulation map. Because it is built from each row’s level relative to its own mean, a faint singer that switches fully on and off scores as strongly as a loud one; the map is not weighted by energy. The lower limit of 2 Hz lets a singer whose strongest rhythm is a chirp repeated a few times a second appear at that rate rather than only through its harmonics.

Peaks are connected regions of the map more than 6 dB above baseline that span at least two carrier rows. Within a region, rows are grouped by the rate bin of their own maximum and the region is divided wherever consecutive rows’ maxima differ by more than two bins (0.34 Hz), since two singers less than a hertz apart in rate otherwise share one region. Each peak’s rate is first taken as the bin of its strongest cell and then refined to about 0.01 Hz by maximizing the contrast of the spectrogram folded at that period: over the loudest 30 s of the peak’s rows for a singer present in more than three quarters of the window, and otherwise over its runs of song of at least 1 s, each folded with its own phase and the runs aligned on their envelopes before they are summed, because a singer that sings in bouts resets its phase at every bout. The refined rate is checked against its sub-multiples R/2 and R/3: the fold at the lower rate replaces R when it is at least 1.25 times deeper, the case of a singer whose pulses alternate in strength or whose chirp rate was first read at a multiple; the check is iterated up to three times. A peak below 8 Hz that shares rows with a faster peak is tested as that singer’s bout rhythm: when the envelope of the fast modulation, folded at the slow rate, falls below 0.4 of its peak, the slow peak is the fast singer’s pattern of chirps and is recorded as its chirp rate rather than as a second singer.

#### Parts and singers

A peak can hold pieces of two singers at nearly the same rate in adjacent frequencies. For each row of the peak and 300 Hz of padding on either side, two features are computed: the syllable profile, the fold of the row at the refined period, scaled to unit peak after removing the row mean; and the singing pattern, the amplitude of the row’s modulation at the refined rate in each 0.5 s of the window, high-passed with a 10 s running median so that slow common trends do not make unrelated singers co-vary. Rows are walked from low to high frequency, and a new part begins where a row’s profile correlates below 0.3 with the previous row’s, or where a row with a bout pattern does not follow the part’s. Each part is seeded by rows with modulation depth of at least 0.15 relative to their own power and then extended row by row into the padding while the row’s absolute in-phase amplitude stays within 15 dB of the part’s peak, the row is modulated at all (depth at least 0.02) and its profile still correlates at 0.3 with the row it extends from; the second stage recovers rows that a louder steady neighbor had hidden from the threshold.

Parts at a rate m·R (m = 2 to 6, within 1.5 %) within 300 Hz of a part at R are the same pulses seen at a harmonic of the pulse rate and are absorbed into it without a test, since any repeating pulse produces them. Remaining parts are joined into one singer when their singing patterns correlate (r > 0.2) and the correlation exceeds the distribution obtained from 60 circular shifts of one series by more than 4.5 standard deviations. Carrier harmonics (n·F at the same rate) join through this test; combination harmonics (n·F, m·R) nested within an existing component are absorbed structurally only when both parts are continuous (singing in more than 75 % of the window). Every part is tested against the singer’s lowest-frequency component alone, never against the other components, so two singers cannot be merged through a third part that resembles both. For each singer the window records its component bands, the centroid of in-phase amplitude over the rows of its lowest band (so that neighbors’ unmodulated energy does not pull it), its refined rate, chirp rate where it has one, modulation depth, the fraction of the window in which it sang, and a per-row profile of rate-locked amplitude used later to measure the carrier over a fixed frequency window. A window takes about 8 s to analyze.

#### Chirpers

A singer that groups its syllables into chirps appears in the map at its chirp rate (2 to 8 Hz), with a weak, broadened peak at the syllable rate inside the chirp that a continuous neighbor in adjacent rows often masks. The syllable rate is therefore read in the time domain. The audio is band-passed to the singer’s band and its envelope taken; chirps are the events in which the envelope, smoothed over 40 ms, clears a threshold between its quiet and loud levels for at least 30 ms, and the pulses inside each chirp are the peaks of the envelope smoothed over 3 ms. The syllable rate is the reciprocal of the median interval between pulses, kept only when the intervals are regular (interquartile range below 35 % of the median, over at least 20 intervals in at least 5 chirps) and the rhythm’s power inside the chirps is at least twice its power between them. Where the chirper’s syllable peak was also found as a separate singer in the same rows, the two are one singer and are merged (rates within 6 %). When no regular pulse train is found the singer is recorded with its chirp rate alone. Chirp rates below the map’s 2 Hz floor are not measured, so the slowest chirpers on the coldest nights appear only at their syllable rate.

#### Flags before linking

Three kinds of detection are flagged and set aside before linking; nothing is deleted. Two singers in one window whose carriers stand in a 2:1 or 3:1 ratio (within 8 %) at the same rate (within 3 %, or 6 % on chirp rate) are one singer when one is at least 6 dB weaker than the other: a weaker upper member is a carrier harmonic that the window analysis did not fold in, a weaker lower member is the louder singer’s energy leaking into the band at a fraction of its carrier. Pairs of comparable level are left as two singers. Where a fundamental was itself not resolved, as happens in a dense chorus, its harmonics escape this flag. Machinery modulates broadband noise at one rate that never moves with temperature; per recorder and month, syllable rates are binned at 0.25 Hz and a bin holding at least 40 detections and 3 % of the month’s, fifteen times its neighbors, from at least five recorder-periods, is a line, and detections within 0.5 Hz of it are flagged. Over the year this found ten lines, all at the House station (an induction motor at 59.2 Hz from November to January, with weaker readings of the same rhythm at 18.9 and 37.9 Hz) and none elsewhere; the motor is a useful negative control, a vertical column in the carrier-rate plane, the one shape no insect line takes. In all, 85 % of night detections and 90 % of day detections passed to the linker; harmonics reported as their own singer were 14.6 % of night detections.

#### Linking a period into tracks

The detections of one recorder-period are joined into tracks by a single optimal assignment rather than by chaining window to window. Because all singers cool together, the rate drift common to the period is removed first: for each pair of consecutive windows every detection is matched to its nearest neighbor in the next window within 150 Hz and 3 % of rate, the median step in log rate is taken, and the steps are integrated and smoothed with a three-window median. A detection in one window may continue a detection up to six windows earlier (30 min) when their detrended log rates agree within 2 %, widened by 0.4 % for each skipped window, and their lowest-frequency bands overlap within 150 Hz. The cost of a link is 1 + min((Δy/0.01)², 6) + (ΔF/σ)² + 0.6 per skipped window, where Δy is the detrended rate step, ΔF the step in carrier centroid and σ is 3 % of the carrier with a floor of 60 Hz; ending one track and starting another costs 10. The minimum-cost set of tracks is the minimum-weight bipartite matching between each detection’s exit and each detection’s entry, solved exactly per connected component, so a link is taken only when it is cheaper than a break. Links are not allowed through harmonics: a 7 kHz singer whose attached 12 kHz harmonic overlaps a lone 12 kHz detection does not continue into it. Detections carrying only a chirp rate link to one another on chirp rate within 6 %. Tracks of fewer than three windows are dropped.

Because the rows that make up a band change from window to window as faint edge rows cross or miss the threshold, a carrier measured from the band itself jumps by tens of hertz for reasons unrelated to the singer. For each track one reference band is fixed, the lowest group of touching component bands present in at least a fifth of its windows, and in every window the carrier is the power-weighted centroid over that band plus 150 Hz on either side, each row weighted by the energy that pulses at the singer’s rate, so that a steady neighbor sharing the rows does not move the estimate. Taking the most frequent band instead of the lowest had placed most of the singer-periods above 10 kHz at a harmonic of their fundamental, because in a dense chorus a harmonic survives in more windows than a masked fundamental.

#### Singer-periods and singer-hours

A track of at least 24 windows (two hours) with a fixed-window carrier is a singer-period: a singer-night or a singer-day. Its windows are summarized to one point per clock hour, the singer-hour, with the median rate, median carrier, median depth, median chirp rate and the share of windows that carried one, and the HRRR temperature of the hour. The year holds 26,116 singer-nights (139,171 singer-hours) and 5,427 singer-days (27,673 singer-hours), at temperatures from −6 to 36 °C (median 22). Each singer-period is further summarized by its median rate and carrier and mean temperature.

#### Song form

Whether a singer-period trills or chirps is read from its own audio, in two windows of its track, the deepest and the one of median depth, so that a chirper that trilled in its loudest window is still caught. In each window the band’s envelope in decibels, smoothed over two syllable periods and with its slow trend removed, is tested for a bout rhythm at 1.5 to 12 Hz: the spectral peak in that range is compared with the median of its octave neighborhood (the contrast), and the strength of the band’s modulation at the track’s syllable rate is correlated with the bout envelope over the loudest 30 s. A chirper’s syllables exist only inside its chirps, so the two rise and fall together; a triller’s syllable modulation is steady while a neighbor’s bursts move the envelope. A singer-period is a chirper when every measured window has a contrast above 50 and a correlation above 0.2, a triller when every window has a contrast below 20, and mixed otherwise. On 17 reference recordings of known form the rule classified every one correctly.

#### Call types: the lines in the carrier-rate plane

In the plane of log carrier against log syllable rate, the singer-nights of one population form a diagonal line, not a cloud, because the night’s temperatures spread them along the line. The clustering therefore works in a frame rotated onto the line. Each singer-night is first brought to 25 °C from its mean temperature with the pooled within-singer slopes, 1.54 % per °C for rate and 0.46 % per °C for carrier, and then placed in a frame whose along-line axis has slope 0.55 in log carrier per log rate below 10 kHz and 0.25 above, the slopes the ridges of the plane themselves show. Singer-nights are clustered by average linkage with a tolerance of 0.8 along the line (log-rate units) and 0.06 across it (log-carrier units), so that a population spread along its line by individual variation and residual temperature effects falls in one cluster while a neighbor a few percent away in carrier does not. Neither the season nor the song’s form enters the clustering.

Two refinements follow. A cluster may hold two populations on one line that differ in where they sit along it at a given temperature: the along-line position of its singer-nights is regressed robustly on temperature (Theil-Sen), moved to 25 °C, and fitted with Gaussian mixtures of one and two components; a split is accepted when the Bayesian information criterion improves by at least 50, each component holds at least 15 singer-nights and 8 % of the cluster, and the components are at least 2.5 pooled standard deviations apart. Conversely, neighboring clusters are merged when they lie within 5 % of each other across the line with along-line ranges overlapping or within 5 %, and a single band explains their pooled positions at 25 °C (no improvement of 50 in the criterion, or components closer than 2 standard deviations, or an across-line offset below 2.5 times the pooled scatter, that scatter capped at 3 % so that a wide group cannot absorb its neighbors). A cluster whose position sits at two or three times another’s carrier at the same rate is flagged as a carrier harmonic and excluded. Every remaining cluster of at least 40 singer-nights is a call type; where a cluster holds both chirping and trilling singers, by their audio, it is divided into a chirp type and a trill type at the same position, and the chirp types are then treated on their own plane (below). Clusters were then inspected type by type on the plane, against temperature and as the paths of individual singers through their nights, and six were split by eye where they held two or more populations (two parallel bands offset in carrier, two rate lines at one temperature, or a group of singers whose rate at a given temperature differed from the rest), with the requirement that individual singers not cross between the proposed parts within a night; the clustering itself was not rerun, since the partition proved sensitive to small changes in the carrier estimate. The year gives 37 trill types, numbered 1 to 37 from the lowest carrier up, holding 69 % of all singer-nights; each is heard at a median of 21 recorders on 67 nights, and the types sit between 2.2 and 9.5 kHz and 20 and 81 syllables per second at 25 °C. For Figure 7C each type’s line is the least-squares fit of log carrier on log rate over its singer-hours; the slopes run from 0.33 to 0.69 with a median of 0.54.

Day singer-periods, defined by the same track rule, are assigned to the night-defined lines afterwards: brought to 25 °C, placed in the same frame, and joined to the nearest type when within 6 % across the line and inside the type’s along-line extent with 5 % of slack; three fifths of them are assigned. Day streaks that are large by day but fall below the 40-singer-night cut at night keep no night number and are labeled D1, D2, D3 by carrier.

#### The chirp plane

Chirping singers are typed on a second plane, carrier against chirp rate, because every chirping track has a chirp rate while the syllable rate inside the chirps is read in only part of its hours. A chirping track is one of at least two hours whose kept windows carry a chirp rate in at least half of their number. Because any sound that pulses in bouts below 8 Hz in a band is a chirper to the detector, only tracks whose syllable rate was read in at least half of their hours are typed: 3,943 tracks and 15,547 singer-hours. The 3,894 tracks with a bout rhythm but no readable syllables (17,361 hours; half of them below 4 kHz, the frogs of late winter among them, and the rasping and broadband songs of some katydids) are kept as a separate, untyped class. Singer-hours are brought to 25 °C with the pooled within-track slopes measured on the chirpers themselves, 1.94 % per °C for chirp rate and 0.91 % per °C for carrier, placed in a frame with slope 0.35 in log carrier per log chirp rate, and clustered and merged as the trill types were, with a tolerance of 1.0 along the line and 0.10 across it. Seven clusters reached 40 singer-nights; one, at 15 kHz with 51 of 63 singer-nights at a single station and tracks of two to six hours at low depth, was found on inspection of its grid to hold several different sounds and was excluded. Two of the remaining six held singers at two distinct syllable rates at the same temperature and were split on it, giving eight chirp types, C1 to C8 by carrier, which hold 95 % of the typed chirping singer-nights. The syllable rate is carried as an attribute of each type and gives the number of syllables per chirp as the chirp’s duty cycle divided by the chirp rate times the syllable rate.

#### Carrier-rate coupling within singers

For every track of at least 12 windows whose rate spanned at least 2 percent of its median, the slope of log carrier on log rate across its windows was fitted by least squares, the percent change in carrier per percent change in rate within one singer’s night. Per type the median and interquartile range over its tracks are reported (Figure 8B; 17,799 tracks of the 37 types). To ask whether the line a population traces over the year is the same as the path a singer traces in a night, pairs of singer-hours of one type recorded in the same hour on the same recorder, hence at one air temperature, were taken, and the slope of their carrier offsets on their rate offsets (each pair’s mean removed) was fitted per type; and the syllable duration of each type’s singers was regressed on their carrier after both were corrected for temperature.

#### Syllable shape and the type reference

The shape of each type’s syllable was measured from fine folds. For a singer-period, the loudest 30 s of its deepest window were analyzed at 5.8 ms frames with a 0.73 ms hop (256-point Hann, zero-padded to 2048 points), folded at the refined rate into 64 phase bins (160 for a chirper folded at its chirp rate), expressed in decibels above each row’s quiet level and aligned at the onset of the pulse. From the fold come the duty (share of the period above half the envelope’s peak), the syllable duration (duty over rate), the glide (the change of the power-weighted frequency centroid from the onset to the end of the pulse), its curvature (the quadratic term of centroid against phase within the pulse), and the bandwidth at half power at the envelope’s peak. For the type reference (Figure 17) up to 40 of each type’s clearest singer-nights with night temperature between 21 and 29 °C were taken (the window widened symmetrically when a type had fewer than 15), the fold of each resampled onto a grid of frequency relative to the carrier (±8 %) and normalized, and the pixel-wise median over folds with a contrast of at least 4 dB drawn as the type’s reference. Types were then classified by rule from the medians of these descriptors: form (the plane the type was clustered on); tick, a trill at 25 syllables per second or slower; glide class, flat when the glide is under 1 % of the carrier, otherwise falling or rising; hooked, a falling glide whose centroid sits more than 0.4 % of the carrier below the chord at mid-pulse, the carrier falling steeply at first and then leveling; and for chirpers the chirp length and syllables per chirp. Separately, for the pulse measurements of individual singers, pulses were taken as peaks of the band-limited power at least 0.6 periods apart and 3 dB above a running-median floor, at least 15 per singer-period, and a singer-period with a pulse signal-to-noise ratio of at least 6 dB is called clear; clear singer-periods are preferred wherever examples are drawn.

#### Reference recordings

The species pages of Singing Insects of North America (SINA) were read for their reference recordings (WAV, 44.1 kHz, median 20 s, each captioned with its recording temperature), for the per-species pulse rate at 25 °C and dominant frequency stated in the text, and for Walker’s per-recording tables where they exist. Each recording was run through the window analysis above, and the component nearest the page’s stated dominant frequency was taken as the species’ point: 157 recordings of 134 species have such a point, 133 of them (111 species) of species recorded in Arkansas or a neighboring state. Each was compared with every trill type whose rate range covers it by two distances. Across the line, z is the recording’s perpendicular distance from the type’s fitted line in units of the type’s own scatter in log carrier (floor 0.03). Along the line, each type’s log rate was regressed on HRRR temperature over its singer-hours, and z_along is the recording’s departure from the rate the type sings at the recording’s temperature, in units of the type’s rate scatter at fixed temperature (floor 0.08 in log rate), widened by the rate change over a 2 °C thermometer disagreement. A recording’s best line is the one with the smallest combined distance; a recording within 2.5 standard deviations across its best line is on it, and within 2.5 along it as well is at the right place for its temperature (90 and 68 of the 157; 70 and 53 of the 133 in range). The chirp types were treated the same way on the chirp plane after each recording’s chirp rate was read with the bout test of the song-form section applied to the whole clip (contrast above 50 between 1.5 and 12 Hz): 45 recordings chirp, 21 lie within 2.5 standard deviations of one of the eight chirp lines and 19 also along it. The species’ published positions were placed on the plane as a second, independent marker. No type is named from these comparisons.

For Figure 14 each type is shown with the recording whose best line it is and whose combined distance is smallest, beside four of the type’s field singers from distinct recorder-nights whose night temperature was within 1.5 °C of the recording’s and whose rate within 15 % (the windows widened to 3 °C, then 30 % in rate, then 6 °C, until twelve candidates were available), the clearest and deepest first; of the twelve analyzed the four shown are those with no other singer in the display range, then those with the strongest band above its surroundings, ordered by temperature. For chirp types the match is on the corrected chirp rate of the singer-night and the cells are 1 s long; otherwise cells are the loudest 0.5 s of the singer’s deepest window. Song form was not required to match, so that a reader can see when a recording and a type share position but not form.

#### Looking at the types

For every type a grid of raw excerpts was drawn: twelve positions along the line from the cold, slow end to the warm, fast end, and at each position five singers from distinct recorder-nights ordered by temperature, each cell the loudest 0.5 s (1 s for chirp types) of the singer’s deepest window that has no other singer in its band, on one log-frequency axis shared by the grid, the detected band in color as decibels above the row’s quiet level and the rest of the spectrum in grayscale. A 5 s listening clip centered on every drawn stretch accompanies each grid. These pages, one per type, were the quality control for the type list: the chirp types of a first pass, clustered without the syllable requirement, were judged from them to mix crickets with frogs and birds, which led to the rule above, and the excluded 15 kHz cluster was judged from its page. Figure 10 is a 6 by 3 sample of three of these grids.

#### Type mix

The activity of a type in a week is its singer-hours divided by the hours recorded that week by the recorders in question, the mean number of its singers tracked at once per recorded hour; weeks ending Saturday. No count of individuals is made. For the per-site panels of Figure 11 the denominator is the site’s own recorded hours, and weeks with under a quarter of the site’s median weekly recording are hatched.

#### Validation and controls

Twenty-three windows from September 2024 were annotated manually before the detector was revised: of 112 annotated singers 104 were covered by a detected singer, and in 21 of 23 windows every rate was right; the misses were faint singers under louder neighbors and the false positives non-insect sound. The House motor is the negative control for the plane. In the September 2024 month a Wildlife Acoustics SM4 recorder 64 m from a BAR-LT placed its detections on the same lines with the same slopes, and the recorders’ sample clocks (0.45 ppm per °C) are five orders of magnitude too stable to produce the rate slopes. A singer-night may belong to a trill type on the main plane and a chirp type on the chirp plane (a chirper whose syllable rate was read); the two catalogs are kept separate and no singer-night is counted twice within a plane.

#### Software

Python 3.11 with NumPy, SciPy (including the linear-sum-assignment solver used for linking and the Theil-Sen estimator), pandas, scikit-learn (Gaussian mixtures) and Matplotlib. Detection over the corpus took 151 hours on 22 cores; everything from the machine-line scan to the figures runs in a few hours. Code and the per-window, per-track and per-type tables will be deposited with the paper. Initial versions of the software were written in JupyterLab. Subsequent versions utilized Claude Code (Anthropic) for code generation, project management and manuscript proofreading.

## Data and Code Availability

The raw data for this paper are over 160 Tb. We are working on a way to make them available. We are happy to share any of the data with anyone and will figure out how to get it to you.

Code, example data and outputs are available at https://github.com/cardinal-acoustics/brooks

## Funding and Acknowledgements

This work is part of a long-term collaboration with Patrick O. Brown of Stanford University to carry out experiments to optimize strategies for large-scale restoration of native habitats and biodiversity on land currently used for animal agriculture, in which the Arkansas site discussed here features prominently.

This work was funded by a Howard Hughes Medical Institute (HHMI) Investigator Award. This funding includes purchasing the recorders used here, funding all of the computational hardware, software and cloud services used in the analysis, and funding travel to the site. MBE is an employee of HHMI. The site where these experiments were conducted is owned and managed by the Impossible Foundation.

MBE thanks Cody and Katlin Lockhart for their work on maintaining the site, Ana Sanz Matias, Nadya Ali and Holli Weld for logistical assistance, and Natasha Mhatre for helpful discussions.

