## Supplemental Figures for "A million orthopteran singers from a year of continuously operating field recorders detected, tracked and organized by song"

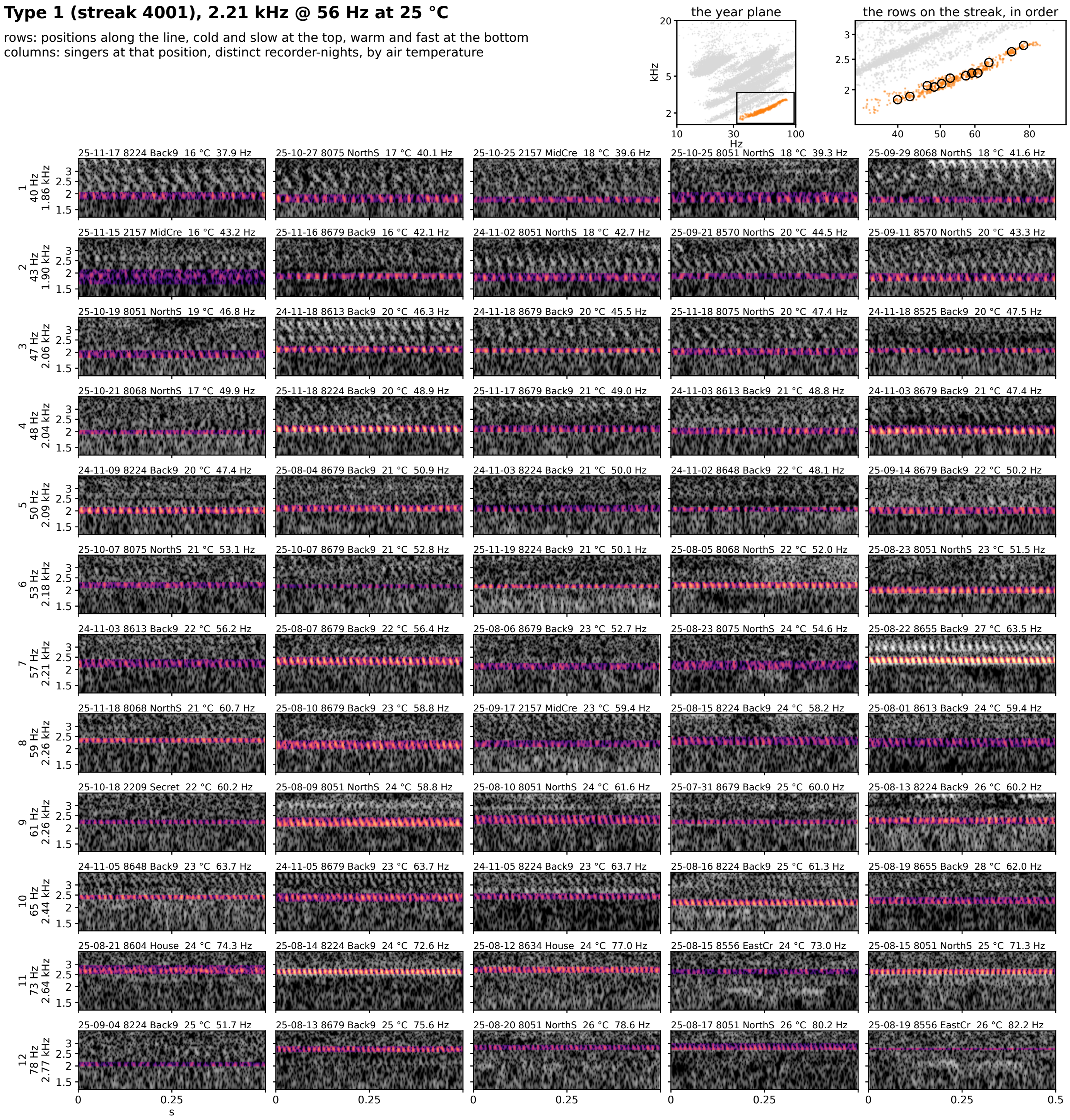


**Figure 7S-1.** Type 1 (streak 4001): 2.21 kHz @ 56 Hz at 25 °C, 132 singer-nights. Rows are positions along the line from the cold, slow end at the top to the warm, fast end at the bottom; columns are singers at that position from distinct recorder-nights, ordered by air temperature. Each cell is 0.5 s of the loudest 30 s of the singer's deepest window on one shared log-frequency axis, the detected band in color (dB above the row's quiet level, 0 to 30) and the rest in grayscale. The cell head gives the night, recorder, site, air temperature and rate (c: chirp rate). Top right: the type on the plane, and the row positions on the streak, in order.

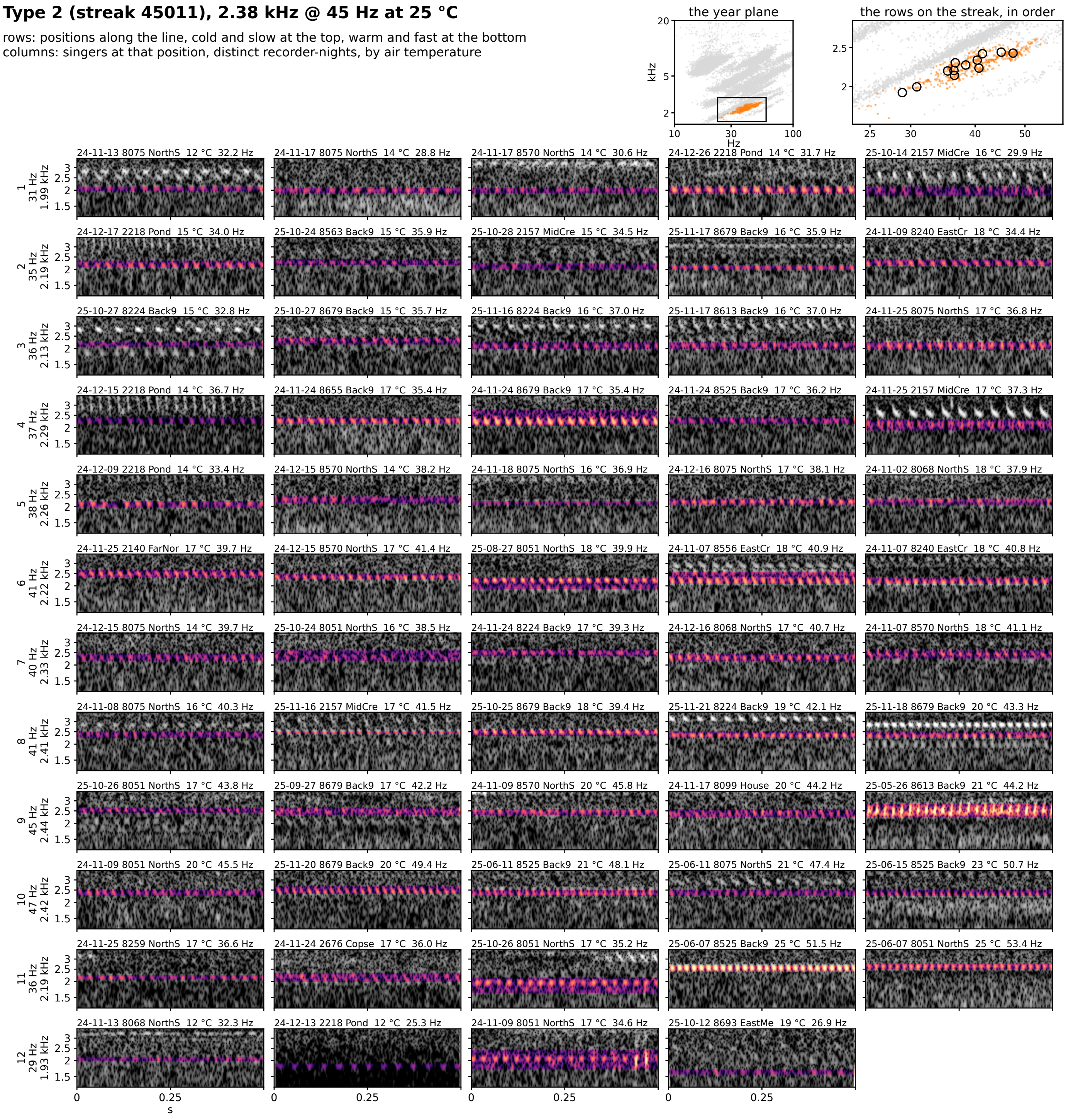


**Figure 7S-2.** Type 2 (streak 45011): 2.38 kHz @ 45 Hz at 25 °C, 109 singer-nights. Rows are positions along the line from the cold, slow end at the top to the warm, fast end at the bottom; columns are singers at that position from distinct recorder-nights, ordered by air temperature. Each cell is 0.5 s of the loudest 30 s of the singer's deepest window on one shared log-frequency axis, the detected band in color (dB above the row's quiet level, 0 to 30) and the rest in grayscale. The cell head gives the night, recorder, site, air temperature and rate (c: chirp rate). Top right: the type on the plane, and the row positions on the streak, in order.

**Type 3 (streak 4301), 2.59 kHz @ 41 Hz at 25 °C**

rows: positions along the line, cold and slow at the top, warm and fast at the bottom
columns: singers at that position, distinct recorder-nights, by air temperature

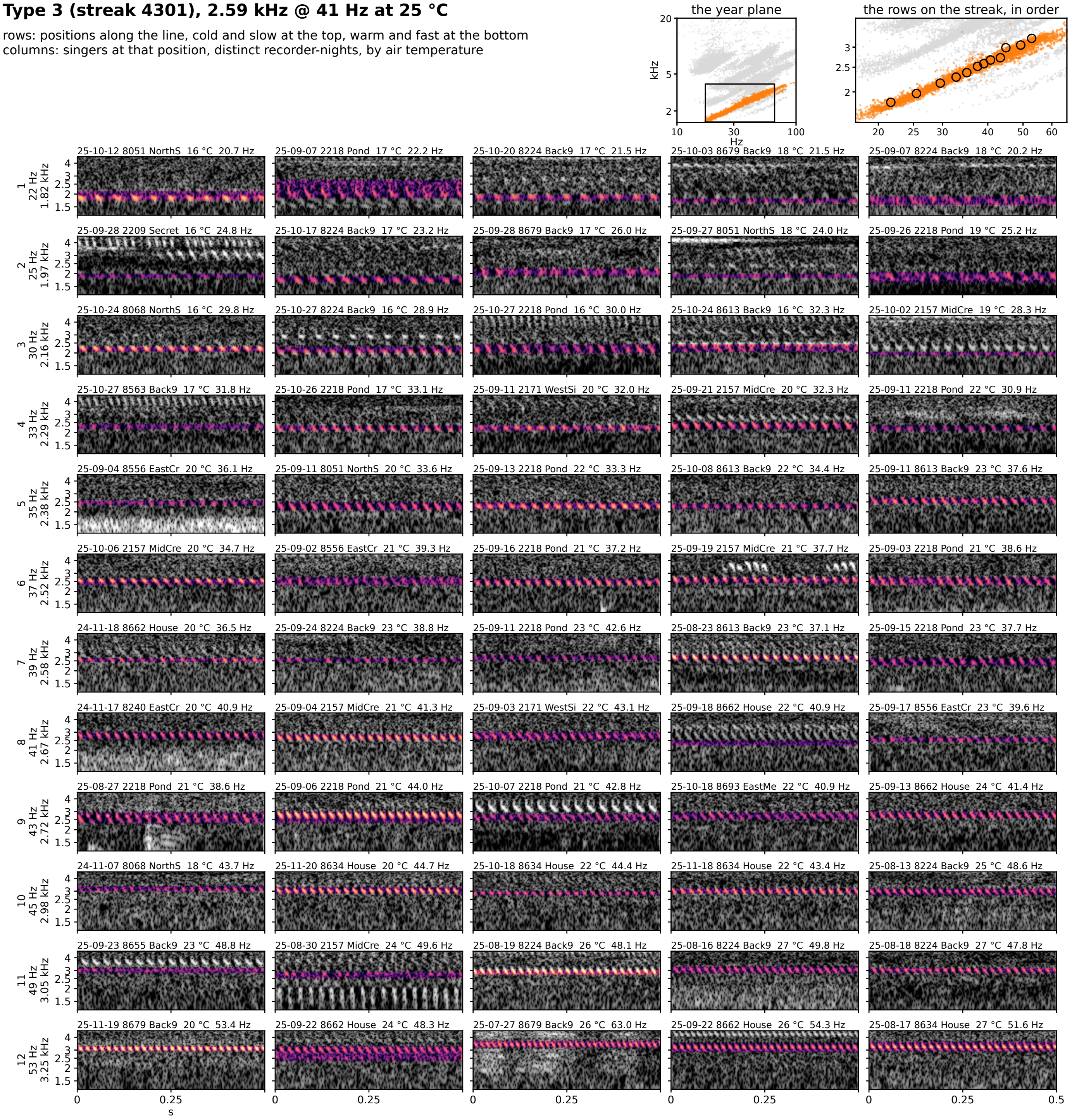


**Figure 7S-3.** Type 3 (streak 4301): 2.59 kHz @ 41 Hz at 25 °C, 1655 singer-nights. Rows are positions along the line from the cold, slow end at the top to the warm, fast end at the bottom; columns are singers at that position from distinct recorder-nights, ordered by air temperature. Each cell is 0.5 s of the loudest 30 s of the singer's deepest window on one shared log-frequency axis, the detected band in color (dB above the row's quiet level, 0 to 30) and the rest in grayscale. The cell head gives the night, recorder, site, air temperature and rate (c: chirp rate). Top right: the type on the plane, and the row positions on the streak, in order.

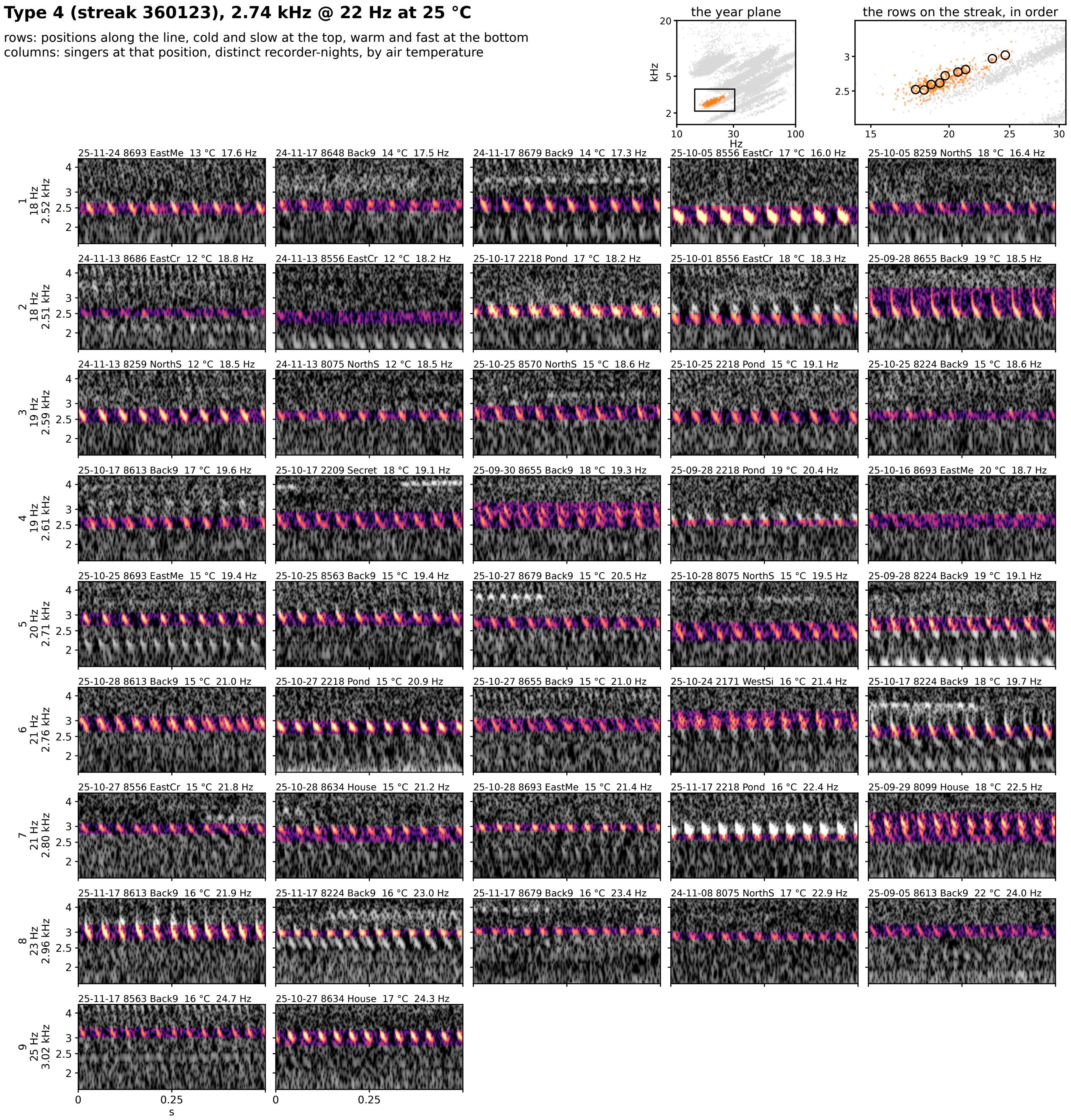


**Figure 7S-4.** Type 4 (streak 360123): 2.74 kHz @ 22 Hz at 25 °C, 76 singer-nights. Rows are positions along the line from the cold, slow end at the top to the warm, fast end at the bottom; columns are singers at that position from distinct recorder-nights, ordered by air temperature. Each cell is 0.5 s of the loudest 30 s of the singer's deepest window on one shared log-frequency axis, the detected band in color (dB above the row's quiet level, 0 to 30) and the rest in grayscale. The cell head gives the night, recorder, site, air temperature and rate (c: chirp rate). Top right: the type on the plane, and the row positions on the streak, in order.

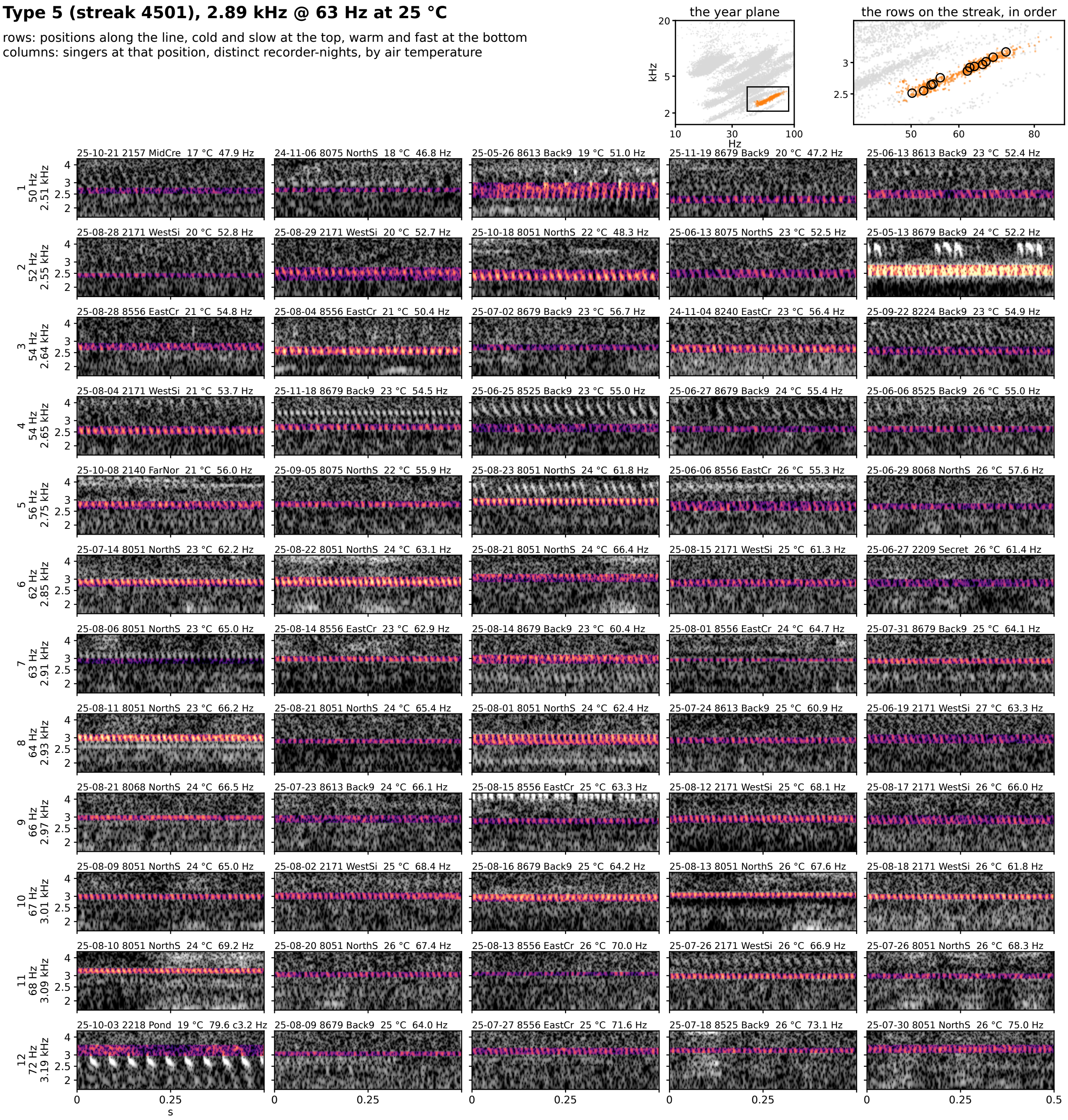


**Figure 7S-5.** Type 5 (streak 4501): 2.89 kHz @ 63 Hz at 25 °C, 144 singer-nights. Rows are positions along the line from the cold, slow end at the top to the warm, fast end at the bottom; columns are singers at that position from distinct recorder-nights, ordered by air temperature. Each cell is 0.5 s of the loudest 30 s of the singer's deepest window on one shared log-frequency axis, the detected band in color (dB above the row's quiet level, 0 to 30) and the rest in grayscale. The cell head gives the night, recorder, site, air temperature and rate (c: chirp rate). Top right: the type on the plane, and the row positions on the streak, in order.

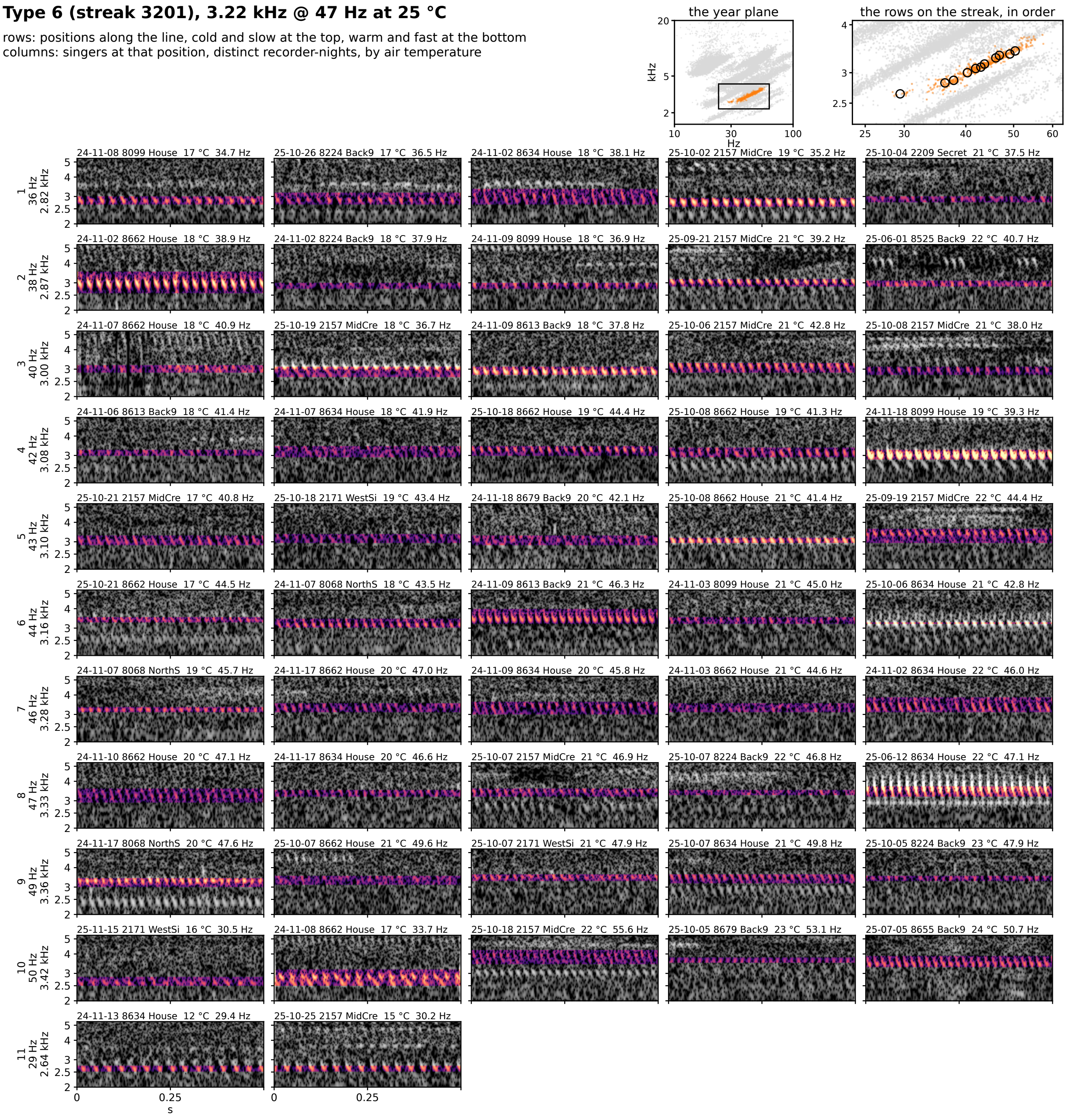


**Figure 7S-6.** Type 6 (streak 3201): 3.22 kHz @ 47 Hz at 25 °C, 106 singer-nights. Rows are positions along the line from the cold, slow end at the top to the warm, fast end at the bottom; columns are singers at that position from distinct recorder-nights, ordered by air temperature. Each cell is 0.5 s of the loudest 30 s of the singer's deepest window on one shared log-frequency axis, the detected band in color (dB above the row's quiet level, 0 to 30) and the rest in grayscale. The cell head gives the night, recorder, site, air temperature and rate (c: chirp rate). Top right: the type on the plane, and the row positions on the streak, in order.

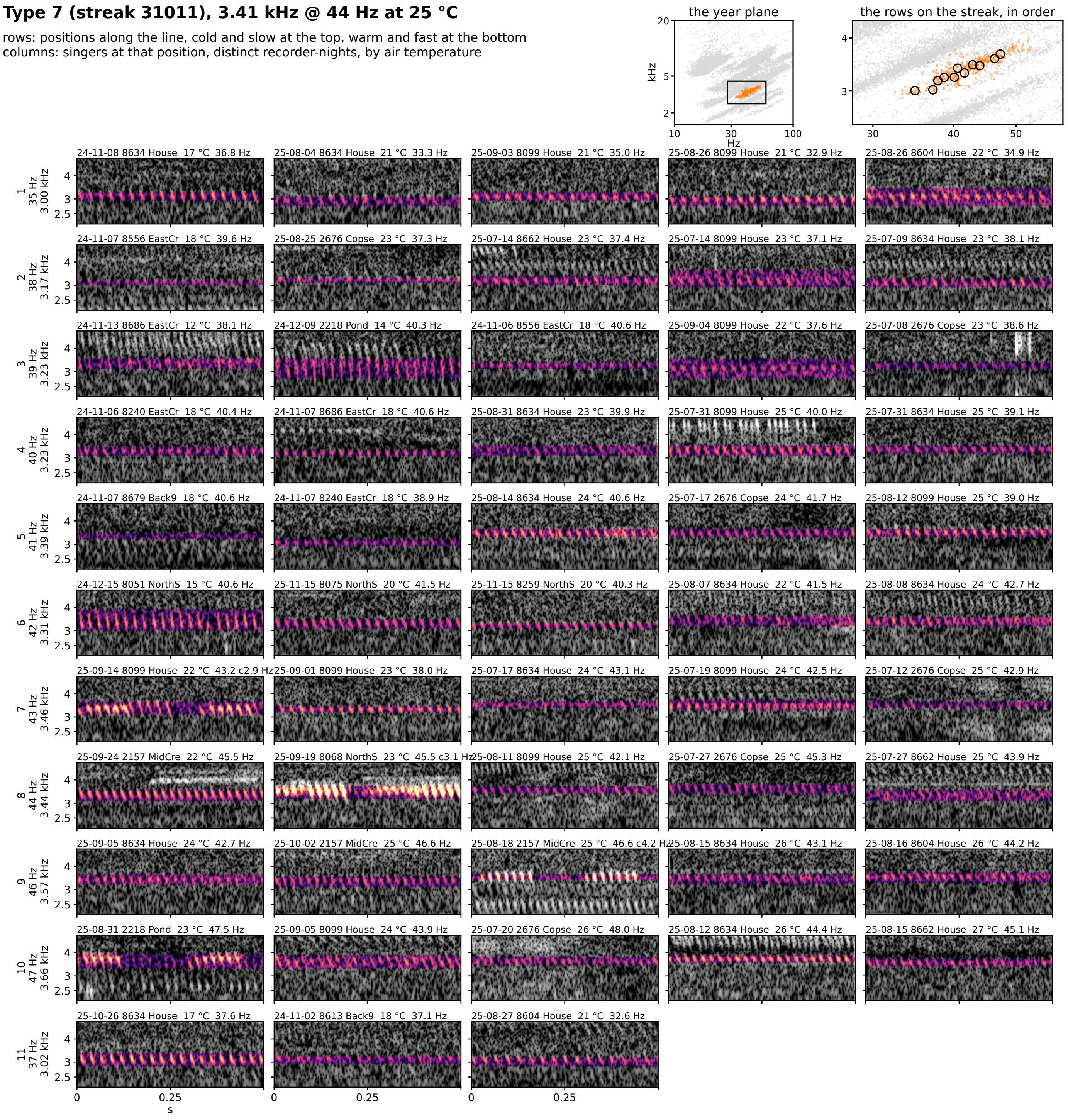


**Figure 7S-7.** Type 7 (streak 31011): 3.41 kHz @ 44 Hz at 25 °C, 90 singer-nights. Rows are positions along the line from the cold, slow end at the top to the warm, fast end at the bottom; columns are singers at that position from distinct recorder-nights, ordered by air temperature. Each cell is 0.5 s of the loudest 30 s of the singer's deepest window on one shared log-frequency axis, the detected band in color (dB above the row's quiet level, 0 to 30) and the rest in grayscale. The cell head gives the night, recorder, site, air temperature and rate (c: chirp rate). Top right: the type on the plane, and the row positions on the streak, in order.

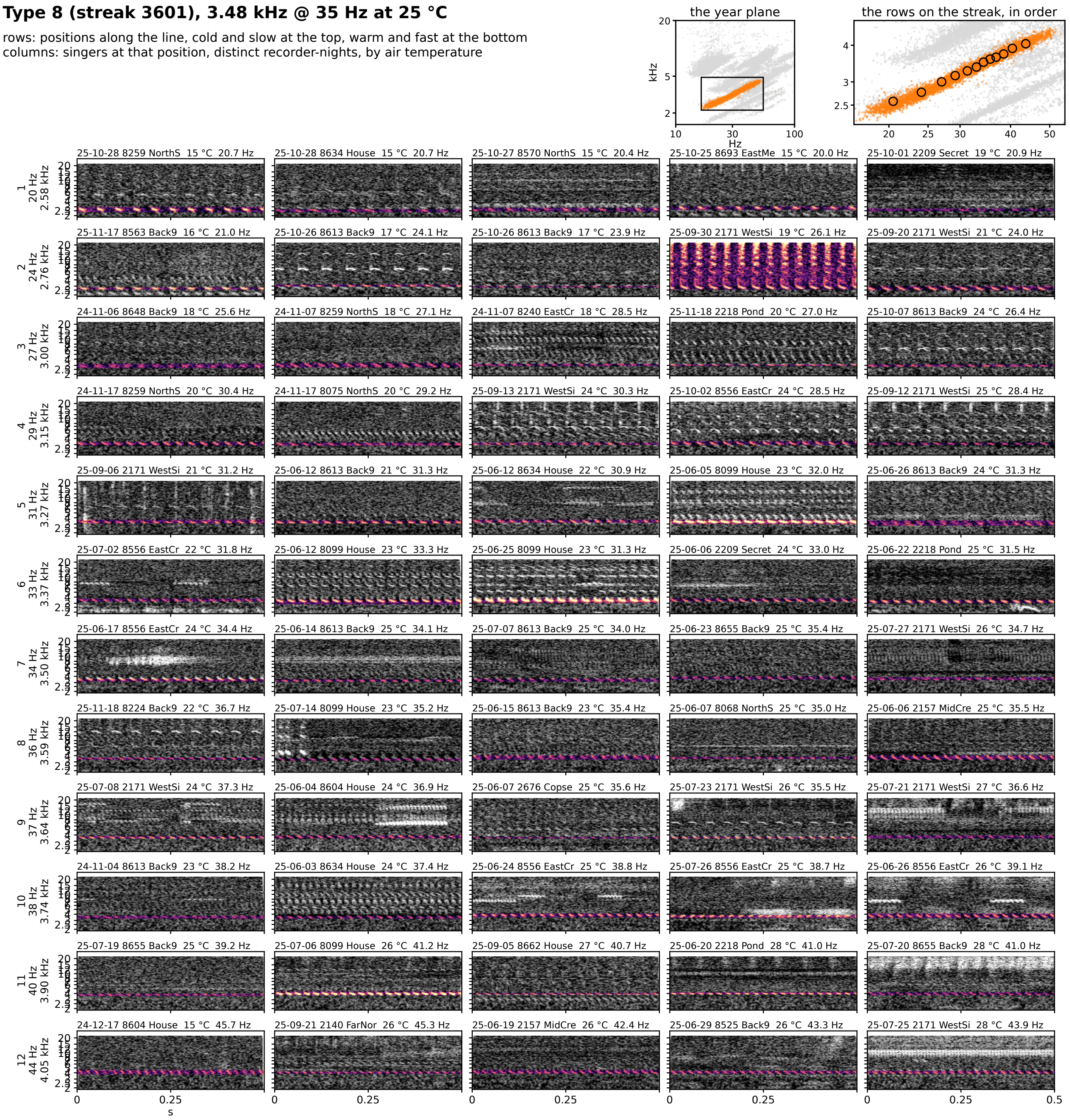


**Figure 7S-8.** Type 8 (streak 3601): 3.48 kHz @ 35 Hz at 25 °C, 1834 singer-nights. Rows are positions along the line from the cold, slow end at the top to the warm, fast end at the bottom; columns are singers at that position from distinct recorder-nights, ordered by air temperature. Each cell is 0.5 s of the loudest 30 s of the singer's deepest window on one shared log-frequency axis, the detected band in color (dB above the row's quiet level, 0 to 30) and the rest in grayscale. The cell head gives the night, recorder, site, air temperature and rate (c: chirp rate). Top right: the type on the plane, and the row positions on the streak, in order.

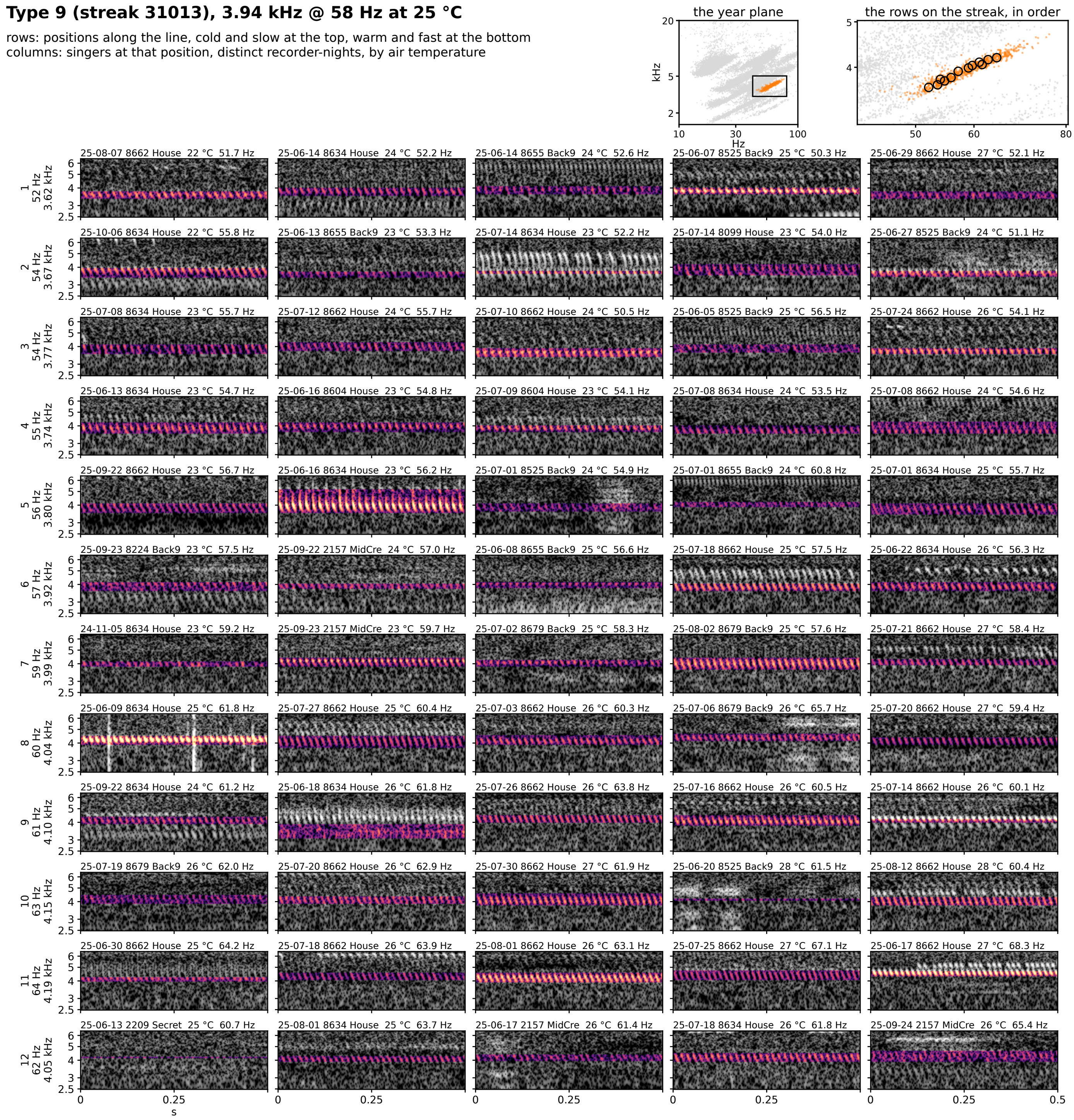


**Figure 7S-9.** Type 9 (streak 31013): 3.94 kHz @ 58 Hz at 25 °C, 167 singer-nights. Rows are positions along the line from the cold, slow end at the top to the warm, fast end at the bottom; columns are singers at that position from distinct recorder-nights, ordered by air temperature. Each cell is 0.5 s of the loudest 30 s of the singer's deepest window on one shared log-frequency axis, the detected band in color (dB above the row's quiet level, 0 to 30) and the rest in grayscale. The cell head gives the night, recorder, site, air temperature and rate (c: chirp rate). Top right: the type on the plane, and the row positions on the streak, in order.

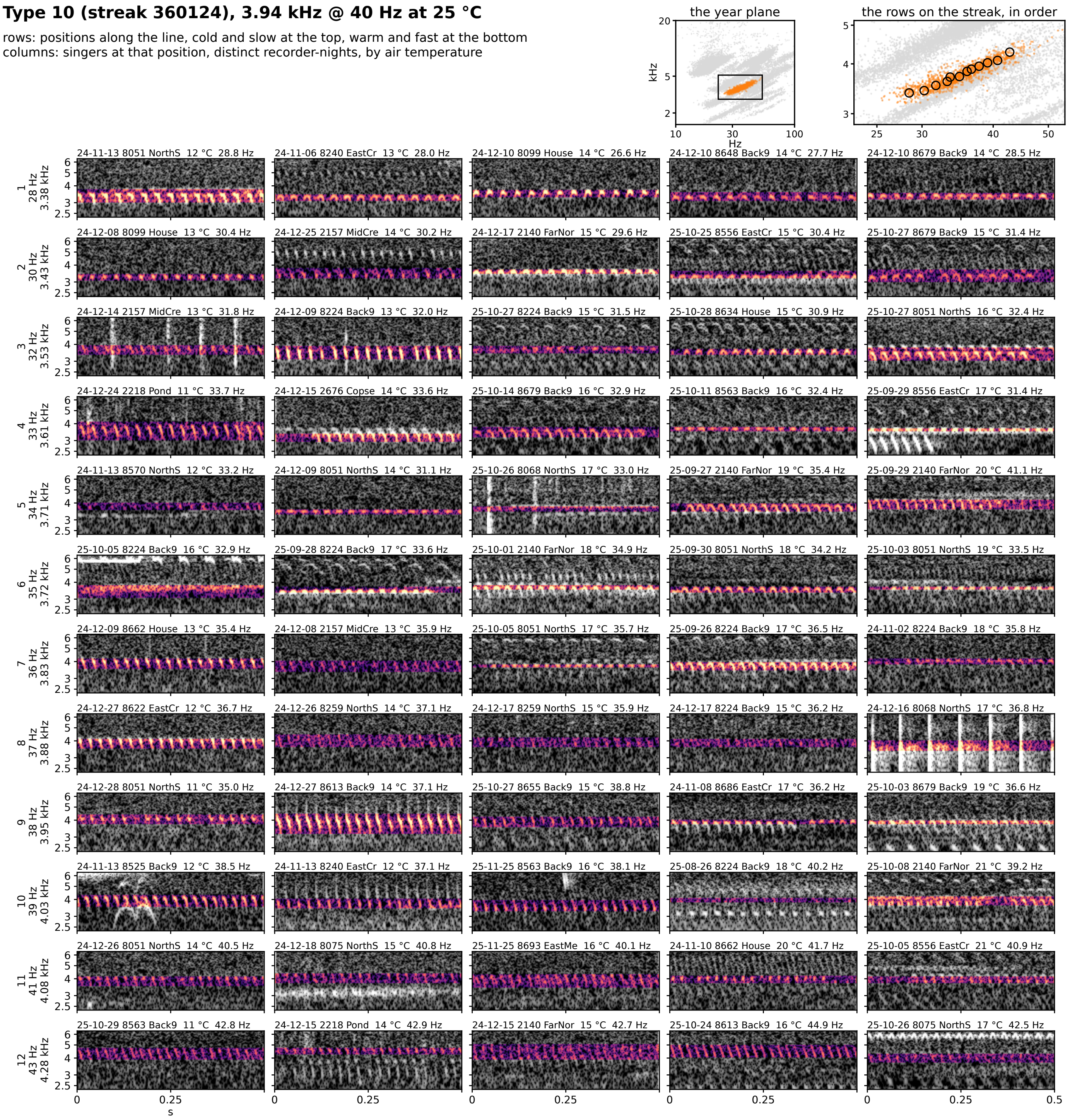


**Figure 7S-10.** Type 10 (streak 360124): 3.94 kHz @ 40 Hz at 25 °C, 231 singer-nights. Rows are positions along the line from the cold, slow end at the top to the warm, fast end at the bottom; columns are singers at that position from distinct recorder-nights, ordered by air temperature. Each cell is 0.5 s of the loudest 30 s of the singer's deepest window on one shared log-frequency axis, the detected band in color (dB above the row's quiet level, 0 to 30) and the rest in grayscale. The cell head gives the night, recorder, site, air temperature and rate (c: chirp rate). Top right: the type on the plane, and the row positions on the streak, in order.

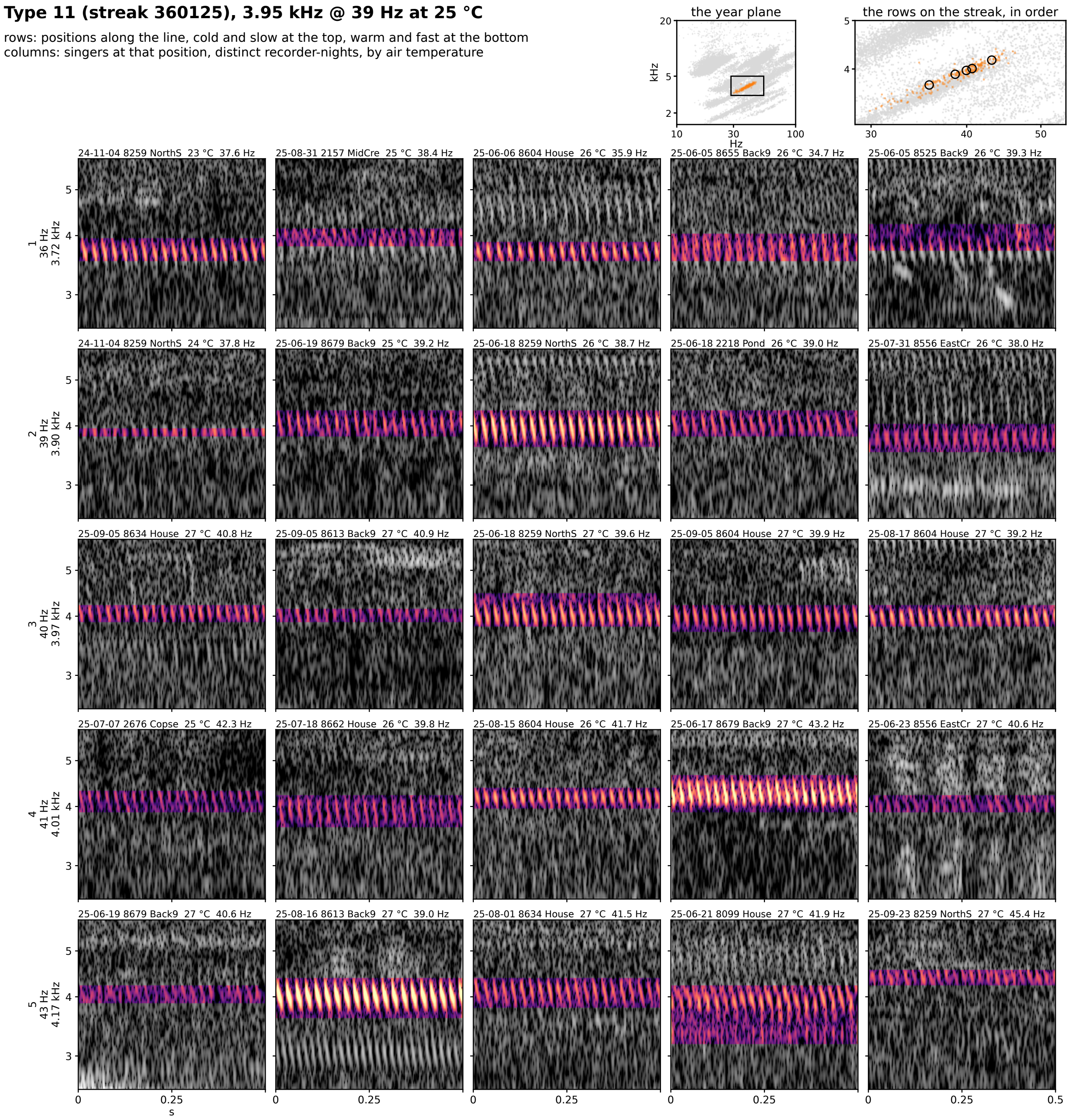


**Figure 7S-11.** Type 11 (streak 360125): 3.95 kHz @ 39 Hz at 25 °C, 42 singer-nights. Rows are positions along the line from the cold, slow end at the top to the warm, fast end at the bottom; columns are singers at that position from distinct recorder-nights, ordered by air temperature. Each cell is 0.5 s of the loudest 30 s of the singer's deepest window on one shared log-frequency axis, the detected band in color (dB above the row's quiet level, 0 to 30) and the rest in grayscale. The cell head gives the night, recorder, site, air temperature and rate (c: chirp rate). Top right: the type on the plane, and the row positions on the streak, in order.

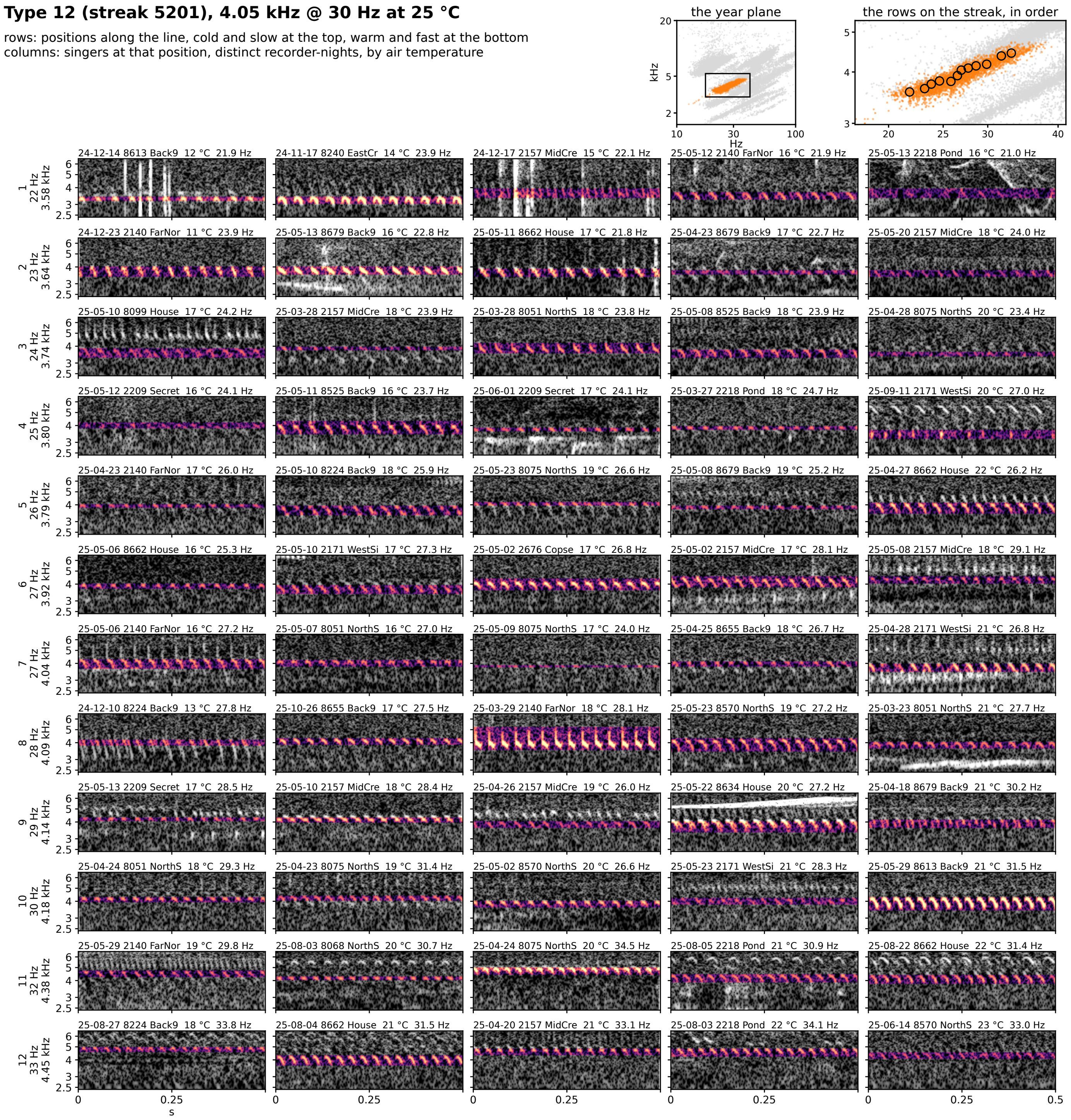


**Figure 7S-12.** Type 12 (streak 5201): 4.05 kHz @ 30 Hz at 25 °C, 630 singer-nights. Rows are positions along the line from the cold, slow end at the top to the warm, fast end at the bottom; columns are singers at that position from distinct recorder-nights, ordered by air temperature. Each cell is 0.5 s of the loudest 30 s of the singer's deepest window on one shared log-frequency axis, the detected band in color (dB above the row's quiet level, 0 to 30) and the rest in grayscale. The cell head gives the night, recorder, site, air temperature and rate (c: chirp rate). Top right: the type on the plane, and the row positions on the streak, in order.

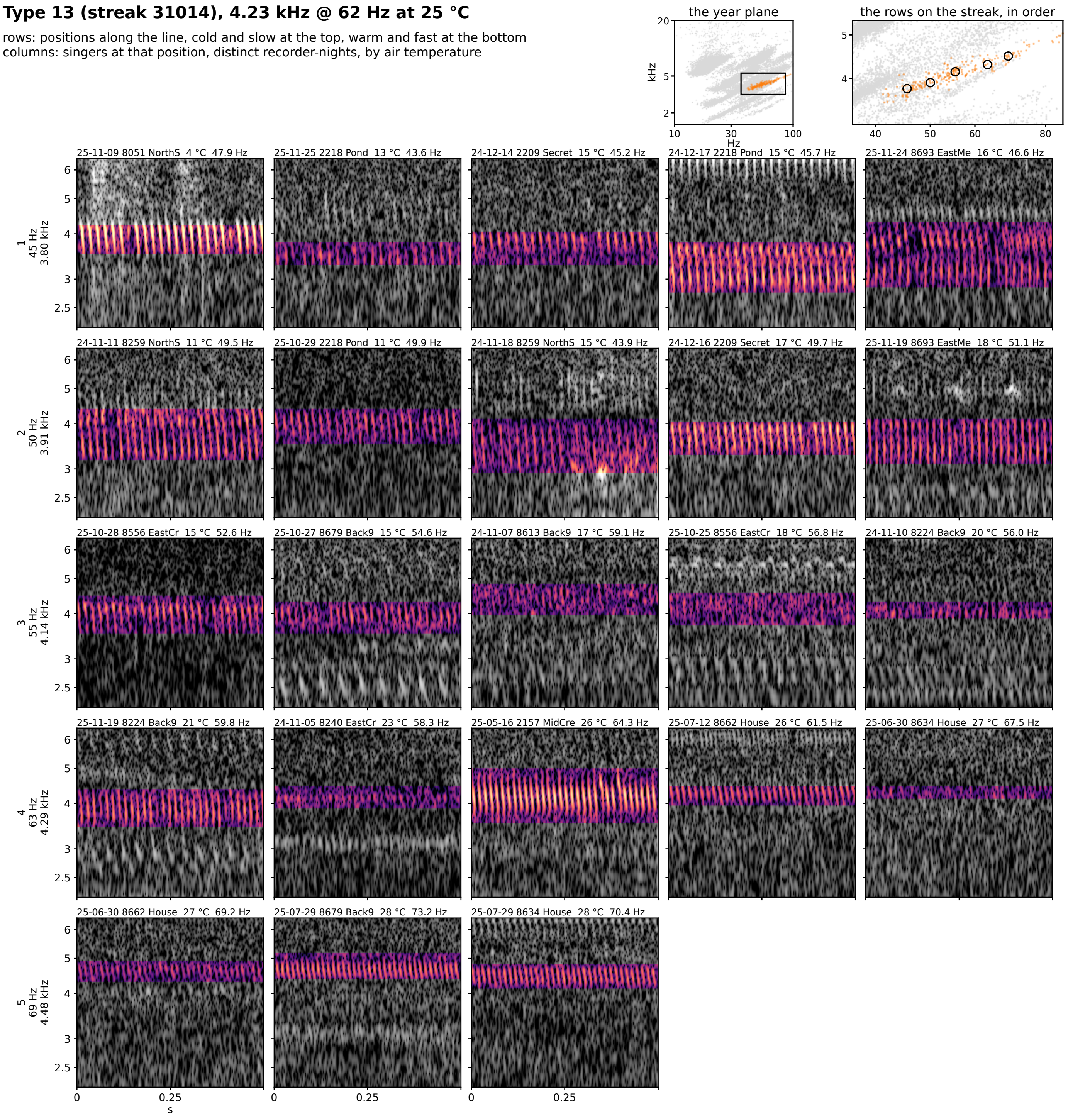


**Figure 7S-13.** Type 13 (streak 31014): 4.23 kHz @ 62 Hz at 25 °C, 44 singer-nights. Rows are positions along the line from the cold, slow end at the top to the warm, fast end at the bottom; columns are singers at that position from distinct recorder-nights, ordered by air temperature. Each cell is 0.5 s of the loudest 30 s of the singer's deepest window on one shared log-frequency axis, the detected band in color (dB above the row's quiet level, 0 to 30) and the rest in grayscale. The cell head gives the night, recorder, site, air temperature and rate (c: chirp rate). Top right: the type on the plane, and the row positions on the streak, in order.

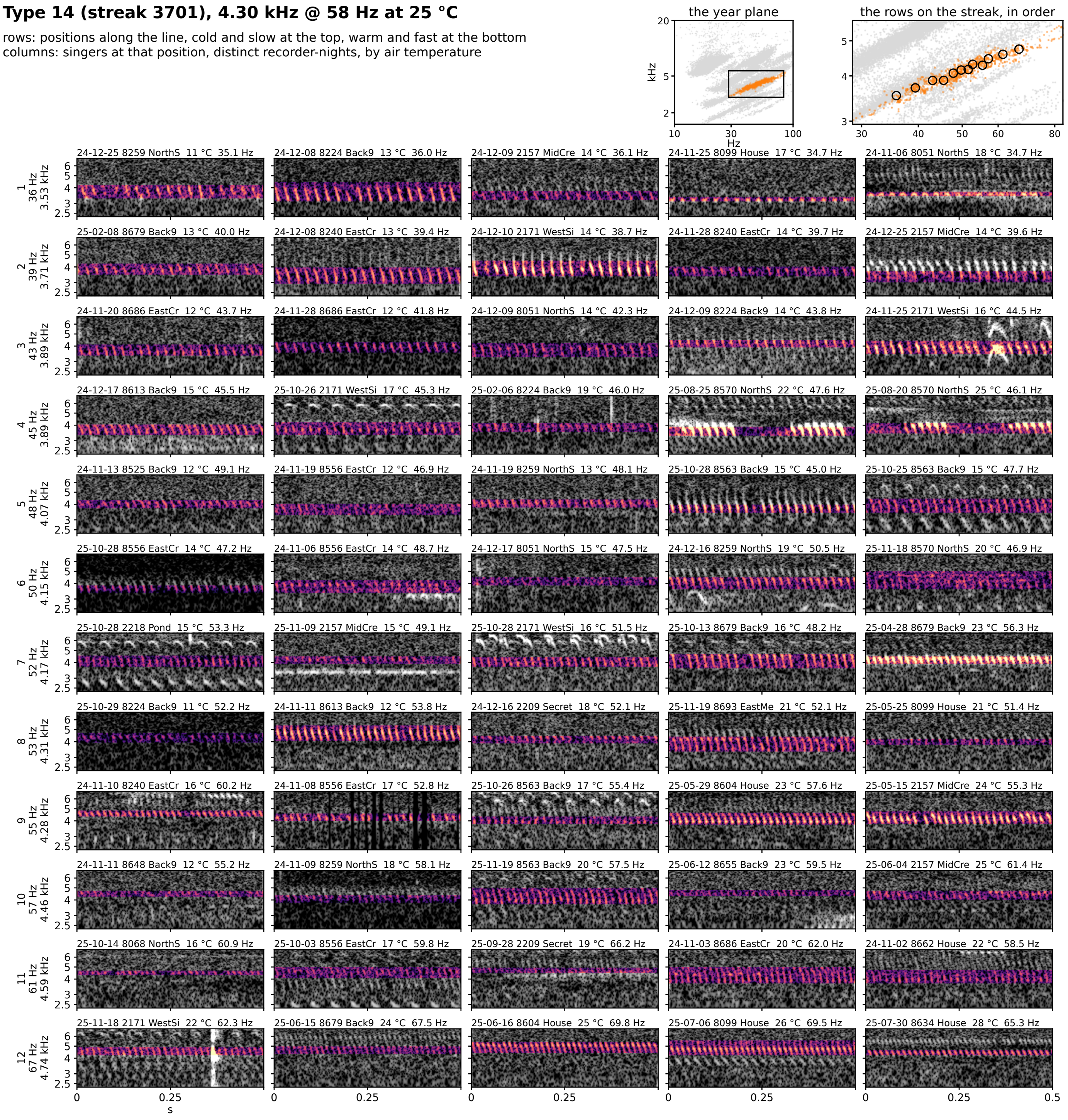


**Figure 7S-14.** Type 14 (streak 3701): 4.30 kHz @ 58 Hz at 25 °C, 166 singer-nights. Rows are positions along the line from the cold, slow end at the top to the warm, fast end at the bottom; columns are singers at that position from distinct recorder-nights, ordered by air temperature. Each cell is 0.5 s of the loudest 30 s of the singer's deepest window on one shared log-frequency axis, the detected band in color (dB above the row's quiet level, 0 to 30) and the rest in grayscale. The cell head gives the night, recorder, site, air temperature and rate (c: chirp rate). Top right: the type on the plane, and the row positions on the streak, in order.

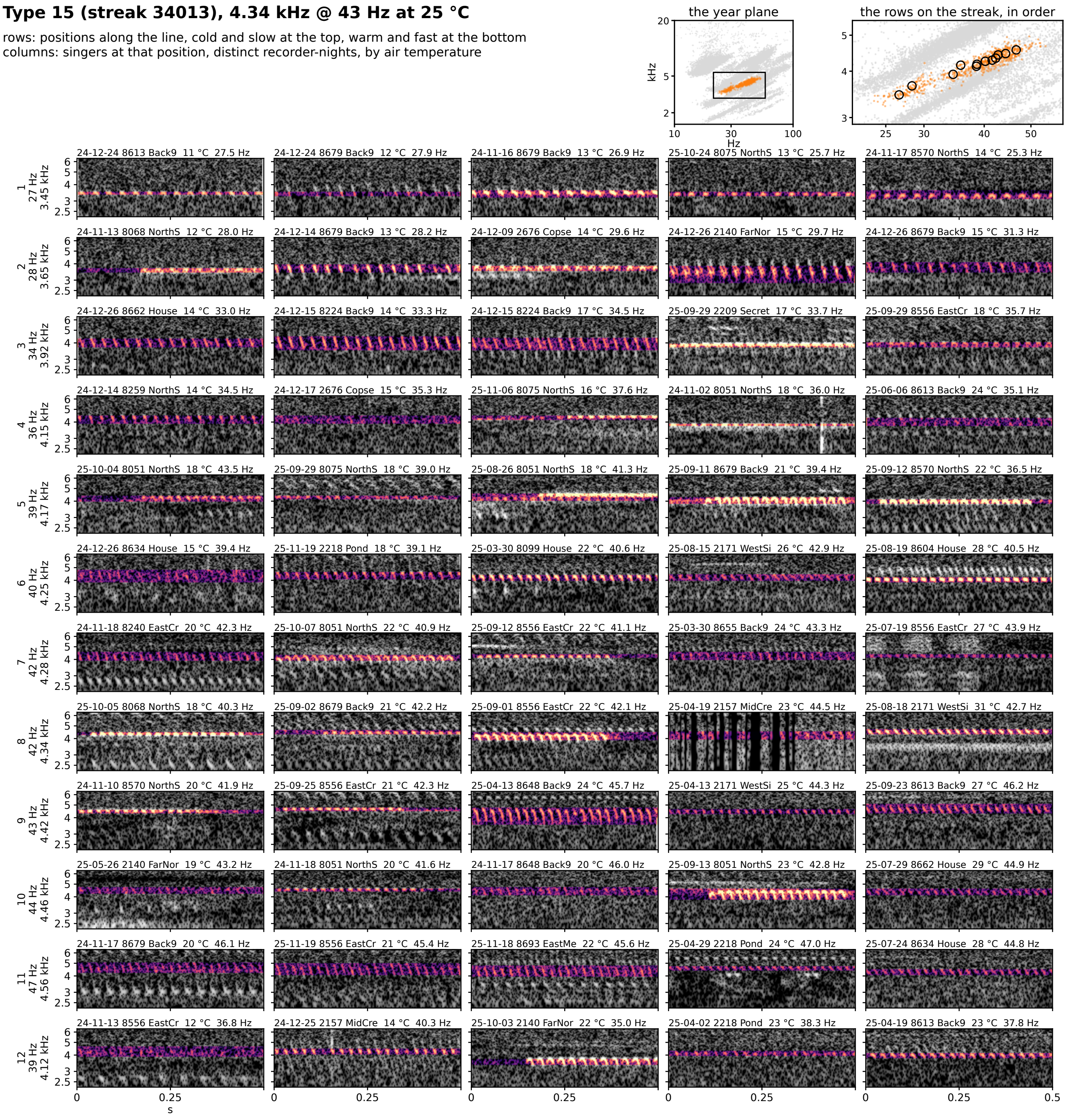


**Figure 7S-15.** Type 15 (streak 34013): 4.34 kHz @ 43 Hz at 25 °C, 124 singer-nights. Rows are positions along the line from the cold, slow end at the top to the warm, fast end at the bottom; columns are singers at that position from distinct recorder-nights, ordered by air temperature. Each cell is 0.5 s of the loudest 30 s of the singer's deepest window on one shared log-frequency axis, the detected band in color (dB above the row's quiet level, 0 to 30) and the rest in grayscale. The cell head gives the night, recorder, site, air temperature and rate (c: chirp rate). Top right: the type on the plane, and the row positions on the streak, in order.

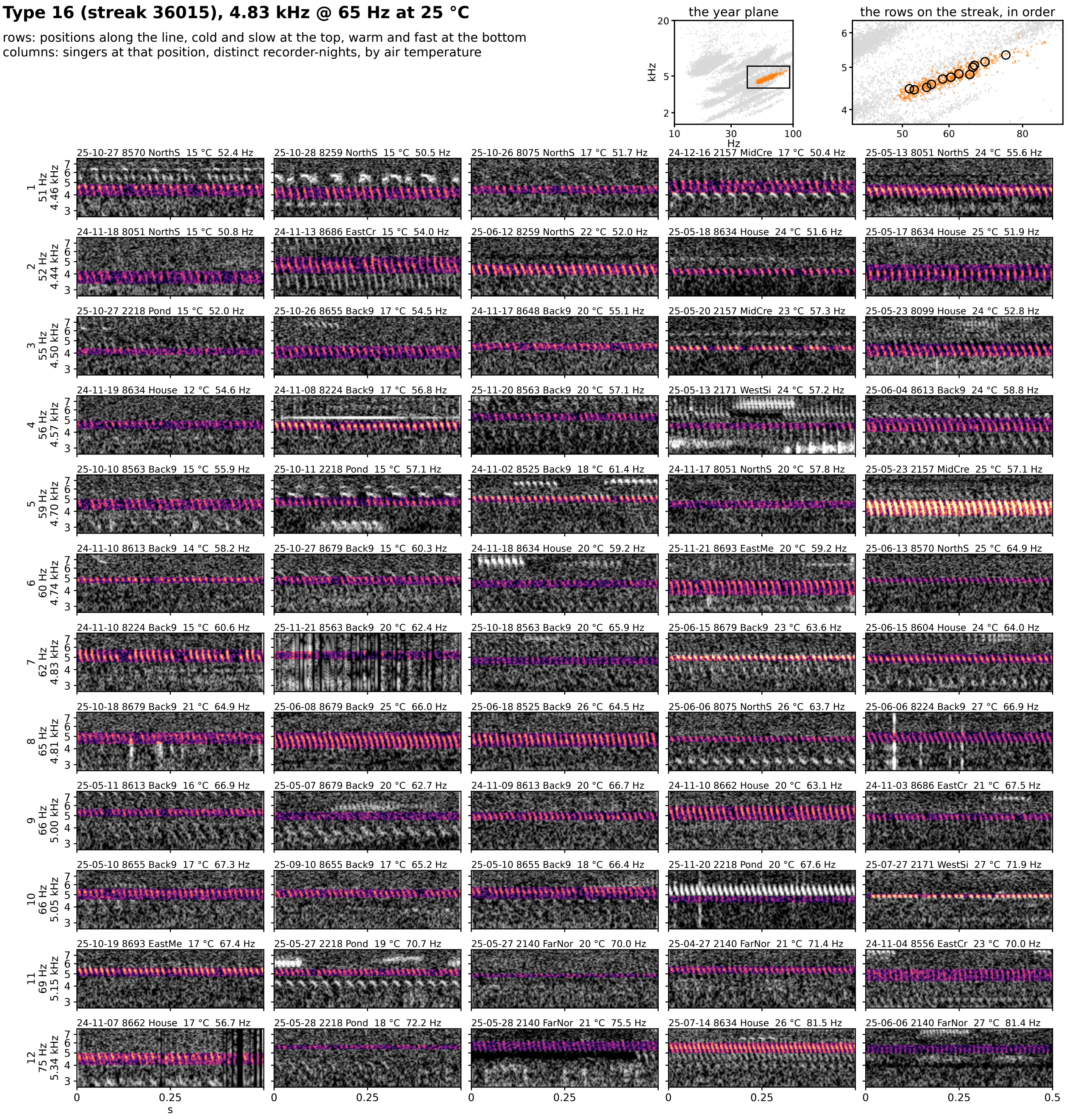


**Figure 7S-16.** Type 16 (streak 36015): 4.83 kHz @ 65 Hz at 25 °C, 114 singer-nights. Rows are positions along the line from the cold, slow end at the top to the warm, fast end at the bottom; columns are singers at that position from distinct recorder-nights, ordered by air temperature. Each cell is 0.5 s of the loudest 30 s of the singer's deepest window on one shared log-frequency axis, the detected band in color (dB above the row's quiet level, 0 to 30) and the rest in grayscale. The cell head gives the night, recorder, site, air temperature and rate (c: chirp rate). Top right: the type on the plane, and the row positions on the streak, in order.

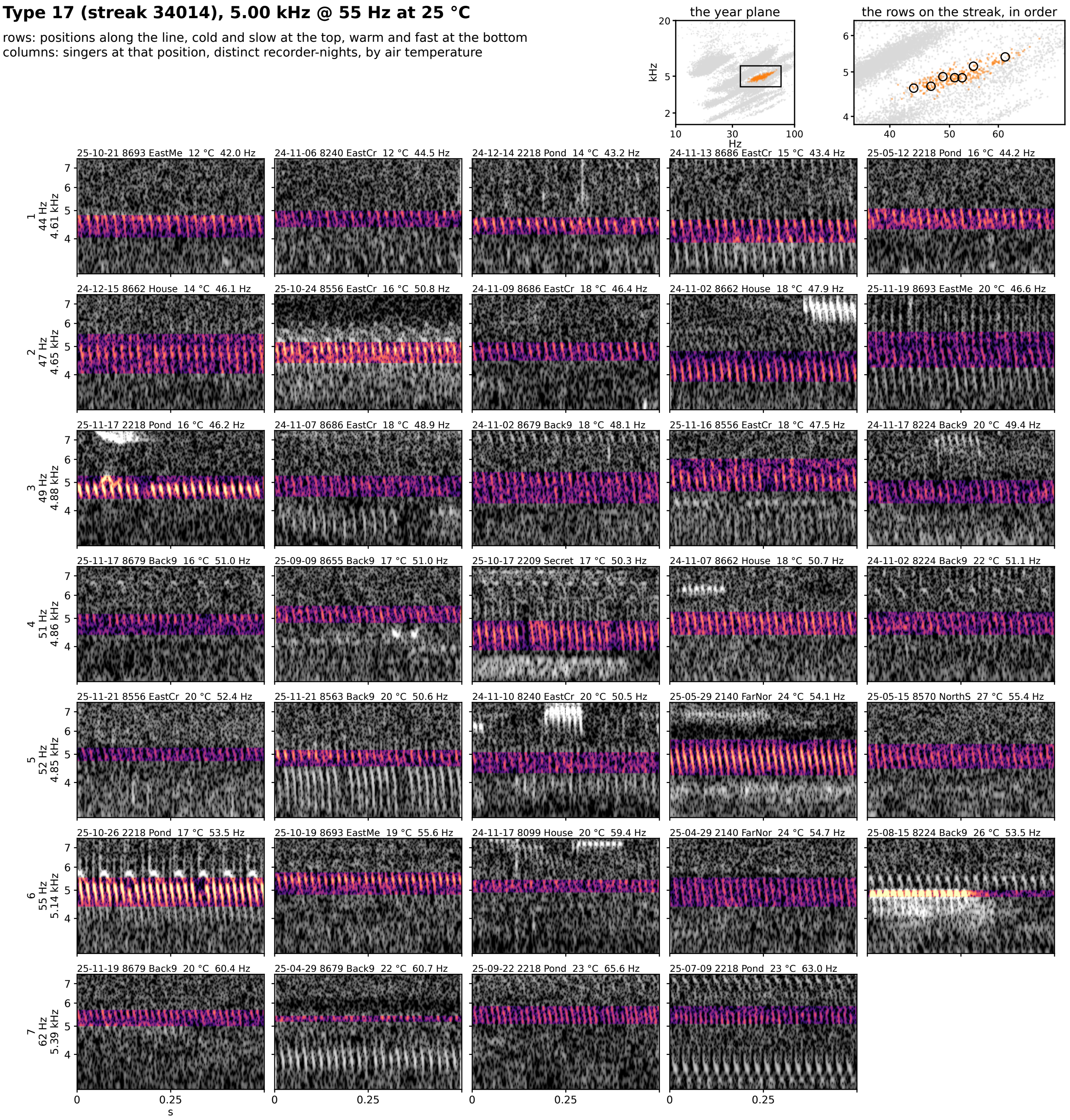


**Figure 7S-17.** Type 17 (streak 34014): 5.00 kHz @ 55 Hz at 25 °C, 62 singer-nights. Rows are positions along the line from the cold, slow end at the top to the warm, fast end at the bottom; columns are singers at that position from distinct recorder-nights, ordered by air temperature. Each cell is 0.5 s of the loudest 30 s of the singer's deepest window on one shared log-frequency axis, the detected band in color (dB above the row's quiet level, 0 to 30) and the rest in grayscale. The cell head gives the night, recorder, site, air temperature and rate (c: chirp rate). Top right: the type on the plane, and the row positions on the streak, in order.

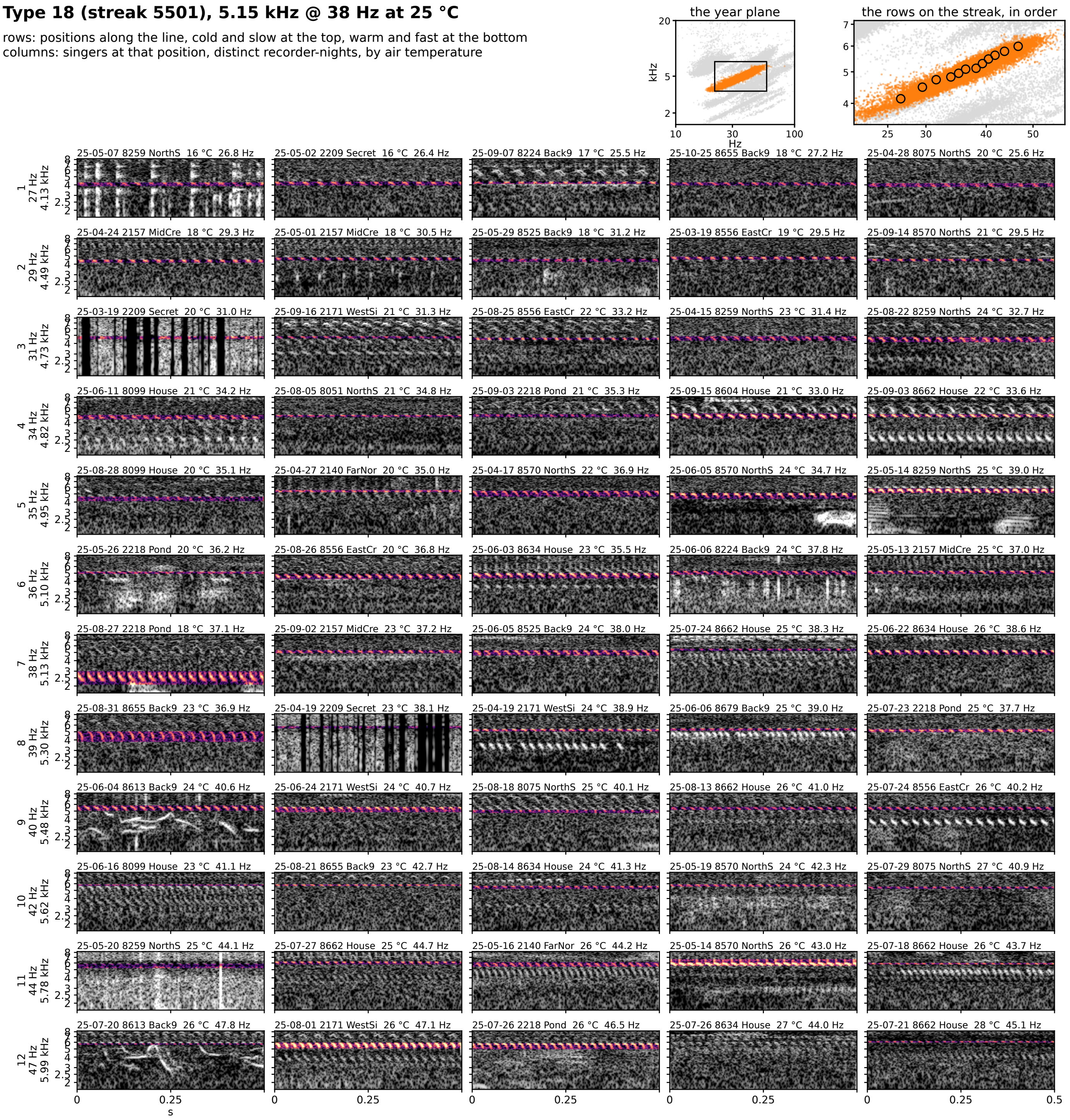


**Figure 7S-18.** Type 18 (streak 5501): 5.15 kHz @ 38 Hz at 25 °C, 4213 singer-nights. Rows are positions along the line from the cold, slow end at the top to the warm, fast end at the bottom; columns are singers at that position from distinct recorder-nights, ordered by air temperature. Each cell is 0.5 s of the loudest 30 s of the singer's deepest window on one shared log-frequency axis, the detected band in color (dB above the row's quiet level, 0 to 30) and the rest in grayscale. The cell head gives the night, recorder, site, air temperature and rate (c: chirp rate). Top right: the type on the plane, and the row positions on the streak, in order.

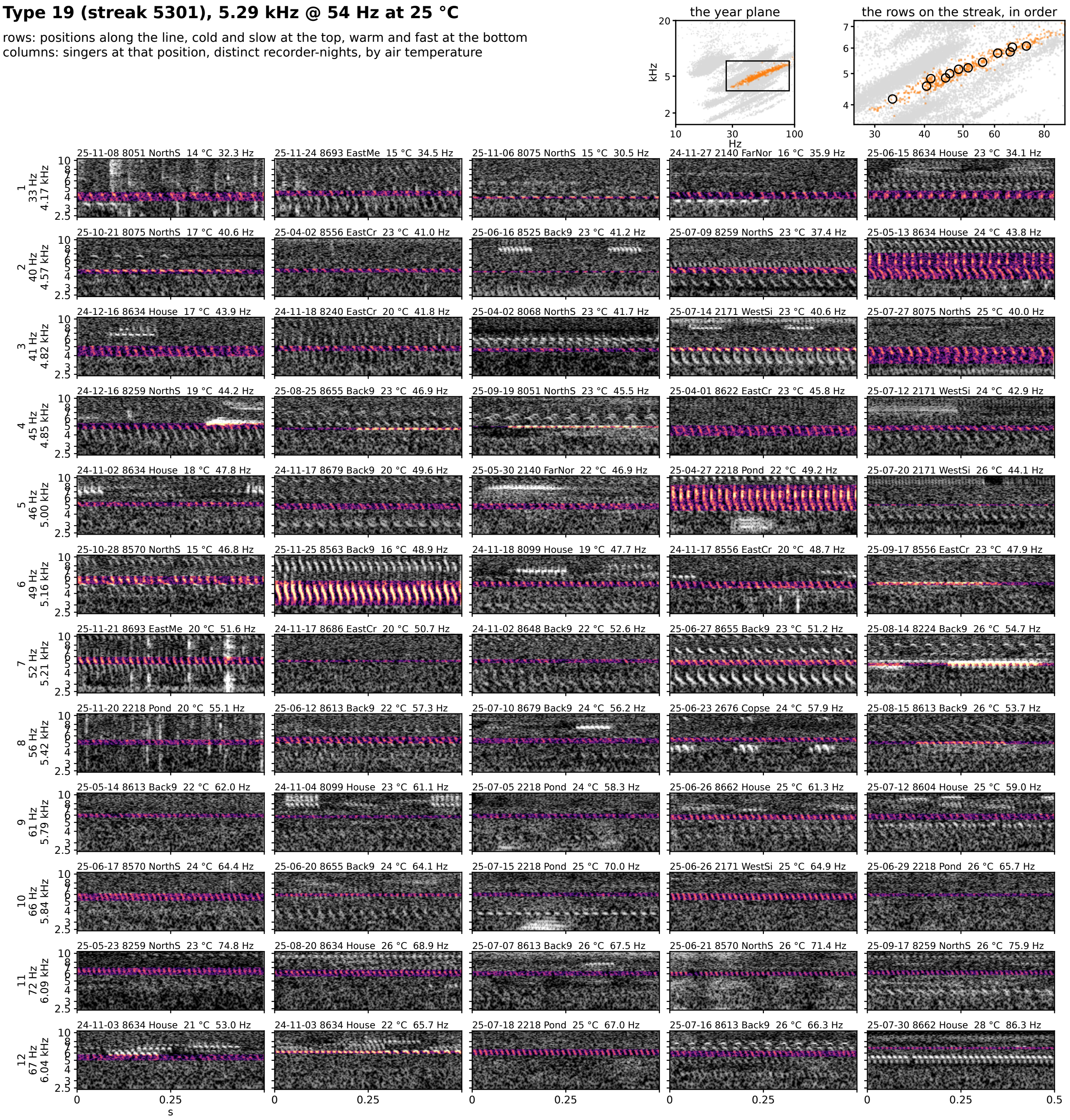


**Figure 7S-19.** Type 19 (streak 5301): 5.29 kHz @ 54 Hz at 25 °C, 117 singer-nights. Rows are positions along the line from the cold, slow end at the top to the warm, fast end at the bottom; columns are singers at that position from distinct recorder-nights, ordered by air temperature. Each cell is 0.5 s of the loudest 30 s of the singer's deepest window on one shared log-frequency axis, the detected band in color (dB above the row's quiet level, 0 to 30) and the rest in grayscale. The cell head gives the night, recorder, site, air temperature and rate (c: chirp rate). Top right: the type on the plane, and the row positions on the streak, in order.

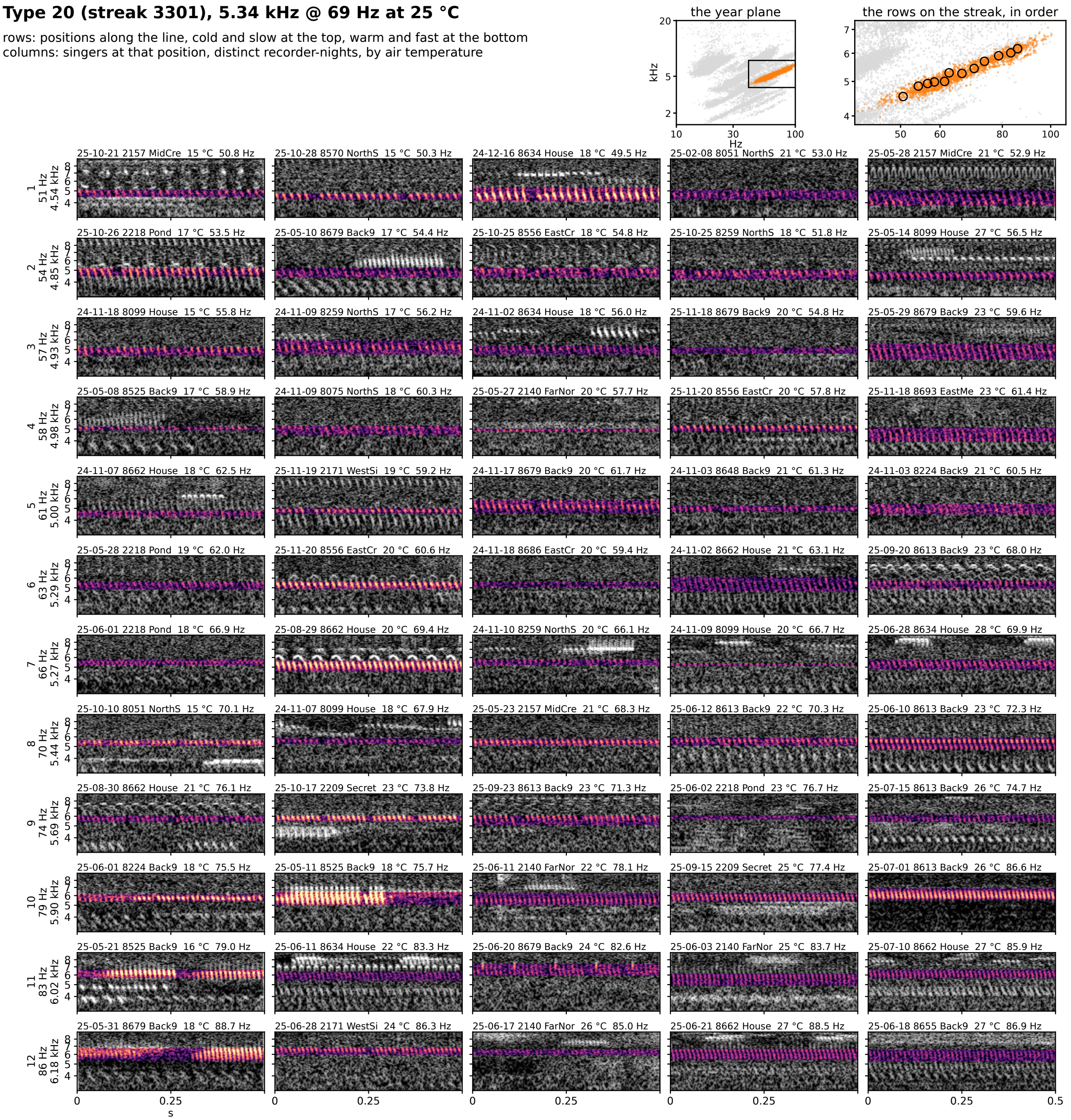


**Figure 7S-20.** Type 20 (streak 3301): 5.34 kHz @ 69 Hz at 25 °C, 374 singer-nights. Rows are positions along the line from the cold, slow end at the top to the warm, fast end at the bottom; columns are singers at that position from distinct recorder-nights, ordered by air temperature. Each cell is 0.5 s of the loudest 30 s of the singer's deepest window on one shared log-frequency axis, the detected band in color (dB above the row's quiet level, 0 to 30) and the rest in grayscale. The cell head gives the night, recorder, site, air temperature and rate (c: chirp rate). Top right: the type on the plane, and the row positions on the streak, in order.

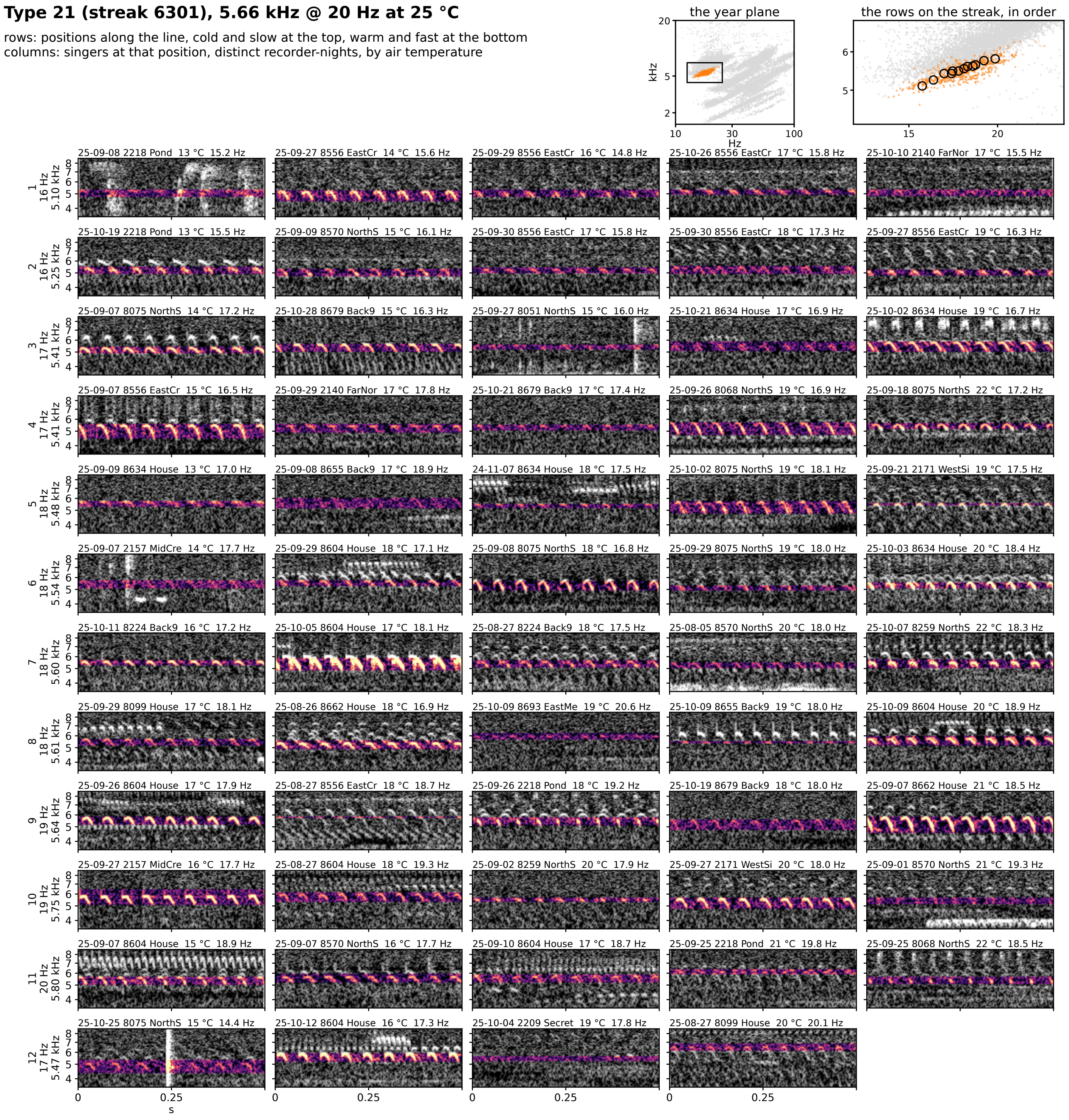


**Figure 7S-21.** Type 21 (streak 6301): 5.66 kHz @ 20 Hz at 25 °C, 117 singer-nights. Rows are positions along the line from the cold, slow end at the top to the warm, fast end at the bottom; columns are singers at that position from distinct recorder-nights, ordered by air temperature. Each cell is 0.5 s of the loudest 30 s of the singer's deepest window on one shared log-frequency axis, the detected band in color (dB above the row's quiet level, 0 to 30) and the rest in grayscale. The cell head gives the night, recorder, site, air temperature and rate (c: chirp rate). Top right: the type on the plane, and the row positions on the streak, in order.

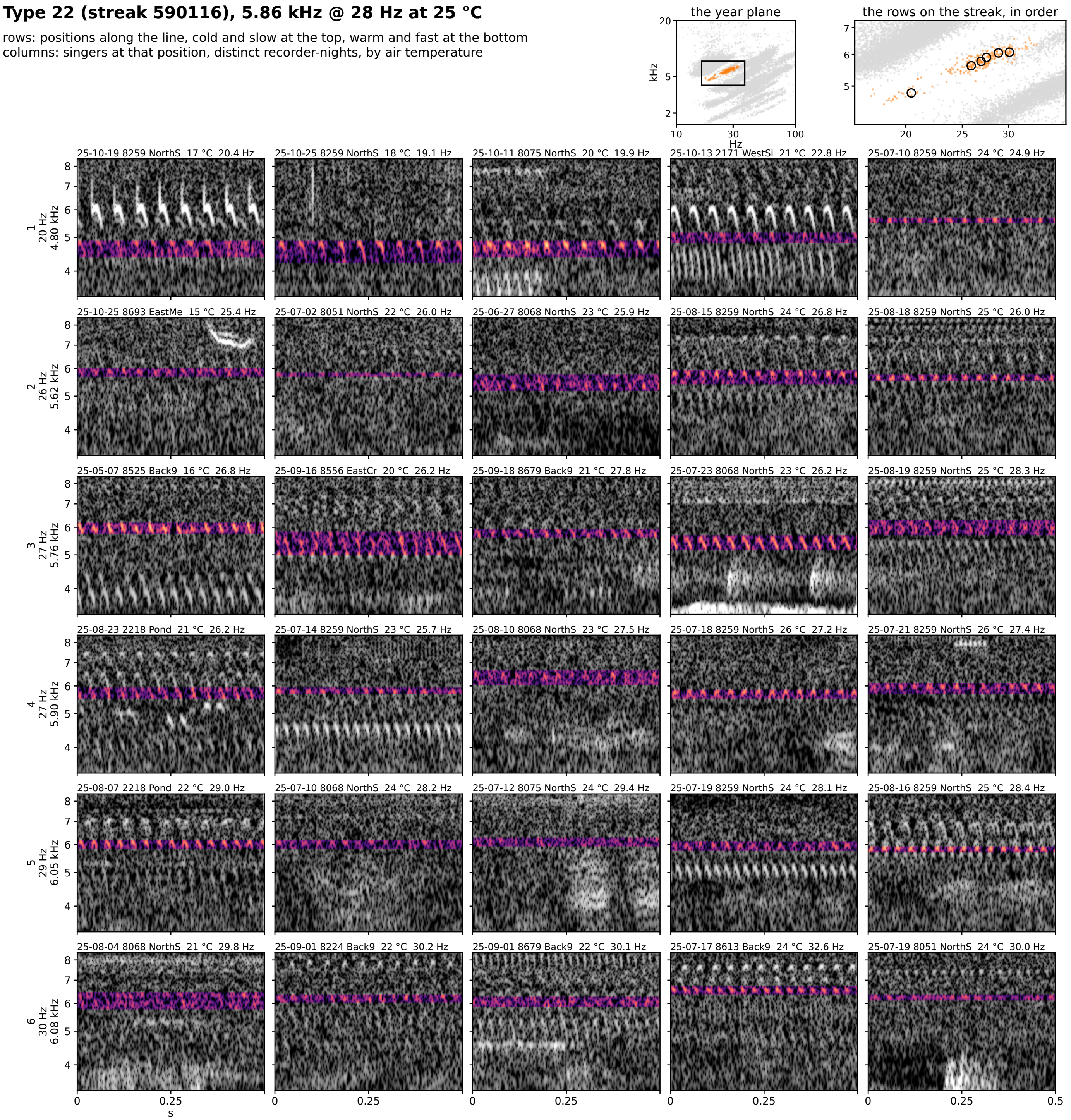


**Figure 7S-22.** Type 22 (streak 590116): 5.86 kHz @ 28 Hz at 25 °C, 54 singer-nights. Rows are positions along the line from the cold, slow end at the top to the warm, fast end at the bottom; columns are singers at that position from distinct recorder-nights, ordered by air temperature. Each cell is 0.5 s of the loudest 30 s of the singer's deepest window on one shared log-frequency axis, the detected band in color (dB above the row's quiet level, 0 to 30) and the rest in grayscale. The cell head gives the night, recorder, site, air temperature and rate (c: chirp rate). Top right: the type on the plane, and the row positions on the streak, in order.

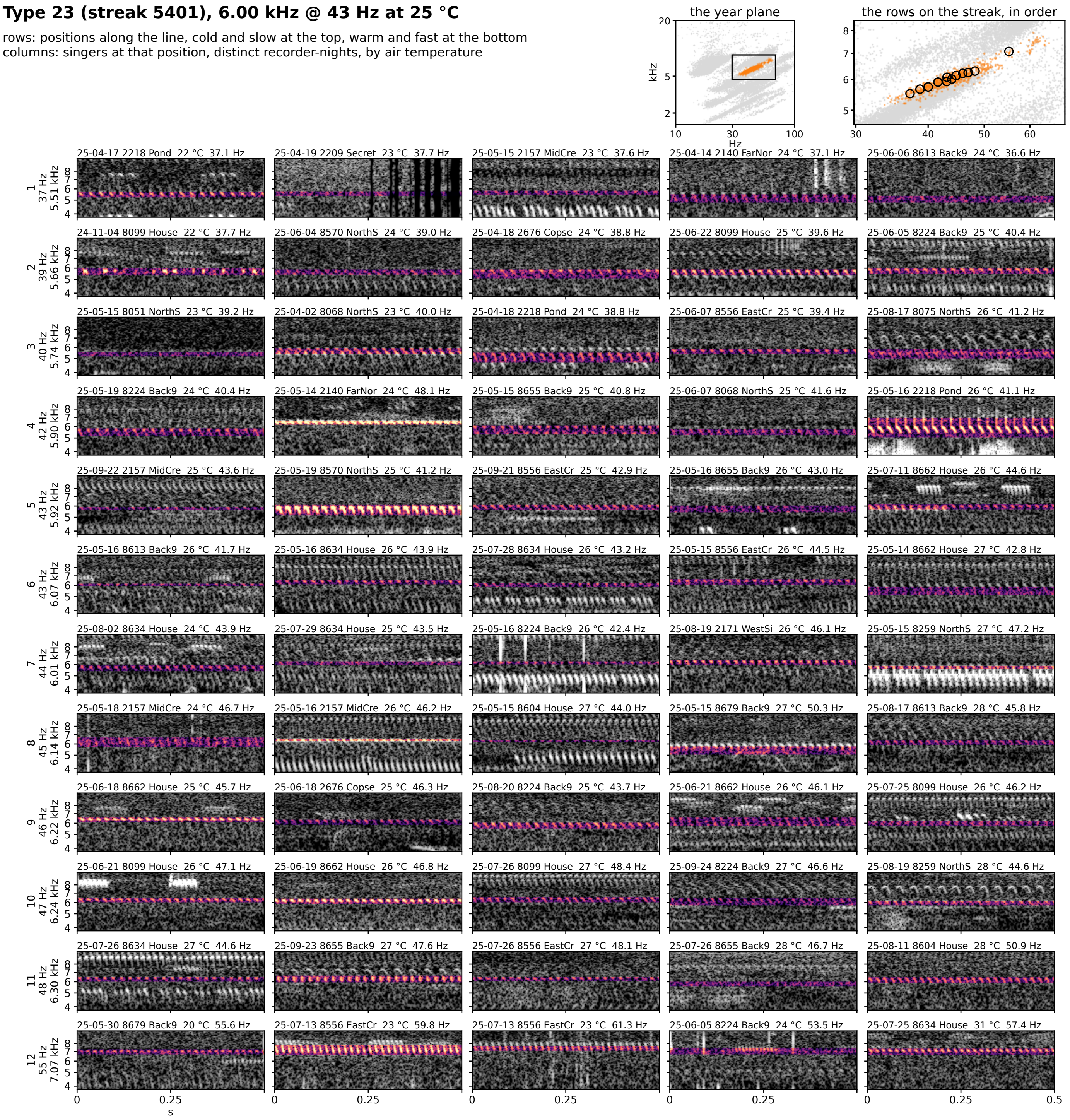


**Figure 7S-23.** Type 23 (streak 5401): 6.00 kHz @ 43 Hz at 25 °C, 143 singer-nights. Rows are positions along the line from the cold, slow end at the top to the warm, fast end at the bottom; columns are singers at that position from distinct recorder-nights, ordered by air temperature. Each cell is 0.5 s of the loudest 30 s of the singer's deepest window on one shared log-frequency axis, the detected band in color (dB above the row's quiet level, 0 to 30) and the rest in grayscale. The cell head gives the night, recorder, site, air temperature and rate (c: chirp rate). Top right: the type on the plane, and the row positions on the streak, in order.

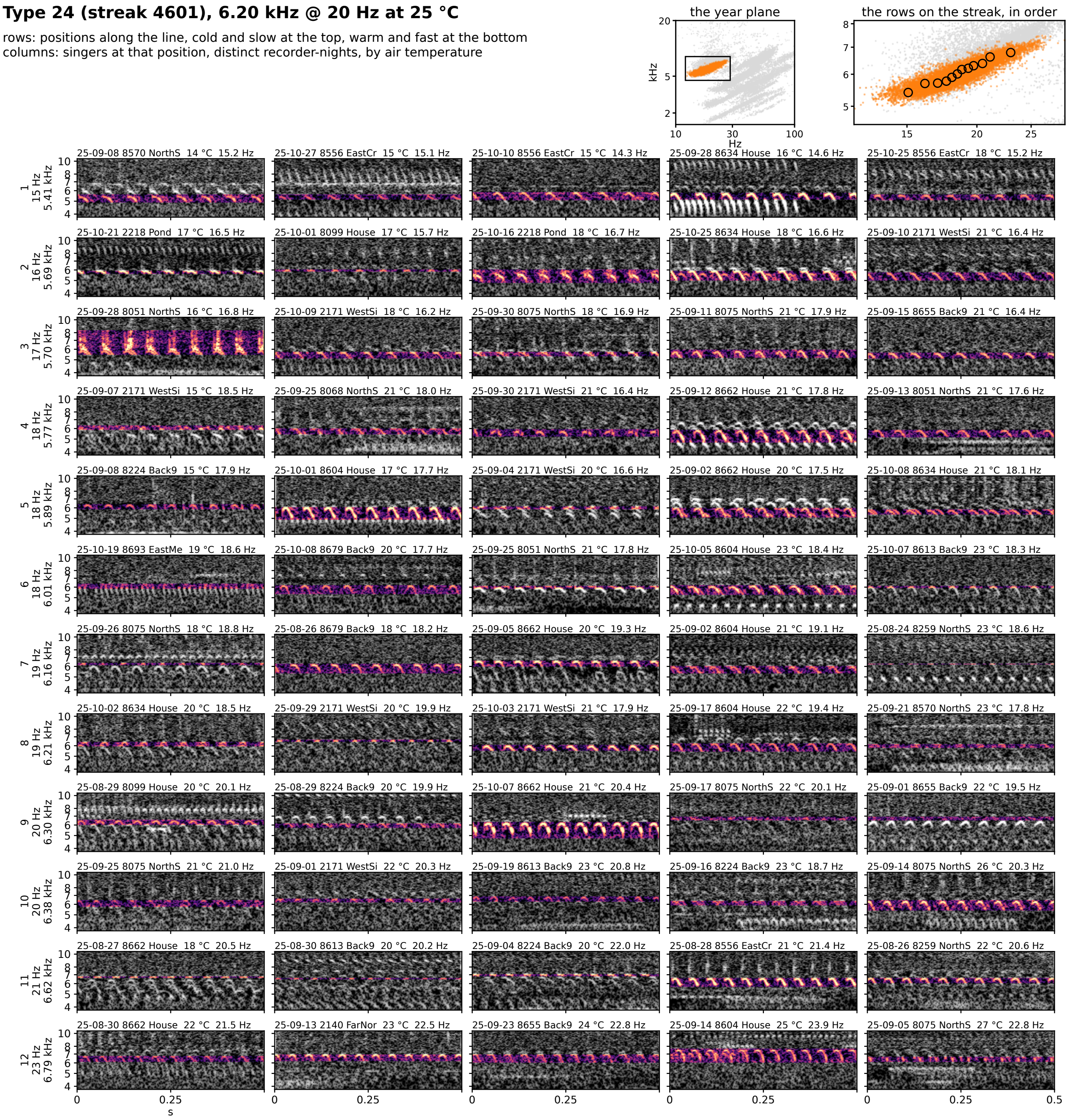


**Figure 7S-24.** Type 24 (streak 4601): 6.20 kHz @ 20 Hz at 25 °C, 1908 singer-nights. Rows are positions along the line from the cold, slow end at the top to the warm, fast end at the bottom; columns are singers at that position from distinct recorder-nights, ordered by air temperature. Each cell is 0.5 s of the loudest 30 s of the singer's deepest window on one shared log-frequency axis, the detected band in color (dB above the row's quiet level, 0 to 30) and the rest in grayscale. The cell head gives the night, recorder, site, air temperature and rate (c: chirp rate). Top right: the type on the plane, and the row positions on the streak, in order.

### Type 25 (streak 3402), 6.21 kHz @ 81 Hz at 25 °C

rows: positions along the line, cold and slow at the top, warm and fast at the bottom
columns: singers at that position, distinct recorder-nights, by air temperature

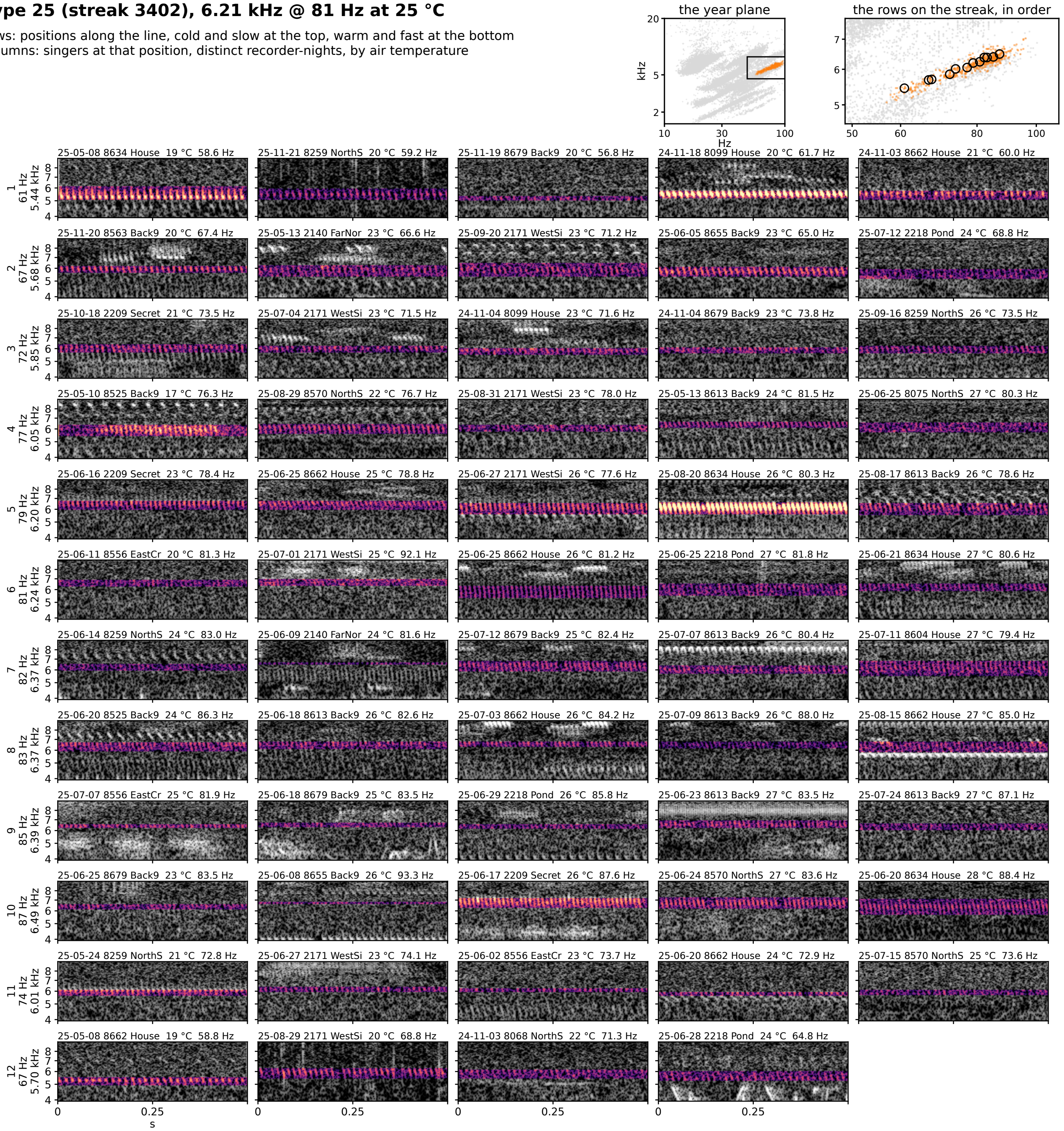


**Figure 7S-25.** Type 25 (streak 3402): 6.21 kHz @ 81 Hz at 25 °C, 105 singer-nights. Rows are positions along the line from the cold, slow end at the top to the warm, fast end at the bottom; columns are singers at that position from distinct recorder-nights, ordered by air temperature. Each cell is 0.5 s of the loudest 30 s of the singer's deepest window on one shared log-frequency axis, the detected band in color (dB above the row's quiet level, 0 to 30) and the rest in grayscale. The cell head gives the night, recorder, site, air temperature and rate (c: chirp rate). Top right: the type on the plane, and the row positions on the streak, in order.

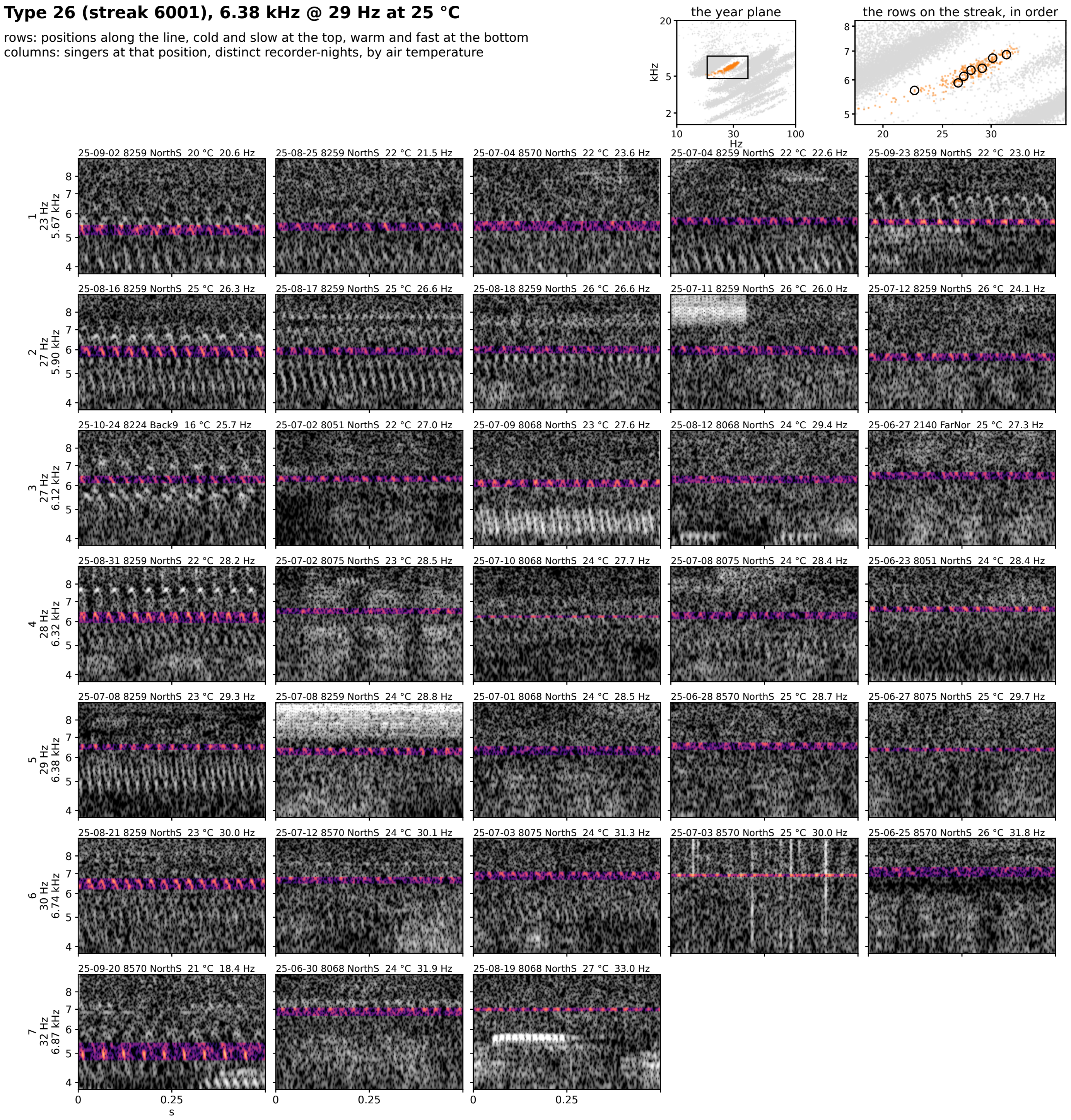


**Figure 7S-26.** Type 26 (streak 6001): 6.38 kHz @ 29 Hz at 25 °C, 61 singer-nights. Rows are positions along the line from the cold, slow end at the top to the warm, fast end at the bottom; columns are singers at that position from distinct recorder-nights, ordered by air temperature. Each cell is 0.5 s of the loudest 30 s of the singer's deepest window on one shared log-frequency axis, the detected band in color (dB above the row's quiet level, 0 to 30) and the rest in grayscale. The cell head gives the night, recorder, site, air temperature and rate (c: chirp rate). Top right: the type on the plane, and the row positions on the streak, in order.

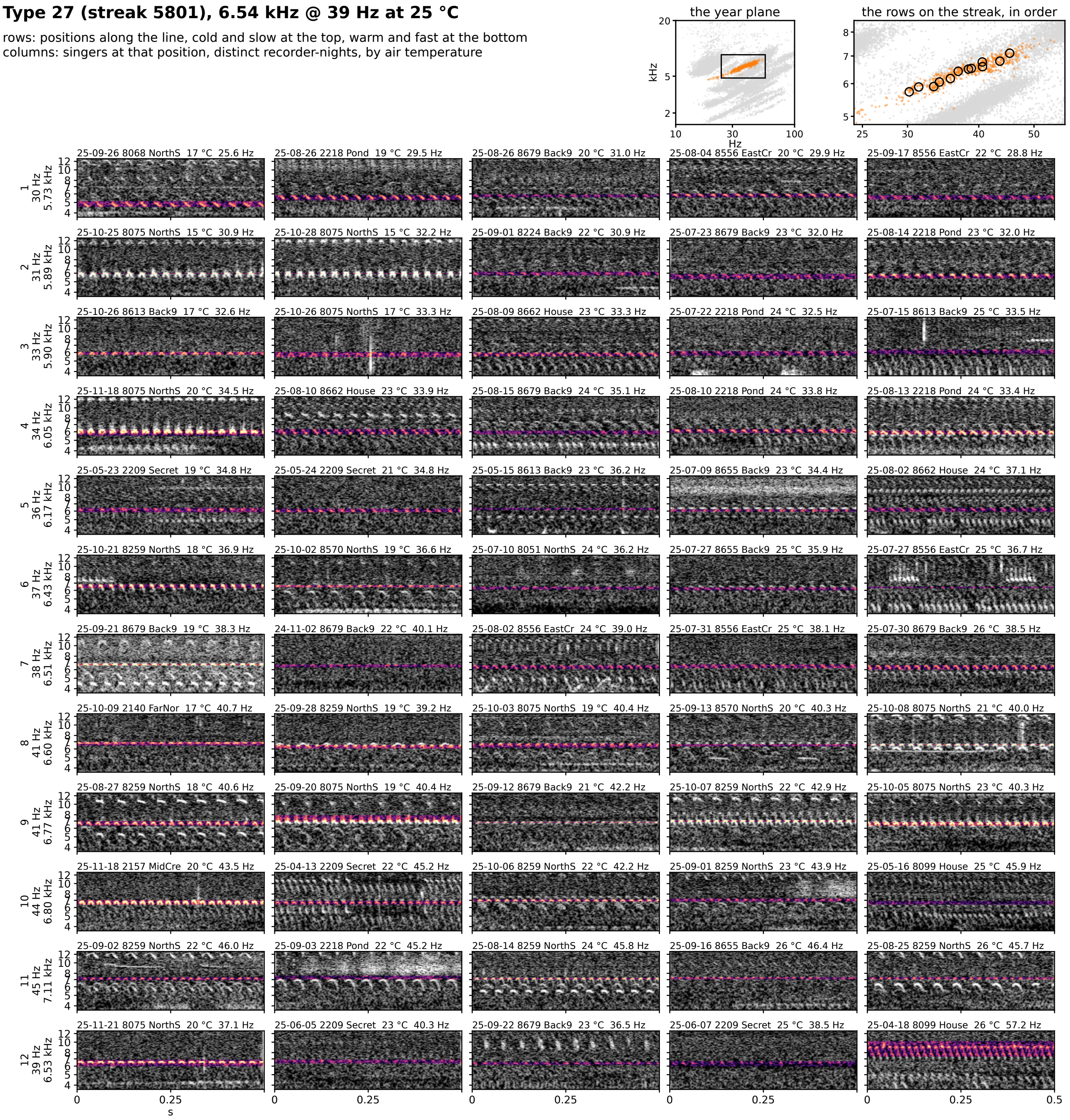


**Figure 7S-27.** Type 27 (streak 5801): 6.54 kHz @ 39 Hz at 25 °C, 124 singer-nights. Rows are positions along the line from the cold, slow end at the top to the warm, fast end at the bottom; columns are singers at that position from distinct recorder-nights, ordered by air temperature. Each cell is 0.5 s of the loudest 30 s of the singer's deepest window on one shared log-frequency axis, the detected band in color (dB above the row's quiet level, 0 to 30) and the rest in grayscale. The cell head gives the night, recorder, site, air temperature and rate (c: chirp rate). Top right: the type on the plane, and the row positions on the streak, in order.

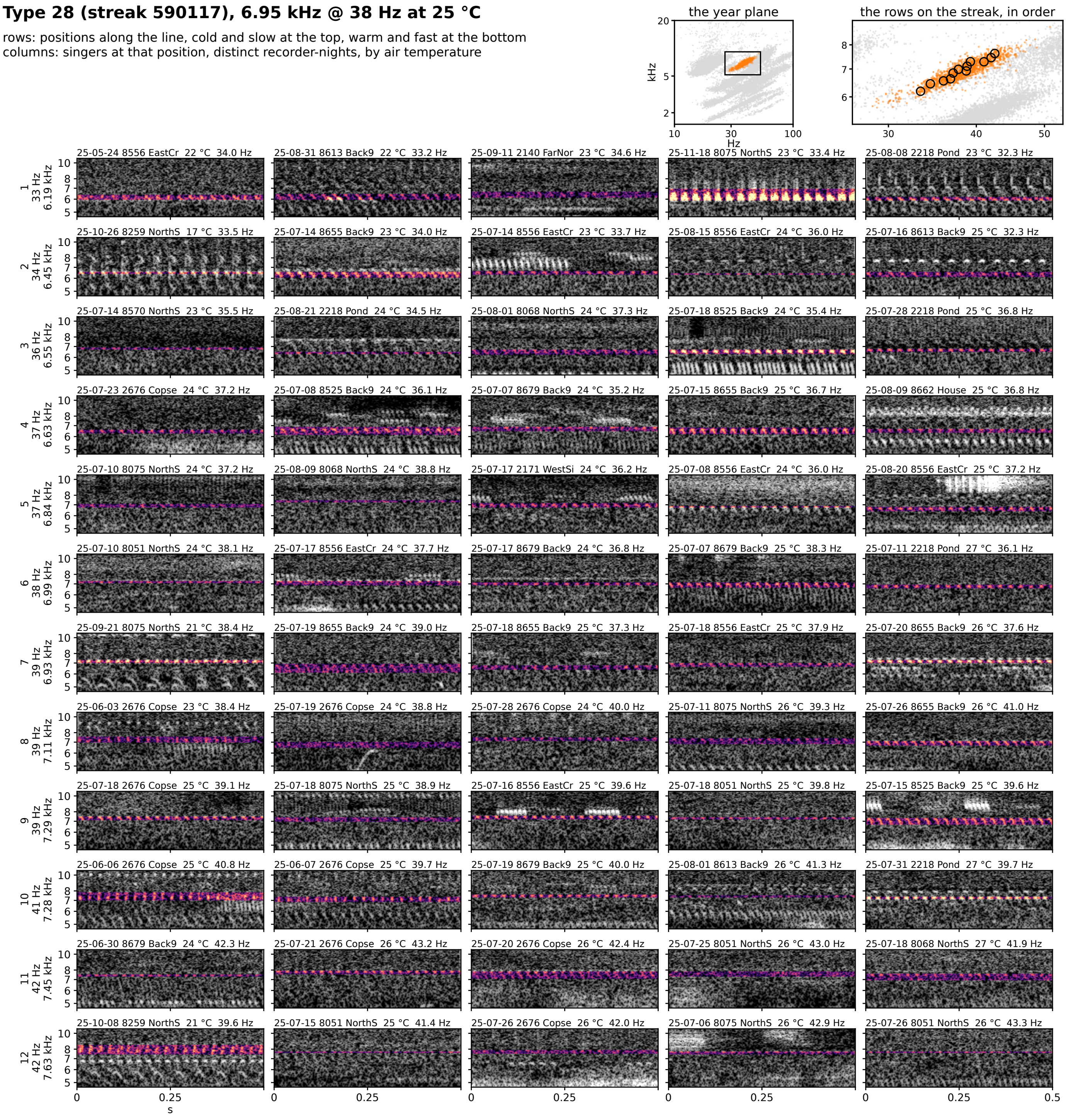


**Figure 7S-28.** Type 28 (streak 590117): 6.95 kHz @ 38 Hz at 25 °C, 273 singer-nights. Rows are positions along the line from the cold, slow end at the top to the warm, fast end at the bottom; columns are singers at that position from distinct recorder-nights, ordered by air temperature. Each cell is 0.5 s of the loudest 30 s of the singer's deepest window on one shared log-frequency axis, the detected band in color (dB above the row's quiet level, 0 to 30) and the rest in grayscale. The cell head gives the night, recorder, site, air temperature and rate (c: chirp rate). Top right: the type on the plane, and the row positions on the streak, in order.

### Type 29 (streak 54012), 6.96 kHz @ 51 Hz at 25 °C

rows: positions along the line, cold and slow at the top, warm and fast at the bottom
columns: singers at that position, distinct recorder-nights, by air temperature

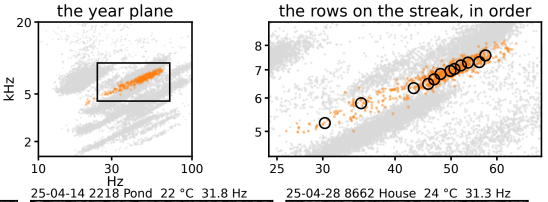


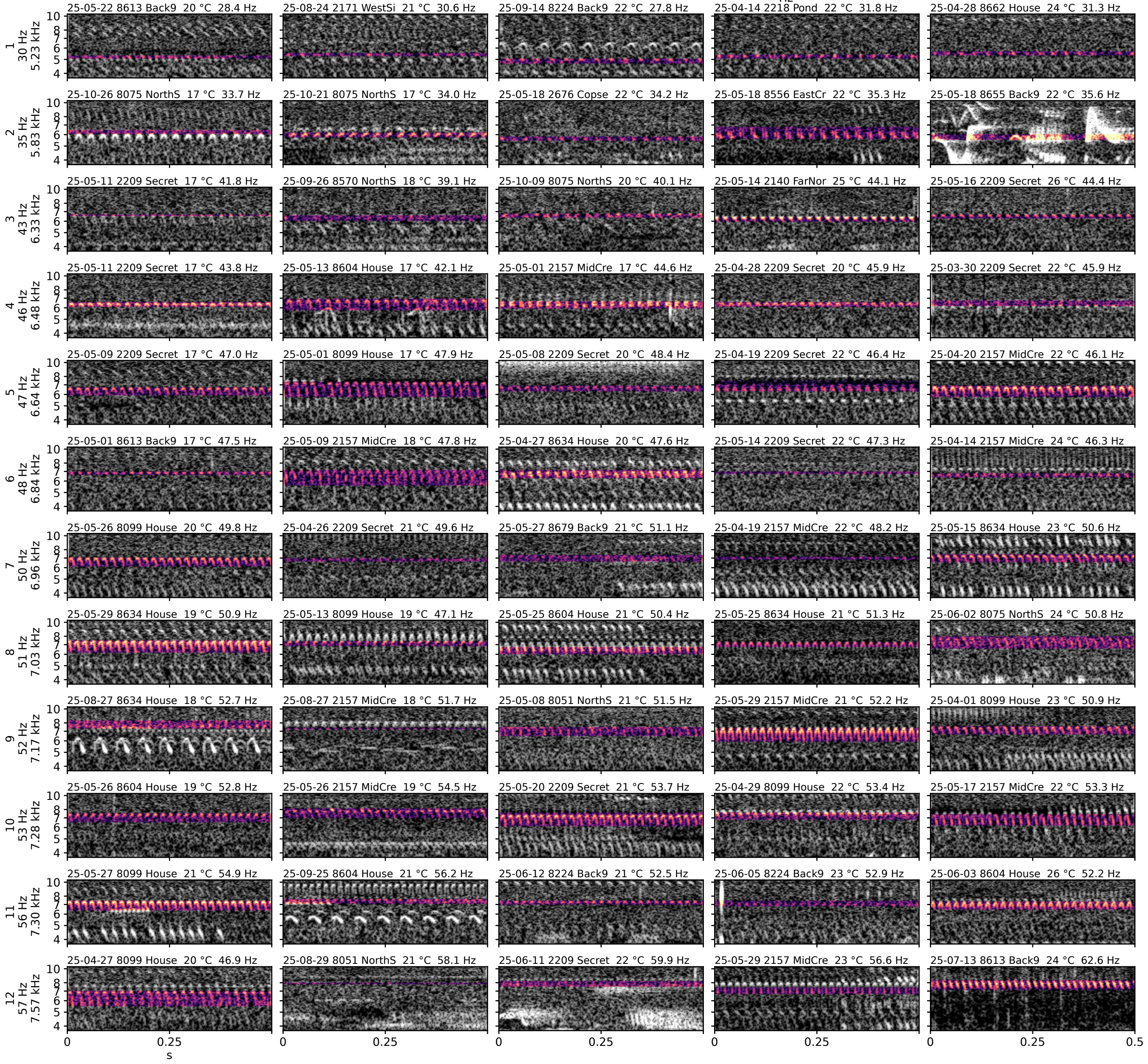


**Figure 7S-29.** Type 29 (streak 54012): 6.96 kHz @ 51 Hz at 25 °C, 128 singer-nights. Rows are positions along the line from the cold, slow end at the top to the warm, fast end at the bottom; columns are singers at that position from distinct recorder-nights, ordered by air temperature. Each cell is 0.5 s of the loudest 30 s of the singer's deepest window on one shared log-frequency axis, the detected band in color (dB above the row's quiet level, 0 to 30) and the rest in grayscale. The cell head gives the night, recorder, site, air temperature and rate (c: chirp rate). Top right: the type on the plane, and the row positions on the streak, in order.

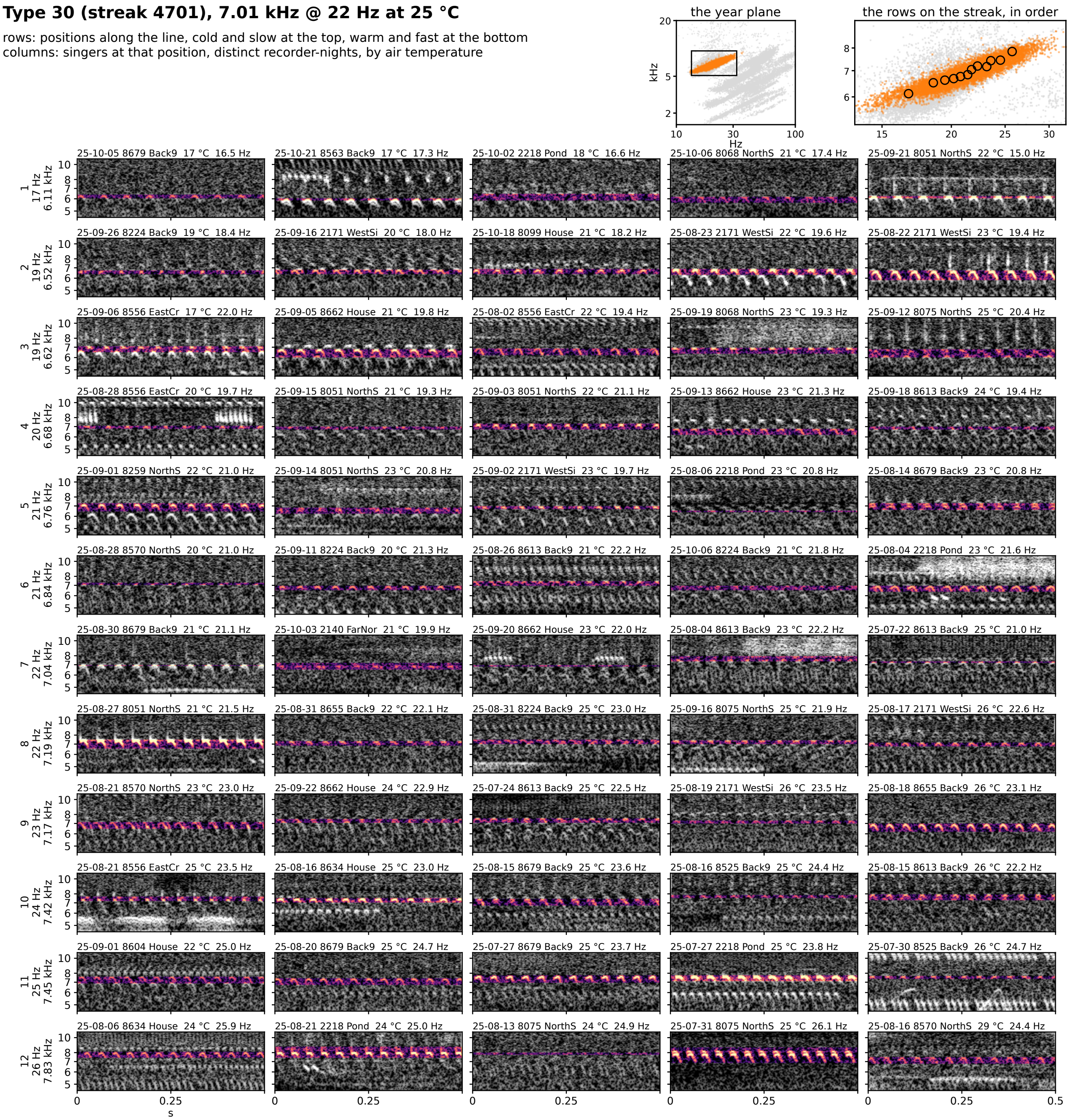

**Figure 7S-30.** Type 30 (streak 4701): 7.01 kHz @ 22 Hz at 25 °C, 2080 singer-nights. Rows are positions along the line from the cold, slow end at the top to the warm, fast end at the bottom; columns are singers at that position from distinct recorder-nights, ordered by air temperature. Each cell is 0.5 s of the loudest 30 s of the singer's deepest window on one shared log-frequency axis, the detected band in color (dB above the row's quiet level, 0 to 30) and the rest in grayscale. The cell head gives the night, recorder, site, air temperature and rate (c: chirp rate). Top right: the type on the plane, and the row positions on the streak, in order.

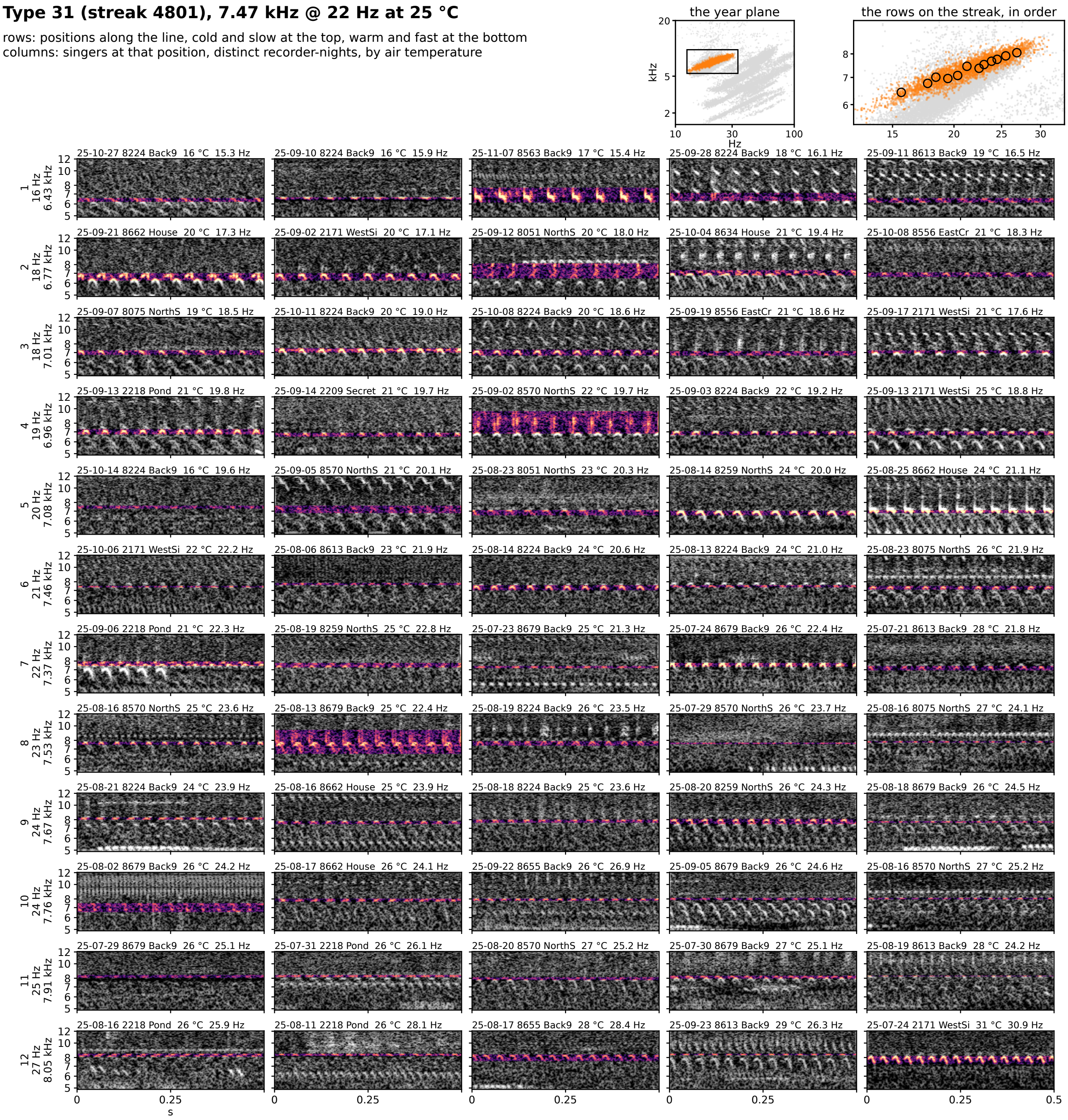


**Figure 7S-31.** Type 31 (streak 4801): 7.47 kHz @ 22 Hz at 25 °C, 623 singer-nights. Rows are positions along the line from the cold, slow end at the top to the warm, fast end at the bottom; columns are singers at that position from distinct recorder-nights, ordered by air temperature. Each cell is 0.5 s of the loudest 30 s of the singer's deepest window on one shared log-frequency axis, the detected band in color (dB above the row's quiet level, 0 to 30) and the rest in grayscale. The cell head gives the night, recorder, site, air temperature and rate (c: chirp rate). Top right: the type on the plane, and the row positions on the streak, in order.

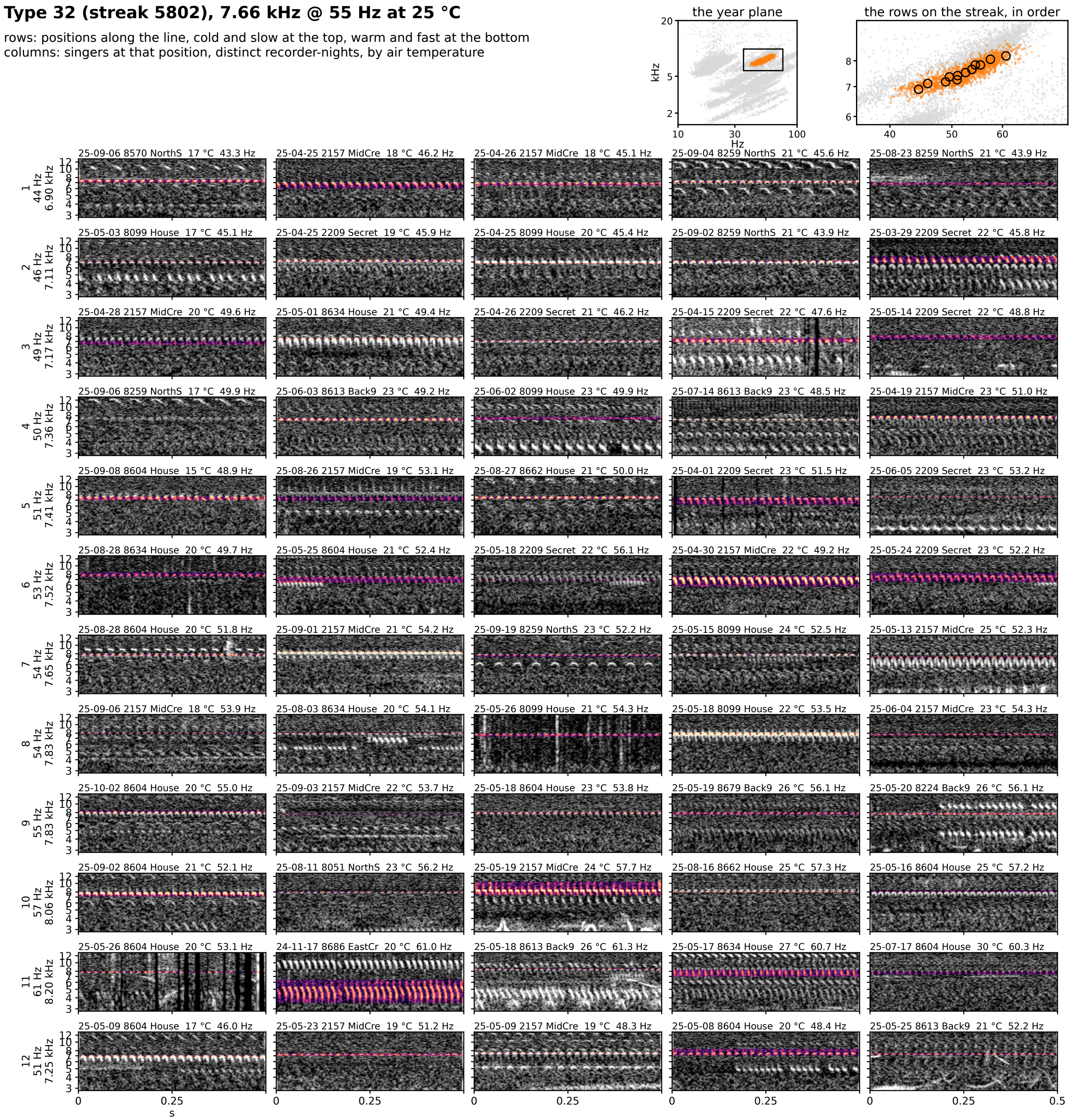


**Figure 7S-32.** Type 32 (streak 5802): 7.66 kHz @ 55 Hz at 25 °C, 375 singer-nights. Rows are positions along the line from the cold, slow end at the top to the warm, fast end at the bottom; columns are singers at that position from distinct recorder-nights, ordered by air temperature. Each cell is 0.5 s of the loudest 30 s of the singer's deepest window on one shared log-frequency axis, the detected band in color (dB above the row's quiet level, 0 to 30) and the rest in grayscale. The cell head gives the night, recorder, site, air temperature and rate (c: chirp rate). Top right: the type on the plane, and the row positions on the streak, in order.

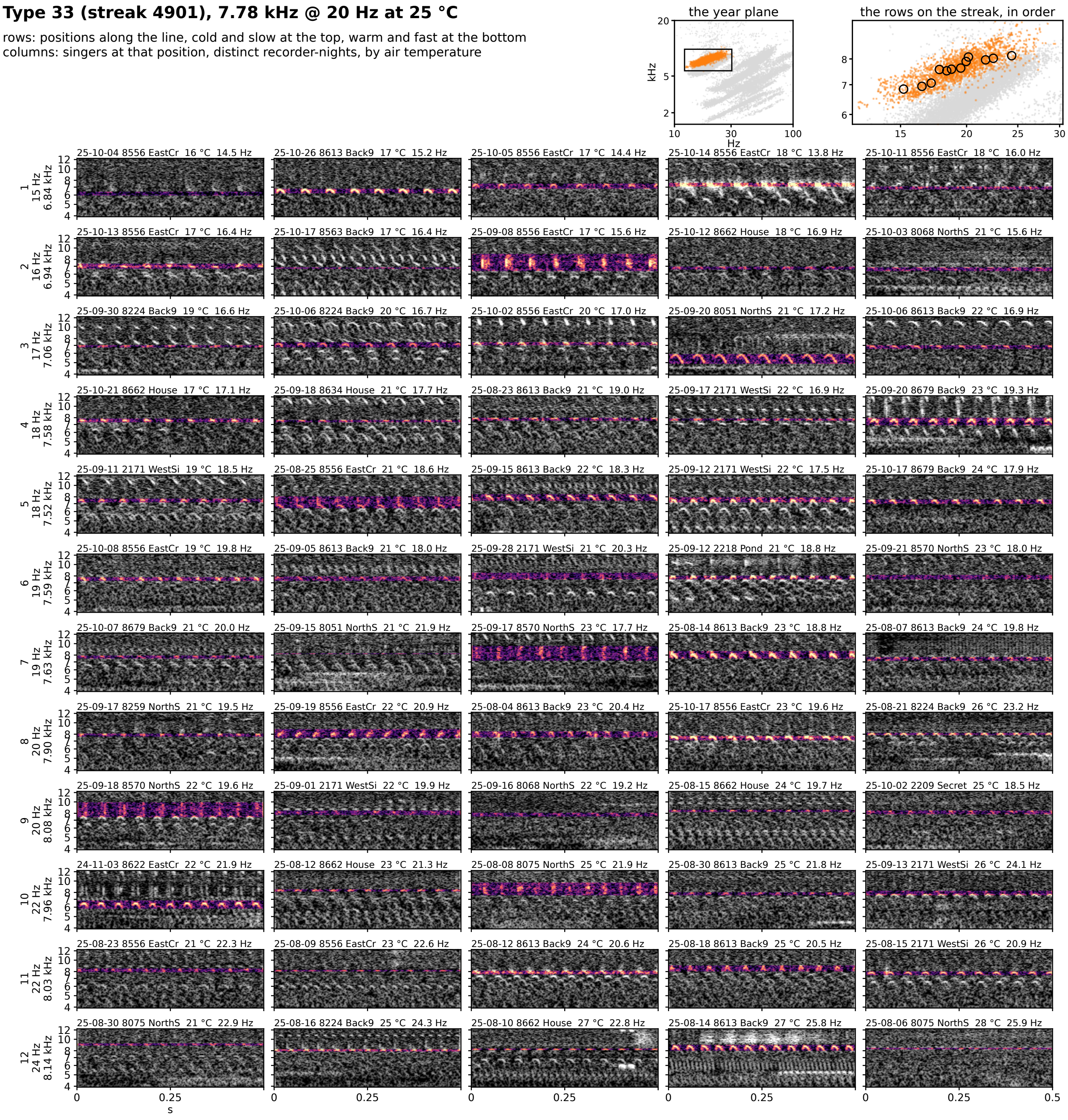


**Figure 7S-33.** Type 33 (streak 4901): 7.78 kHz @ 20 Hz at 25 °C, 391 singer-nights. Rows are positions along the line from the cold, slow end at the top to the warm, fast end at the bottom; columns are singers at that position from distinct recorder-nights, ordered by air temperature. Each cell is 0.5 s of the loudest 30 s of the singer's deepest window on one shared log-frequency axis, the detected band in color (dB above the row's quiet level, 0 to 30) and the rest in grayscale. The cell head gives the night, recorder, site, air temperature and rate (c: chirp rate). Top right: the type on the plane, and the row positions on the streak, in order.

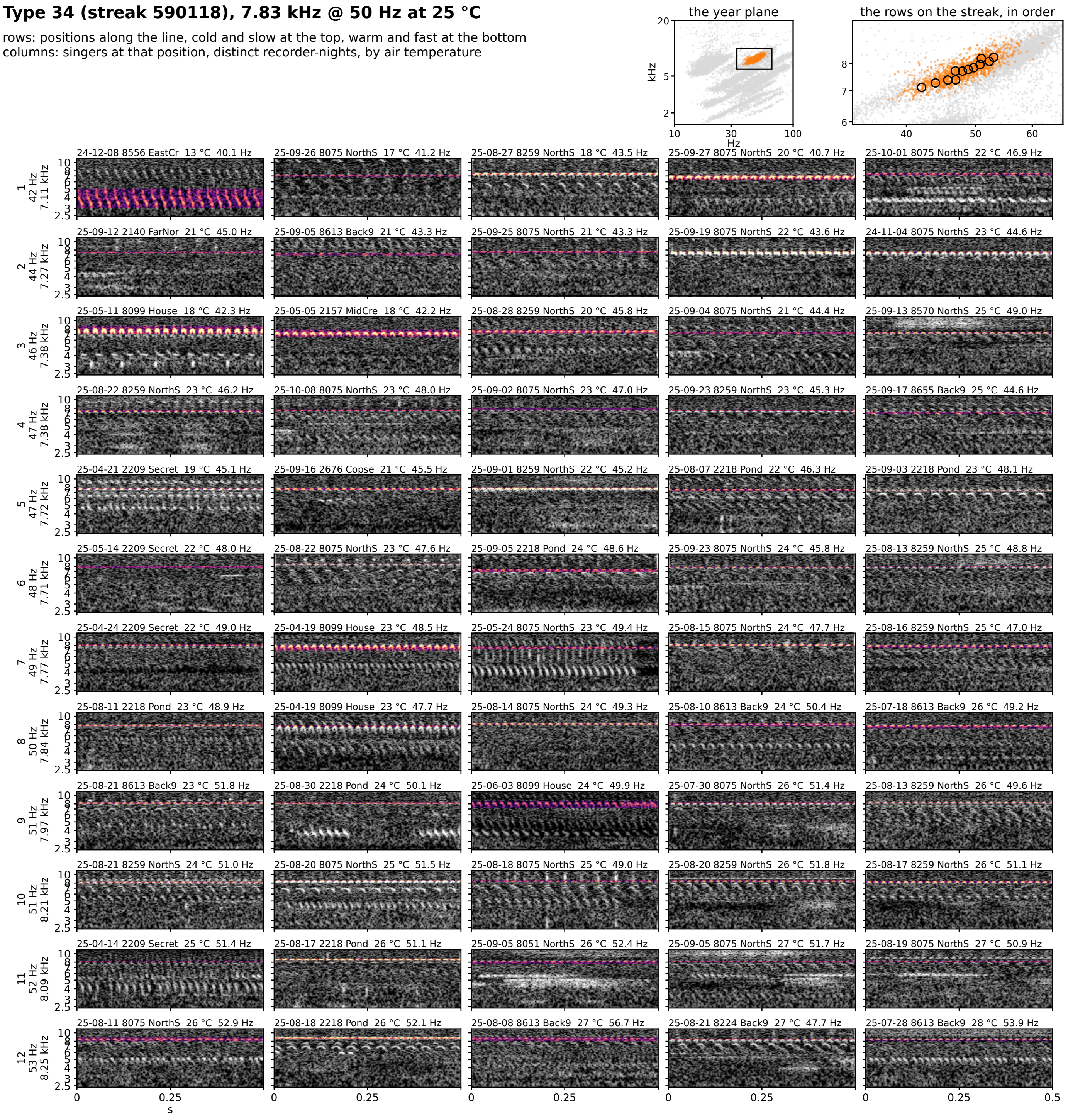


**Figure 7S-34.** Type 34 (streak 590118): 7.83 kHz @ 50 Hz at 25 °C, 300 singer-nights. Rows are positions along the line from the cold, slow end at the top to the warm, fast end at the bottom; columns are singers at that position from distinct recorder-nights, ordered by air temperature. Each cell is 0.5 s of the loudest 30 s of the singer's deepest window on one shared log-frequency axis, the detected band in color (dB above the row's quiet level, 0 to 30) and the rest in grayscale. The cell head gives the night, recorder, site, air temperature and rate (c: chirp rate). Top right: the type on the plane, and the row positions on the streak, in order.

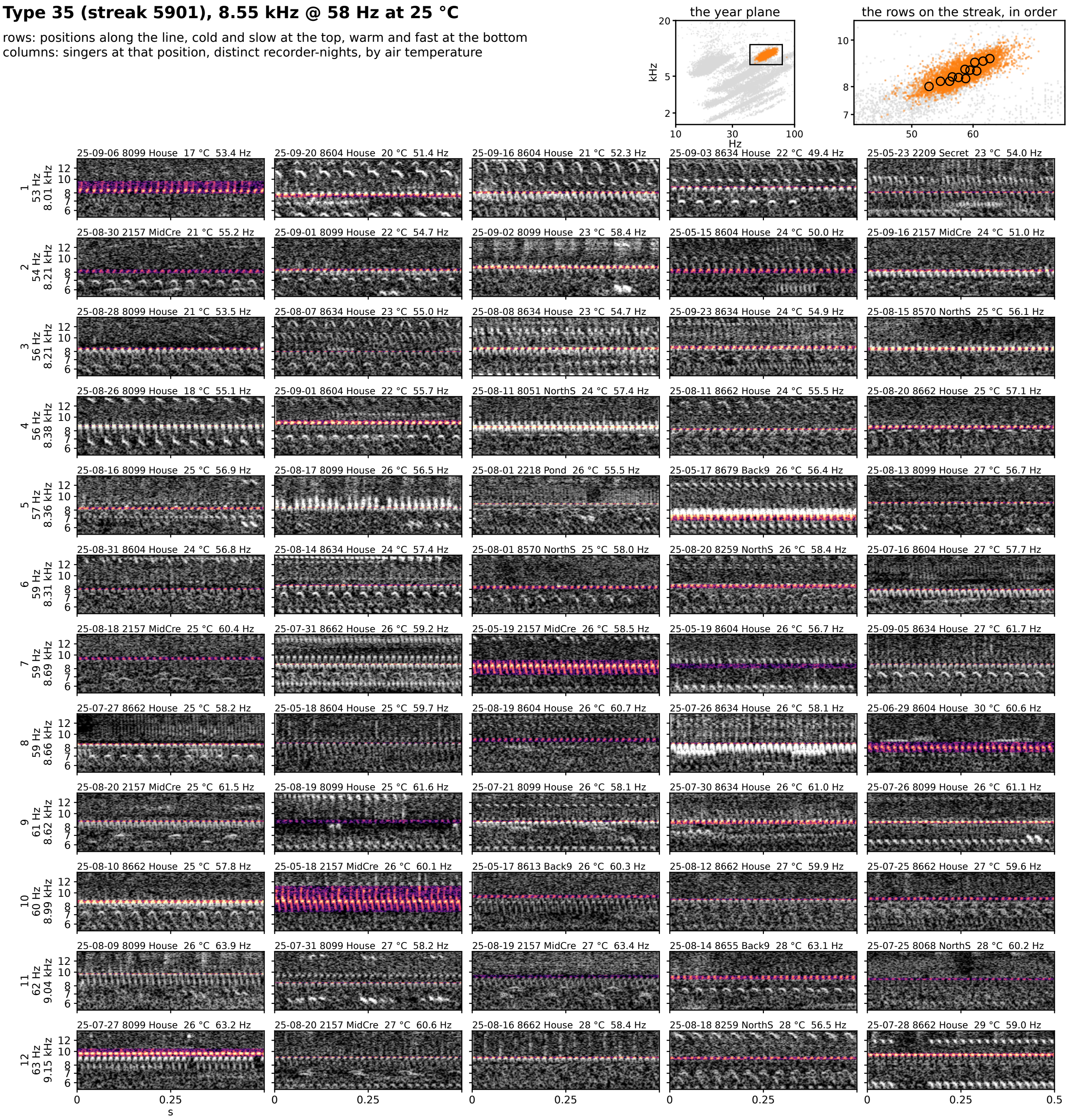


**Figure 7S-35.** Type 35 (streak 5901): 8.55 kHz @ 58 Hz at 25 °C, 826 singer-nights. Rows are positions along the line from the cold, slow end at the top to the warm, fast end at the bottom; columns are singers at that position from distinct recorder-nights, ordered by air temperature. Each cell is 0.5 s of the loudest 30 s of the singer's deepest window on one shared log-frequency axis, the detected band in color (dB above the row's quiet level, 0 to 30) and the rest in grayscale. The cell head gives the night, recorder, site, air temperature and rate (c: chirp rate). Top right: the type on the plane, and the row positions on the streak, in order.

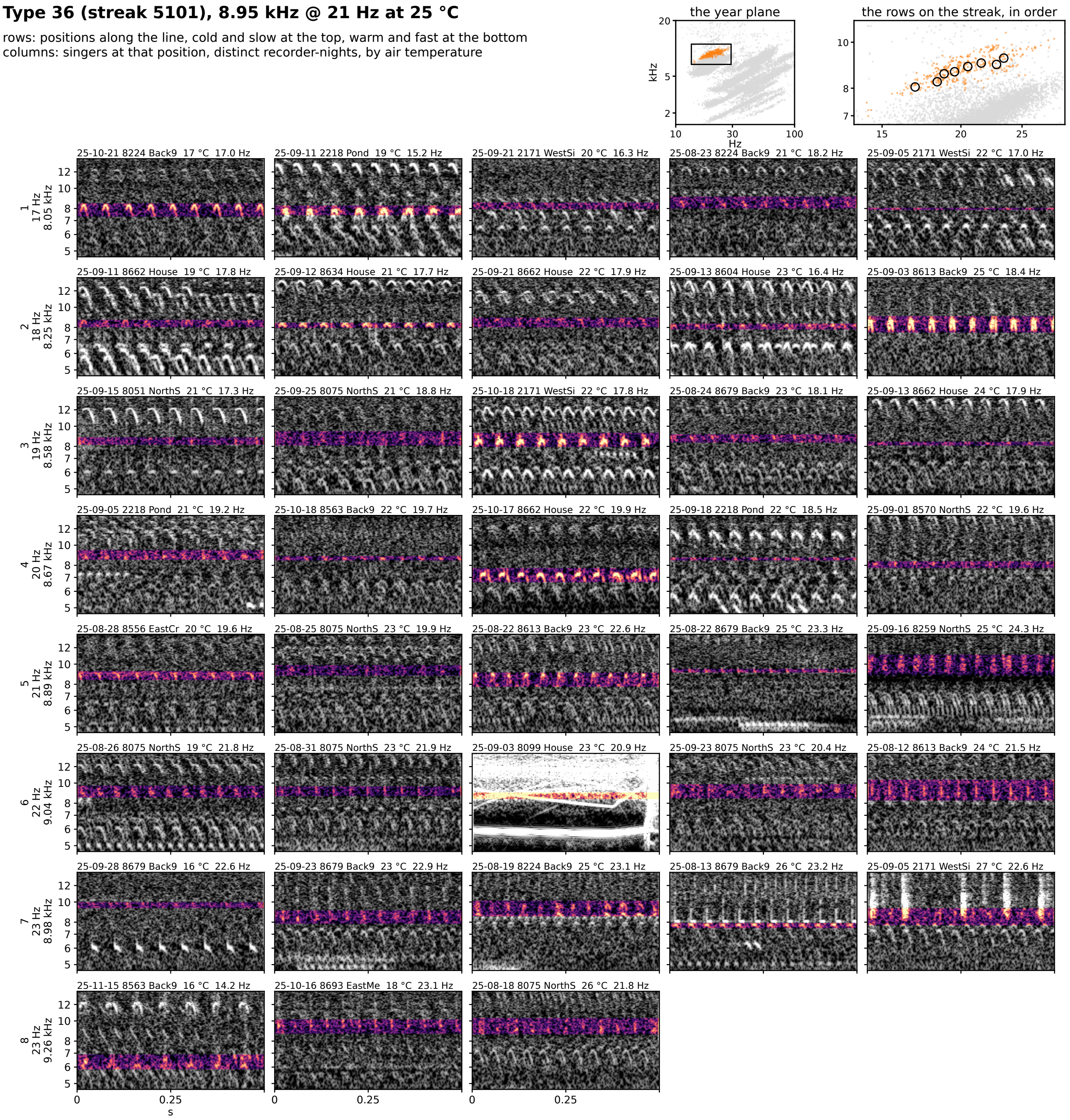


**Figure 7S-36.** Type 36 (streak 5101): 8.95 kHz @ 21 Hz at 25 °C, 69 singer-nights. Rows are positions along the line from the cold, slow end at the top to the warm, fast end at the bottom; columns are singers at that position from distinct recorder-nights, ordered by air temperature. Each cell is 0.5 s of the loudest 30 s of the singer's deepest window on one shared log-frequency axis, the detected band in color (dB above the row's quiet level, 0 to 30) and the rest in grayscale. The cell head gives the night, recorder, site, air temperature and rate (c: chirp rate). Top right: the type on the plane, and the row positions on the streak, in order.

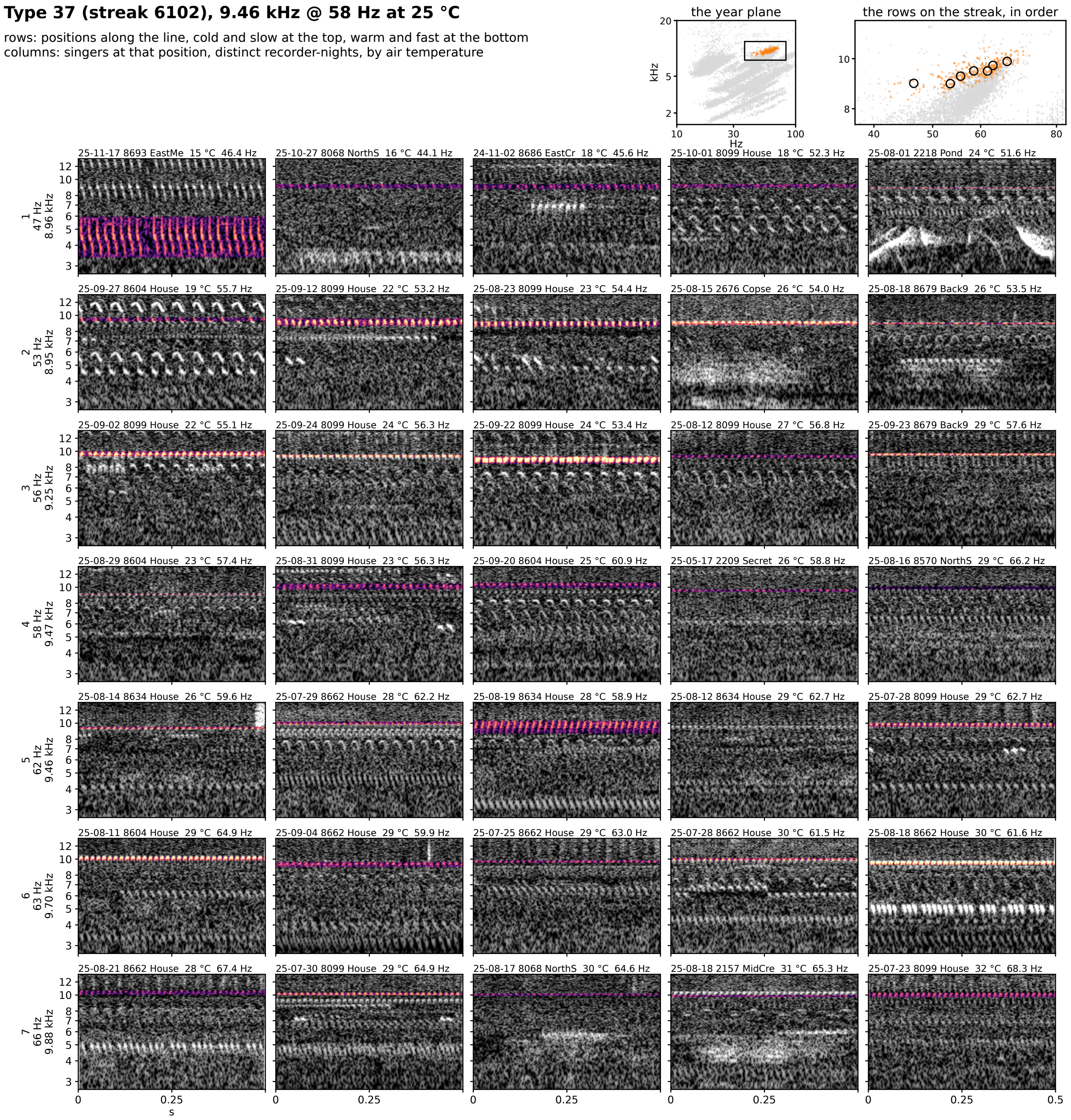


**Figure 7S-37.** Type 37 (streak 6102): 9.46 kHz @ 58 Hz at 25 °C, 64 singer-nights. Rows are positions along the line from the cold, slow end at the top to the warm, fast end at the bottom; columns are singers at that position from distinct recorder-nights, ordered by air temperature. Each cell is 0.5 s of the loudest 30 s of the singer's deepest window on one shared log-frequency axis, the detected band in color (dB above the row's quiet level, 0 to 30) and the rest in grayscale. The cell head gives the night, recorder, site, air temperature and rate (c: chirp rate). Top right: the type on the plane, and the row positions on the streak, in order.

**Streak 3151 (chirp-form; its type is on the chirp plane), 3.50 kHz @ 47 Hz at 25 °C**

rows: positions along the line, cold and slow at the top, warm and fast at the bottom
columns: singers at that position, distinct recorder-nights, by air temperature

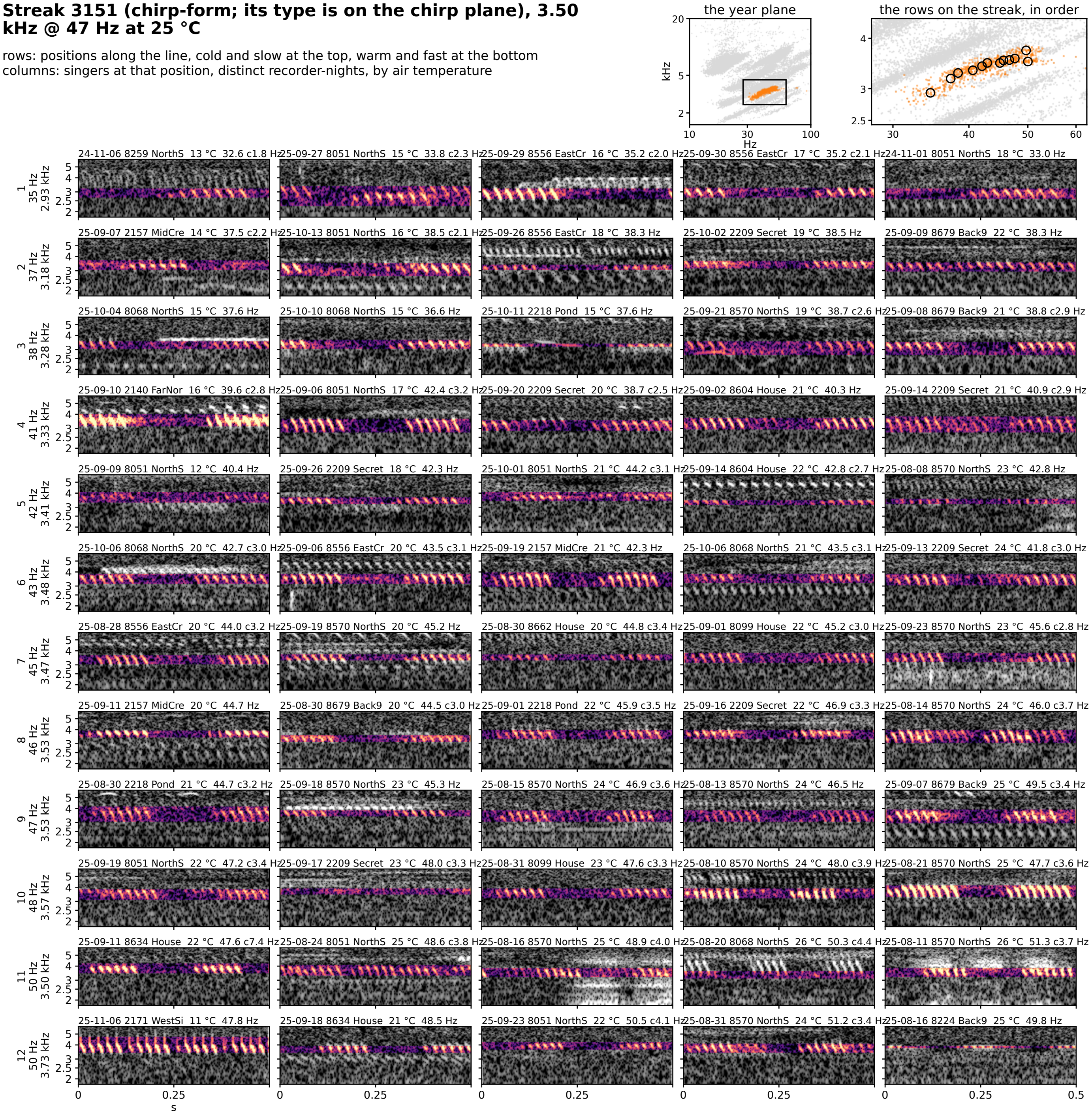


**Figure 7S-streak-3151.** Streak 3151 (chirp-form; its type is on the chirp plane): 3.50 kHz @ 47 Hz at 25 °C, 177 singer-nights. Rows are positions along the line from the cold, slow end at the top to the warm, fast end at the bottom; columns are singers at that position from distinct recorder-nights, ordered by air temperature. Each cell is 0.5 s of the loudest 30 s of the singer's deepest window on one shared log-frequency axis, the detected band in color (dB above the row's quiet level, 0 to 30) and the rest in grayscale. The cell head gives the night, recorder, site, air temperature and rate (c: chirp rate). Top right: the type on the plane, and the row positions on the streak, in order.

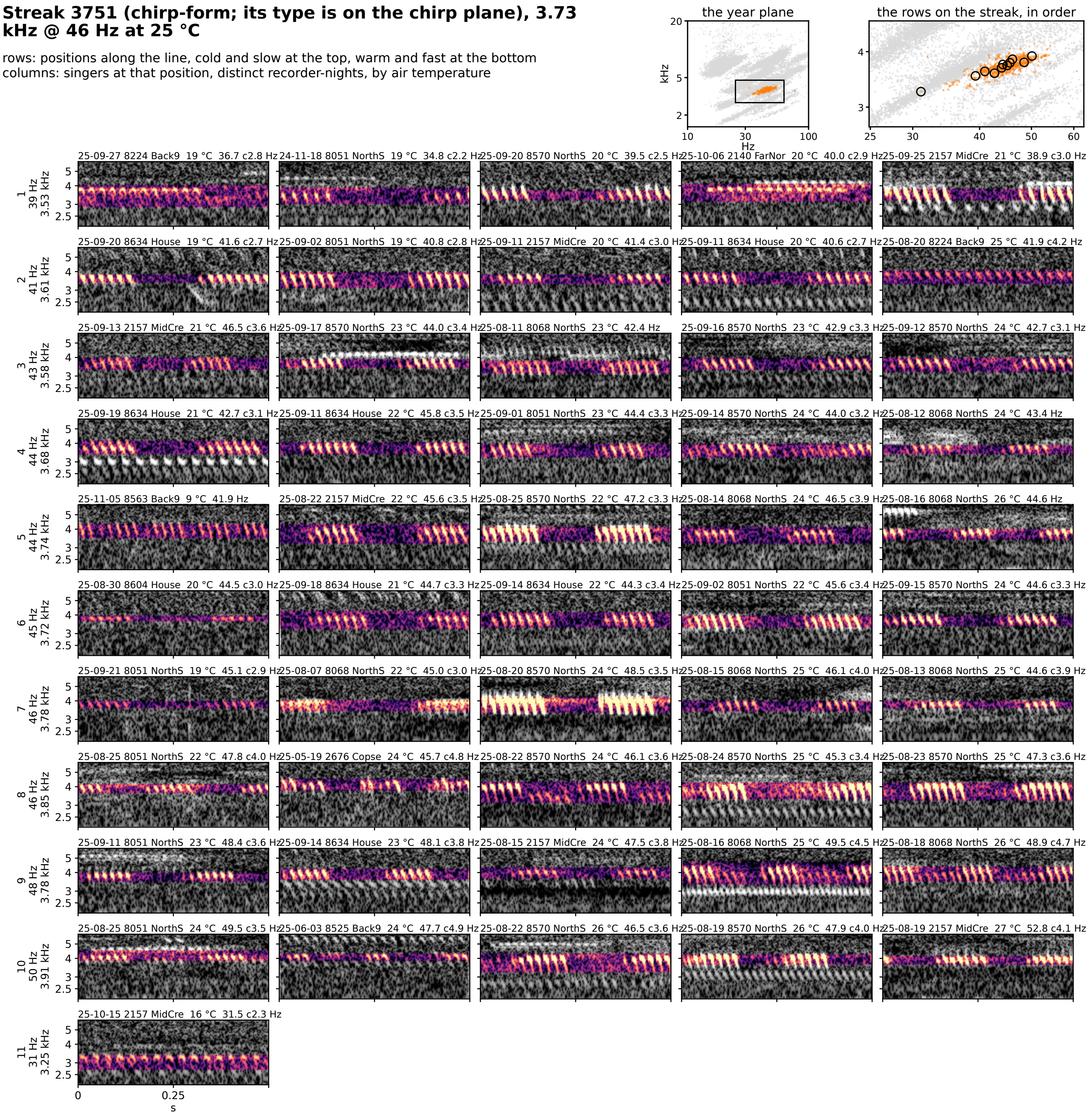


**Figure 7S-streak-3751.** Streak 3751 (chirp-form; its type is on the chirp plane): 3.73 kHz @ 46 Hz at 25 °C, 144 singer-nights. Rows are positions along the line from the cold, slow end at the top to the warm, fast end at the bottom; columns are singers at that position from distinct recorder-nights, ordered by air temperature. Each cell is 0.5 s of the loudest 30 s of the singer's deepest window on one shared log-frequency axis, the detected band in color (dB above the row's quiet level, 0 to 30) and the rest in grayscale. The cell head gives the night, recorder, site, air temperature and rate (c: chirp rate). Top right: the type on the plane, and the row positions on the streak, in order.

**Streak 5751 (chirp-form; its type is on the chirp plane), 8.10 kHz @ 85 Hz at 25 °C**

rows: positions along the line, cold and slow at the top, warm and fast at the bottom
columns: singers at that position, distinct recorder-nights, by air temperature

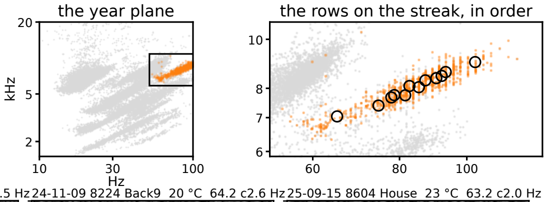


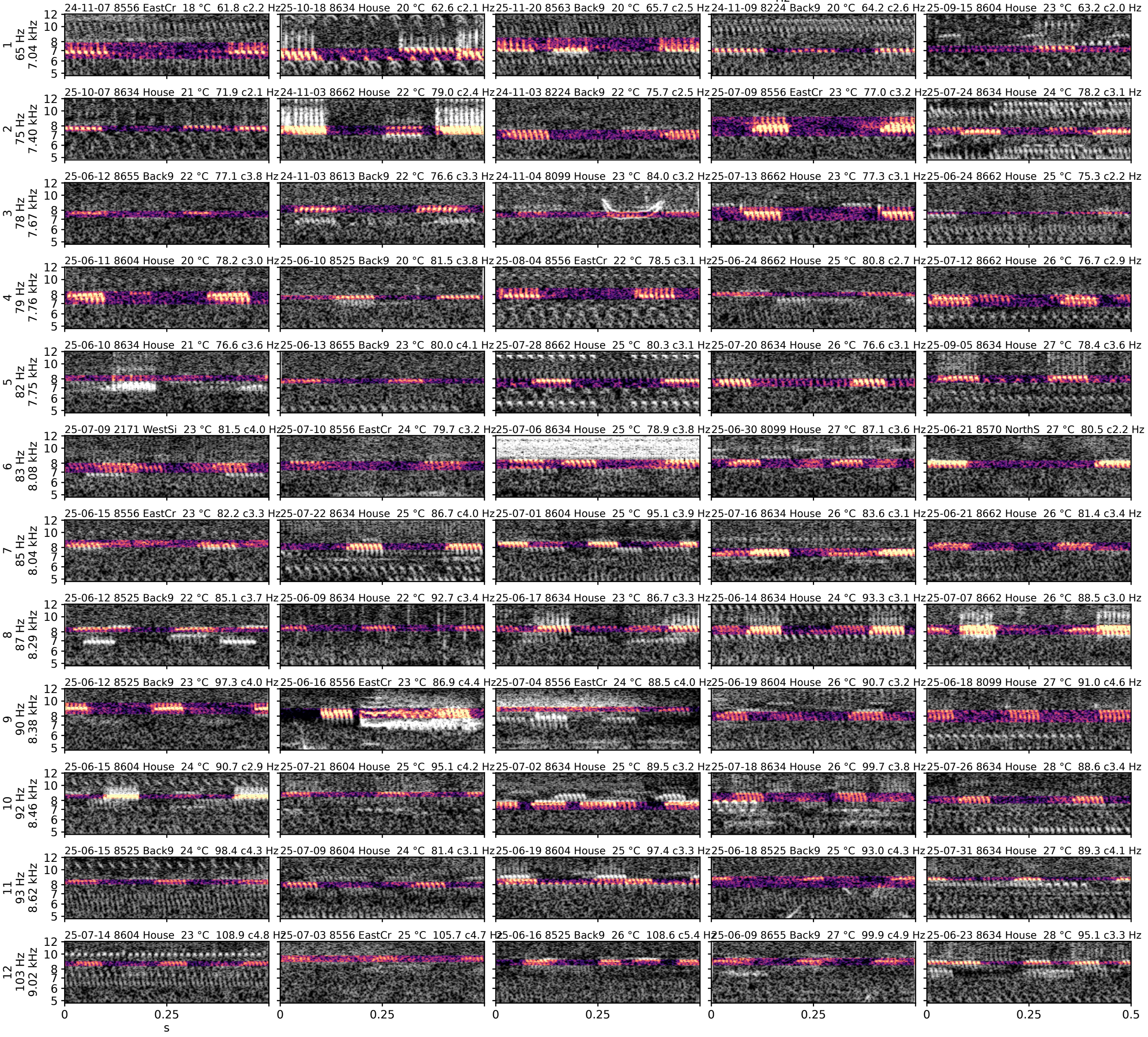


**Figure 7S-streak-5751.** Streak 5751 (chirp-form; its type is on the chirp plane): 8.10 kHz @ 85 Hz at 25 °C, 142 singer-nights. Rows are positions along the line from the cold, slow end at the top to the warm, fast end at the bottom; columns are singers at that position from distinct recorder-nights, ordered by air temperature. Each cell is 0.5 s of the loudest 30 s of the singer's deepest window on one shared log-frequency axis, the detected band in color (dB above the row's quiet level, 0 to 30) and the rest in grayscale. The cell head gives the night, recorder, site, air temperature and rate (c: chirp rate). Top right: the type on the plane, and the row positions on the streak, in order.

**Streak 5851 (chirp-form; its type is on the chirp plane), 6.54 kHz @ 39 Hz at 25 °C**

rows: positions along the line, cold and slow at the top, warm and fast at the bottom
columns: singers at that position, distinct recorder-nights, by air temperature

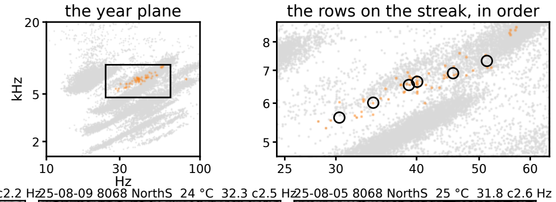


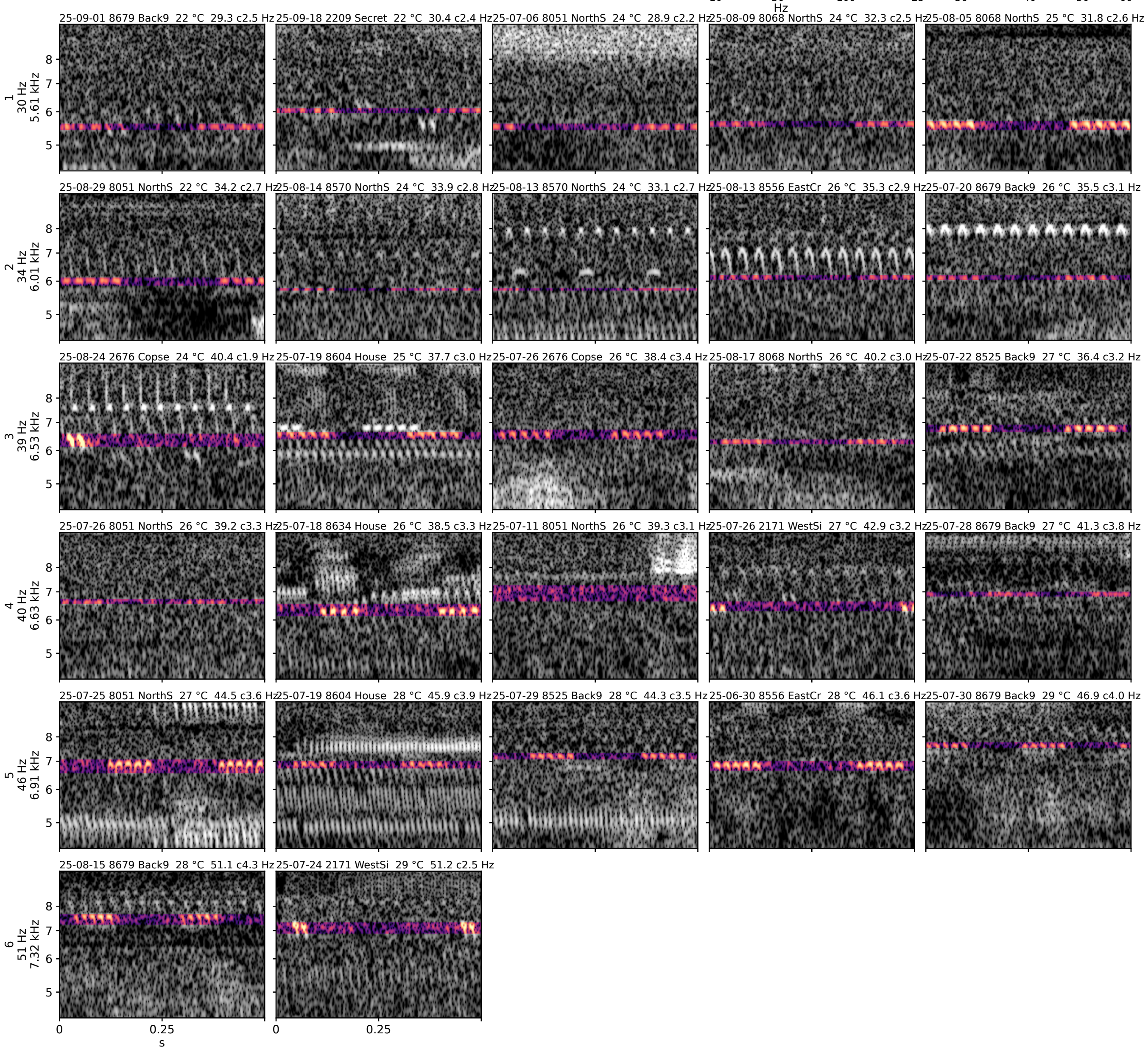


**Figure 7S-streak-5851.** Streak 5851 (chirp-form; its type is on the chirp plane): 6.54 kHz @ 39 Hz at 25 °C, 46 singer-nights. Rows are positions along the line from the cold, slow end at the top to the warm, fast end at the bottom; columns are singers at that position from distinct recorder-nights, ordered by air temperature. Each cell is 0.5 s of the loudest 30 s of the singer's deepest window on one shared log-frequency axis, the detected band in color (dB above the row's quiet level, 0 to 30) and the rest in grayscale. The cell head gives the night, recorder, site, air temperature and rate (c: chirp rate). Top right: the type on the plane, and the row positions on the streak, in order.

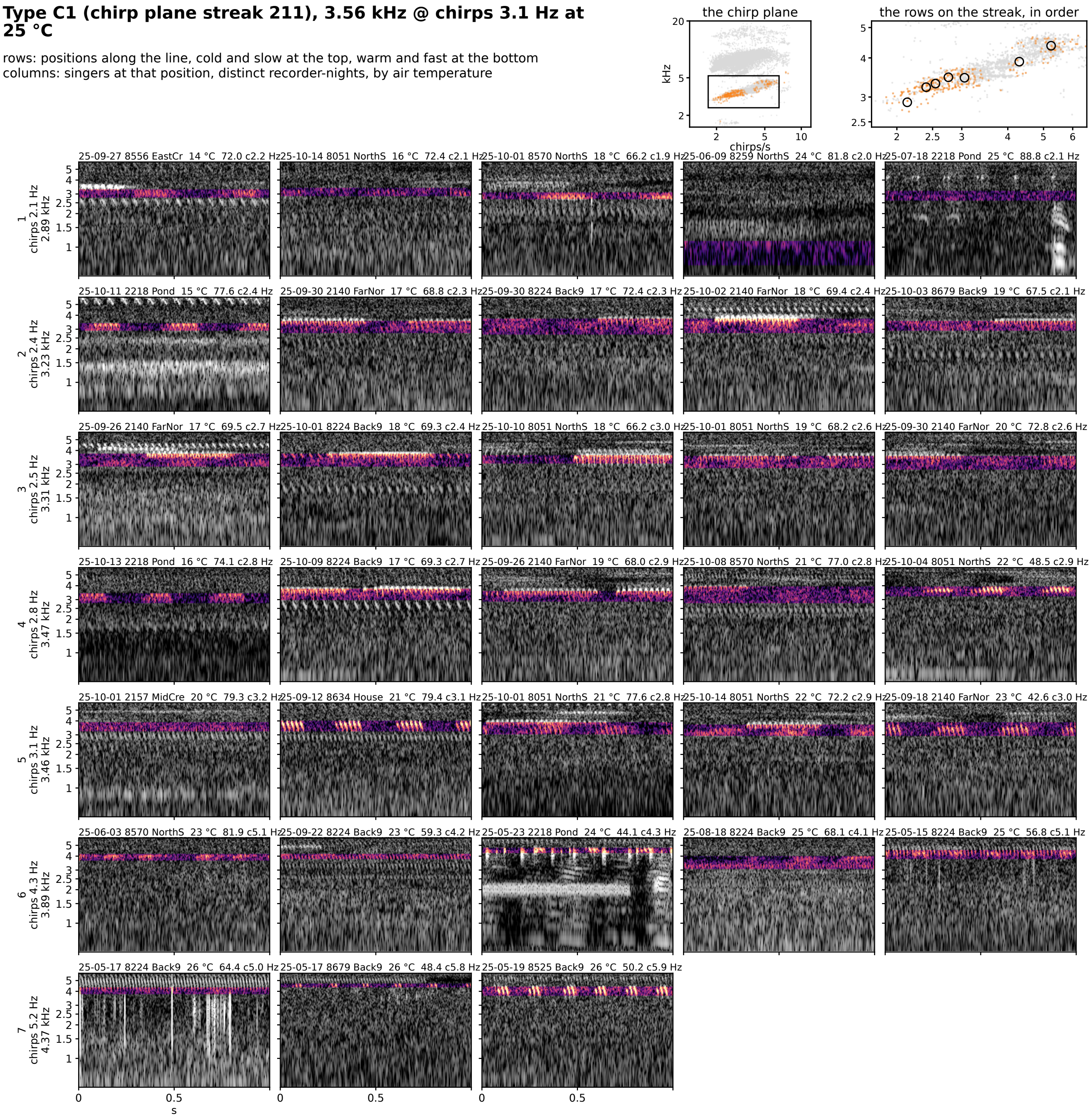


**Figure 7S-C1.** Type C1 (chirp plane streak 211): 3.56 kHz @ chirps 3.1 Hz at 25 °C, 62 singer-nights. Rows are positions along the line from the cold, slow end at the top to the warm, fast end at the bottom; columns are singers at that position from distinct recorder-nights, ordered by air temperature. Each cell is 1 s of the loudest 30 s of the singer's deepest window on one shared log-frequency axis, the detected band in color (dB above the row's quiet level, 0 to 30) and the rest in grayscale; a cell whose singer had no syllable rate in that window is folded at its chirp rate. The cell head gives the night, recorder, site, air temperature and rate (c: chirp rate). Top right: the type on the plane, and the row positions on the streak, in order.

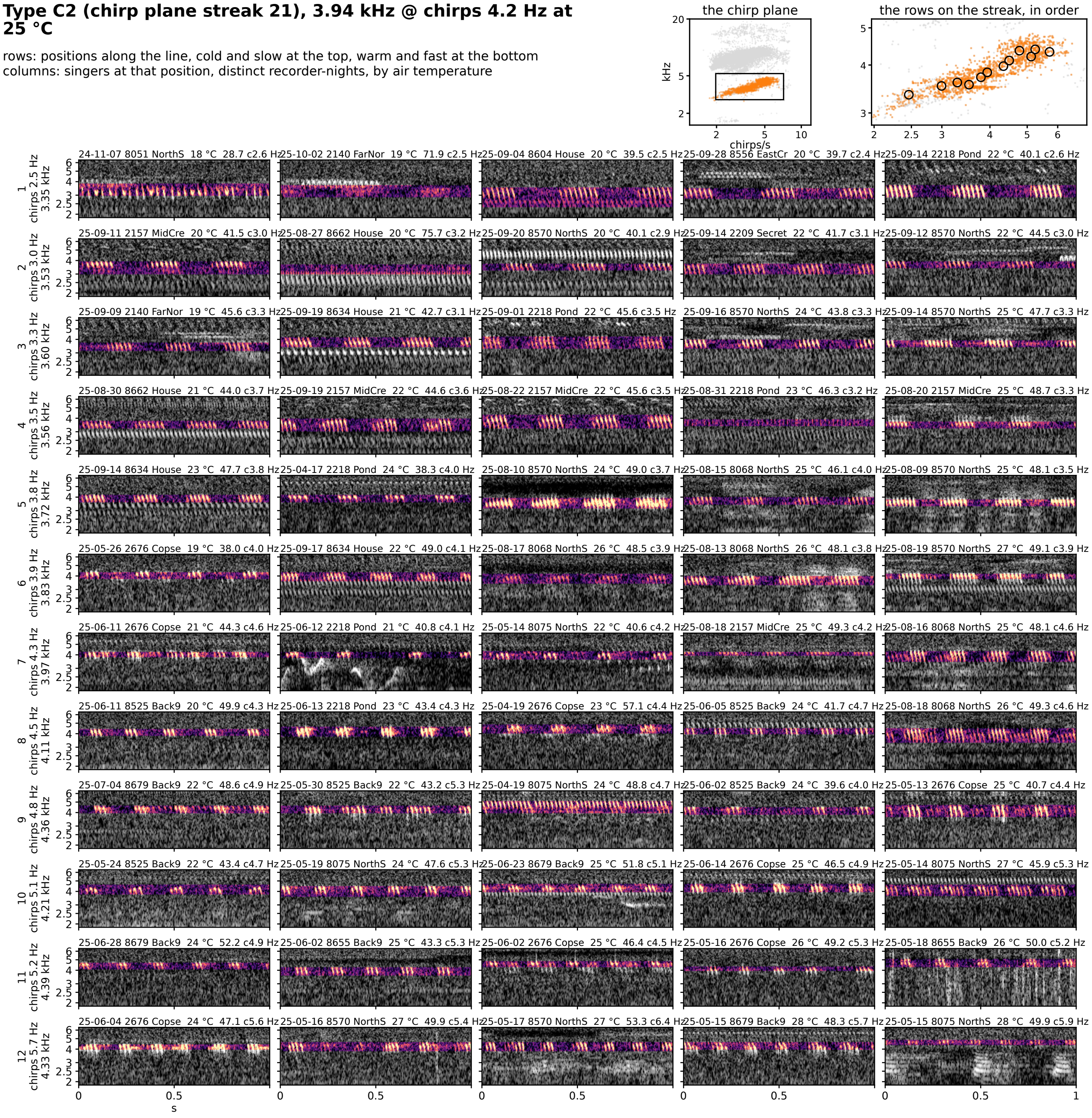


**Figure 7S-C2.** Type C2 (chirp plane streak 21): 3.94 kHz @ chirps 4.2 Hz at 25 °C, 351 singer-nights. Rows are positions along the line from the cold, slow end at the top to the warm, fast end at the bottom; columns are singers at that position from distinct recorder-nights, ordered by air temperature. Each cell is 1 s of the loudest 30 s of the singer's deepest window on one shared log-frequency axis, the detected band in color (dB above the row's quiet level, 0 to 30) and the rest in grayscale; a cell whose singer had no syllable rate in that window is folded at its chirp rate. The cell head gives the night, recorder, site, air temperature and rate (c: chirp rate). Top right: the type on the plane, and the row positions on the streak, in order.

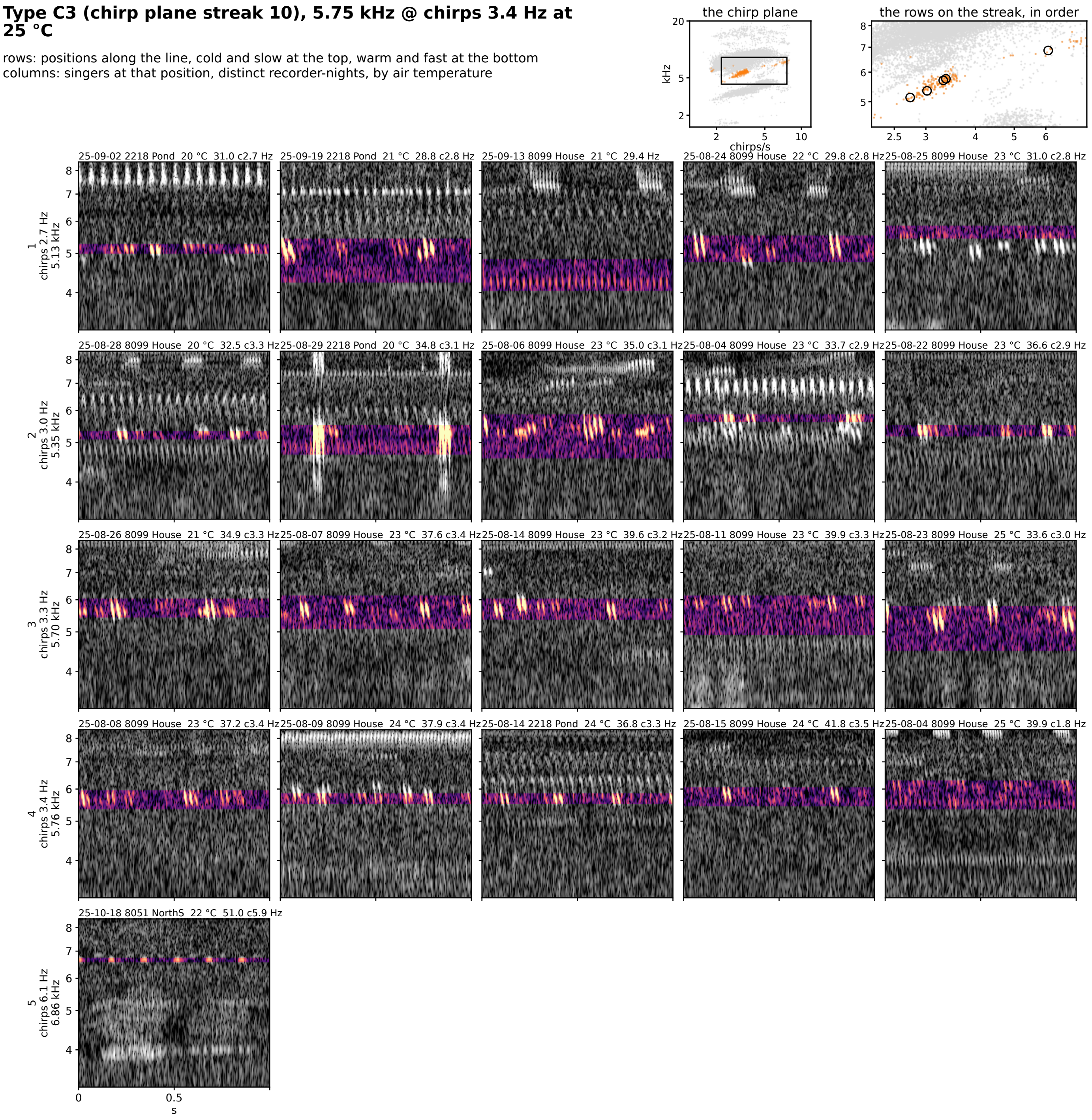


**Figure 7S-C3.** Type C3 (chirp plane streak 10): 5.75 kHz @ chirps 3.4 Hz at 25 °C, 55 singer-nights. Rows are positions along the line from the cold, slow end at the top to the warm, fast end at the bottom; columns are singers at that position from distinct recorder-nights, ordered by air temperature. Each cell is 1 s of the loudest 30 s of the singer's deepest window on one shared log-frequency axis, the detected band in color (dB above the row's quiet level, 0 to 30) and the rest in grayscale; a cell whose singer had no syllable rate in that window is folded at its chirp rate. The cell head gives the night, recorder, site, air temperature and rate (c: chirp rate). Top right: the type on the plane, and the row positions on the streak, in order.

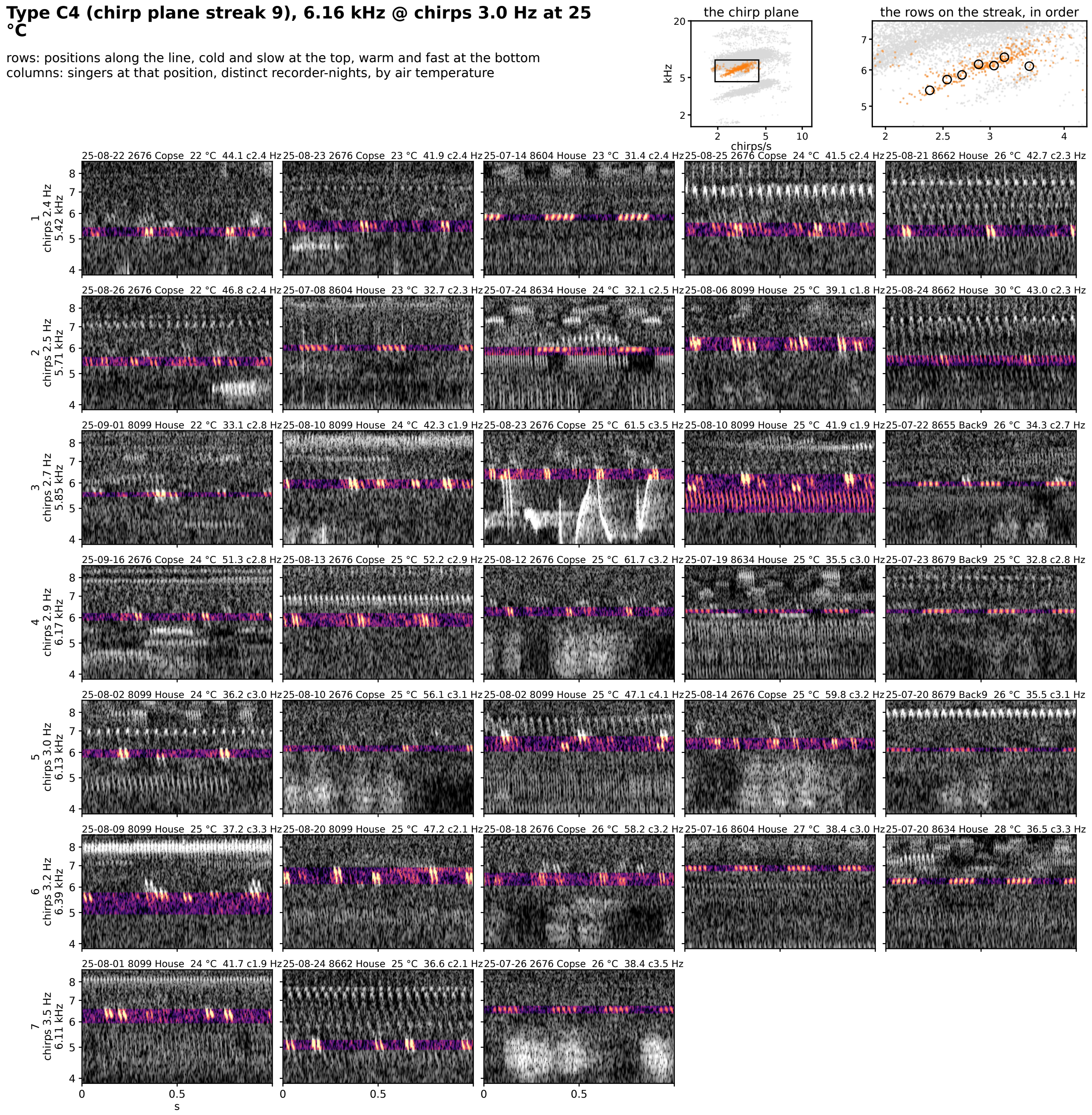


**Figure 7S-C4.** Type C4 (chirp plane streak 9): 6.16 kHz @ chirps 3.0 Hz at 25 °C, 90 singer-nights. Rows are positions along the line from the cold, slow end at the top to the warm, fast end at the bottom; columns are singers at that position from distinct recorder-nights, ordered by air temperature. Each cell is 1 s of the loudest 30 s of the singer's deepest window on one shared log-frequency axis, the detected band in color (dB above the row's quiet level, 0 to 30) and the rest in grayscale; a cell whose singer had no syllable rate in that window is folded at its chirp rate. The cell head gives the night, recorder, site, air temperature and rate (c: chirp rate). Top right: the type on the plane, and the row positions on the streak, in order.

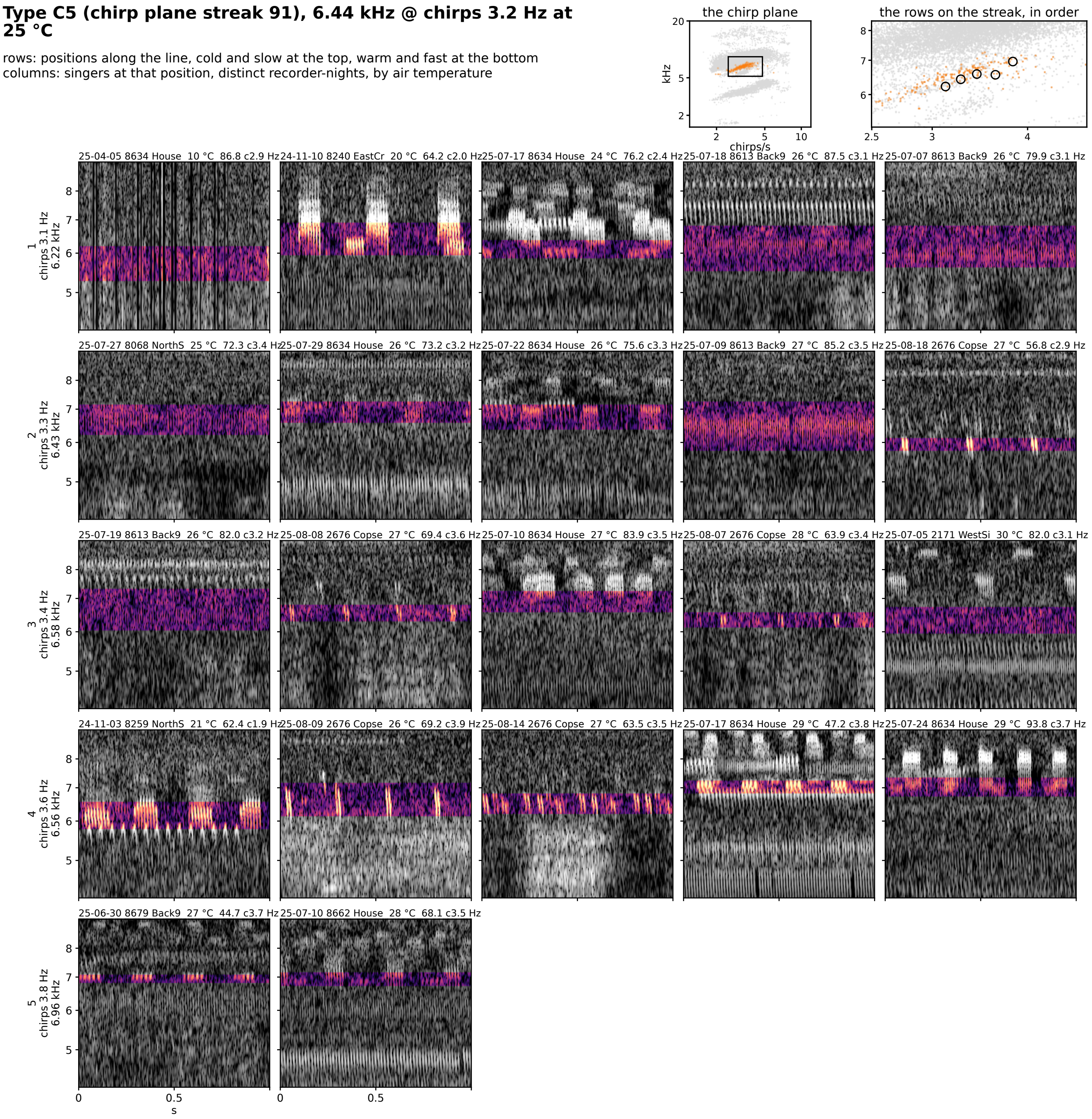


**Figure 7S-C5.** Type C5 (chirp plane streak 91): 6.44 kHz @ chirps 3.2 Hz at 25 °C, 48 singer-nights. Rows are positions along the line from the cold, slow end at the top to the warm, fast end at the bottom; columns are singers at that position from distinct recorder-nights, ordered by air temperature. Each cell is 1 s of the loudest 30 s of the singer's deepest window on one shared log-frequency axis, the detected band in color (dB above the row's quiet level, 0 to 30) and the rest in grayscale; a cell whose singer had no syllable rate in that window is folded at its chirp rate. The cell head gives the night, recorder, site, air temperature and rate (c: chirp rate). Top right: the type on the plane, and the row positions on the streak, in order.

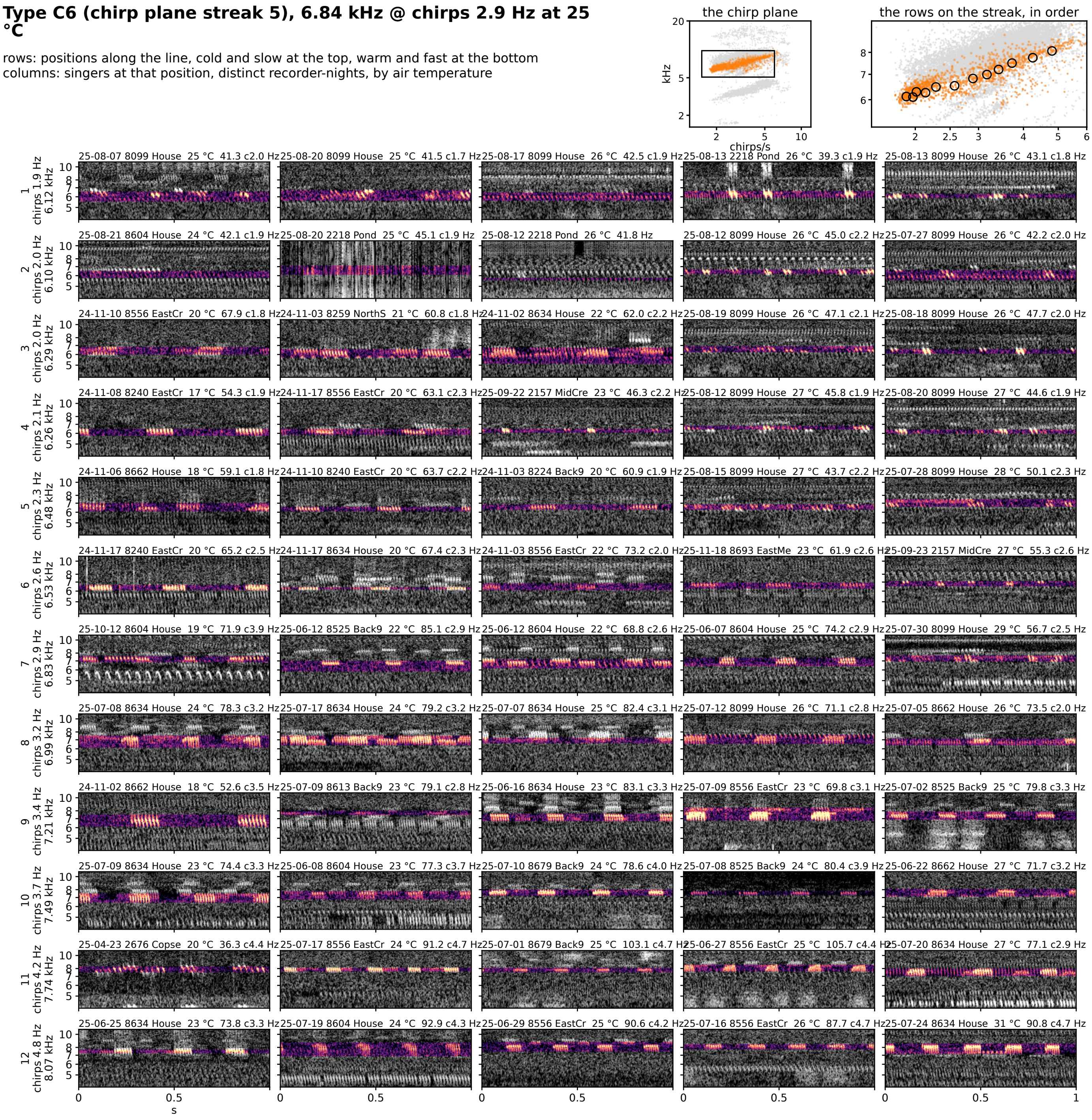


**Figure 7S-C6.** Type C6 (chirp plane streak 5): 6.84 kHz @ chirps 2.9 Hz at 25 °C, 365 singer-nights. Rows are positions along the line from the cold, slow end at the top to the warm, fast end at the bottom; columns are singers at that position from distinct recorder-nights, ordered by air temperature. Each cell is 1 s of the loudest 30 s of the singer's deepest window on one shared log-frequency axis, the detected band in color (dB above the row's quiet level, 0 to 30) and the rest in grayscale; a cell whose singer had no syllable rate in that window is folded at its chirp rate. The cell head gives the night, recorder, site, air temperature and rate (c: chirp rate). Top right: the type on the plane, and the row positions on the streak, in order.

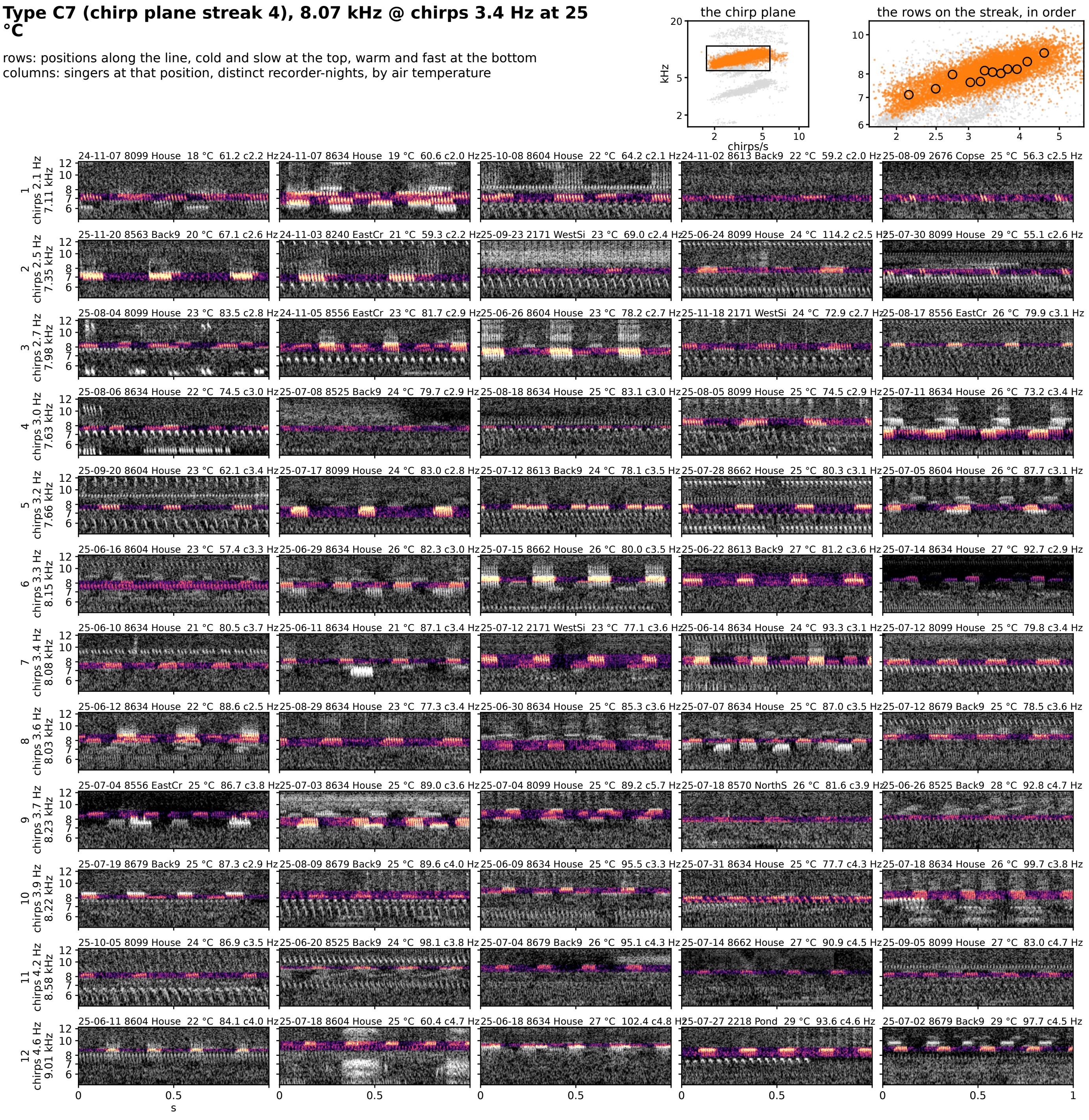


**Figure 7S-C7.** Type C7 (chirp plane streak 4): 8.07 kHz @ chirps 3.4 Hz at 25 °C, 2659 singer-nights. Rows are positions along the line from the cold, slow end at the top to the warm, fast end at the bottom; columns are singers at that position from distinct recorder-nights, ordered by air temperature. Each cell is 1 s of the loudest 30 s of the singer's deepest window on one shared log-frequency axis, the detected band in color (dB above the row's quiet level, 0 to 30) and the rest in grayscale; a cell whose singer had no syllable rate in that window is folded at its chirp rate. The cell head gives the night, recorder, site, air temperature and rate (c: chirp rate). Top right: the type on the plane, and the row positions on the streak, in order.

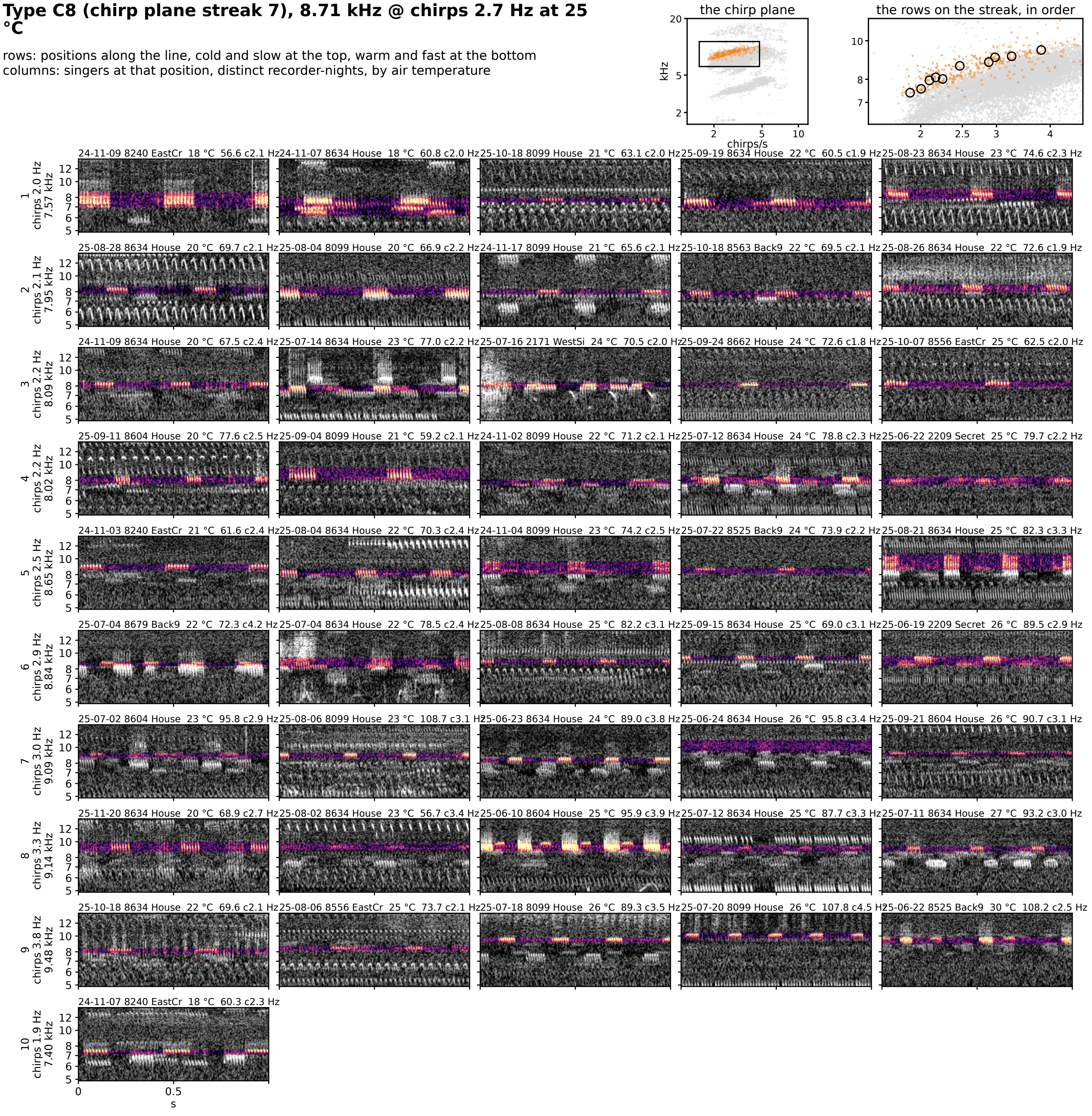


**Figure 7S-C8.** Type C8 (chirp plane streak 7): 8.71 kHz @ chirps 2.7 Hz at 25 °C, 101 singer-nights. Rows are positions along the line from the cold, slow end at the top to the warm, fast end at the bottom; columns are singers at that position from distinct recorder-nights, ordered by air temperature. Each cell is 1 s of the loudest 30 s of the singer's deepest window on one shared log-frequency axis, the detected band in color (dB above the row's quiet level, 0 to 30) and the rest in grayscale; a cell whose singer had no syllable rate in that window is folded at its chirp rate. The cell head gives the night, recorder, site, air temperature and rate (c: chirp rate). Top right: the type on the plane, and the row positions on the streak, in order.

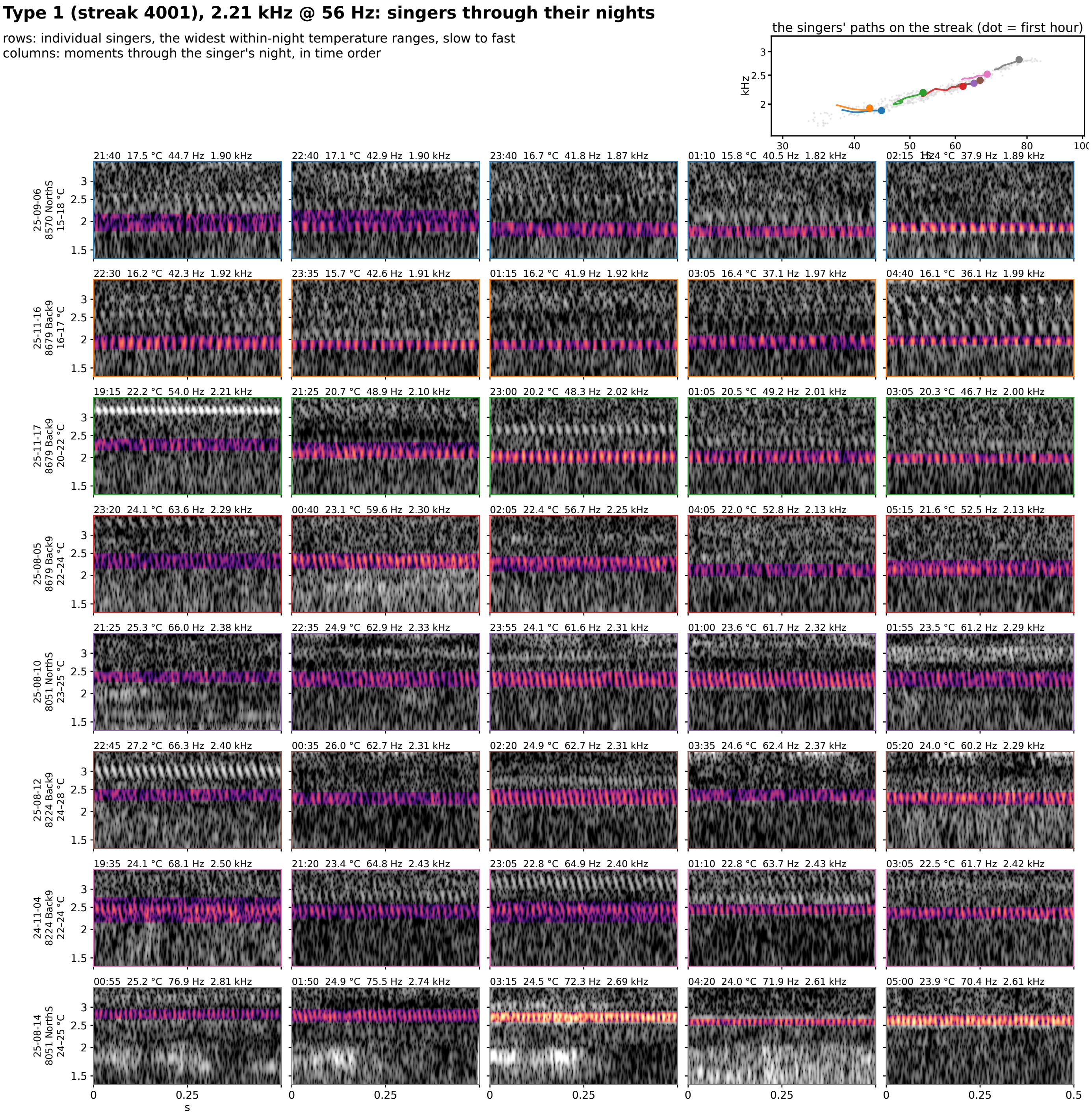


**Figure S8-1.** Type 1 (streak 4001): 2.21 kHz @ 56 Hz at 25 °C, eight singers through their nights. Rows are individual singers of the type, the eight with the widest temperature range within one night (top to bottom by the singer's median rate); columns are moments through that singer's night in time order, 5 windows at even steps through its span. Each cell is 0.5 s of the loudest 30 s of that window on one shared log-frequency axis, the detected band in color (dB above the row's quiet level, 0 to 30) and the rest in grayscale. The cell head gives the local time, the air temperature, the syllable rate and the carrier. Top right: each singer's hourly path on the plane, in the row's color, over the year's points in gray.

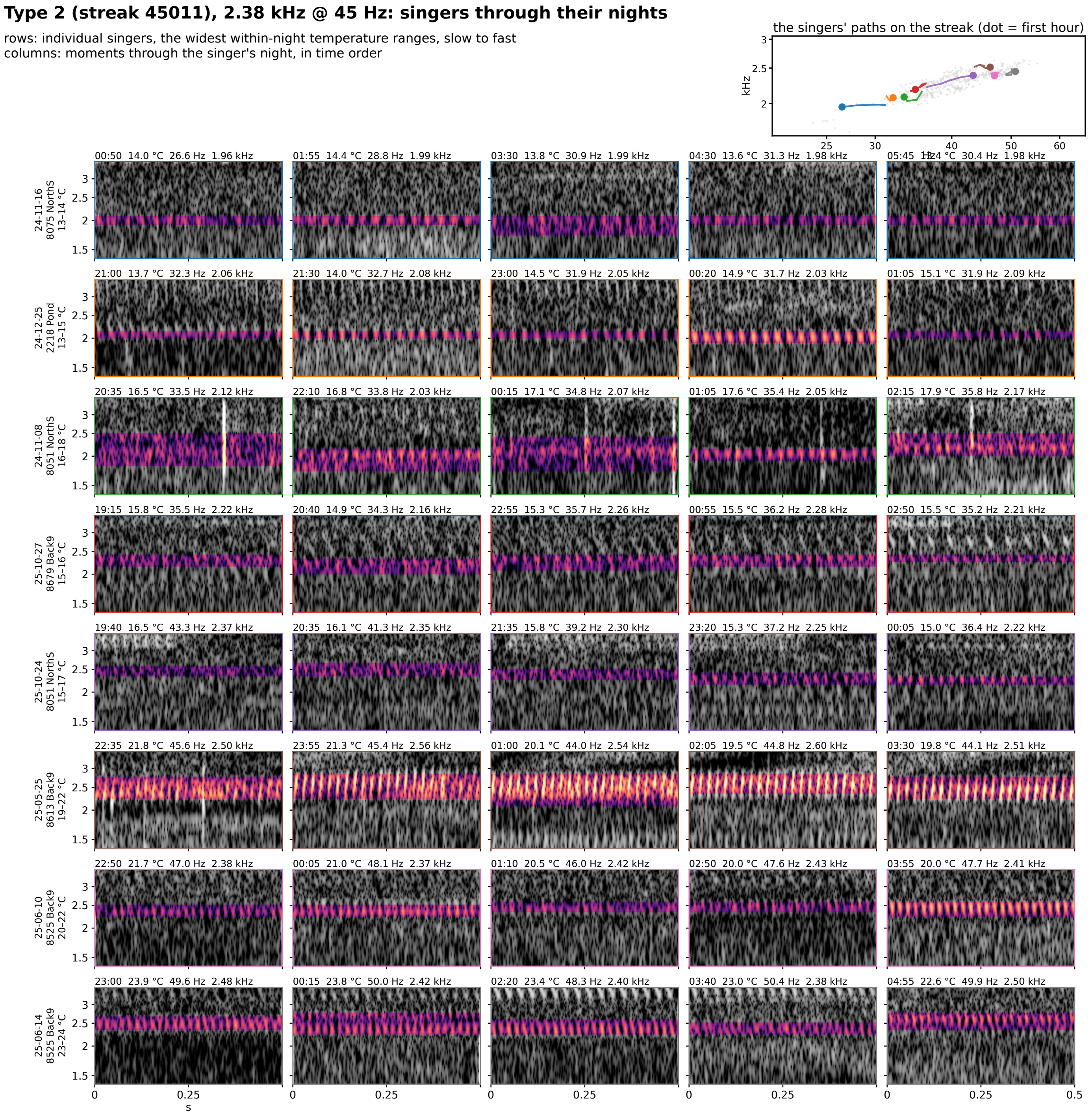


**Figure S8-2.** Type 2 (streak 45011): 2.38 kHz @ 45 Hz at 25 °C, eight singers through their nights. Rows are individual singers of the type, the eight with the widest temperature range within one night (top to bottom by the singer's median rate); columns are moments through that singer's night in time order, 5 windows at even steps through its span. Each cell is 0.5 s of the loudest 30 s of that window on one shared log-frequency axis, the detected band in color (dB above the row's quiet level, 0 to 30) and the rest in grayscale. The cell head gives the local time, the air temperature, the syllable rate and the carrier. Top right: each singer's hourly path on the plane, in the row's color, over the year's points in gray.

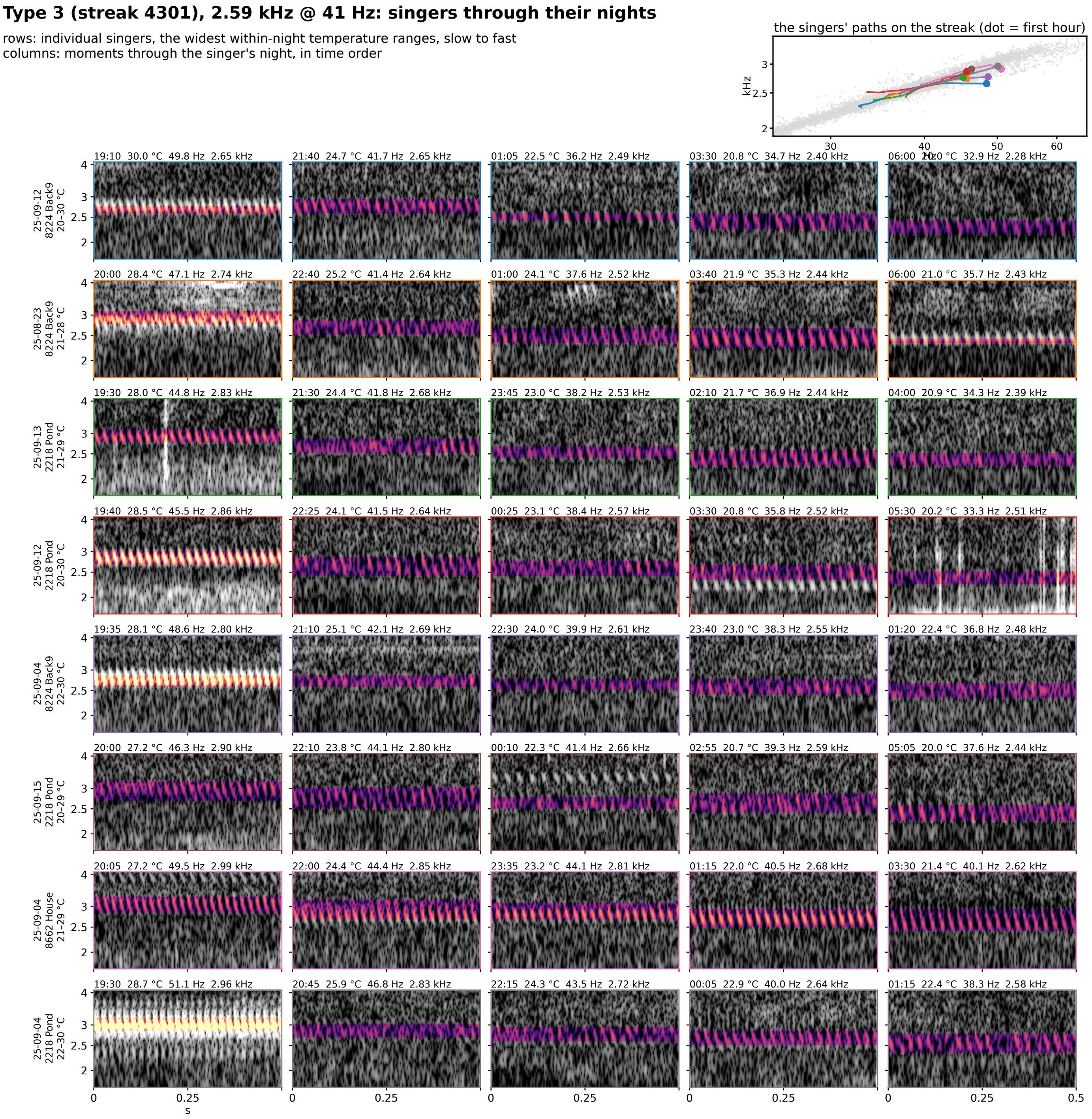


**Figure S8-3.** Type 3 (streak 4301): 2.59 kHz @ 41 Hz at 25 °C, eight singers through their nights. Rows are individual singers of the type, the eight with the widest temperature range within one night (top to bottom by the singer's median rate); columns are moments through that singer's night in time order, 5 windows at even steps through its span. Each cell is 0.5 s of the loudest 30 s of that window on one shared log-frequency axis, the detected band in color (dB above the row's quiet level, 0 to 30) and the rest in grayscale. The cell head gives the local time, the air temperature, the syllable rate and the carrier. Top right: each singer's hourly path on the plane, in the row's color, over the year's points in gray.

**Type 4 (streak 360123), 2.74 kHz @ 22 Hz: singers through their nights**

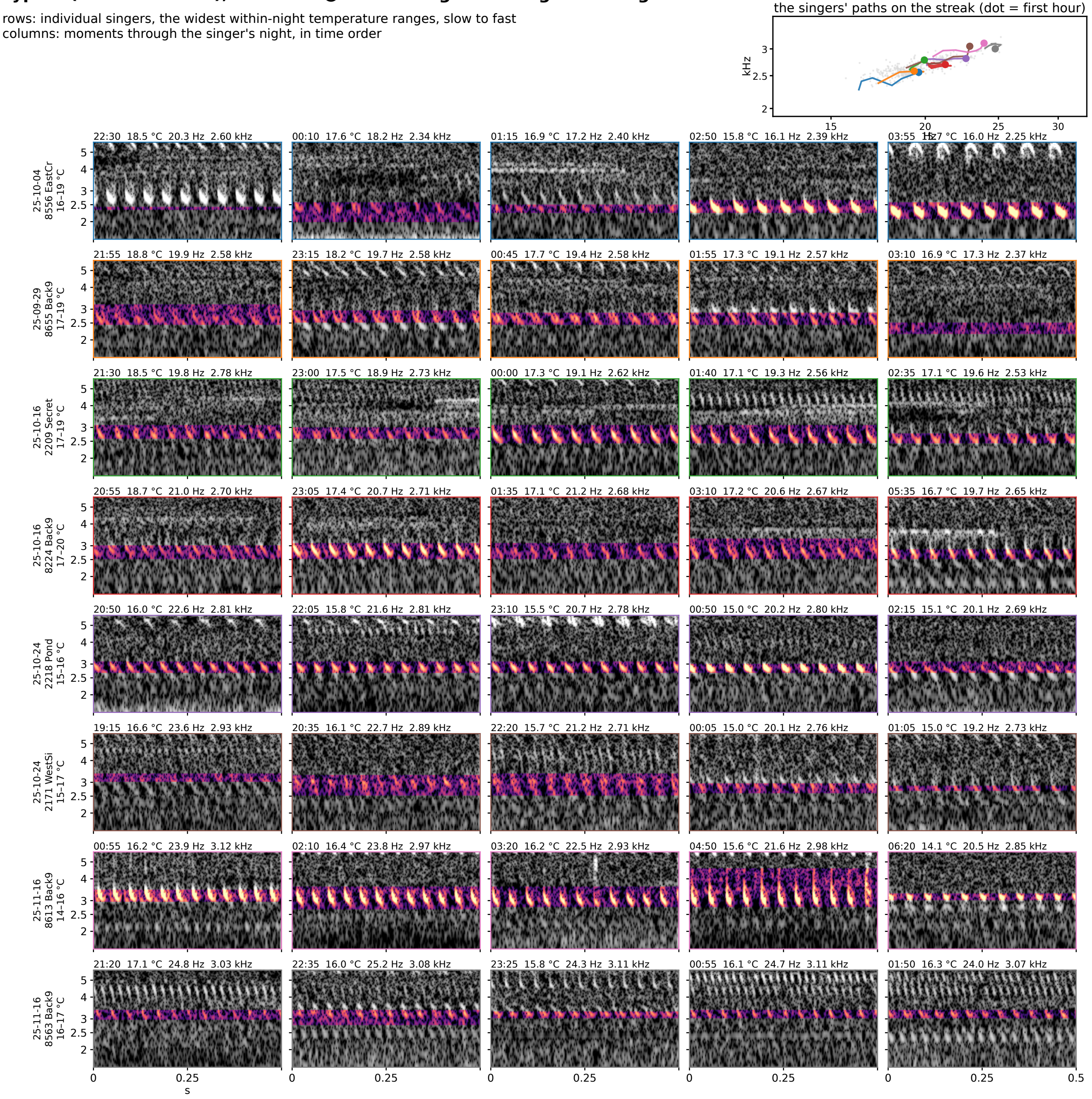


**Figure S8-4.** Type 4 (streak 360123): 2.74 kHz @ 22 Hz at 25 °C, eight singers through their nights. Rows are individual singers of the type, the eight with the widest temperature range within one night (top to bottom by the singer's median rate); columns are moments through that singer's night in time order, 5 windows at even steps through its span. Each cell is 0.5 s of the loudest 30 s of that window on one shared log-frequency axis, the detected band in color (dB above the row's quiet level, 0 to 30) and the rest in grayscale. The cell head gives the local time, the air temperature, the syllable rate and the carrier. Top right: each singer's hourly path on the plane, in the row's color, over the year's points in gray.

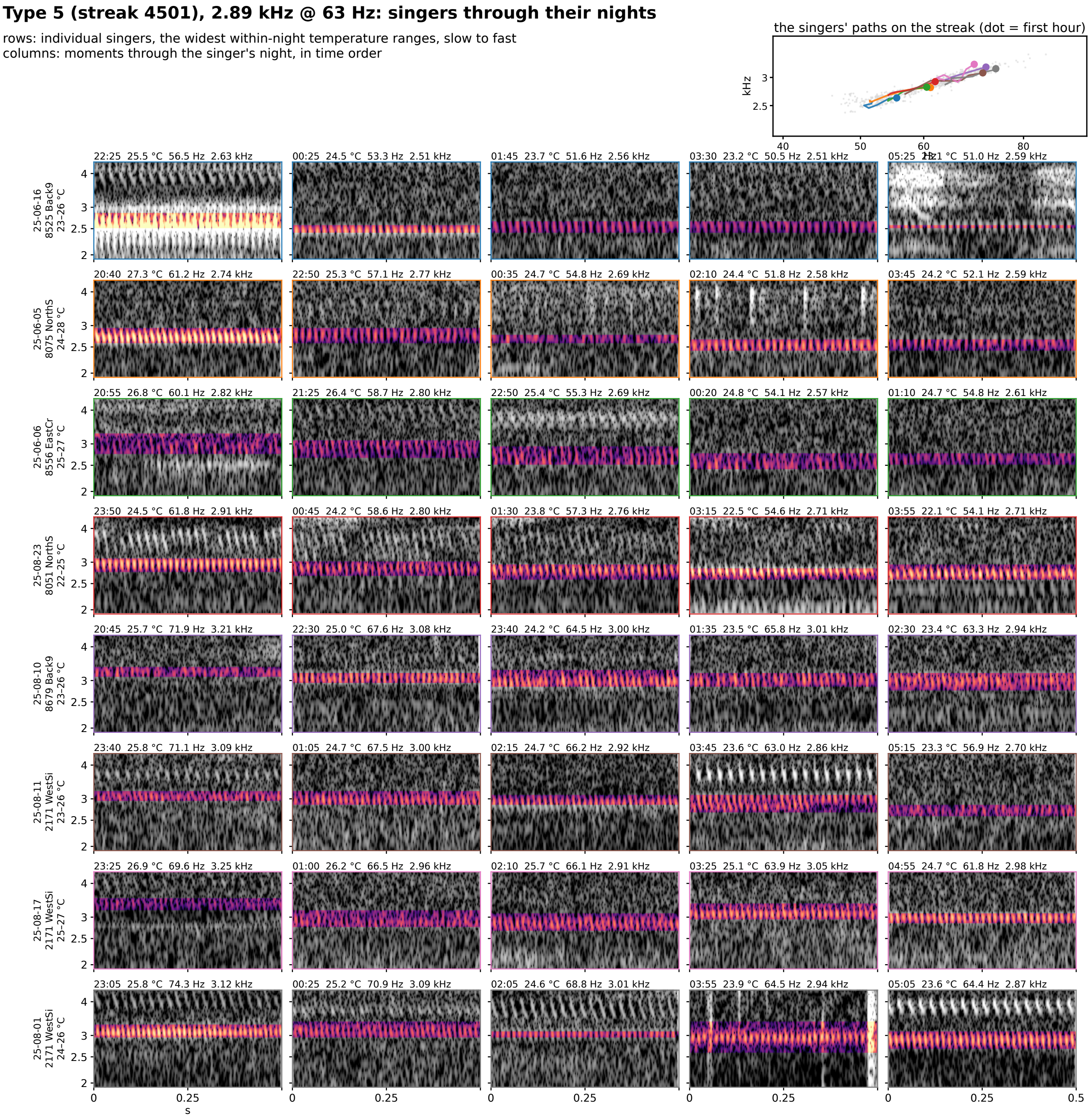


**Figure S8-5.** Type 5 (streak 4501): 2.89 kHz @ 63 Hz at 25 °C, eight singers through their nights. Rows are individual singers of the type, the eight with the widest temperature range within one night (top to bottom by the singer's median rate); columns are moments through that singer's night in time order, 5 windows at even steps through its span. Each cell is 0.5 s of the loudest 30 s of that window on one shared log-frequency axis, the detected band in color (dB above the row's quiet level, 0 to 30) and the rest in grayscale. The cell head gives the local time, the air temperature, the syllable rate and the carrier. Top right: each singer's hourly path on the plane, in the row's color, over the year's points in gray.

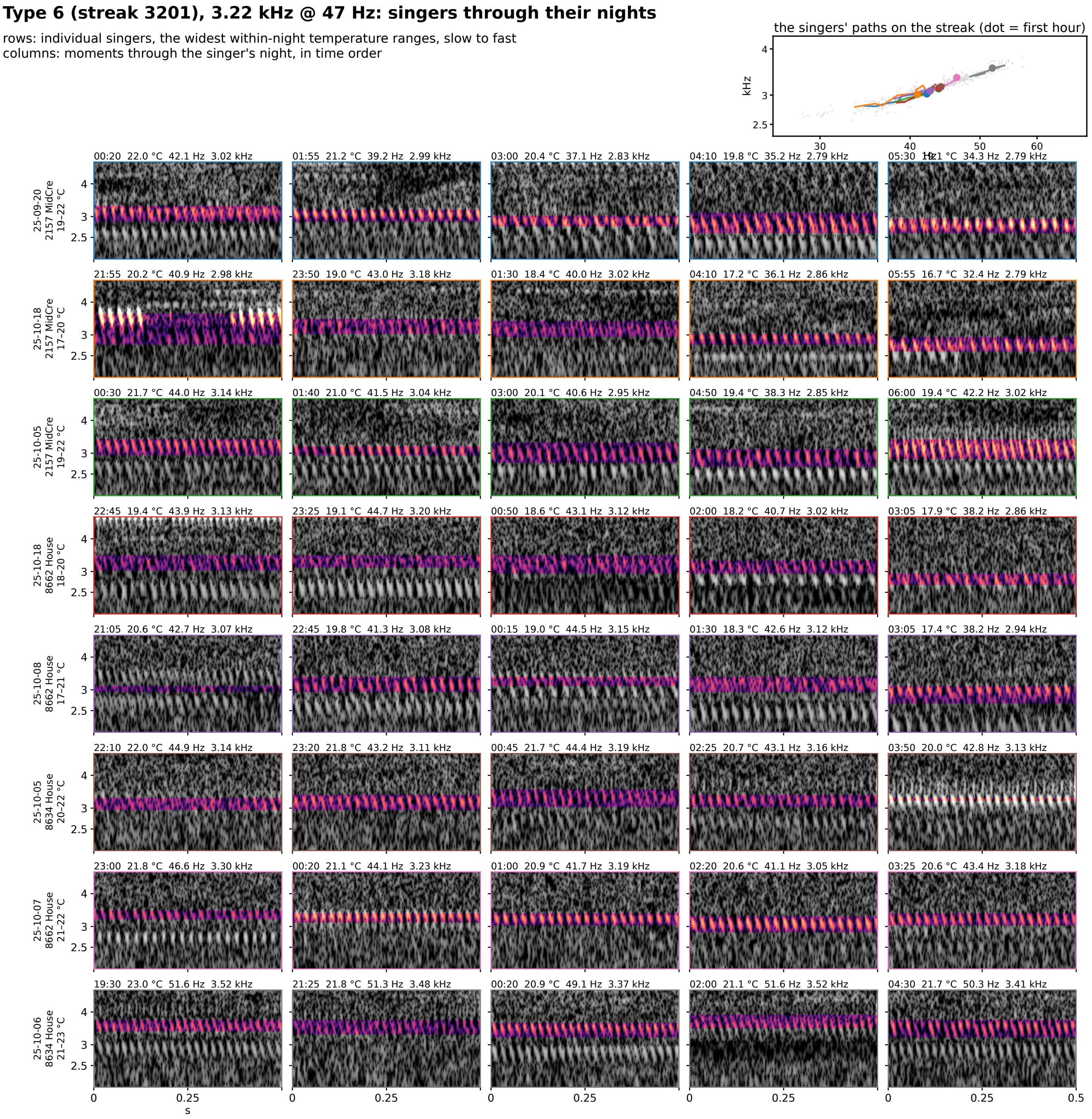


**Figure S8-6.** Type 6 (streak 3201): 3.22 kHz @ 47 Hz at 25 °C, eight singers through their nights. Rows are individual singers of the type, the eight with the widest temperature range within one night (top to bottom by the singer's median rate); columns are moments through that singer's night in time order, 5 windows at even steps through its span. Each cell is 0.5 s of the loudest 30 s of that window on one shared log-frequency axis, the detected band in color (dB above the row's quiet level, 0 to 30) and the rest in grayscale. The cell head gives the local time, the air temperature, the syllable rate and the carrier. Top right: each singer's hourly path on the plane, in the row's color, over the year's points in gray.

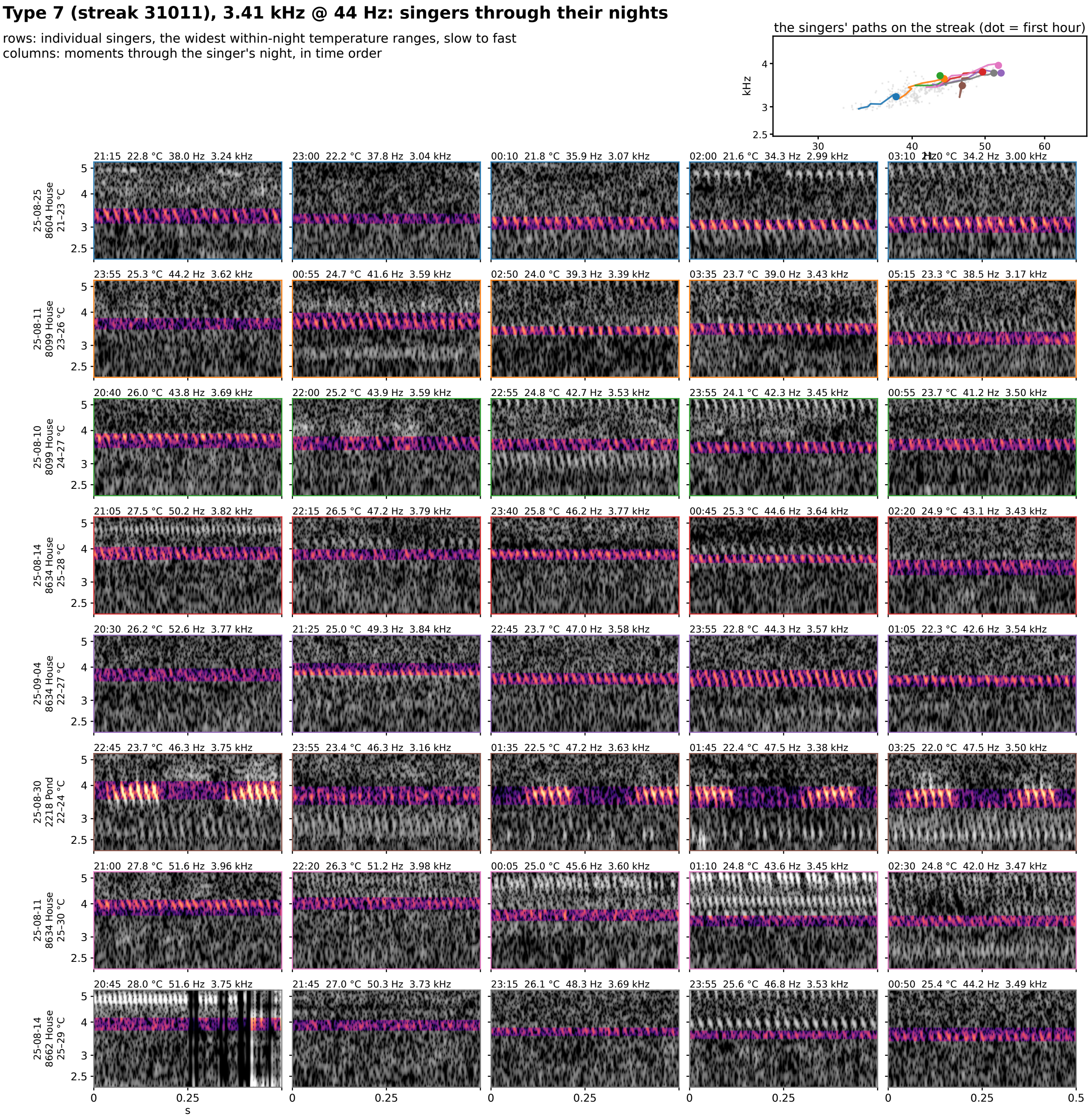


**Figure S8-7.** Type 7 (streak 31011): 3.41 kHz @ 44 Hz at 25 °C, eight singers through their nights. Rows are individual singers of the type, the eight with the widest temperature range within one night (top to bottom by the singer's median rate); columns are moments through that singer's night in time order, 5 windows at even steps through its span. Each cell is 0.5 s of the loudest 30 s of that window on one shared log-frequency axis, the detected band in color (dB above the row's quiet level, 0 to 30) and the rest in grayscale. The cell head gives the local time, the air temperature, the syllable rate and the carrier. Top right: each singer's hourly path on the plane, in the row's color, over the year's points in gray.

### Type 8 (streak 3601), 3.48 kHz @ 35 Hz: singers through their nights

rows: individual singers, the widest within-night temperature ranges, slow to fast
columns: moments through the singer's night, in time order

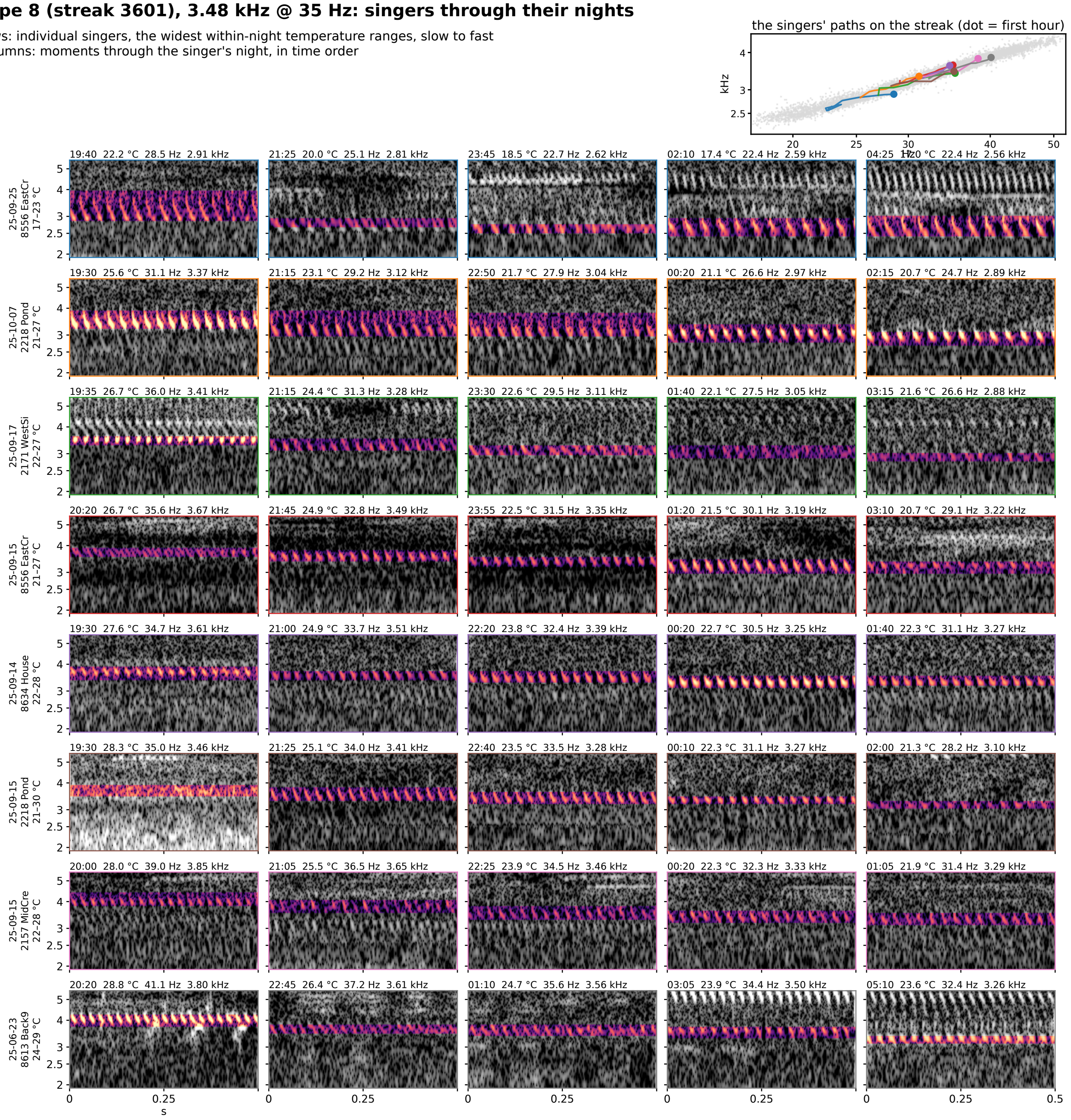

**Figure S8-8.** Type 8 (streak 3601): 3.48 kHz @ 35 Hz at 25 °C, eight singers through their nights. Rows are individual singers of the type, the eight with the widest temperature range within one night (top to bottom by the singer's median rate); columns are moments through that singer's night in time order, 5 windows at even steps through its span. Each cell is 0.5 s of the loudest 30 s of that window on one shared log-frequency axis, the detected band in color (dB above the row's quiet level, 0 to 30) and the rest in grayscale. The cell head gives the local time, the air temperature, the syllable rate and the carrier. Top right: each singer's hourly path on the plane, in the row's color, over the year's points in gray.

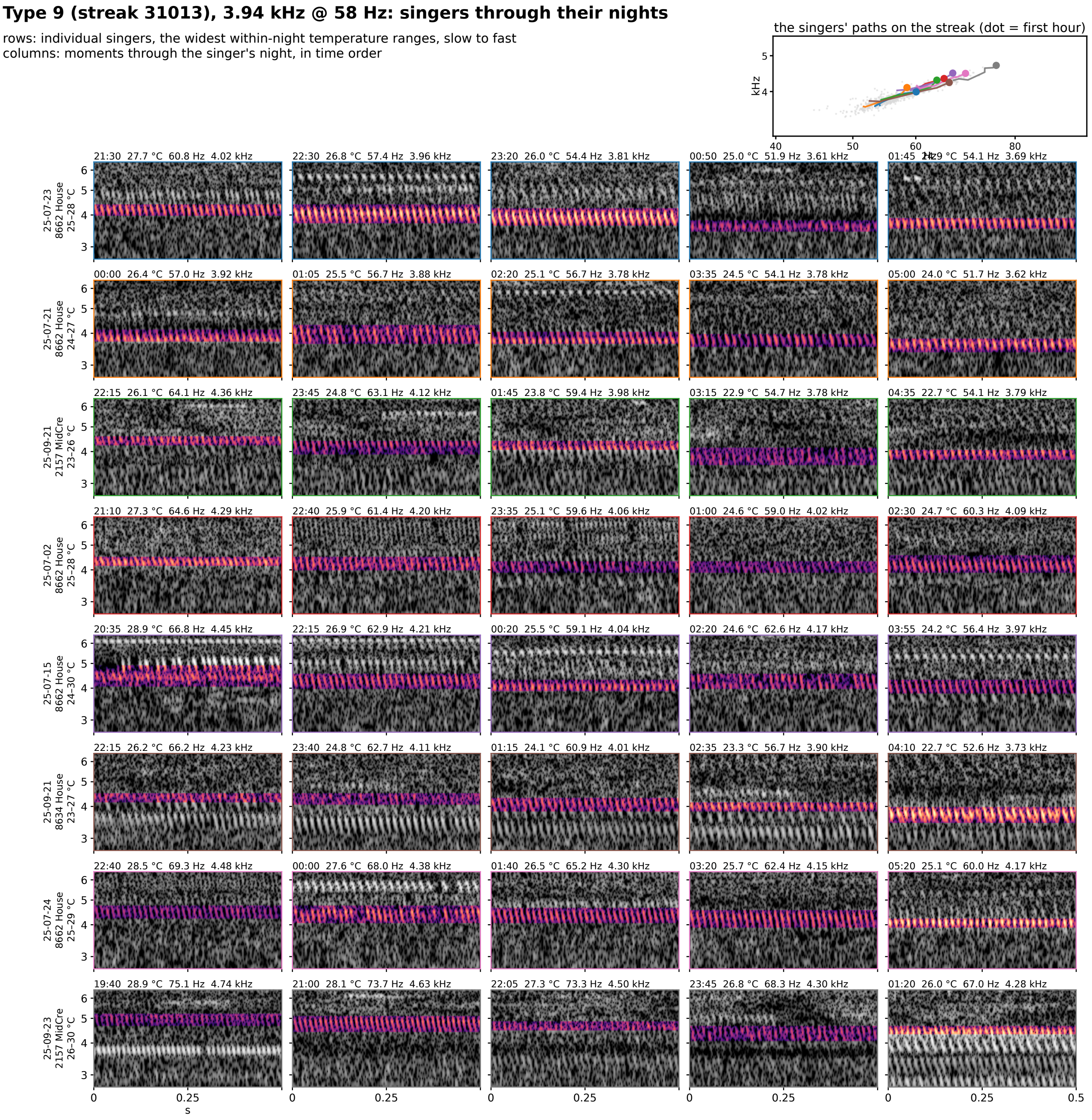


**Figure S8-9.** Type 9 (streak 31013): 3.94 kHz @ 58 Hz at 25 °C, eight singers through their nights. Rows are individual singers of the type, the eight with the widest temperature range within one night (top to bottom by the singer's median rate); columns are moments through that singer's night in time order, 5 windows at even steps through its span. Each cell is 0.5 s of the loudest 30 s of that window on one shared log-frequency axis, the detected band in color (dB above the row's quiet level, 0 to 30) and the rest in grayscale. The cell head gives the local time, the air temperature, the syllable rate and the carrier. Top right: each singer's hourly path on the plane, in the row's color, over the year's points in gray.

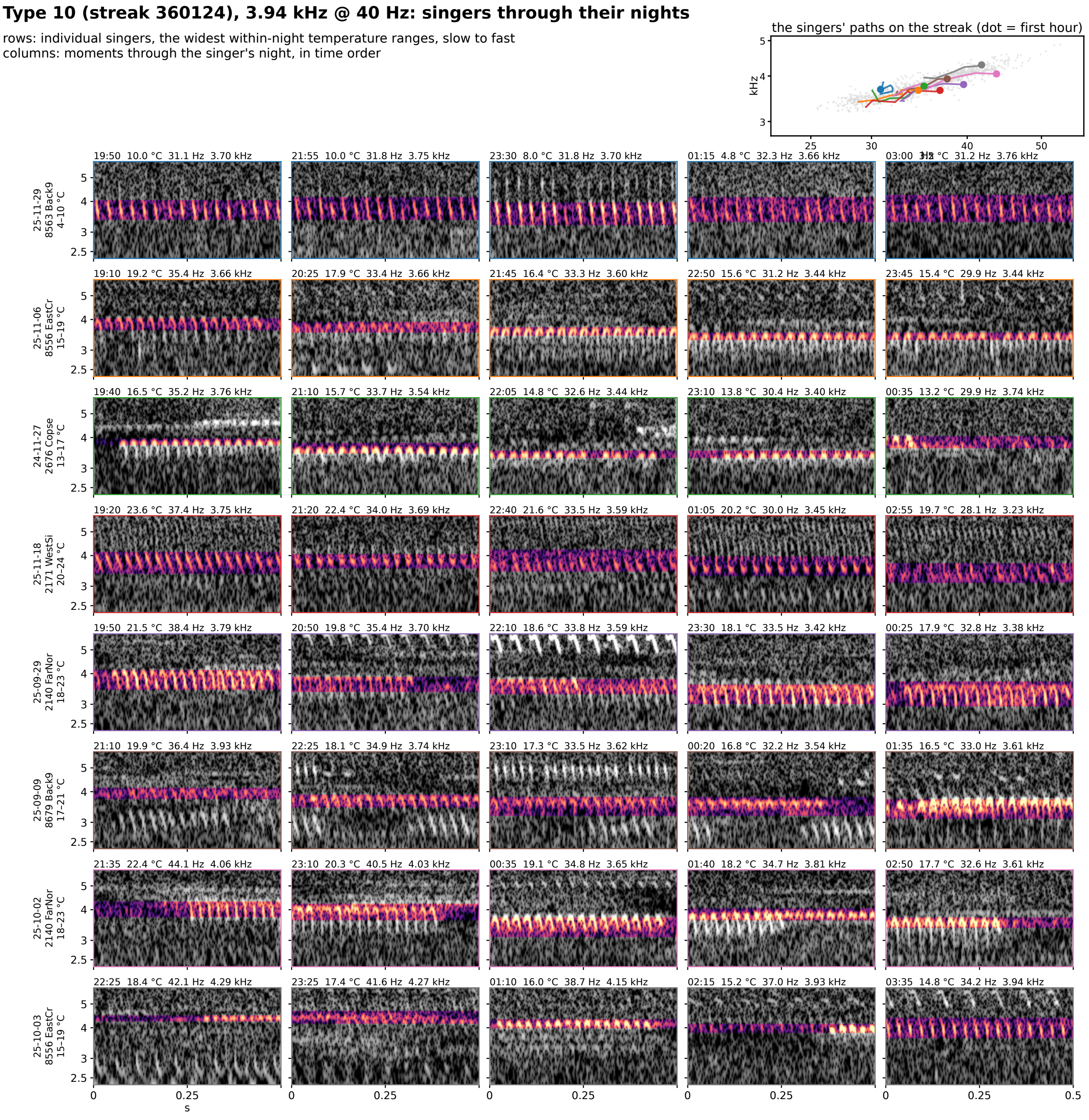


**Figure S8-10.** Type 10 (streak 360124): 3.94 kHz @ 40 Hz at 25 °C, eight singers through their nights. Rows are individual singers of the type, the eight with the widest temperature range within one night (top to bottom by the singer's median rate); columns are moments through that singer's night in time order, 5 windows at even steps through its span. Each cell is 0.5 s of the loudest 30 s of that window on one shared log-frequency axis, the detected band in color (dB above the row's quiet level, 0 to 30) and the rest in grayscale. The cell head gives the local time, the air temperature, the syllable rate and the carrier. Top right: each singer's hourly path on the plane, in the row's color, over the year's points in gray.

### Type 11 (streak 360125), 3.95 kHz @ 39 Hz: singers through their nights

rows: individual singers, the widest within-night temperature ranges, slow to fast
columns: moments through the singer's night, in time order

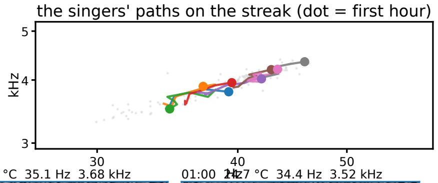


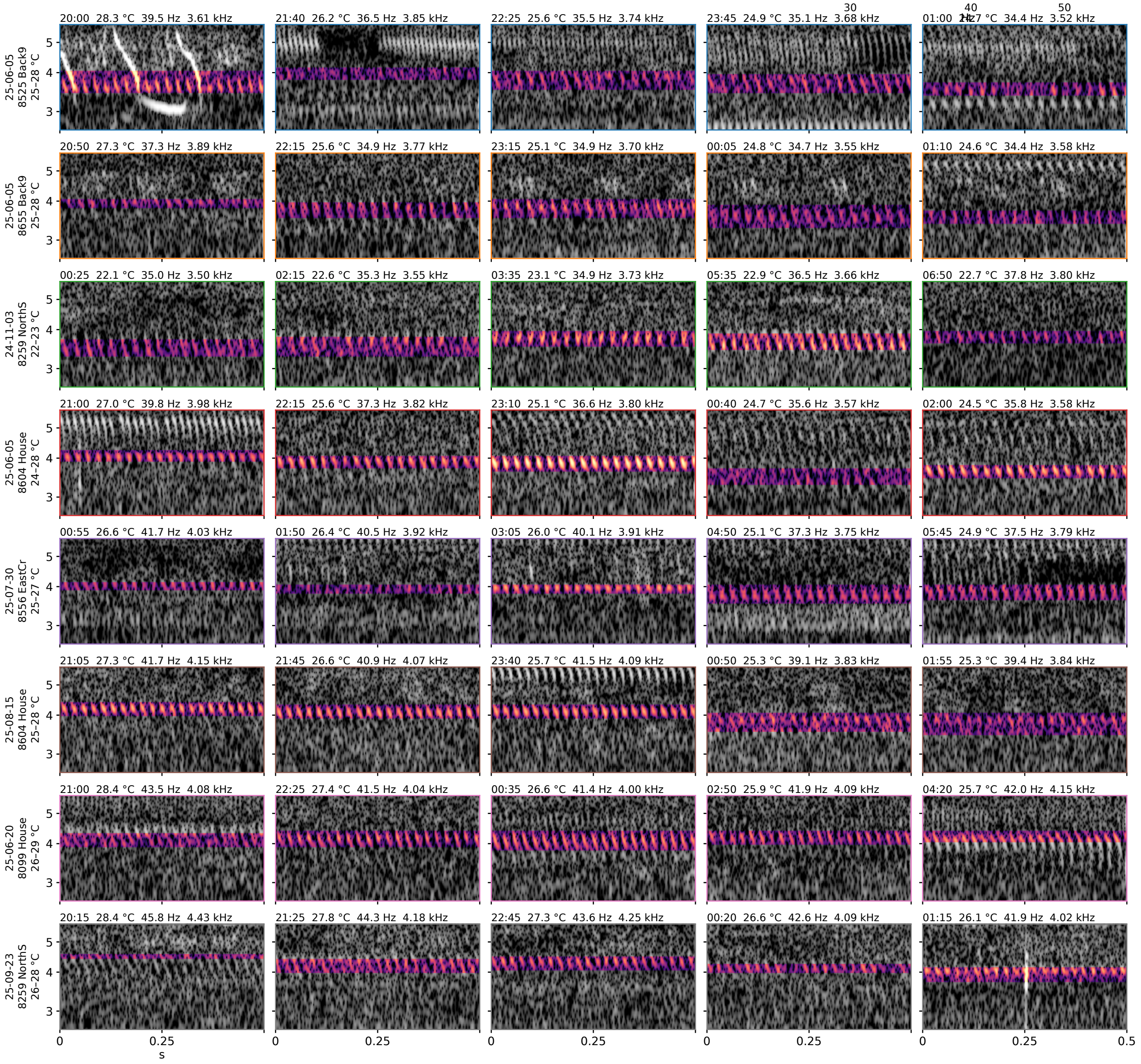


**Figure S8-11.** Type 11 (streak 360125): 3.95 kHz @ 39 Hz at 25 °C, eight singers through their nights. Rows are individual singers of the type, the eight with the widest temperature range within one night (top to bottom by the singer's median rate); columns are moments through that singer's night in time order, 5 windows at even steps through its span. Each cell is 0.5 s of the loudest 30 s of that window on one shared log-frequency axis, the detected band in color (dB above the row's quiet level, 0 to 30) and the rest in grayscale. The cell head gives the local time, the air temperature, the syllable rate and the carrier. Top right: each singer's hourly path on the plane, in the row's color, over the year's points in gray.

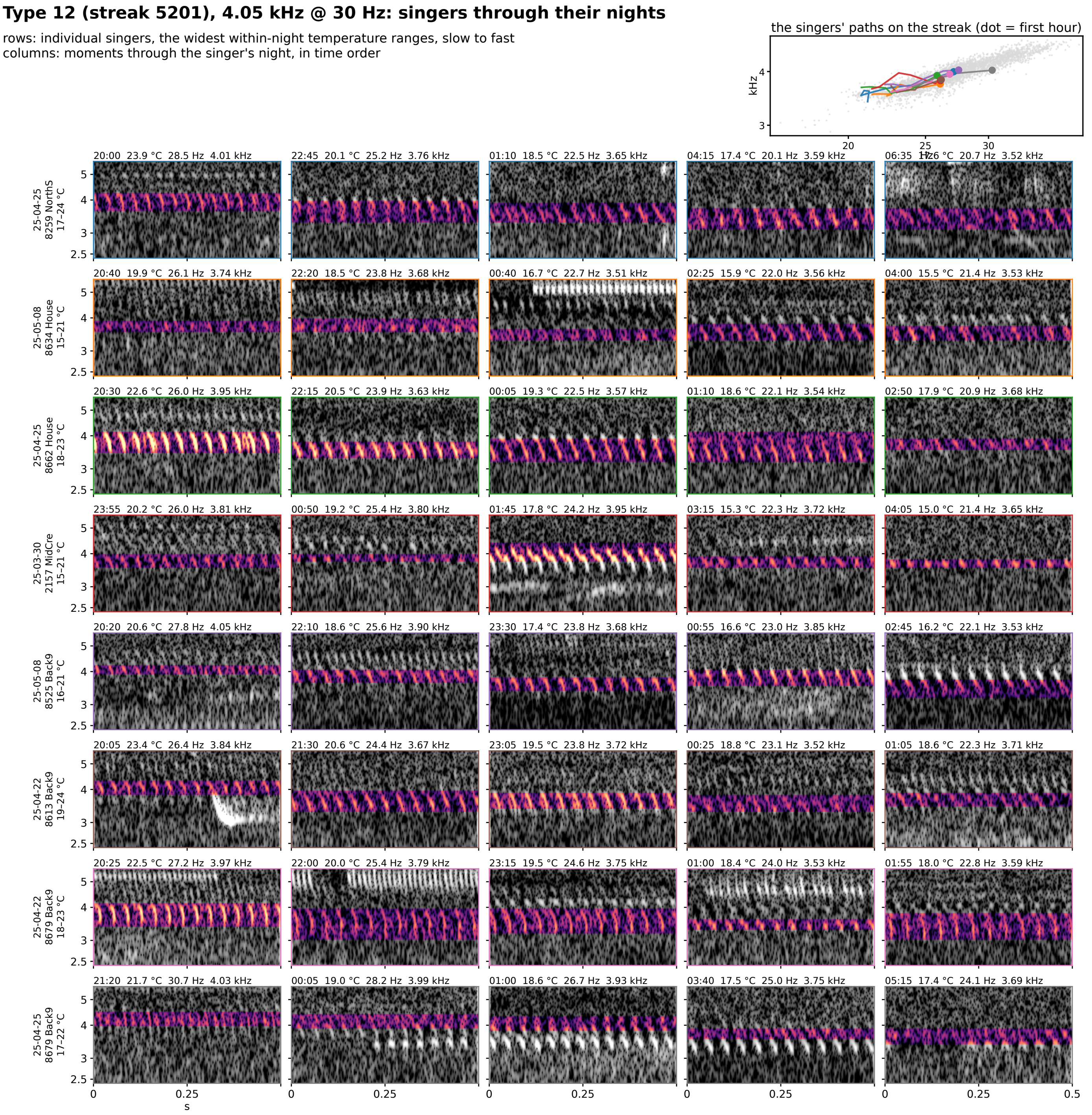


**Figure S8-12.** Type 12 (streak 5201): 4.05 kHz @ 30 Hz at 25 °C, eight singers through their nights. Rows are individual singers of the type, the eight with the widest temperature range within one night (top to bottom by the singer's median rate); columns are moments through that singer's night in time order, 5 windows at even steps through its span. Each cell is 0.5 s of the loudest 30 s of that window on one shared log-frequency axis, the detected band in color (dB above the row's quiet level, 0 to 30) and the rest in grayscale. The cell head gives the local time, the air temperature, the syllable rate and the carrier. Top right: each singer's hourly path on the plane, in the row's color, over the year's points in gray.

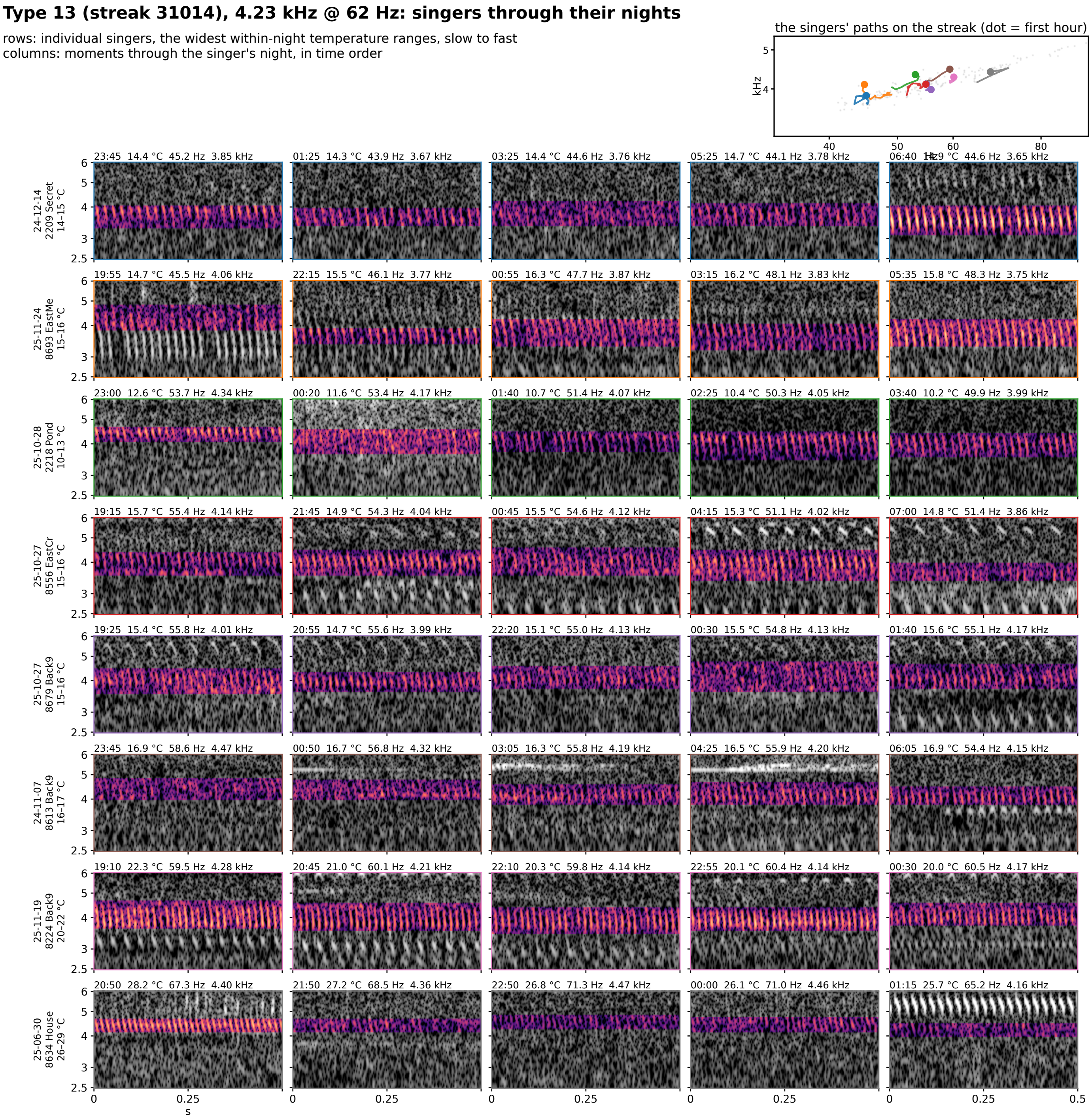


**Figure S8-13.** Type 13 (streak 31014): 4.23 kHz @ 62 Hz at 25 °C, eight singers through their nights. Rows are individual singers of the type, the eight with the widest temperature range within one night (top to bottom by the singer's median rate); columns are moments through that singer's night in time order, 5 windows at even steps through its span. Each cell is 0.5 s of the loudest 30 s of that window on one shared log-frequency axis, the detected band in color (dB above the row's quiet level, 0 to 30) and the rest in grayscale. The cell head gives the local time, the air temperature, the syllable rate and the carrier. Top right: each singer's hourly path on the plane, in the row's color, over the year's points in gray.

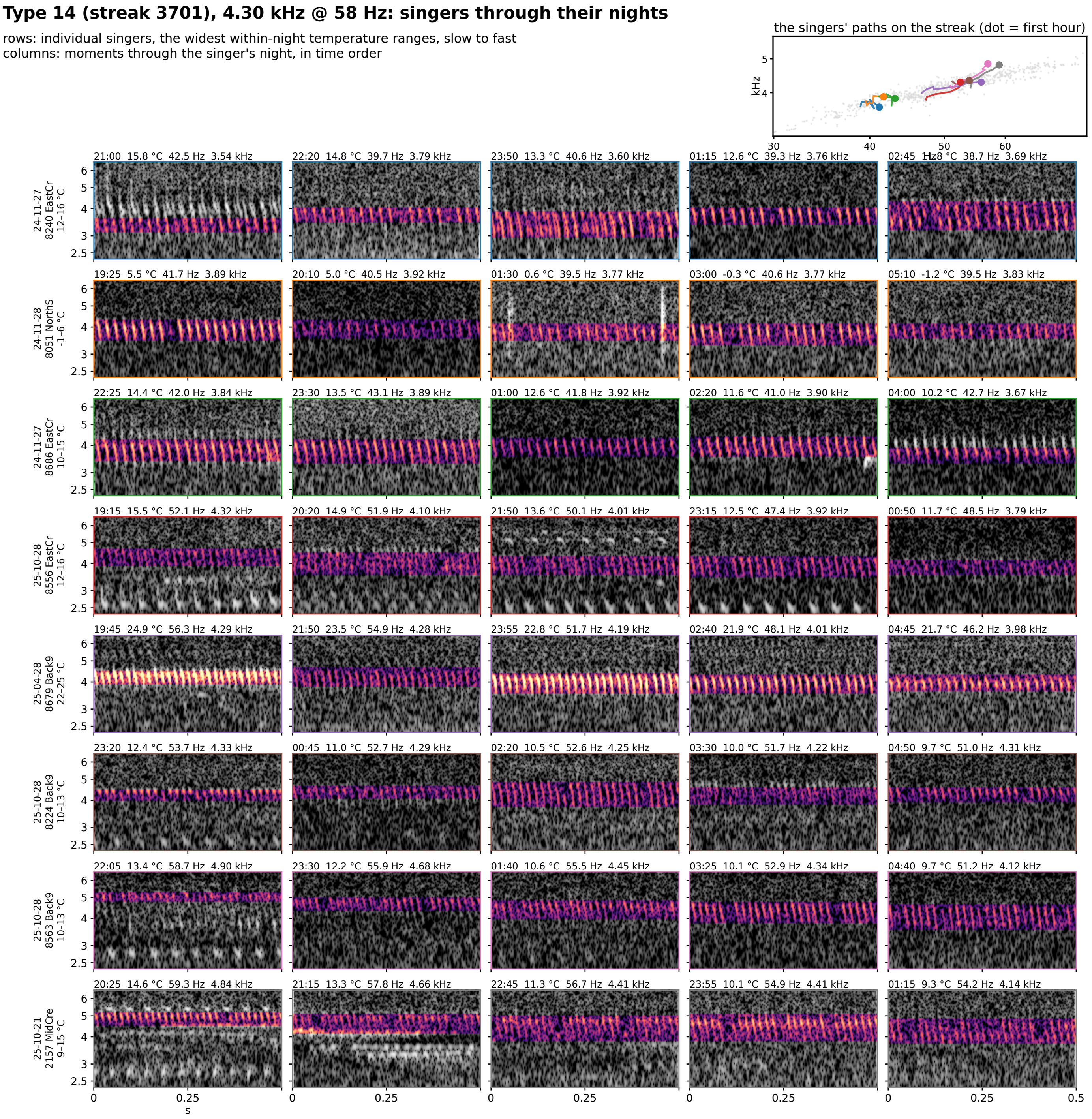


**Figure S8-14.** Type 14 (streak 3701): 4.30 kHz @ 58 Hz at 25 °C, eight singers through their nights. Rows are individual singers of the type, the eight with the widest temperature range within one night (top to bottom by the singer's median rate); columns are moments through that singer's night in time order, 5 windows at even steps through its span. Each cell is 0.5 s of the loudest 30 s of that window on one shared log-frequency axis, the detected band in color (dB above the row's quiet level, 0 to 30) and the rest in grayscale. The cell head gives the local time, the air temperature, the syllable rate and the carrier. Top right: each singer's hourly path on the plane, in the row's color, over the year's points in gray.

### Type 15 (streak 34013), 4.34 kHz @ 43 Hz: singers through their nights

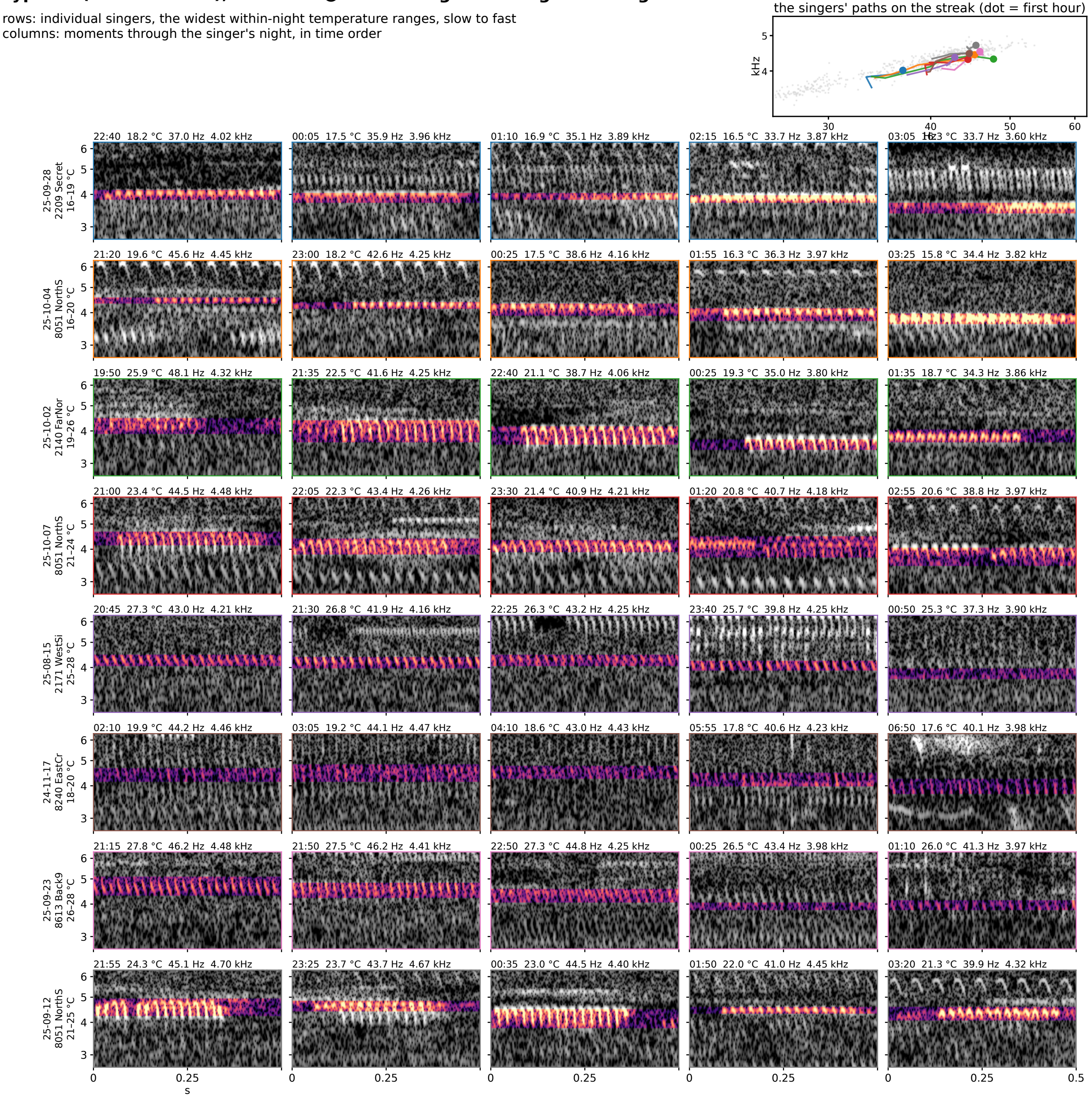


**Figure S8-15.** Type 15 (streak 34013): 4.34 kHz @ 43 Hz at 25 °C, eight singers through their nights. Rows are individual singers of the type, the eight with the widest temperature range within one night (top to bottom by the singer's median rate); columns are moments through that singer's night in time order, 5 windows at even steps through its span. Each cell is 0.5 s of the loudest 30 s of that window on one shared log-frequency axis, the detected band in color (dB above the row's quiet level, 0 to 30) and the rest in grayscale. The cell head gives the local time, the air temperature, the syllable rate and the carrier. Top right: each singer's hourly path on the plane, in the row's color, over the year's points in gray.

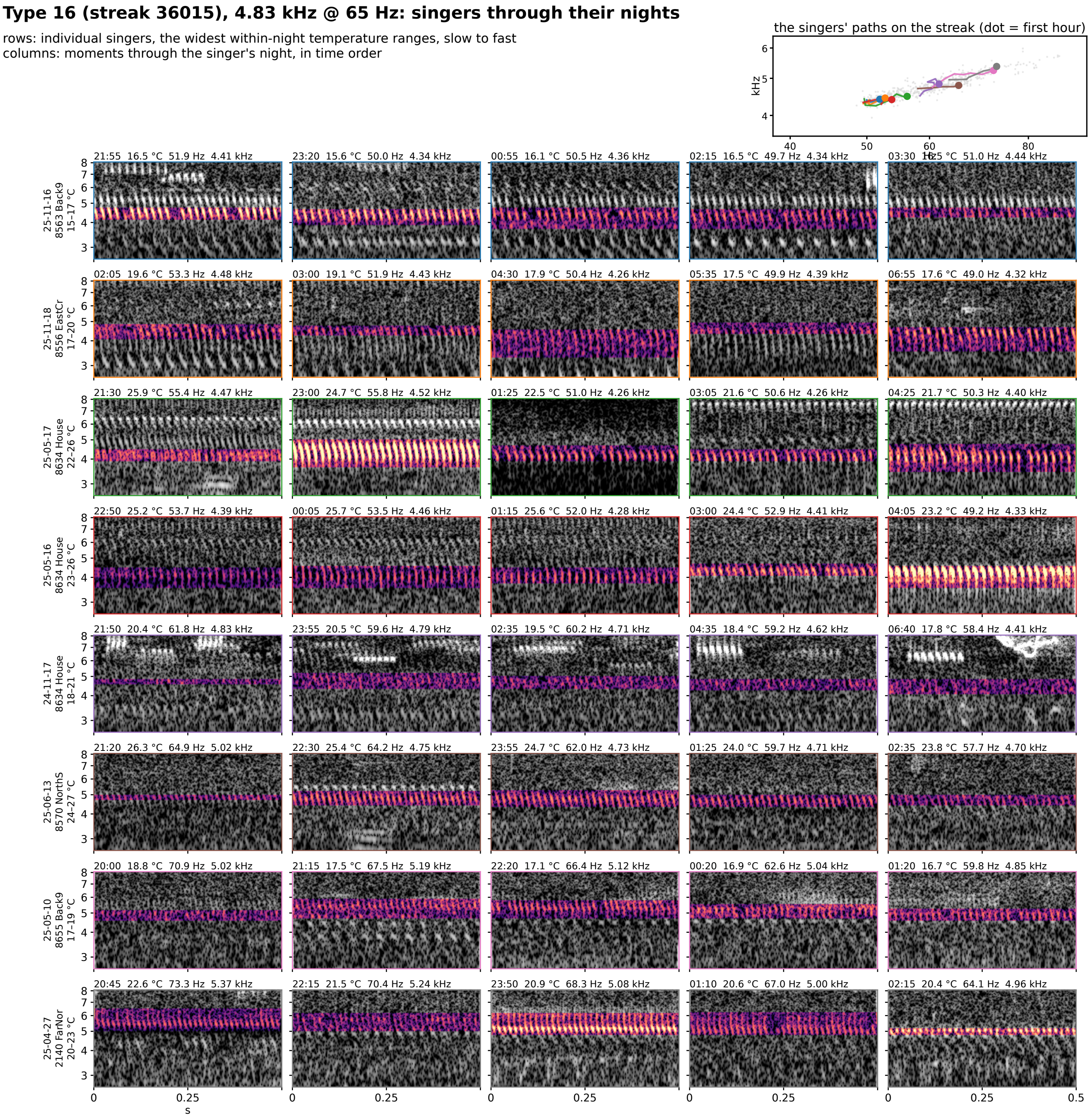


**Figure S8-16.** Type 16 (streak 36015): 4.83 kHz @ 65 Hz at 25 °C, eight singers through their nights. Rows are individual singers of the type, the eight with the widest temperature range within one night (top to bottom by the singer's median rate); columns are moments through that singer's night in time order, 5 windows at even steps through its span. Each cell is 0.5 s of the loudest 30 s of that window on one shared log-frequency axis, the detected band in color (dB above the row's quiet level, 0 to 30) and the rest in grayscale. The cell head gives the local time, the air temperature, the syllable rate and the carrier. Top right: each singer's hourly path on the plane, in the row's color, over the year's points in gray.

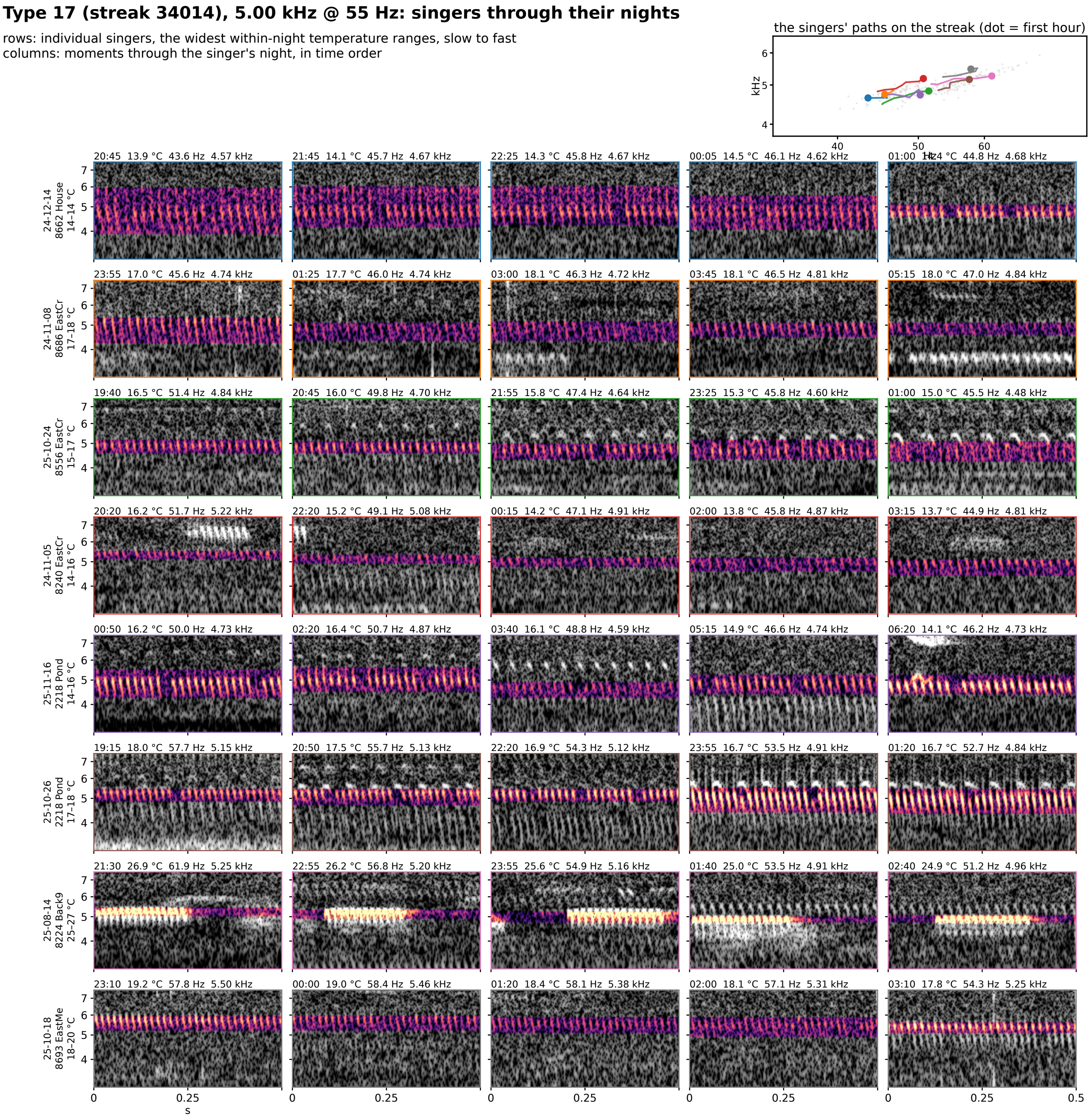


**Figure S8-17.** Type 17 (streak 34014): 5.00 kHz @ 55 Hz at 25 °C, eight singers through their nights. Rows are individual singers of the type, the eight with the widest temperature range within one night (top to bottom by the singer's median rate); columns are moments through that singer's night in time order, 5 windows at even steps through its span. Each cell is 0.5 s of the loudest 30 s of that window on one shared log-frequency axis, the detected band in color (dB above the row's quiet level, 0 to 30) and the rest in grayscale. The cell head gives the local time, the air temperature, the syllable rate and the carrier. Top right: each singer's hourly path on the plane, in the row's color, over the year's points in gray.

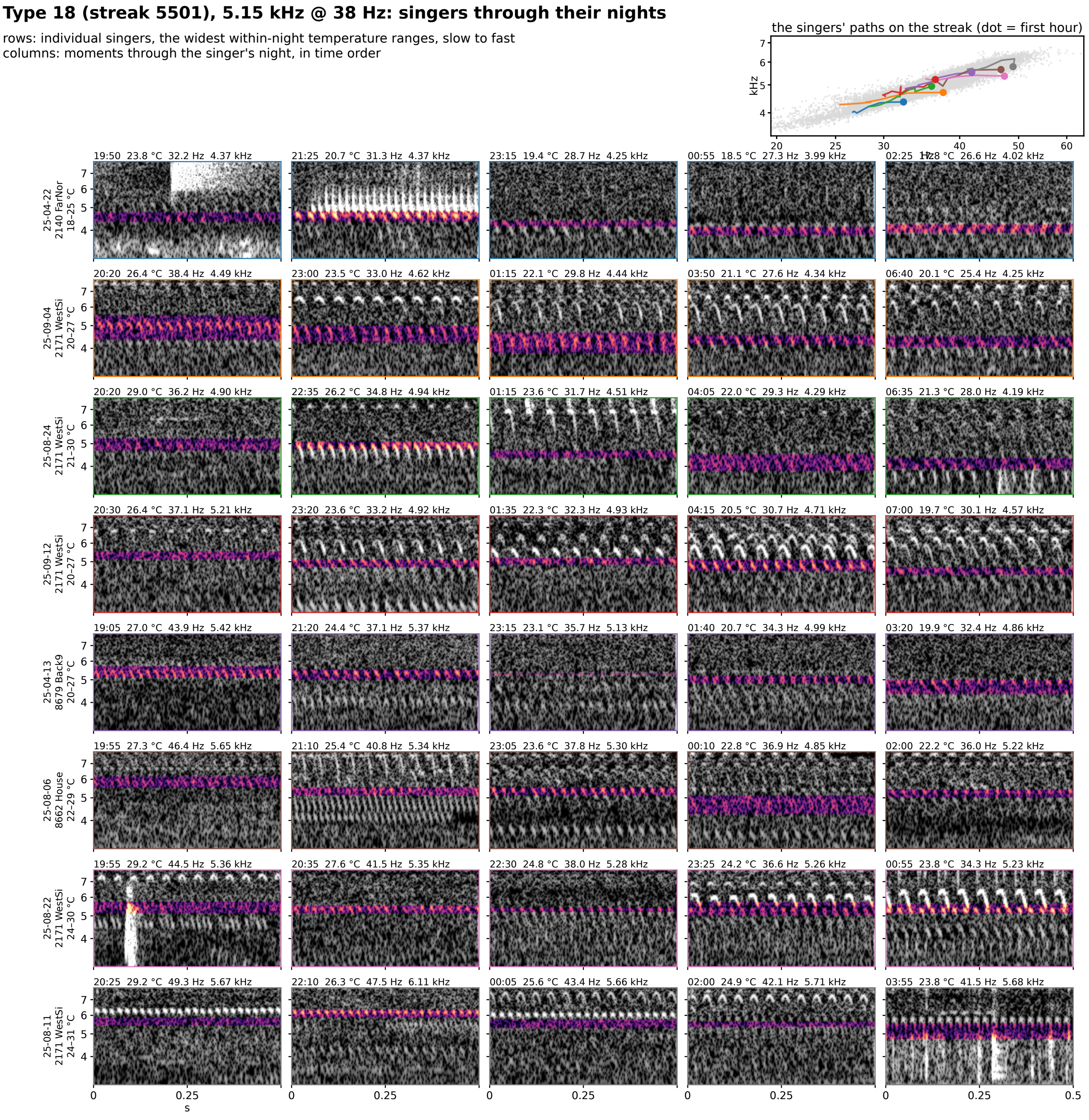


**Figure S8-18.** Type 18 (streak 5501): 5.15 kHz @ 38 Hz at 25 °C, eight singers through their nights. Rows are individual singers of the type, the eight with the widest temperature range within one night (top to bottom by the singer's median rate); columns are moments through that singer's night in time order, 5 windows at even steps through its span. Each cell is 0.5 s of the loudest 30 s of that window on one shared log-frequency axis, the detected band in color (dB above the row's quiet level, 0 to 30) and the rest in grayscale. The cell head gives the local time, the air temperature, the syllable rate and the carrier. Top right: each singer's hourly path on the plane, in the row's color, over the year's points in gray.

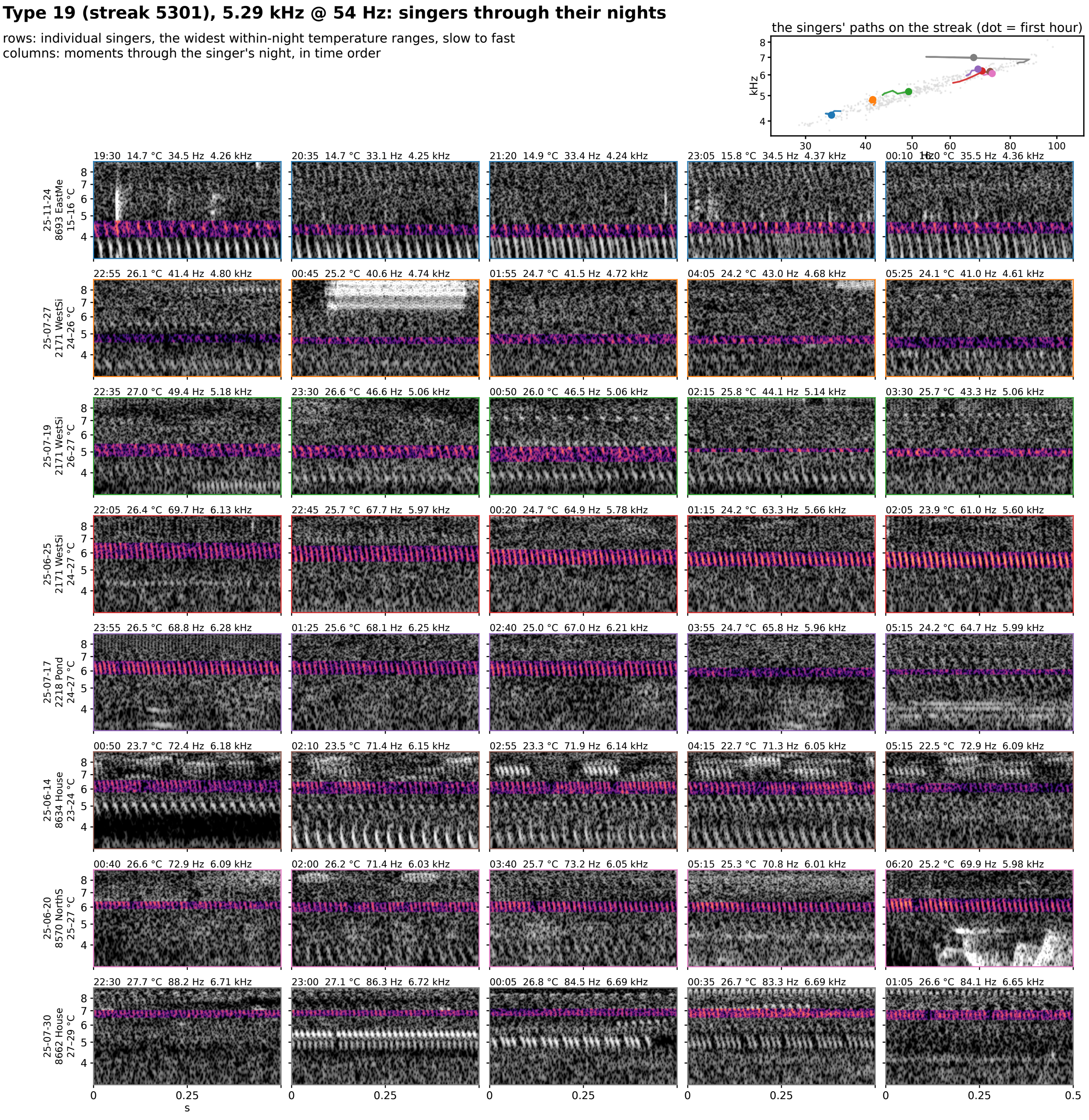


**Figure S8-19.** Type 19 (streak 5301): 5.29 kHz @ 54 Hz at 25 °C, eight singers through their nights. Rows are individual singers of the type, the eight with the widest temperature range within one night (top to bottom by the singer's median rate); columns are moments through that singer's night in time order, 5 windows at even steps through its span. Each cell is 0.5 s of the loudest 30 s of that window on one shared log-frequency axis, the detected band in color (dB above the row's quiet level, 0 to 30) and the rest in grayscale. The cell head gives the local time, the air temperature, the syllable rate and the carrier. Top right: each singer's hourly path on the plane, in the row's color, over the year's points in gray.

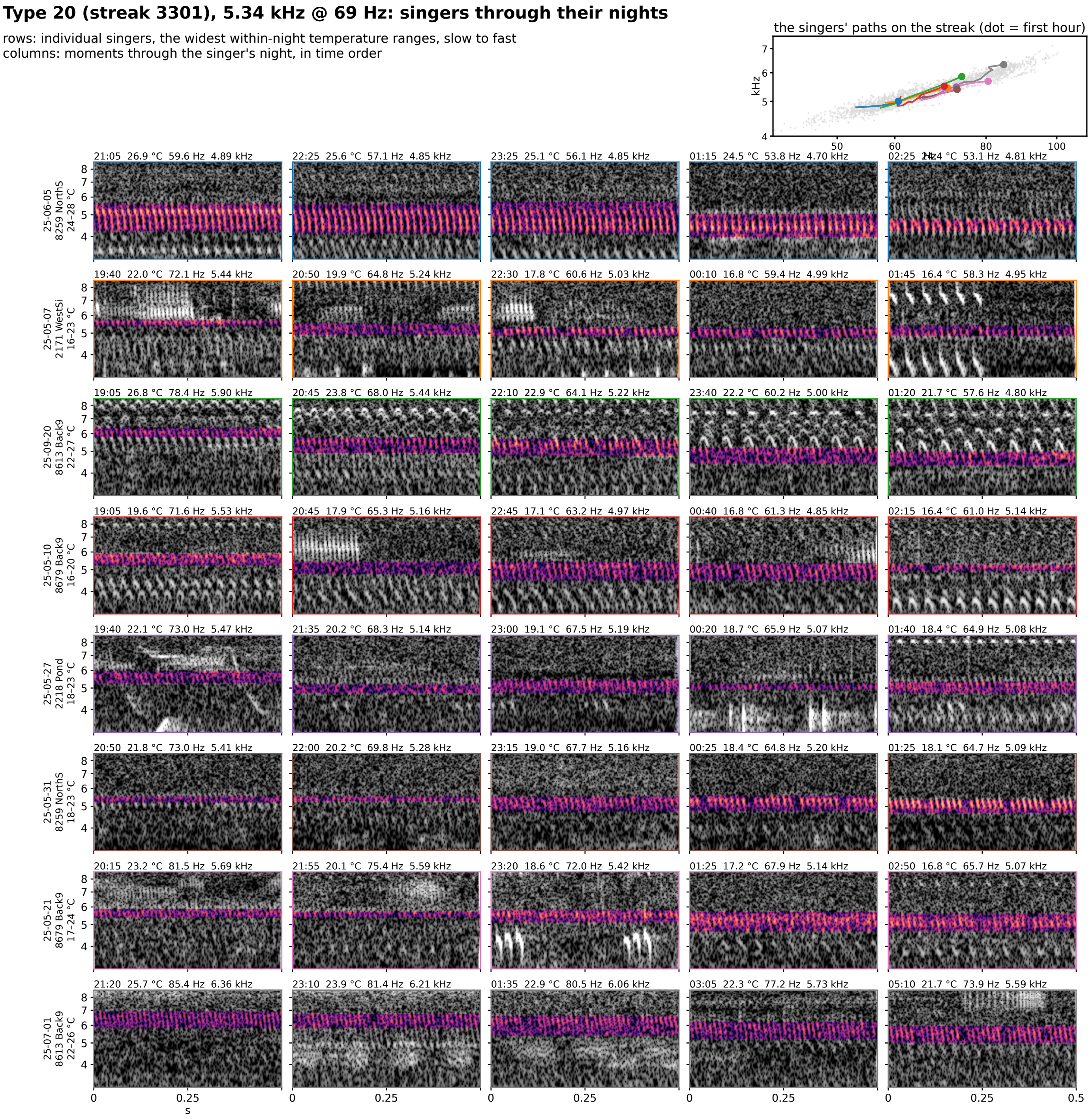


**Figure S8-20.** Type 20 (streak 3301): 5.34 kHz @ 69 Hz at 25 °C, eight singers through their nights. Rows are individual singers of the type, the eight with the widest temperature range within one night (top to bottom by the singer's median rate); columns are moments through that singer's night in time order, 5 windows at even steps through its span. Each cell is 0.5 s of the loudest 30 s of that window on one shared log-frequency axis, the detected band in color (dB above the row's quiet level, 0 to 30) and the rest in grayscale. The cell head gives the local time, the air temperature, the syllable rate and the carrier. Top right: each singer's hourly path on the plane, in the row's color, over the year's points in gray.

**Type 21 (streak 6301), 5.66 kHz @ 20 Hz: singers through their nights**

rows: individual singers, the widest within-night temperature ranges, slow to fast
columns: moments through the singer's night, in time order

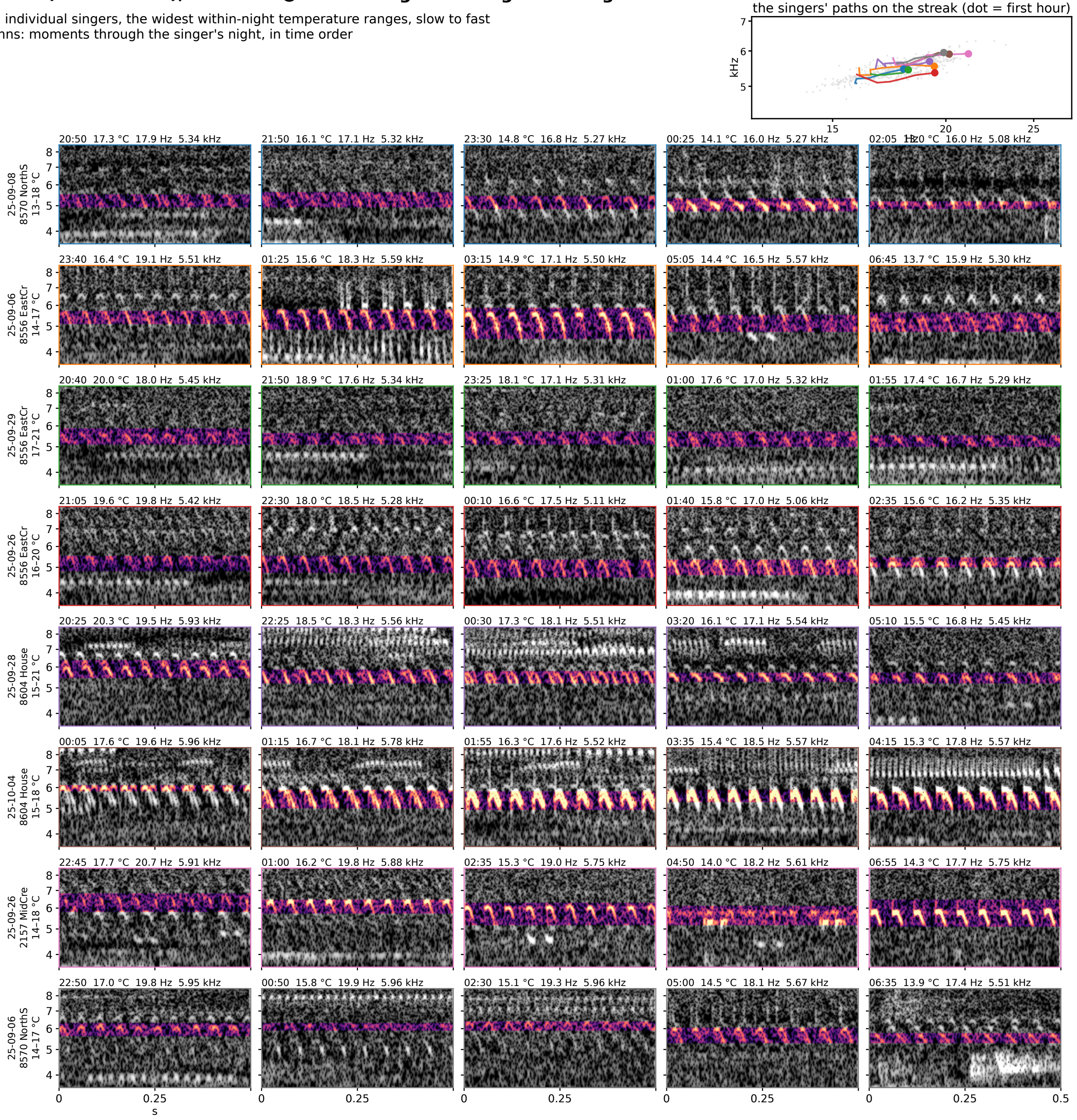


**Figure S8-21.** Type 21 (streak 6301): 5.66 kHz @ 20 Hz at 25 °C, eight singers through their nights. Rows are individual singers of the type, the eight with the widest temperature range within one night (top to bottom by the singer's median rate); columns are moments through that singer's night in time order, 5 windows at even steps through its span. Each cell is 0.5 s of the loudest 30 s of that window on one shared log-frequency axis, the detected band in color (dB above the row's quiet level, 0 to 30) and the rest in grayscale. The cell head gives the local time, the air temperature, the syllable rate and the carrier. Top right: each singer's hourly path on the plane, in the row's color, over the year's points in gray.

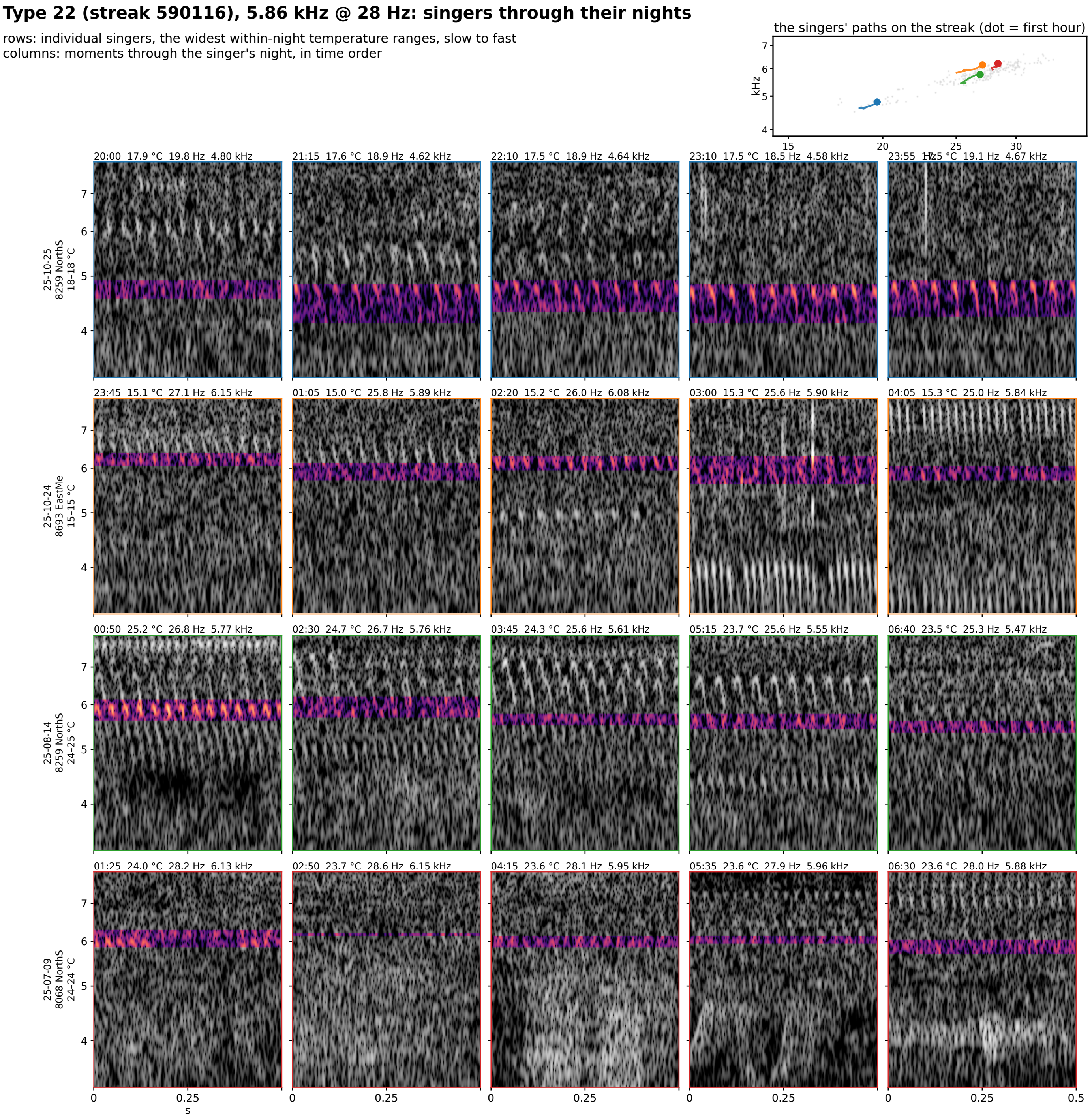


**Figure S8-22.** Type 22 (streak 590116): 5.86 kHz @ 28 Hz at 25 °C, eight singers through their nights. Rows are individual singers of the type, the eight with the widest temperature range within one night (top to bottom by the singer's median rate); columns are moments through that singer's night in time order, 5 windows at even steps through its span. Each cell is 0.5 s of the loudest 30 s of that window on one shared log-frequency axis, the detected band in color (dB above the row's quiet level, 0 to 30) and the rest in grayscale. The cell head gives the local time, the air temperature, the syllable rate and the carrier. Top right: each singer's hourly path on the plane, in the row's color, over the year's points in gray.

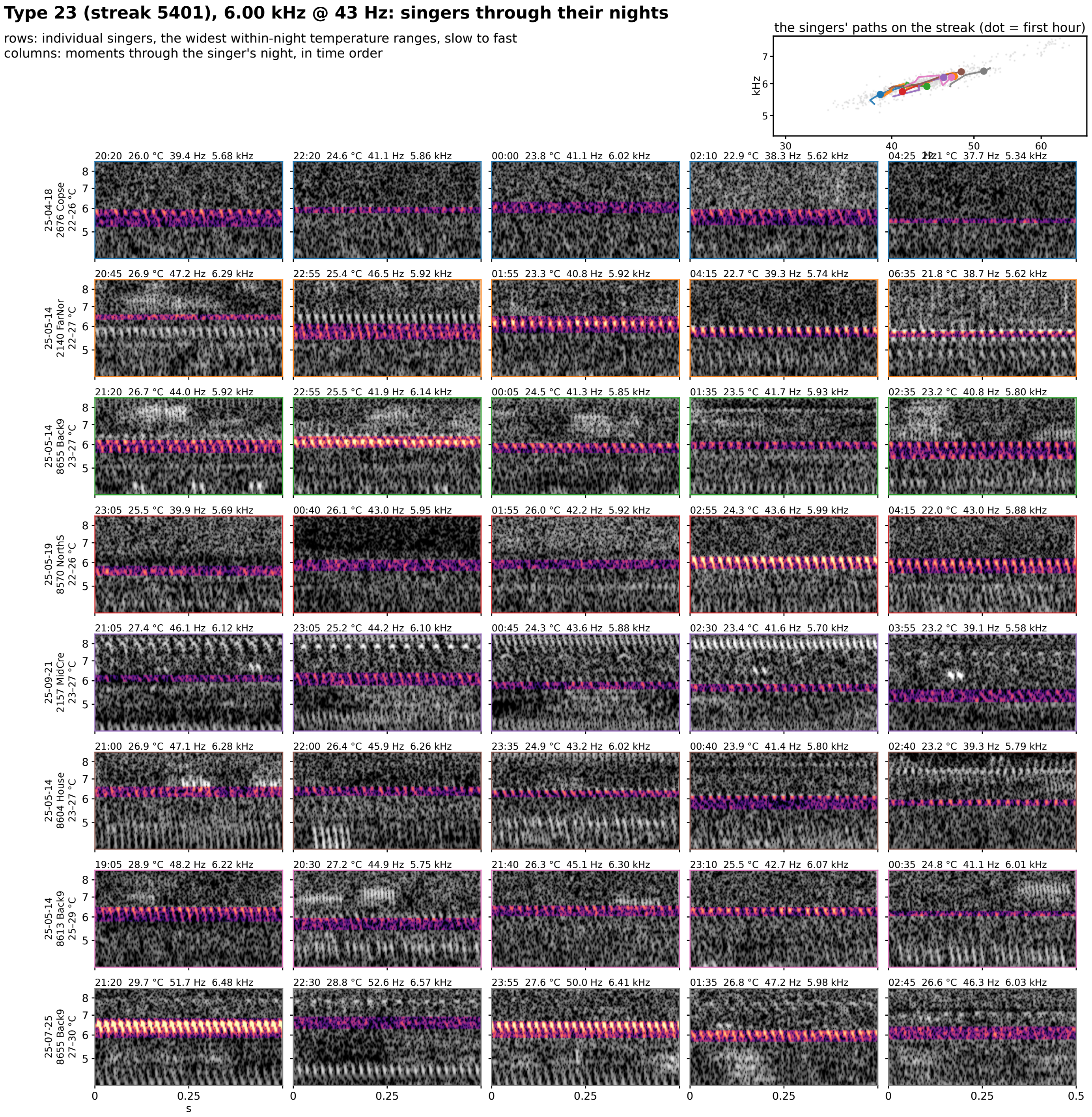


**Figure S8-23.** Type 23 (streak 5401): 6.00 kHz @ 43 Hz at 25 °C, eight singers through their nights. Rows are individual singers of the type, the eight with the widest temperature range within one night (top to bottom by the singer's median rate); columns are moments through that singer's night in time order, 5 windows at even steps through its span. Each cell is 0.5 s of the loudest 30 s of that window on one shared log-frequency axis, the detected band in color (dB above the row's quiet level, 0 to 30) and the rest in grayscale. The cell head gives the local time, the air temperature, the syllable rate and the carrier. Top right: each singer's hourly path on the plane, in the row's color, over the year's points in gray.

### Type 24 (streak 4601), 6.20 kHz @ 20 Hz: singers through their nights

rows: individual singers, the widest within-night temperature ranges, slow to fast
columns: moments through the singer's night, in time order

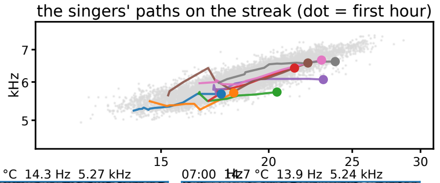


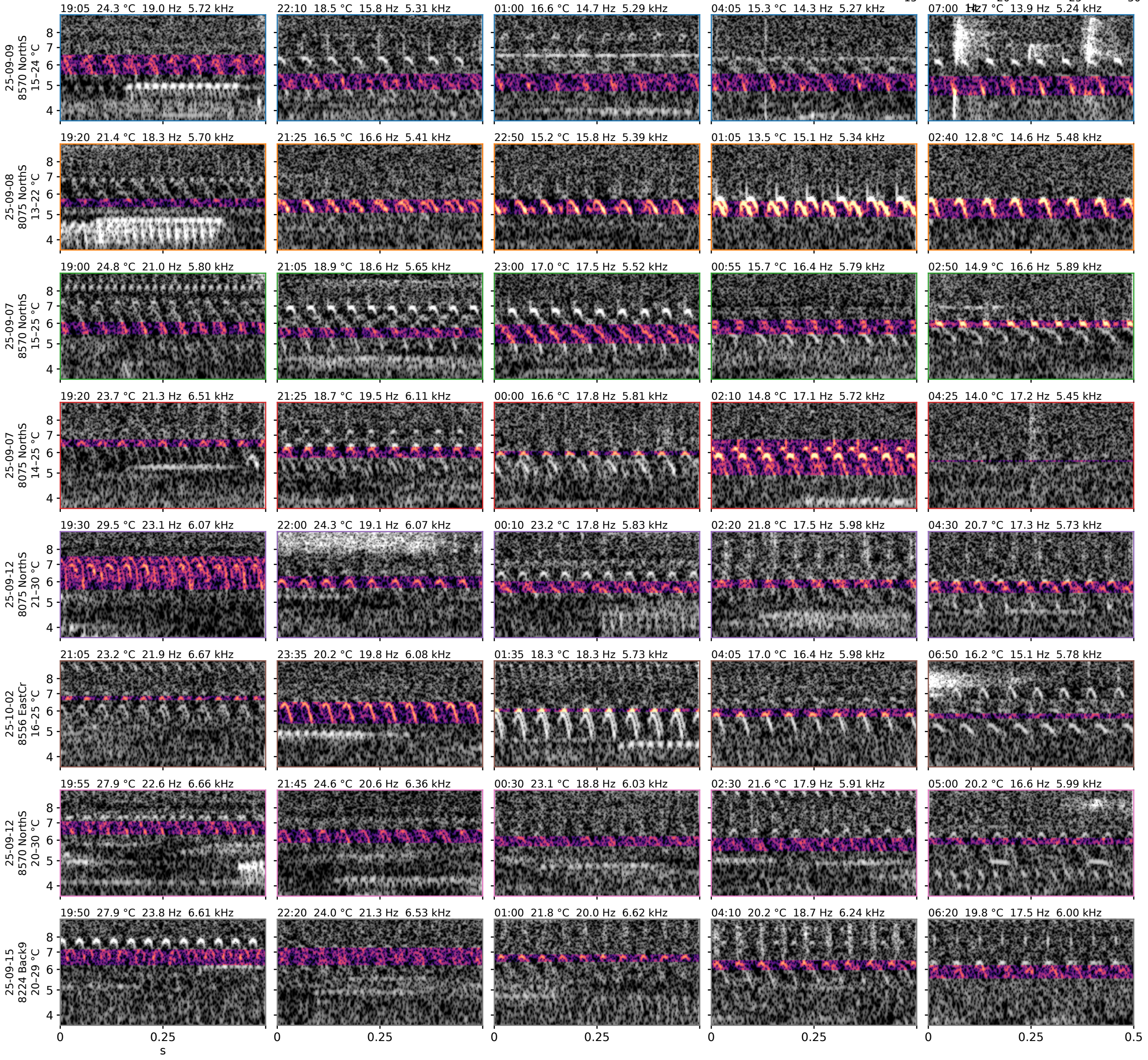


**Figure S8-24.** Type 24 (streak 4601): 6.20 kHz @ 20 Hz at 25 °C, eight singers through their nights. Rows are individual singers of the type, the eight with the widest temperature range within one night (top to bottom by the singer's median rate); columns are moments through that singer's night in time order, 5 windows at even steps through its span. Each cell is 0.5 s of the loudest 30 s of that window on one shared log-frequency axis, the detected band in color (dB above the row's quiet level, 0 to 30) and the rest in grayscale. The cell head gives the local time, the air temperature, the syllable rate and the carrier. Top right: each singer's hourly path on the plane, in the row's color, over the year's points in gray.

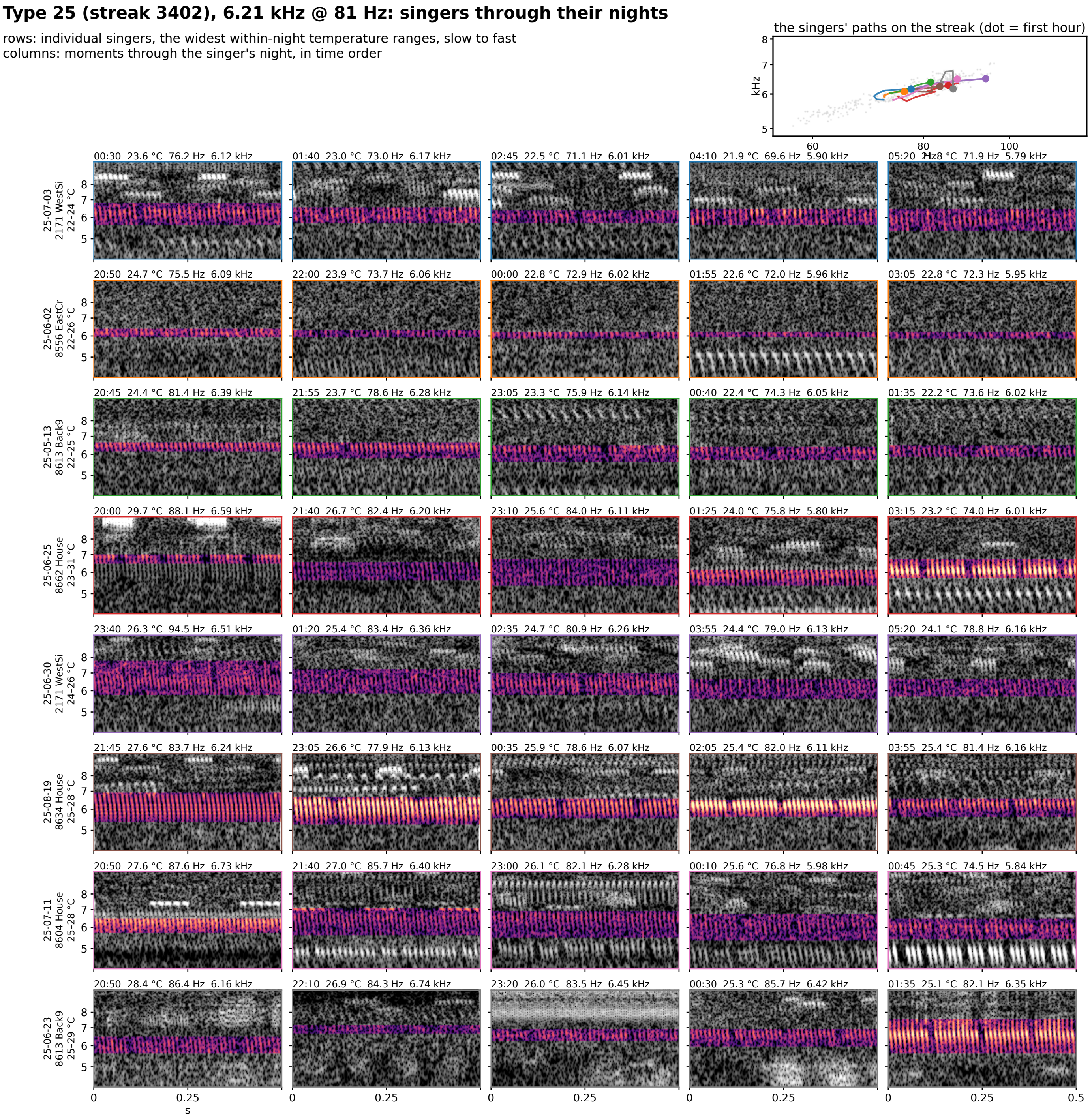


**Figure S8-25.** Type 25 (streak 3402): 6.21 kHz @ 81 Hz at 25 °C, eight singers through their nights. Rows are individual singers of the type, the eight with the widest temperature range within one night (top to bottom by the singer's median rate); columns are moments through that singer's night in time order, 5 windows at even steps through its span. Each cell is 0.5 s of the loudest 30 s of that window on one shared log-frequency axis, the detected band in color (dB above the row's quiet level, 0 to 30) and the rest in grayscale. The cell head gives the local time, the air temperature, the syllable rate and the carrier. Top right: each singer's hourly path on the plane, in the row's color, over the year's points in gray.

### Type 26 (streak 6001), 6.38 kHz @ 29 Hz: singers through their nights

rows: individual singers, the widest within-night temperature ranges, slow to fast
columns: moments through the singer's night, in time order

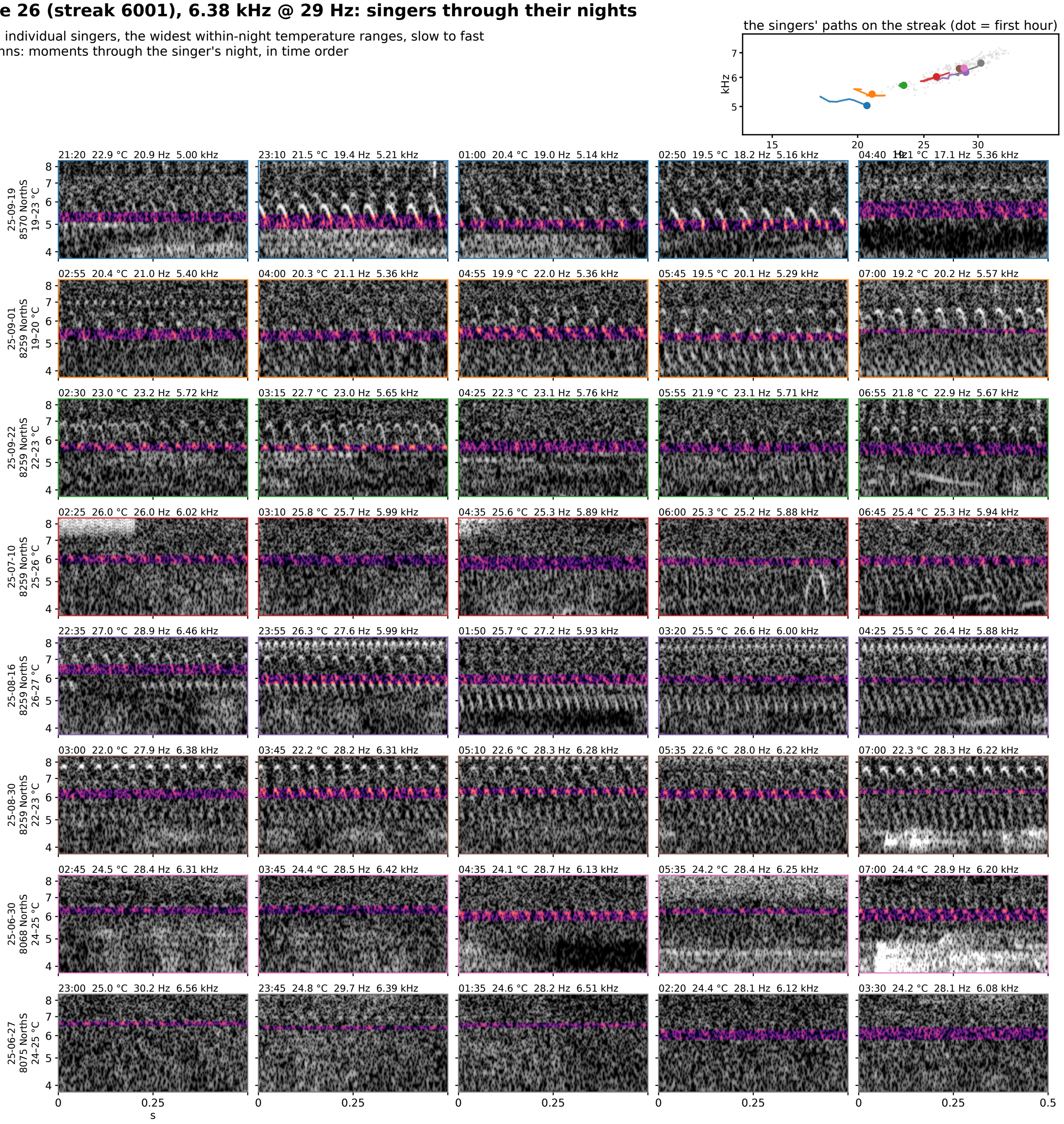


**Figure S8-26.** Type 26 (streak 6001): 6.38 kHz @ 29 Hz at 25 °C, eight singers through their nights. Rows are individual singers of the type, the eight with the widest temperature range within one night (top to bottom by the singer's median rate); columns are moments through that singer's night in time order, 5 windows at even steps through its span. Each cell is 0.5 s of the loudest 30 s of that window on one shared log-frequency axis, the detected band in color (dB above the row's quiet level, 0 to 30) and the rest in grayscale. The cell head gives the local time, the air temperature, the syllable rate and the carrier. Top right: each singer's hourly path on the plane, in the row's color, over the year's points in gray.

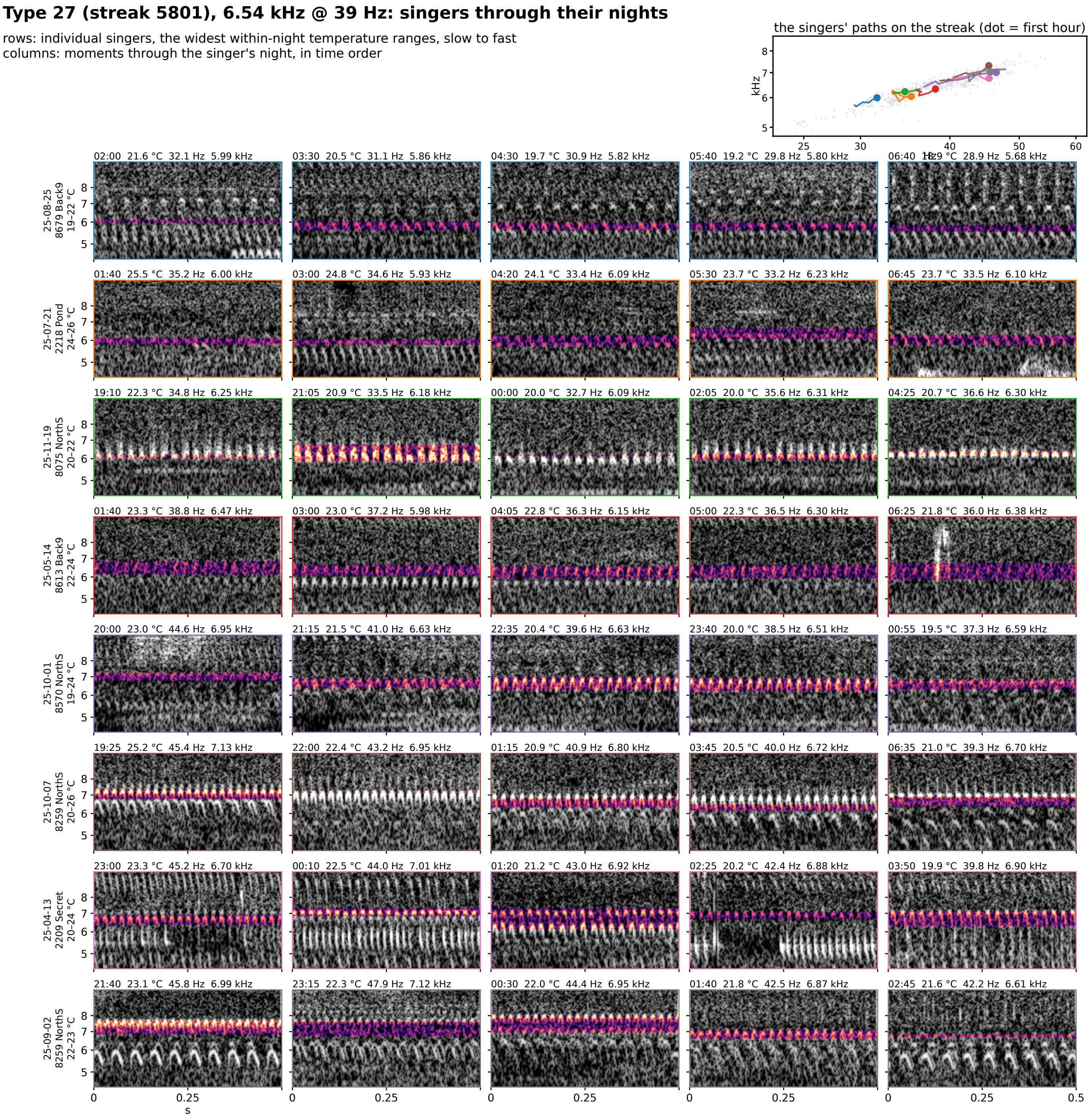


**Figure S8-27.** Type 27 (streak 5801): 6.54 kHz @ 39 Hz at 25 °C, eight singers through their nights. Rows are individual singers of the type, the eight with the widest temperature range within one night (top to bottom by the singer's median rate); columns are moments through that singer's night in time order, 5 windows at even steps through its span. Each cell is 0.5 s of the loudest 30 s of that window on one shared log-frequency axis, the detected band in color (dB above the row's quiet level, 0 to 30) and the rest in grayscale. The cell head gives the local time, the air temperature, the syllable rate and the carrier. Top right: each singer's hourly path on the plane, in the row's color, over the year's points in gray.

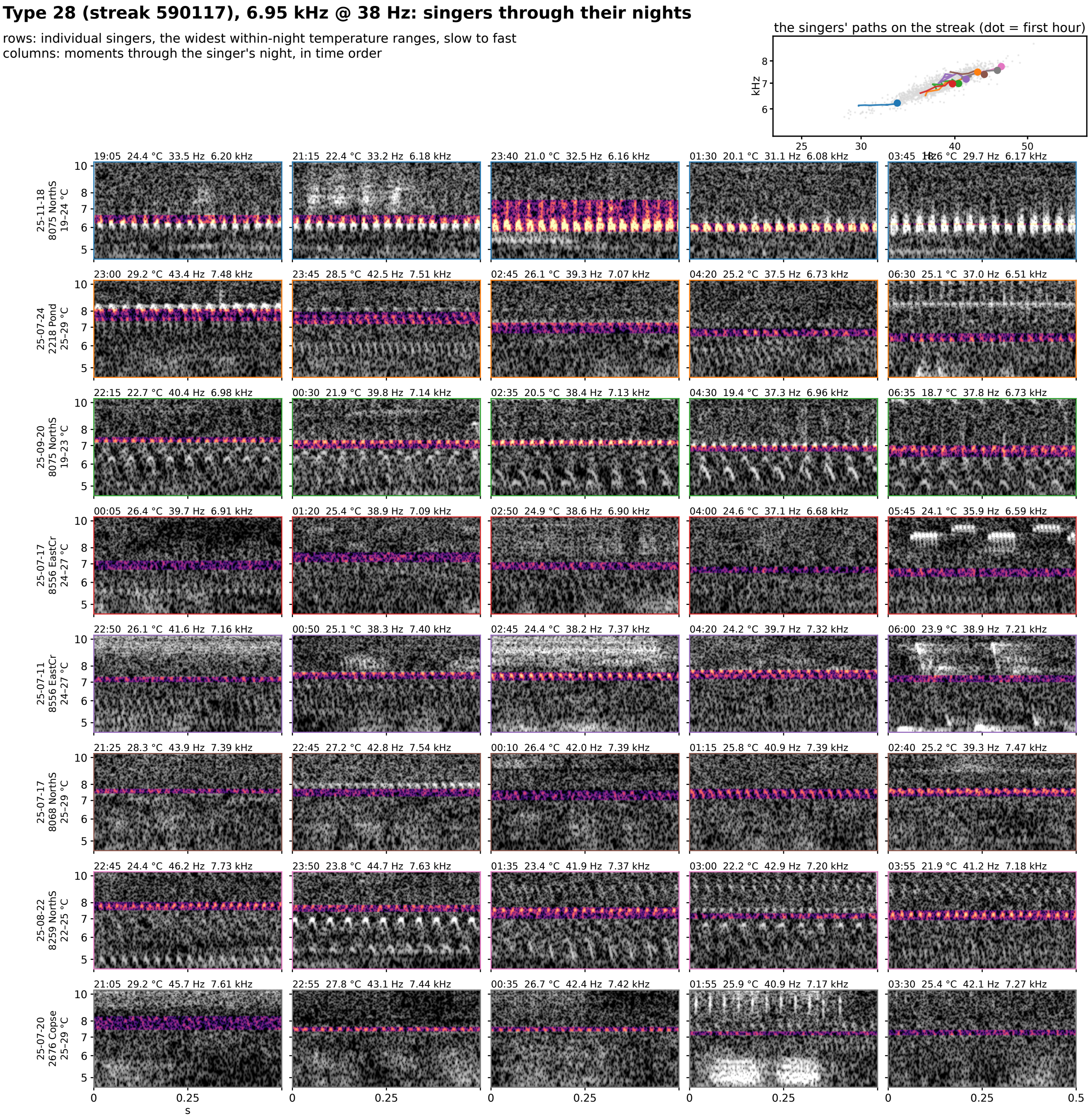


**Figure S8-28.** Type 28 (streak 590117): 6.95 kHz @ 38 Hz at 25 °C, eight singers through their nights. Rows are individual singers of the type, the eight with the widest temperature range within one night (top to bottom by the singer's median rate); columns are moments through that singer's night in time order, 5 windows at even steps through its span. Each cell is 0.5 s of the loudest 30 s of that window on one shared log-frequency axis, the detected band in color (dB above the row's quiet level, 0 to 30) and the rest in grayscale. The cell head gives the local time, the air temperature, the syllable rate and the carrier. Top right: each singer's hourly path on the plane, in the row's color, over the year's points in gray.

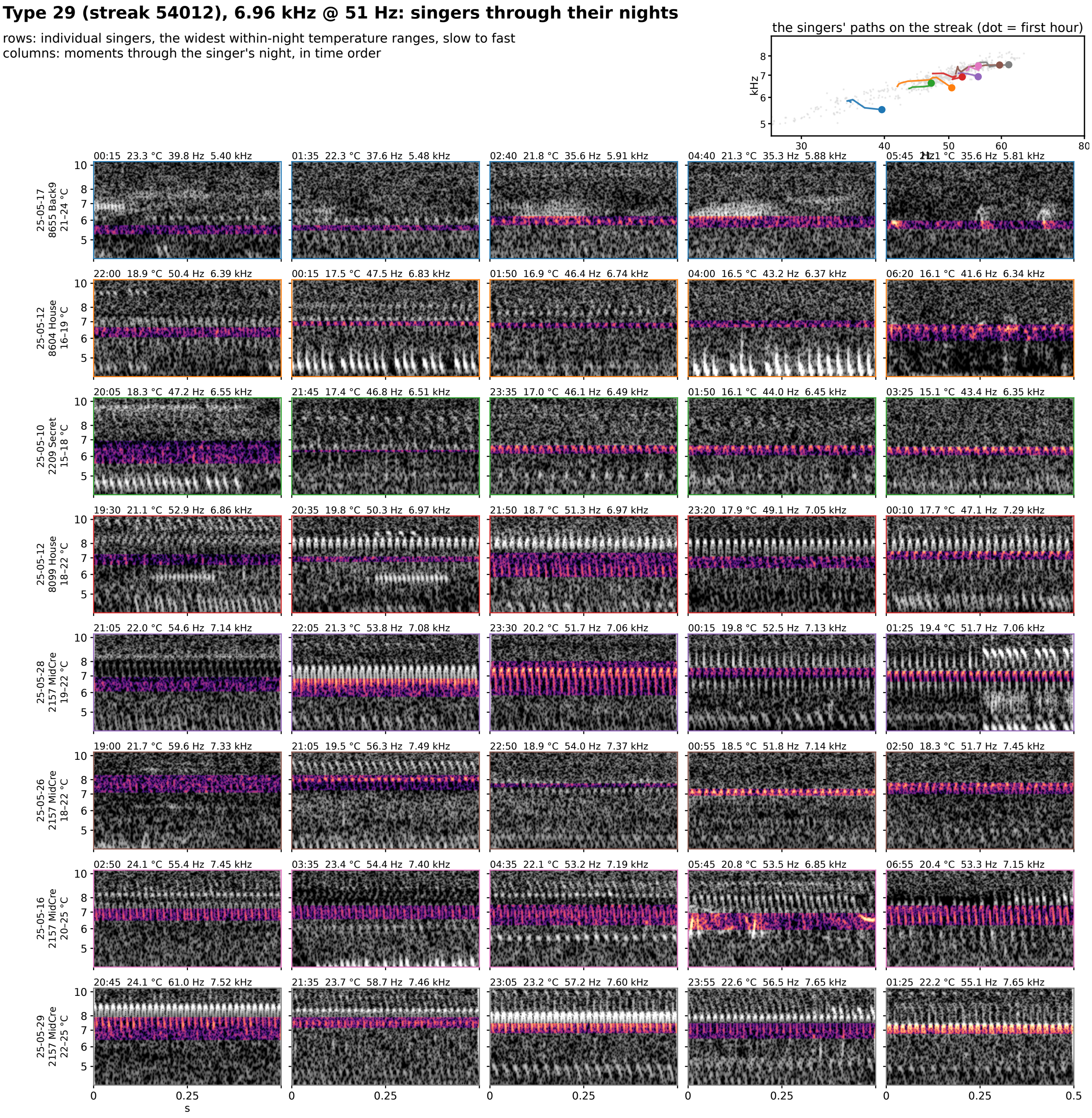


**Figure S8-29.** Type 29 (streak 54012): 6.96 kHz @ 51 Hz at 25 °C, eight singers through their nights. Rows are individual singers of the type, the eight with the widest temperature range within one night (top to bottom by the singer's median rate); columns are moments through that singer's night in time order, 5 windows at even steps through its span. Each cell is 0.5 s of the loudest 30 s of that window on one shared log-frequency axis, the detected band in color (dB above the row's quiet level, 0 to 30) and the rest in grayscale. The cell head gives the local time, the air temperature, the syllable rate and the carrier. Top right: each singer's hourly path on the plane, in the row's color, over the year's points in gray.

### Type 30 (streak 4701), 7.01 kHz @ 22 Hz: singers through their nights

rows: individual singers, the widest within-night temperature ranges, slow to fast
columns: moments through the singer's night, in time order

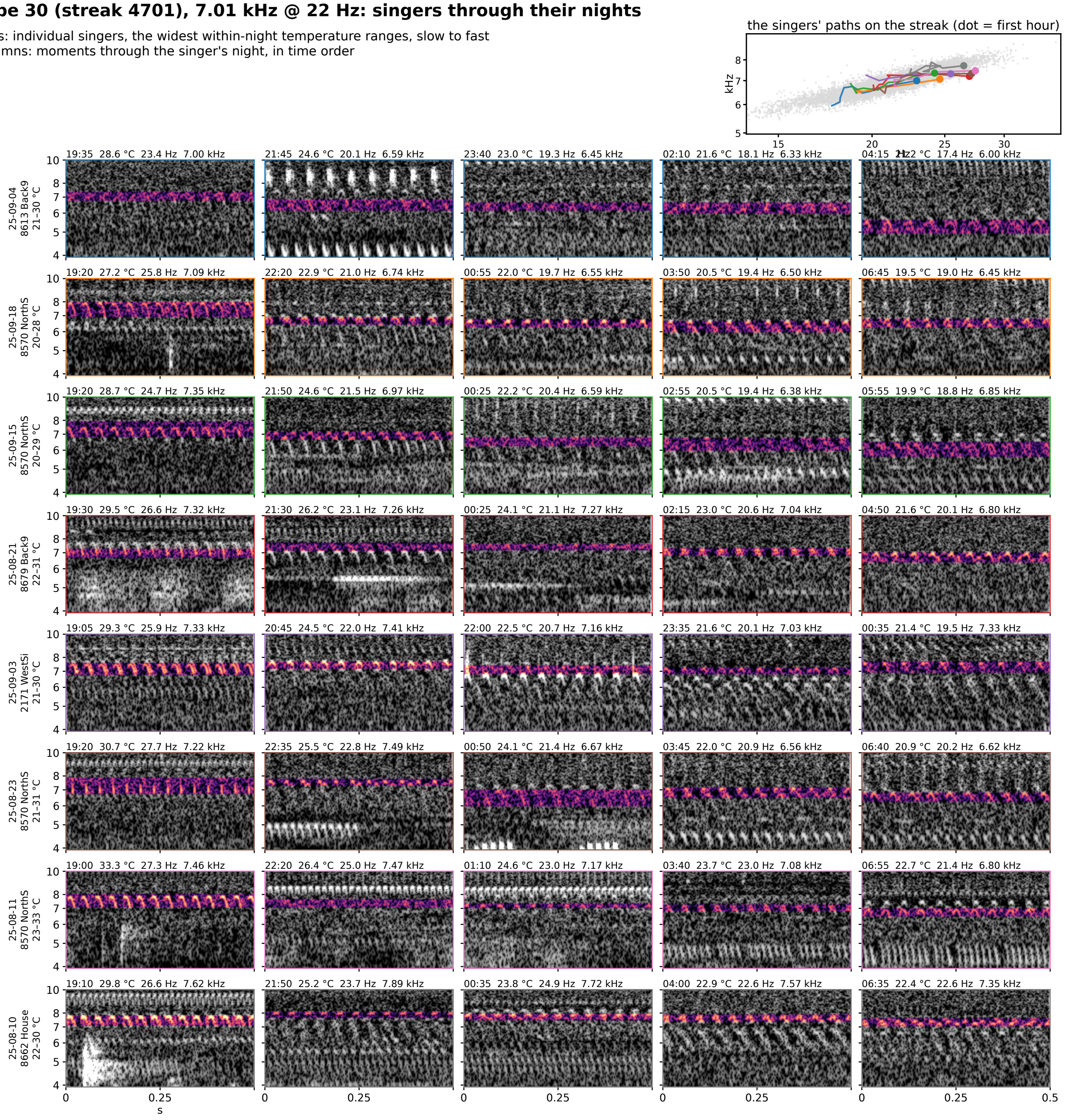


**Figure S8-30.** Type 30 (streak 4701): 7.01 kHz @ 22 Hz at 25 °C, eight singers through their nights. Rows are individual singers of the type, the eight with the widest temperature range within one night (top to bottom by the singer's median rate); columns are moments through that singer's night in time order, 5 windows at even steps through its span. Each cell is 0.5 s of the loudest 30 s of that window on one shared log-frequency axis, the detected band in color (dB above the row's quiet level, 0 to 30) and the rest in grayscale. The cell head gives the local time, the air temperature, the syllable rate and the carrier. Top right: each singer's hourly path on the plane, in the row's color, over the year's points in gray.

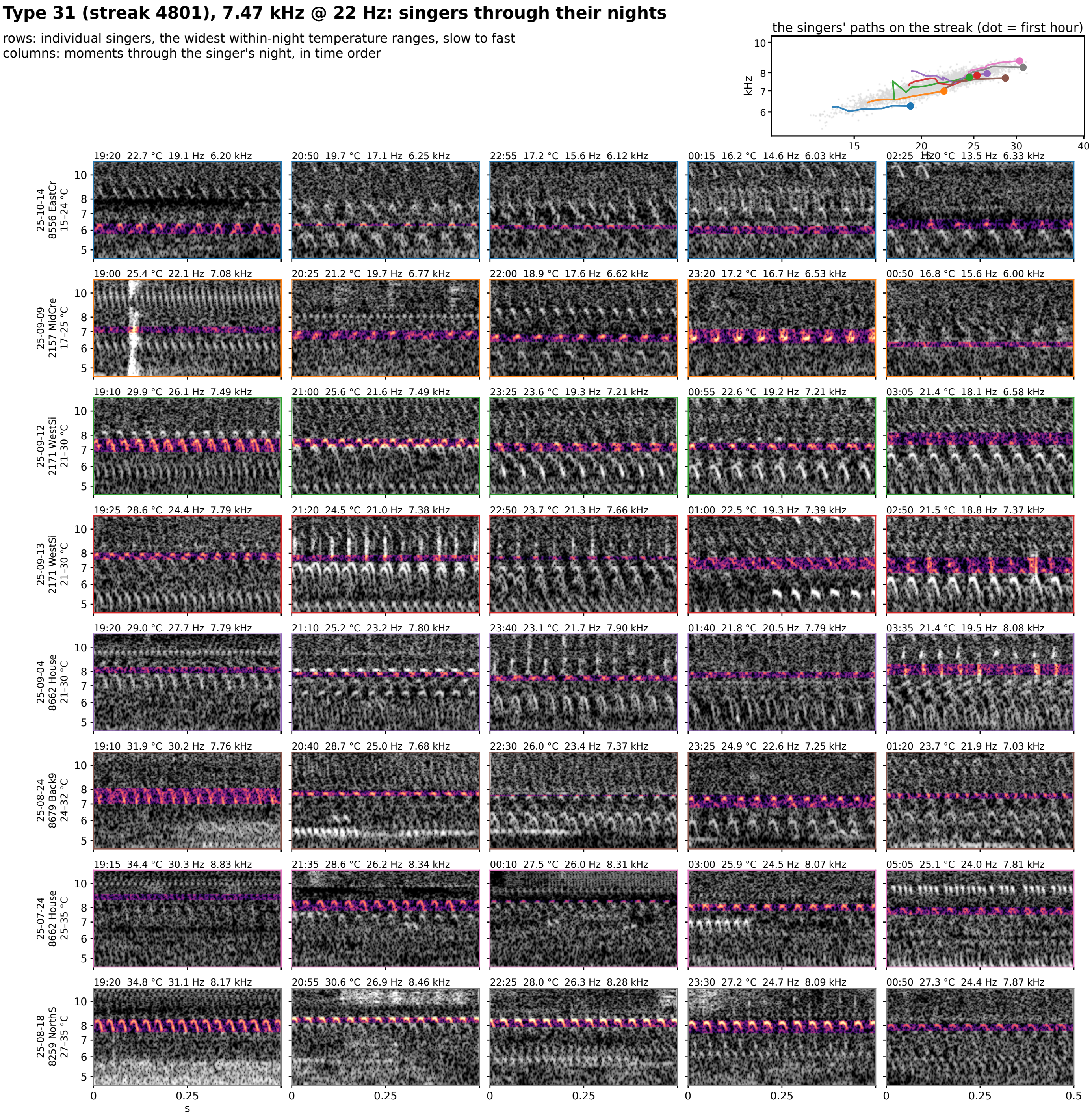


**Figure S8-31.** Type 31 (streak 4801): 7.47 kHz @ 22 Hz at 25 °C, eight singers through their nights. Rows are individual singers of the type, the eight with the widest temperature range within one night (top to bottom by the singer's median rate); columns are moments through that singer's night in time order, 5 windows at even steps through its span. Each cell is 0.5 s of the loudest 30 s of that window on one shared log-frequency axis, the detected band in color (dB above the row's quiet level, 0 to 30) and the rest in grayscale. The cell head gives the local time, the air temperature, the syllable rate and the carrier. Top right: each singer's hourly path on the plane, in the row's color, over the year's points in gray.

**Type 32 (streak 5802), 7.66 kHz @ 55 Hz: singers through their nights**

rows: individual singers, the widest within-night temperature ranges, slow to fast
columns: moments through the singer's night, in time order

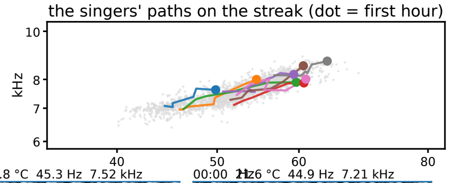


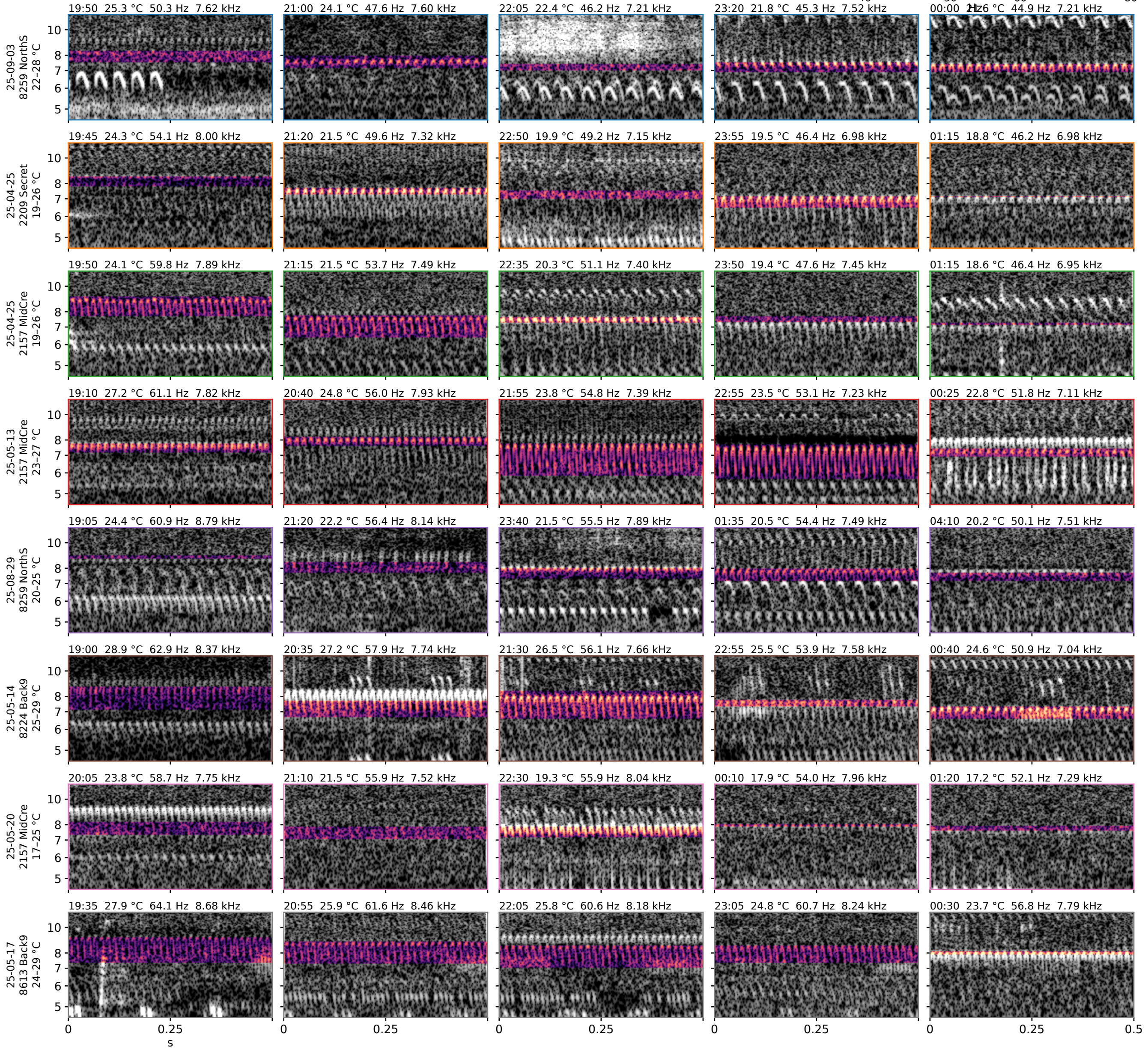


**Figure S8-32.** Type 32 (streak 5802): 7.66 kHz @ 55 Hz at 25 °C, eight singers through their nights. Rows are individual singers of the type, the eight with the widest temperature range within one night (top to bottom by the singer's median rate); columns are moments through that singer's night in time order, 5 windows at even steps through its span. Each cell is 0.5 s of the loudest 30 s of that window on one shared log-frequency axis, the detected band in color (dB above the row's quiet level, 0 to 30) and the rest in grayscale. The cell head gives the local time, the air temperature, the syllable rate and the carrier. Top right: each singer's hourly path on the plane, in the row's color, over the year's points in gray.

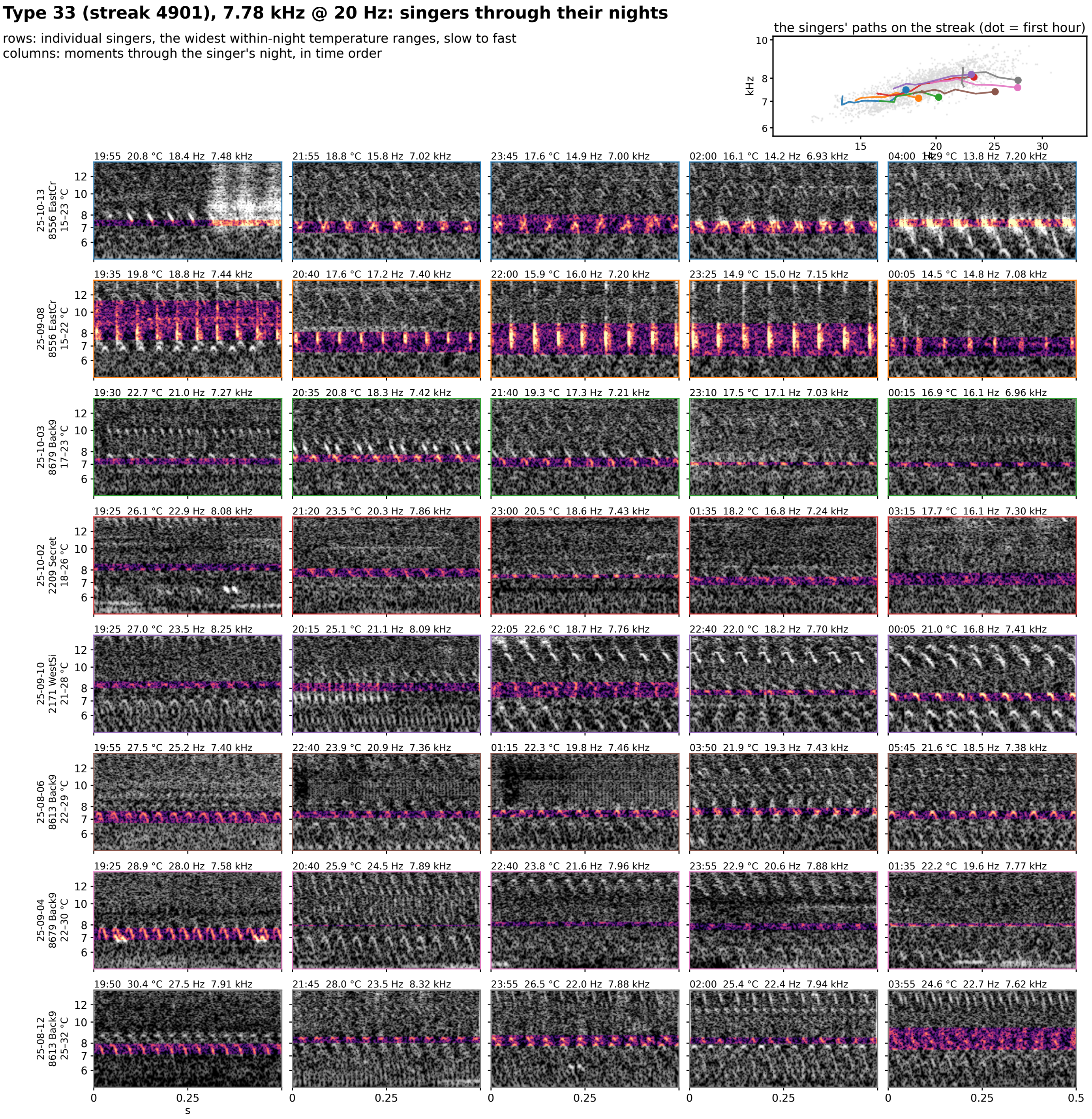


**Figure S8-33.** Type 33 (streak 4901): 7.78 kHz @ 20 Hz at 25 °C, eight singers through their nights. Rows are individual singers of the type, the eight with the widest temperature range within one night (top to bottom by the singer's median rate); columns are moments through that singer's night in time order, 5 windows at even steps through its span. Each cell is 0.5 s of the loudest 30 s of that window on one shared log-frequency axis, the detected band in color (dB above the row's quiet level, 0 to 30) and the rest in grayscale. The cell head gives the local time, the air temperature, the syllable rate and the carrier. Top right: each singer's hourly path on the plane, in the row's color, over the year's points in gray.

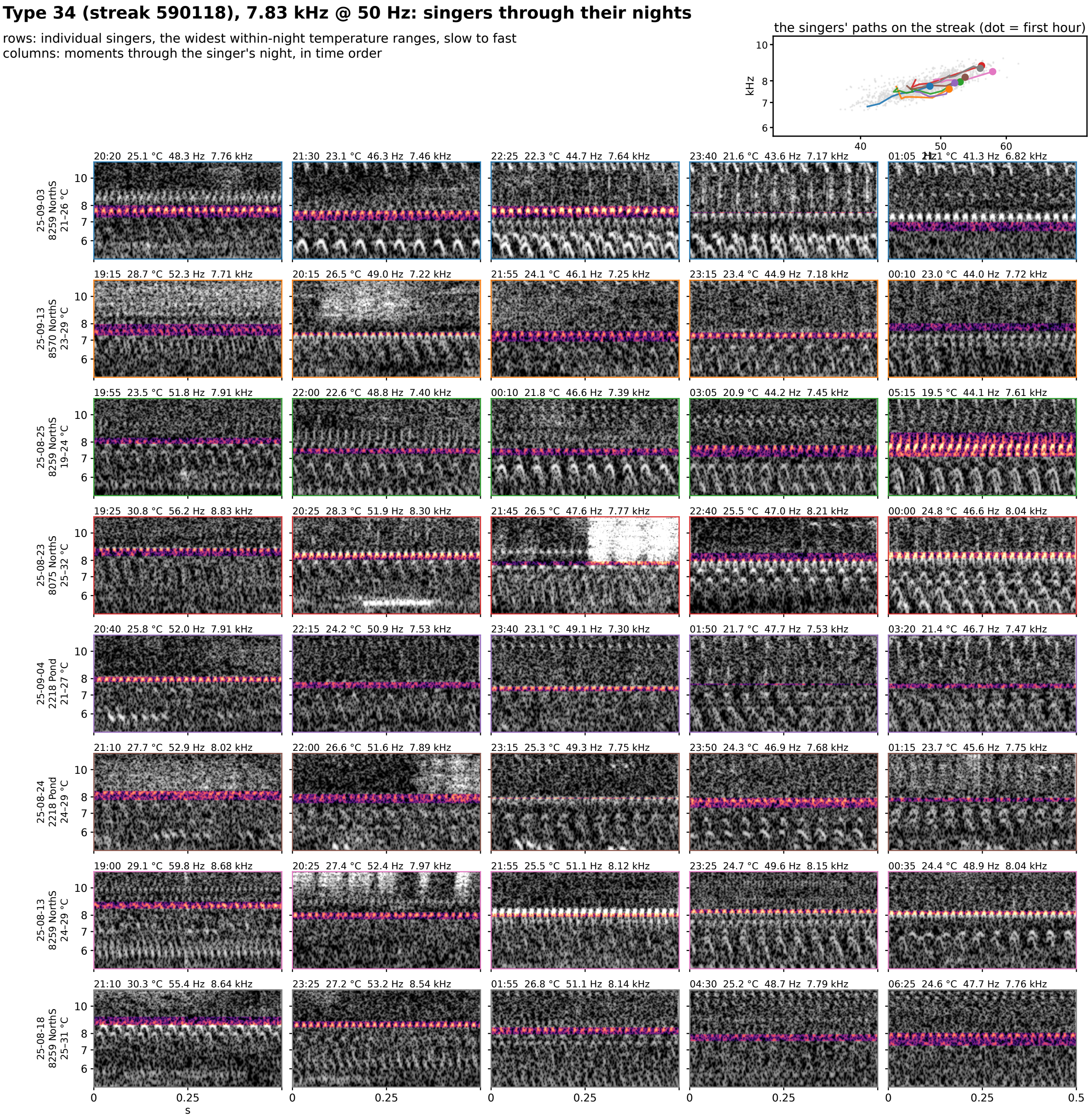


**Figure S8-34.** Type 34 (streak 590118): 7.83 kHz @ 50 Hz at 25 °C, eight singers through their nights. Rows are individual singers of the type, the eight with the widest temperature range within one night (top to bottom by the singer's median rate); columns are moments through that singer's night in time order, 5 windows at even steps through its span. Each cell is 0.5 s of the loudest 30 s of that window on one shared log-frequency axis, the detected band in color (dB above the row's quiet level, 0 to 30) and the rest in grayscale. The cell head gives the local time, the air temperature, the syllable rate and the carrier. Top right: each singer's hourly path on the plane, in the row's color, over the year's points in gray.

### Type 35 (streak 5901), 8.55 kHz @ 58 Hz: singers through their nights

rows: individual singers, the widest within-night temperature ranges, slow to fast
columns: moments through the singer's night, in time order

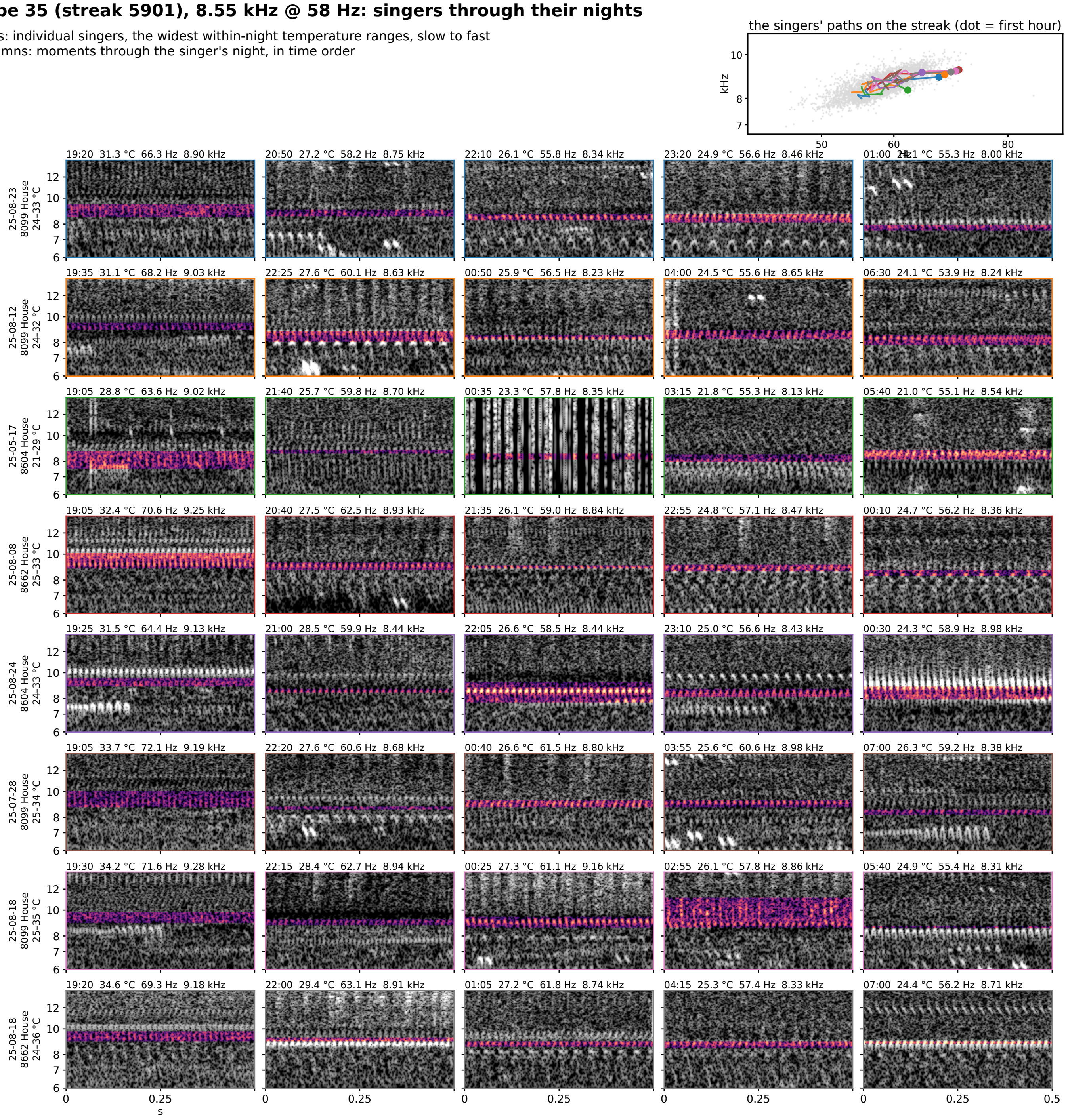


**Figure S8-35.** Type 35 (streak 5901): 8.55 kHz @ 58 Hz at 25 °C, eight singers through their nights. Rows are individual singers of the type, the eight with the widest temperature range within one night (top to bottom by the singer's median rate); columns are moments through that singer's night in time order, 5 windows at even steps through its span. Each cell is 0.5 s of the loudest 30 s of that window on one shared log-frequency axis, the detected band in color (dB above the row's quiet level, 0 to 30) and the rest in grayscale. The cell head gives the local time, the air temperature, the syllable rate and the carrier. Top right: each singer's hourly path on the plane, in the row's color, over the year's points in gray.

### Type 36 (streak 5101), 8.95 kHz @ 21 Hz: singers through their nights

rows: individual singers, the widest within-night temperature ranges, slow to fast
columns: moments through the singer's night, in time order

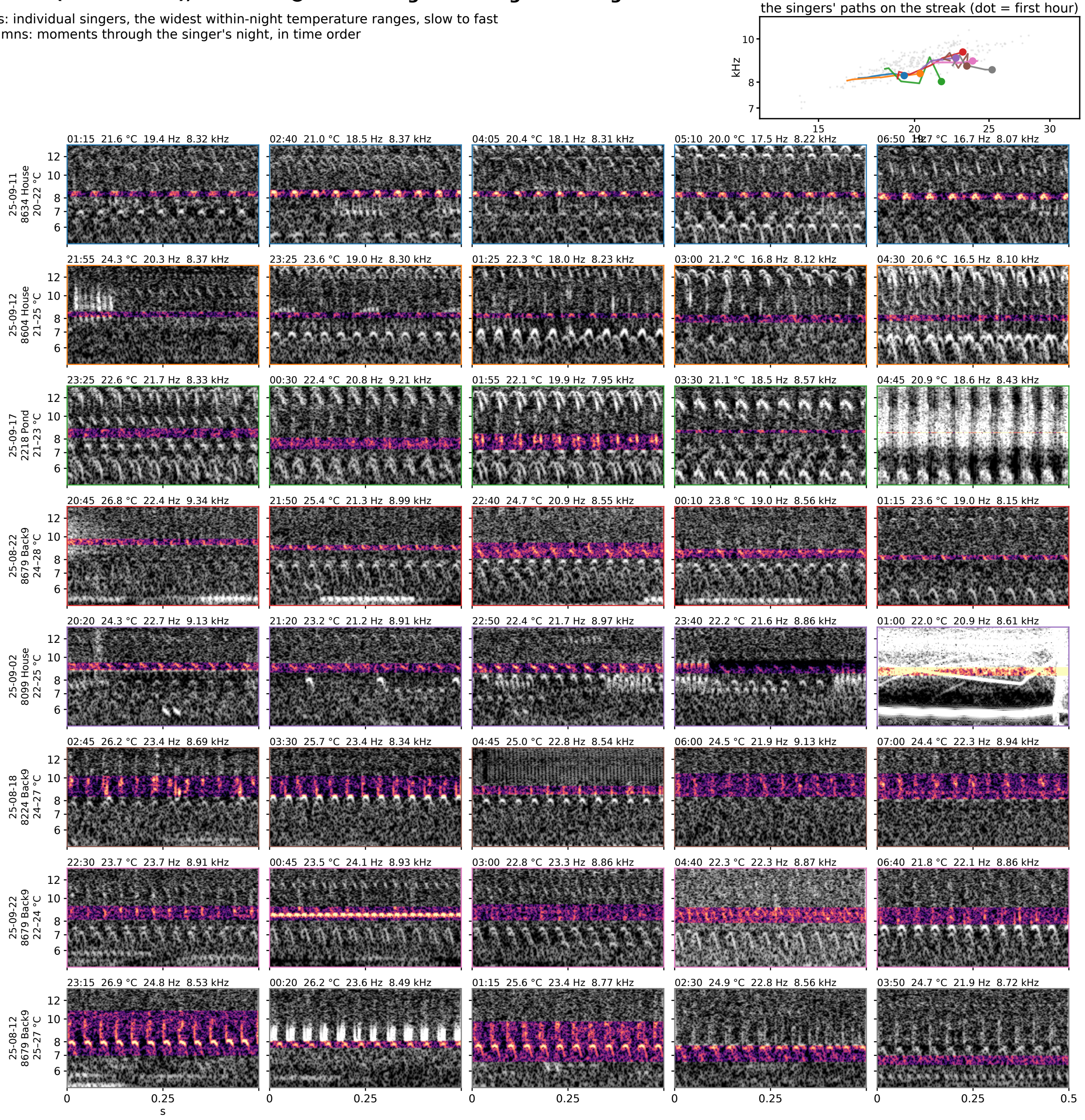


**Figure S8-36.** Type 36 (streak 5101): 8.95 kHz @ 21 Hz at 25 °C, eight singers through their nights. Rows are individual singers of the type, the eight with the widest temperature range within one night (top to bottom by the singer's median rate); columns are moments through that singer's night in time order, 5 windows at even steps through its span. Each cell is 0.5 s of the loudest 30 s of that window on one shared log-frequency axis, the detected band in color (dB above the row's quiet level, 0 to 30) and the rest in grayscale. The cell head gives the local time, the air temperature, the syllable rate and the carrier. Top right: each singer's hourly path on the plane, in the row's color, over the year's points in gray.

### Type 37 (streak 6102), 9.46 kHz @ 58 Hz: singers through their nights

rows: individual singers, the widest within-night temperature ranges, slow to fast
columns: moments through the singer's night, in time order

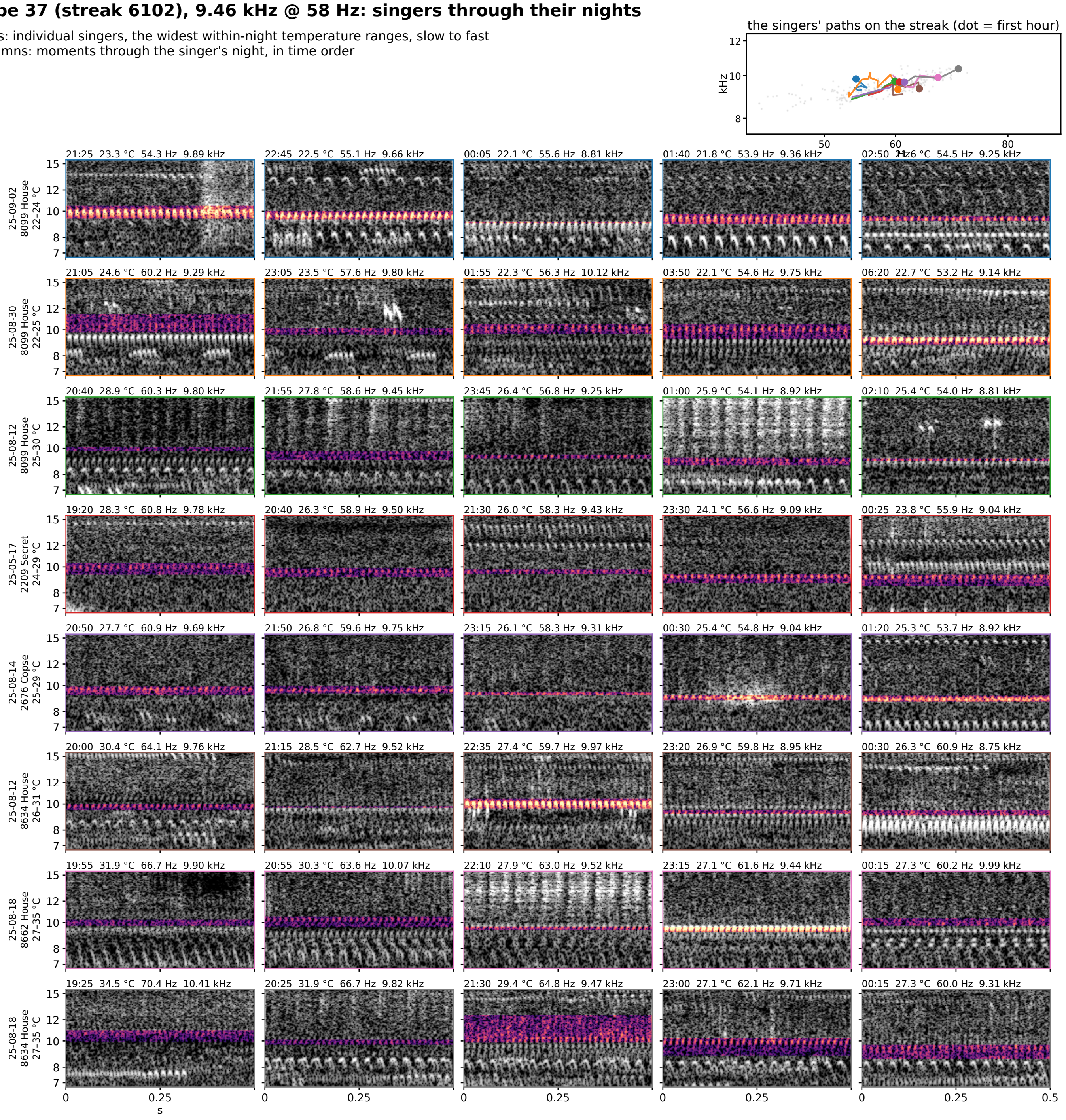


**Figure S8-37.** Type 37 (streak 6102): 9.46 kHz @ 58 Hz at 25 °C, eight singers through their nights. Rows are individual singers of the type, the eight with the widest temperature range within one night (top to bottom by the singer's median rate); columns are moments through that singer's night in time order, 5 windows at even steps through its span. Each cell is 0.5 s of the loudest 30 s of that window on one shared log-frequency axis, the detected band in color (dB above the row's quiet level, 0 to 30) and the rest in grayscale. The cell head gives the local time, the air temperature, the syllable rate and the carrier. Top right: each singer's hourly path on the plane, in the row's color, over the year's points in gray.
